# Evidence for Numerous Embedded Antisense Overlapping Genes in Diverse *E. coli* Strains

**DOI:** 10.1101/2020.11.18.388249

**Authors:** Barbara Zehentner, Zachary Ardern, Michaela Kreitmeier, Siegfried Scherer, Klaus Neuhaus

## Abstract

The genetic code allows six reading frames at a double-stranded DNA locus, and many open reading frames (ORFs) overlap extensively with ORFs of annotated genes (e.g., at least 30 bp or having an embedded ORF). Currently, bacterial genome annotation systematically discards embedded overlapping ORFs of genes (OLGs) due to an assumed information-content constraint, and, consequently, very few OLGs are known. Here we use strand-specific RNAseq and ribosome profiling, detecting about 200 embedded or partially overlapping ORFs of gene candidates in the pathogen *E. coli* O157:H7 EDL933. These are typically short, many of them show clear promoter motifs as determined by Cappable-seq, indistinguishable from those of annotated genes, and are expressed at a low level. We could express most of them as stable proteins, and 49 displayed a potential phenotype. Ribosome profiling analyses in three other *E. coli* strains predicted between 84 and 190 embedded antisense OLGs per strain except in *E. coli* K-12, which is an atypical lab strain. We also found evidence of homology to annotated genes for 100 to 300 OLGs per *E. coli* strain investigated. Based on this evidence we suggest that bacterial OLGs deserve attention with respect to genome annotation and coding complexity of bacterial genomes. Such sequences may constitute an important coding reserve, opening up new research in genetics and evolutionary biology.

## INTRODUCTION

Double-stranded DNA using a triplet code has six possible reading frames. In bacteria, very few extensively overlapping pairs of open reading frames (ORFs) in alternative reading frames are annotated as genes. While differential transcription of regions overlapping to ORFs of annotated genes has been known for quite a while (Figure 1A; data from Landstorfer et al., 2014), implying regulation for at least some of those regions, it remains largely unknown whether such RNAs are also translated. Usually, translation of regions substantially overlapping with ORFs of annotated genes has been dismissed due to several assumptions. For instance, the dictum “one gene - one enzyme” has been associated with “one locus - one gene”, and has typically been assumed for prokaryotes (Meydan et al., 2018). As overlaps place a significant constraint on evolution it is assumed that they should be rare (Johnson and Chisholm, 2004). Consequently, genome-annotation programs discard overlapping ORFs of genes (abbreviated OLGs) above particular length thresholds of the overlapping part (e.g., Delcher et al., 2007). Of note, throughout this manuscript, the OLG is by definition always in antisense to an already annotated gene. However, bacterial genomes contain more long ORFs in alternative frames than expected given codon usage (Mir et al., 2012), and this could be due to selection acting on OLGs. The occurrence of OLGs is accepted in viral genomes (Pavesi et al., 2018) and since there is a constant gene flow between bacteria and their viruses (Kirchberger et al., 2020), shaping bacterial genome evolution (Koskella and Brockhurst, 2014), we propose that long overlaps of ORFs or embedded gene-ORFs, hence, overlapping genes, should also be expected in bacteria (also see Meydan et al., 2018). This hypothesis is supported by several observations: As mentioned, differential transcription of overlapping genome regions (Figure 1A) and translation of so-called non-coding RNAs (Neuhaus et al., 2017) has been reported, also in archaea (Gelsinger et al., 2020). Further, at least a few OLGs in other bacterial species have been described (e.g., Balabanov et al., 2012; Kim et al., 2009).

**Figure 1.**
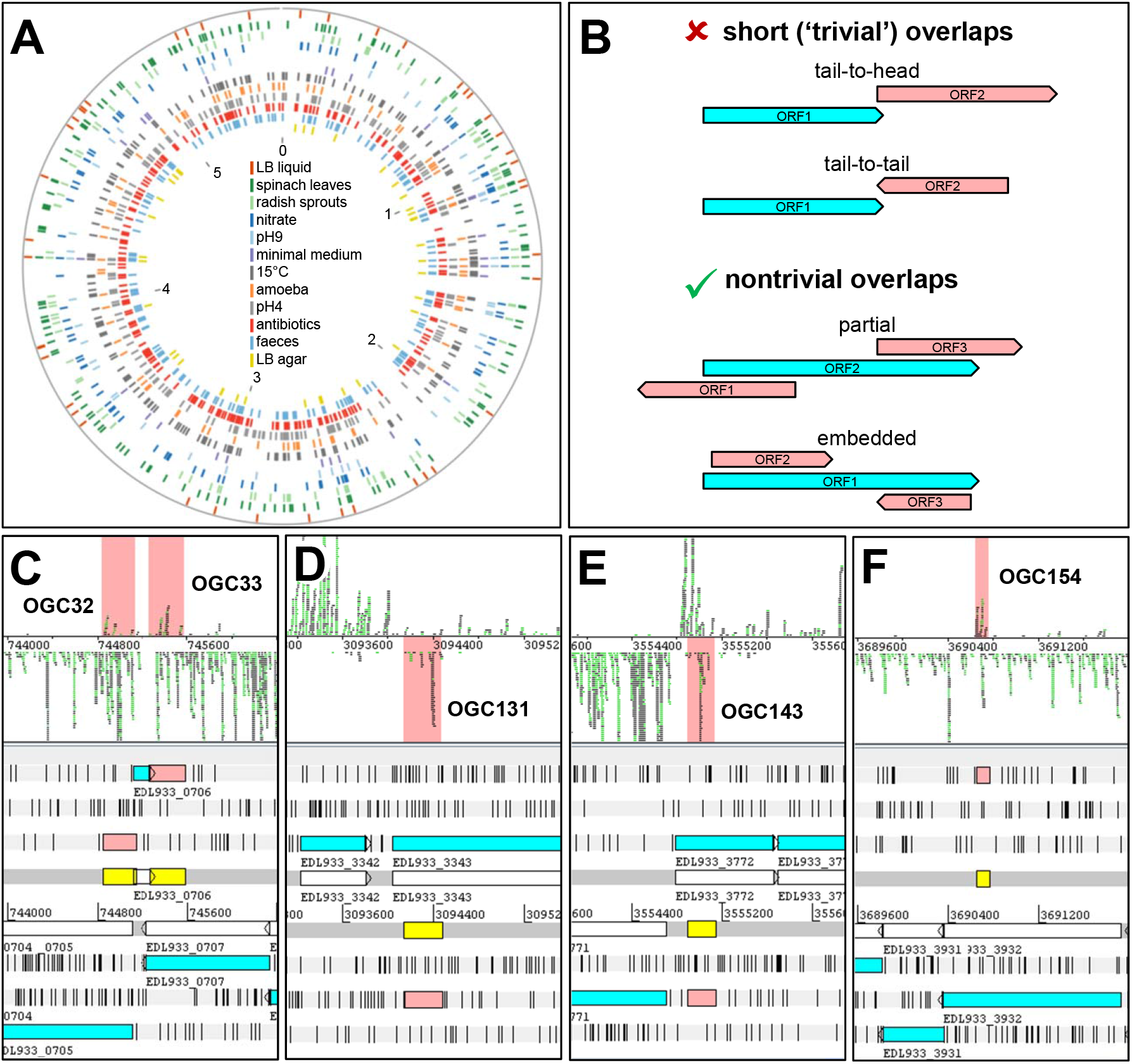
Embedded Overlapping ORFs of *E. coli* EDL933 are Regulated and Also Translated. (A) Differential transcription of 864 embedded ORFs. Colored bars indicate each ORFs’ transcription in the conditions specified. (B) Short and non-trivially OLGs. Gene-ORFs overlap with short regions up to 50 bp mainly (Saha et al., 2015) and most are same-strand allowing translational coupling, but are ‘trivial’ for the encoded proteins’ function. Here, non-trivial partial overlaps and embedded ORFs (100% overlap) sharing at least 93 bp were examined. (C to F) Panels’ upper sections show individual ribosome profiling reads in black (replicate No. 1) and green (No. 2) above (forward) or below (reverse) the DNA (black line). Panels’ lower sections show open reading frames of annotated genes in blue with stop codons (black vertical lines) and the DNA (with the gene-ORFs in white); numbers indicate genomic positions. Reads matching embedded novel ORFs (pink and yellow) are highlighted pink. Exact values for all gene-ORFs are found in Table S6. OGC, overlapping gene candidate. (C) OGC32 and OGC33 overlap to *dacA* (EDL933_0705; D-alanine-carboxypeptidase) and *rlpA*(EDL933_0707; minor lipoprotein). Interestingly, the hypothetical protein EDL933_0706 falls exactly in the gap, presumably avoiding substantial overlaps in the annotation process. (D) OGC131 is embedded in *rtn* (EDL933_3343; EAL-domain bearing Rtn of phage defense). (E) OCG143 overlaps to *napD* (EDL933_3772; NapD nitrate-reductase complex). (F) The very short OGC154 overlaps to a CRISPR-associated Cse1/CasA protein (EDL933_3932).

ORFs of genes overlapping by a few base pairs are recognized as short overlaps. Such short overlaps can be involved, e.g., in expression regulation via translational coupling and have been examined extensively (e.g., Saha et al., 2015). Concerning the encoded proteins, short overlaps are considered ‘trivial’, because very few amino acids (mainly unimportant for protein function) at either end of the proteins are coded by the overlapping region. We excluded short overlaps (‘trivial overlaps’) from the following analysis, focusing instead on overlaps of at least 30 codons. Such substantive overlaps have so far received little attention in bacteria. Non-trivially overlapping ORFs have sufficient overlap for functional domains to be encoded in the overlapping sequence, or are even fully embedded within another ORF. Such overlaps can appear on the same strand or on opposite strands (Figure 1B). Here, we focus on antisense overlaps as they can be detected in ribosome profiling data. The potential widespread functionality of such overlapping ORFs has recently been proposed as a hypothesis to investigate, in light of evidence across diverse bacteria (Ardern et al., 2020). However, only very few cases of such gene pairs, in which their ORFs overlap extensively, have been characterized phenotypically in *E. coli* (Fellner et al., 2014; Fellner et al., 2015; Haycocks and Grainger, 2016; Hücker et al., 2018a; Hücker et al., 2018b; Vanderhaeghen et al., 2018; Zehentner et al., 2020) in strong contrast to more than 128.000 non-overlapping genes known in the *E. coli* pan-genome (Park et al., 2019).

Ribosome profiling detects translated mRNA with unprecedented precision and developing improvements in the method for its use in prokaryotes is an active field of research (Glaub et al., 2020; Ingolia et al., 2019; Wang and Gu, 2018). Using this technique, we previously reported over 400 novel intergenic protein coding ORFs in enterohemorrhagic *E. coli* (EHEC) strains (Hücker et al., 2017; Neuhaus et al., 2016). We also noted that additional translation signals covered the antisense strand of annotated genes (i.e., their ORFs). While the significance of such signals has been disputed in eukaryotes (Ingolia et al., 2014), translational start sites of overlapping protein-encoding ORFs have been reported very recently in bacteria (Weaver et al., 2019). Based on our own ribosome-profiling experiments, we recognized that many signals are specific, reproducible, and regulated (Neuhaus et al., 2016). Consequently, many OLGs could be present in each prokaryote genome. However, do putative antisense reading frames indeed code for proteins? We searched for expressed overlapping ORFs in three pathogenic *E. coli* strains (EHEC O157:H7 strain EDL933 (Riley et al., 1983), EHEC O157:H7 strain Sakai (Michino et al., 1999), and adherent-invasive *E. coli* (AIEC) strain LF82 (Darfeuille-Michaud et al., 1998)) and two lab-strains of *E. coli* (K-12 strain MG1655, Blattner et al., 1997; B-strain BL21(DE3), Daegelen et al., 2009). For a conservative analysis, short overlapping ORFs below 93 nt (i.e., 30 aa) were excluded, despite shorter peptide-coding ORFs existing (Baek et al., 2017; Neuhaus et al., 2017); OLG candidates present in plasmids were also excluded. We present evidence for at least one hundred OLGs in each of the diverse *E. coli* strains studied, with the exception of *E. coli* K12, forming a hitherto unknown large coding reserve.

## RESULTS AND DISCUSSION

### Ribosome Profiling Reveals Translated Overlapping ORF-mRNAs

We initially analyzed strand-specific transcriptomes of EDL933 (Landstorfer et al., 2014; Neuhaus et al., 2016) and found 864 transcribed antisense embedded ORFs. We verified differential expression of those between twelve growth conditions, implying regulation (Figure 1A, Table S4). However, transcriptomics cannot exclude non-coding RNAs (ncRNAs; Neuhaus et al., 2017) and, therefore, the question remained open whether such overlapping-encoded RNAs are translated (Figure 1B). In contrast, ribosome profiling displays the ‘translatome’ (Ingolia et al., 2009), which is the subset of mRNAs actively being translated in the experimental conditions (Ndah et al., 2017). Ribosome profiling data of EDL933 grown in standard LB-medium revealed translation of several overlapping ORFs (for examples, see Figure 1C-F). In total, 216 overlapping open reading frames (subsequently OLG candidates = OGCs) were predicted to be translated in standard LB medium using the ribosome profiling data (Figure S1). Of note, these 216 OGCs have been examined in the following in more detail. To give the reader a better overview, we would like to point to Table S5 for a general overview of all experiments conducted and their results concerning all 216 OGCs. However, the length of the 216 OCGs ranges up to 1497 bp (498 aa) with a mean of ~250 bp (~83 aa). Thus, most ORFs are short compared to annotated gene-ORFs (Figure 2A, Table S6). Interestingly, expression of 137 of the OGCs was detected also in the differential ORF expression data mentioned above (Figure 1A; Table S6). For the 216 OGCs, the number of translational reads per gene-ORF (expressed as RPKM of ribosome profiling) falls within the range observed for annotated gene-ORFs (Figure 2B). The ribosome coverage value (RCV; i.e., normalized ribosome profiling per RNAseq reads) indicates the translational activity of each mRNA (Neuhaus et al., 2017). Annotated genes and overlapping ORFs have a mean RCV of 1.16 and 1.22 respectively (Figure 2C), demonstrating that both groups have a similar translational efficiency.

**Figure 2.**
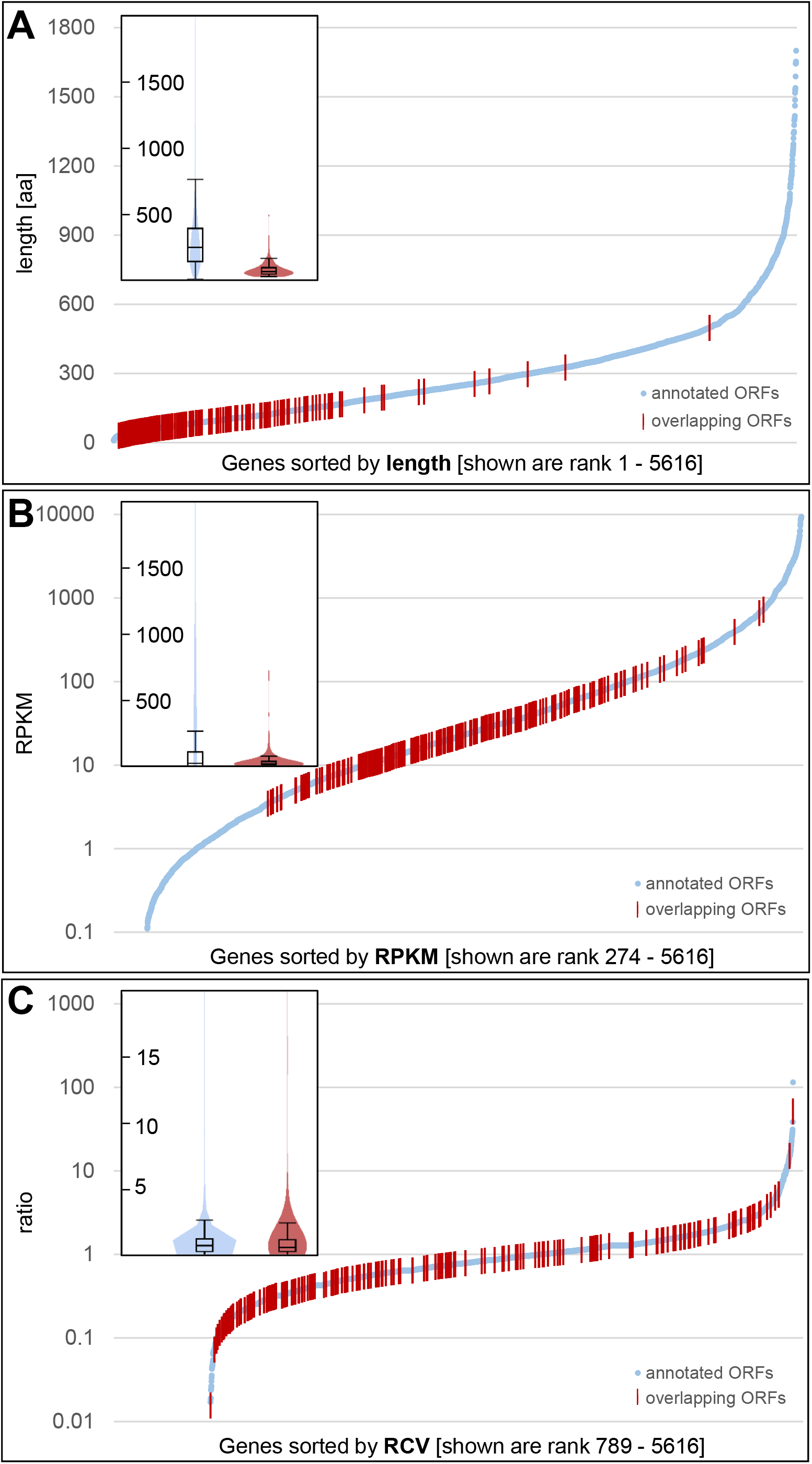
Characteristics of Overlapping Antisense ORFs in OGC Compared to Annotated Gene-ORFs in *E. coli* EDL933. (A to C) Ribosome profiling revealed 216 translated overlapping antisense ORFs. Each panel shows the annotated (blue dots) and antisense overlapping gene-ORFs (red bars) sorted for rank. To complement the overview, each insert represents the same data as its panel but as violin plots in the same color. (A) Ranking by length. The overlapping antisense ORFs tend to be shorter than annotated gene-ORFs. However, a few OGCs reach at least median length. (B) Ranking by RPKM of ribosome profiling. Overlapping ORFs tend to have lower RPKMs compared to annotated gene-ORFs, but some have values above 100. (C) Ranking by the ribosomal coverage value (RCV). The RCV indicates the translational activity of each mRNA (i.e., ribosome coverage). The RCV is comparable between annotated and overlapping gene-ORFs, except for few high values of annotated gene-ORFs.

### Novel Overlapping Genes Can be Expressed and Confer Phenotypes

The 216 OGCs of EDL933 were assessed by Cappable-seq, which allows determination of the transcriptional start (Ettwiller et al., 2016). In total, 148 transcriptional starts within 250 bp upstream of the start codon were detected for 112 OGCs in at least one of eight tested growth conditions. Some +1 sites might be too far upstream and others could not be detected with confidence in our data. An alignment of the promoter regions revealed a clear Pribnow box consensus motif TATAAT (Figure 3A). Functional annotated genes were analyzed in comparison and 2571 genes (57%) had transcription start sites in our data set, displaying an almost identical Pribnow box (Figure 3A). In contrast, the −35 position of both gene sets showed no clear consensus, which is expected for the spacing of the −35 regions is more variable; thus, it will not show up in the analysis used. However, a motif spanning the −2 to +1 region is also very similar between both gene sets (Figure 3A).

**Figure 3.**
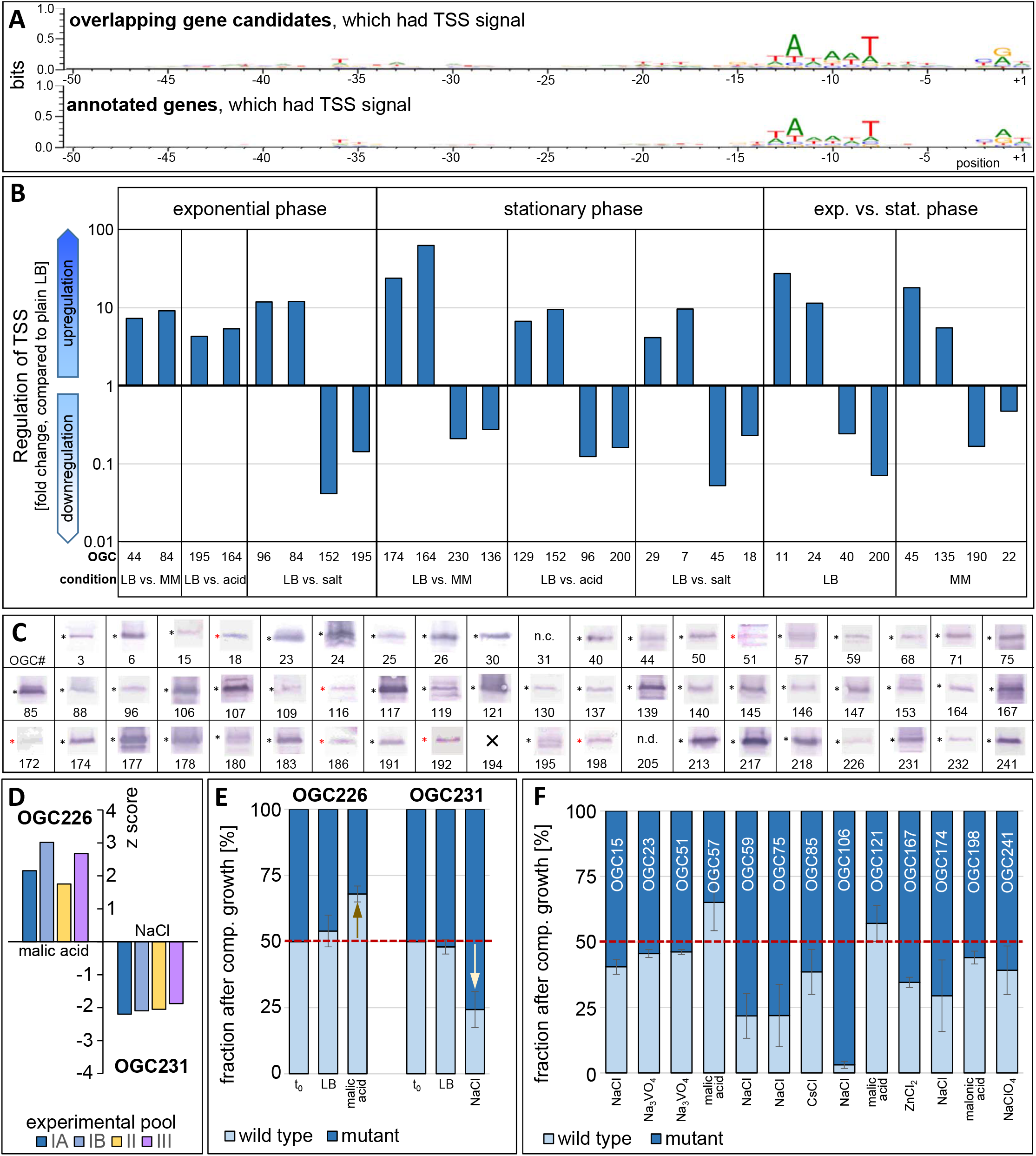
Expression and Phenotypes of Translated Overlapping Antisense ORFs of *E. coli* EDL933. (A) Upstream regions of the transcription start site (TSS =for OGCs (upper panel) and 2571 annotated genes (lower panel) were superimposed according to their +1 (determined by Cappable-seq) and visualized using a sequence logo. For both gene sets, a Pribnow box and a common motif between −2 and +1 appeared. (B) Some significant fold-changes of exemplary TSS for some OGCs in different growth phases and conditions as indicated. Differential up- and downregulation is indicated above or below 1 (i.e., no regulation), respectively. For individual standard deviations, see Figure S2. (C) Western blot snippets for 59 OGCs displaying a phenotype (see text). “n.c.”, not clonable; “×”, not available; “n.d.”, not detectable. Asterisks mark the product of expected size. Red indicates contrast and brightness adjustments for weak products. Complete Western blots of all 210 overlapping gene candidates (OGCs) available are presented in Figure S3. (D) For a global competition experiment, 206 out of 216 OGCs were cloned in an overexpression plasmid. EHEC with plasmids were mixed equally and induced. Each pool (IA, IB, II, and III) was grown in LB (control) and 18 stress conditions (Table S3). z-scores of ±2σ indicate OGCs conferring a phenotype. Exemplary, EHEC expressing OGC226 (left) grew better in malic acid compared to control, while OGC231 (right) grew worse in NaCl. Of 206 OCGs tested, 59 showed a phenotype (Table S7). (E) OGC’s were translationally arrested. EHEC harboring either the wild-type (light blue) or mutant gene (dark blue) were equally mixed (t_0_ and red dashed line) and grown in the respective stress condition. As expected, the wild type of OGC226 grew better in malic acid (left, arrow up), while for OGC231, is grew worse in salt (right, arrow down). In plain LB, the change in ratios was insignificant. Mean values were calculated for four and three biological replicates, respectively. Error bars indicate standard deviations. (F) From 59 OGCs displaying a phenotype (examples in C), 49 were used in a single-competitive assay. After growth (example in D), 15 genes as indicated were found conferring a phenotype in at least one stress condition (indicated), i.e. significant growth difference between wild-type (light blue) and mutant gene (dark blue). Mean values were calculated for three biological replicates, except for OGC59 and OGC241, where eleven and six replicates were used, respectively. Error bars indicate standard deviations. Of note, OGC57 and OGC59, designated *pop* and *asa*, respectively, were studied in detail elsewhere (Vanderhaeghen et al., 2018; Zehentner et al., 2020).

Different TSS signal strength was detected for 74 % of the 148 TSS between either growth conditions (with respect to LB), growth phases (exponential vs. stationary growth) or in both categories (logFC > |1|, FDR ≤0.05, for details see Figure S2). In Figure 3B, the fold change difference of the transcription-start site signal of some OGC examples is shown. Generally, many OGCs are differentially expressed when exponential phase and stationary phase are compared, a phenomenon which is well known for many annotated genes. As expected, only a few OGC’s TSS are influenced by acid or NaCl while more OGC’s TSS are sensitive towards a change of the growth medium (minimal medium MM instead of LB). Fold changes of 5 to 10 are common; in a few cases, even higher differences up to 90-fold were observed. The differential strength of TSS signals depends on growth conditions, which is clear evidence for a specific transcriptional regulation of many of these overlapping genes.

Next, the ability of each OGC to produce a protein was tested by Western blots using SPA-tag fusions. Of 210 clones achieved, 202 produced a protein of the expected size (Figure S3). Western-blot snippets for those 59 OGCs, which also conferred a phenotype (see below) are shown in Figure 3C.

To detect phenotypes, competitive growth was first tested in a pooled overexpression experiment. The coding sequences of 206 out of 216 OGCs were cloned separately on an inducible plasmid and transformed into EHEC. These strains were grown separately initially and pooled in equal cell numbers subsequently to be grown together under various stress conditions, while slightly inducing the OGCs. We expect overexpression of non-functional ORFs to confer a rather minimal effect, while proteins with evidence of function should cause significant differences (Moriya, 2015). In *E. coli*, overexpression of the majority of proteins is mildly to severely deleterious (Kitagawa et al., 2005), an effect which we use to detect a phenotype for the novel gene candidates proposed here (Bhattacharyya et al., 2016). Interestingly, OCG60, OCG95, and OGC223 could be cloned neither for Western blot analysis nor overexpression. This may indicate a toxic phenotype. Three biological (pool I-III) and one technical replicate (pool IB) were conducted. One culture consisted of plain LB with the inducer added. The other batches are again LB with inducer, but plus 1 out of 18 different supplements in each batch (Table S3). After growth, plasmids were isolated and sequenced to determine OGC abundance for each supplement. We deemed OGCs significant that had z-scores of ±2σ in at least two biological pools (Table S7). For example, overexpression of OGC226 led to increased growth in malic acid (positive z-scores), while OGC231 caused diminished growth in salt (negative z-scores; Figure 3D). We found 59 OGCs causing significantly changed growth in at least one stress condition (Table S7), while we would expect up to 22 false positive results for the above thresholds, which is more than twice-as many OGCs displaying a difference in growth between the LB-media with a supplement and plain LB medium.

Next, 49 candidates with a putative growth phenotype in the pooled experiment were subjected to individual competitive growth experiments. Translationally arrested OGC mutants (i.e., a stop codon was introduced in the N-terminal part of the OGC) were compared to the wild type. EHEC harboring plasmids with the arrested OGC or its wild type were mixed 1:1. Using this approach, almost-identical RNAs are expressed, which should not disturb an ncRNA’s function (Bobrovskyy and Vanderpool, 2013), while production of the protein ceases in the mutant. We have tested whether the plasmid copy number changes between a plasmid with the wild-type gene or the mutated gene while growing in competition, showing that this is not the case (Zehentner et al., 2020). The phenotype for the candidate genes was tested in conditions effective in the pooled experiment. In total, 15 mutants showed a significant overexpression phenotype in this type of competition (Figure 3E-F; Table S8). For instance, mutants of OGC226 had lower growth relative to the wild type in malic acid, but mutants of OGC231 had higher relative growth in salt (Figure 3E) corroborating the overexpression phenotypes in the pooled experiment (Figure 3D). Presumably, the number of OGCs detected would be higher if more conditions were tested (e.g., Hillenmeyer et al., 2008).

One interesting example is OGC51, which shows a phenotype in vanadate, an analogue of inorganic phosphate (Rehder, 2020), in the pooled and individual experiments, and has BLASTp hits in *E. coli* and beyond (Table S9). For instance, three codons downstream of the predicted start site is a methionine residue, and a homolog beginning with this residue (WP_000180465.1) is 100% identical to the OLG and is found across *E. coli* and various *Shigella* species. A search of the nr database finds further hits in *Escherichia albertii.* All homologs are labelled as hypothetical proteins. The annotated gene in antisense and its close homologs are also all labelled as hypothetical, with a close homolog also found in *E. albertii.* Comparison to Uniprot protein profiles using HHblits (Remmert et al., 2012) shows further possible homologs for both proteins (Table S9). Such findings add additional evidence for possible functionality at the protein level (for further BLASTp analyses see Table 1, below).

**Table 1.** Embedded antisense overlapping ORFs detected in ribosome profiling data and BLASTp database searches for different *E. coli* strains

| Strain | GenBank | Unique loci – highly expressed in ribosome profiling | BLASTp hits to RefSeq DB |
| --- | --- | --- | --- |
| EDL933 | NZ_CP008957.1 | 84 | 304 |
| Sakai | NC_002695.1 | 190 | 326 |
| K12 | NC_000913.3 | 4 | 125 |
| BL21 | NC_012971.2 | 98 | 89 |
| LF82 | NC_011993.1 | 95 | 87 |

Nevertheless, overexpression phenotypes, especially negative ones, are sometimes disputed, despite it being known that in *E. coli* overexpression for most proteins causes a deleterious phenotype (Kitagawa et al., 2005). According to Moriya (2015), there are four possible mechanisms for an overexpression phenotype: (i) resource overload due to the protein induced, (ii) stoichiometric imbalance, (iii) promiscuous interactions, and (iv) pathway modulations. However, only the first mechanism would be regarded as causing a false phenotype, the other three have biological explanations due to a functional protein product. Our results have shown that only one particular stress condition causes a phenotype, indicating that a general overload does not exist, not even in stress *per se.* Similarly, we only see one or few conditions in which an overexpressed overlapping ORF causes a phenotype in the bulk experiment with 206 different strains in 18 conditions. No phenotype is observed in most of the other conditions, which are - except plain LB - all stressful for the bacteria (i.e. increase doubling time in wild type bacteria).

### Expression and Bioinformatic Evidence for at Least One Hundred OLGs in Typical *E. coli* Strains

In addition to EDL933, we searched for ORFs embedded in antisense to annotated gene-ORFs, the clearest examples of undiscovered OLGs, in four further *E. coli* strains (EHEC Sakai, K-12, BL21, and LF82; Figure 4A). Ribosome profiling data was examined for translation of embedded antisense ORFs, now using even more conservative thresholds of 1 read per million and 60% coverage - thresholds at which only 74% of annotated gene-ORFs are detected, and 57% of annotated gene-ORFs less than 300 nucleotides. The number of expressed gene-ORFs predicted varied with strain and sequencing depth (Table S10). A number of the gene-ORFs were clearly expressed across strains, for instance 43 in distinct loci meet high thresholds of 1 RPM and 60% coverage in at least two strains of the five, and 25 meet a slightly lower threshold of 1 RPM and 30% coverage in at least three strains of the five (Figure 4B). We expect that these numbers would increase with higher read depth samples (particularly for EDL933) and the addition of other environments. Thus, this analysis provides a conservative initial overview warranting further investigation as bioinformatics methods for the investigation of small ORFs, and OLGs specifically (Nelson et al., 2019), continue to improve. Homologs of the set expressed in EDL933 typically had more ribosome profiling reads in the other four strains compared to a length-matched negative control set of homologs of embedded ORFs without footprints in EDL933 (Figure 4C, Table S2). As expected given the strains’ high physiological and genomic similarity, homologs of annotated Sakai genes show the highest expression. Expression in the two laboratory strains, K-12 and BL21, was lower than in the much less closely related pathogenic strain LF82.

**Figure 4.**
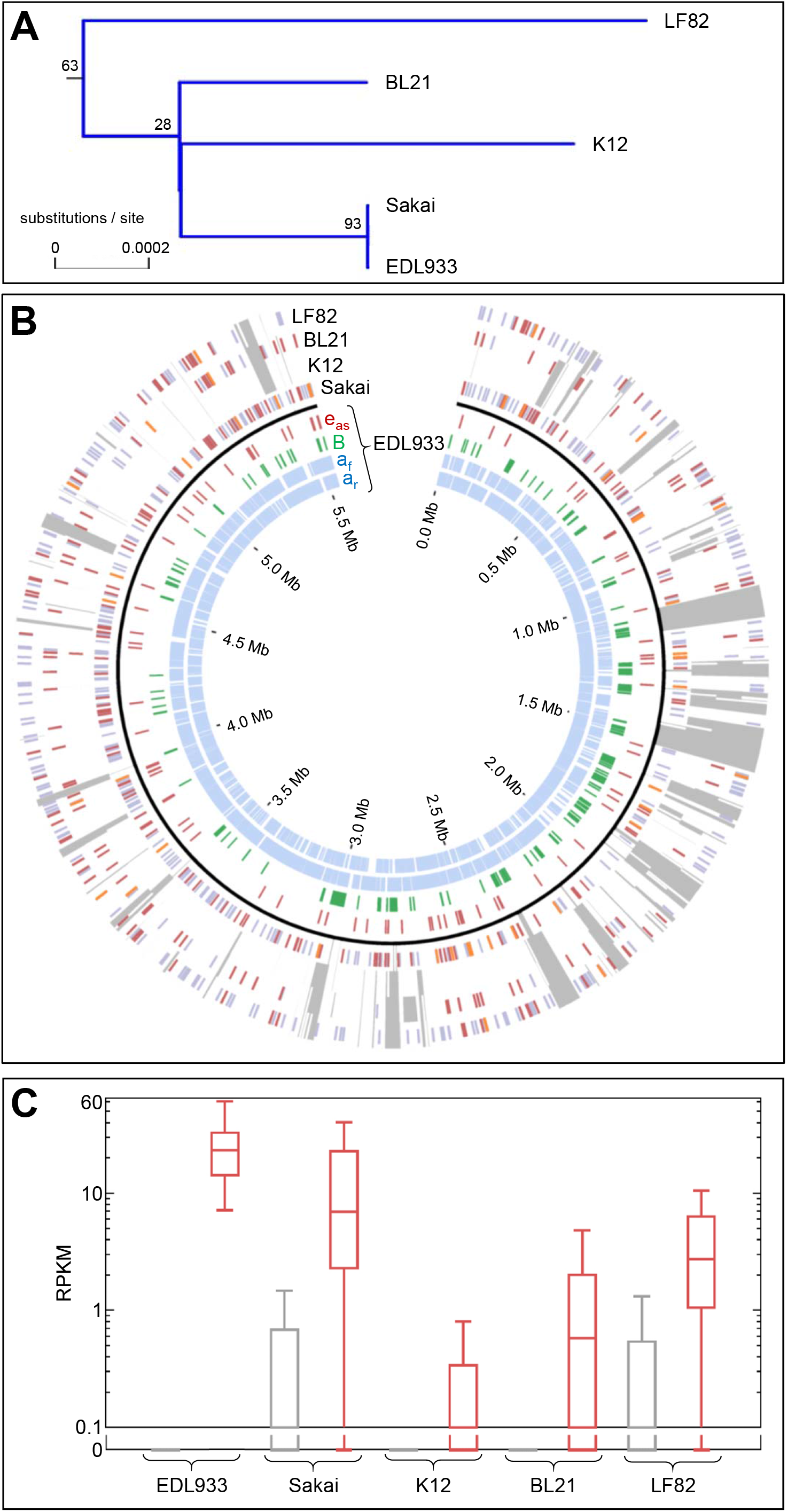
Comparison of Embedded ORFs for Different *E. coli.* (A) Taxonomic tree of the *E. coli* strains used in this study. *E. fergusonii* was used as outgroup (not shown). K-12 and BL21 are common lab strains, while EDL933 and Sakai are closely related enterohemorrhagic *E. coli* (EHEC) of the serogroup O157:H7. Strain LF82 is a pathogenic AIEC. (B) Circos plot showing expressed antisense embedded ORFs in four *Escherichia coli* strains, in ORFs homologous to antisense embedded ORFs in EDL933. In each strain, ORFs are shown which either have at least 1 read per million mapped reads (RPM) and 30% coverage (green), at least 1 RPM and 60% coverage (blue) or at least 1 RPM, 60% coverage, and also homologous to an ORF expressed above these thresholds in EDL933 (orange). (C) Comparing expression (i.e. coverage with ribosome footprints) for two groups of ORFs in the different strains. For strain EDL933, the first group of ORFs has zero reads (grey box plot, i.e., a line since zero); the second group are antisense embedded ORFs, which overlap to the ORF of annotated genes (red box plot) and showing expression (≥ 1 RPK, ≥ 60% coverage). The box plots for the following strains (as indicated) show the data for the homologous ORFs (if present) for each group matching to the ORFs of EDL933. In all cases, the ORFs that showed expression in strain EDL933 and have a homolog in the other strain show also higher expression compared to the ORFs which were zero in EDL933. This indicates that the expression patterns are non-random, i.e., certain frames are expressed across strains.

A common objection to these findings is that many regulatory antisense RNAs exist (Brophy and Voigt, 2016) and the detected novel RNA species are just regulators of the opposing gene but not mRNAs of overlapping genes. While regulatory effects caused by antisense RNA and converging transcription could also play a role here, transcription of antisense RNAs generally is widespread in bacteria (Georg and Hess, 2018). In contrast, only a minority of antisense RNAs produce a pattern of ribosome footprints, which is, in principle, not distinguishable from the pattern formed for annotated genes (Figure 1C-F, Figure S1). The majority of truly ncRNAs are not detected in ribosome profiling (Neuhaus et al., 2017) and additional methods have verified ribosome-profiling results. In a study in *Salmonella*,27 novel ORFs were tested using a translational tag and 25 were confirmed to be expressed (Baek et al., 2017). Similarly, in a study using *E. coli* K-12, 41 small ORFs were tested using chromosomal tagging and 38 were confirmed (Weaver et al., 2019). Both studies suggest a false-positive detection rate of about 10%. Therefore, approximately 90% of the ORFs indicated by ribosomal profiling to be translated should correspond to a true translational signal. Further indications of genuine translation can come from for instance i) differential and specific regulation, ii) detection of a reading frame and iii) retapamulin-assisted ribosome profiling. Regulation indicates functionality. It was found that the set of embedded antisense overlapping gene-ORFs expressed differs according to the culture conditions, assessed with data from media conditions of LB, BHI, and BHI plus cold and salt stress (Figure 5A) for strain Sakai and for aerobic versus anaerobic conditions in Schaedler broth for LF82 (Figure 5B). Most overlapping gene ORFs for Sakai were expressed in the full medium BHI, and for anaerobic conditions in LF82, but each condition was associated with unique genes as well, strongly suggesting that more overlapping gene-ORFs remain to be discovered in other conditions.

Translating mRNA would furthermore be indicated by the codon-wise progression of the ribosome, e.g., the coordinates for the ribosome footprints mapping to a translated ORF differ by multiple of three nucleotides. Re-analyzing this data, we find that, for example, the BL21 data without retapamulin treatment maps with a strong bias associated with the reading frame of annotated gene-ORFs (although not for all gene-ORFs) - we show this for short gene-ORFs of 300 nucleotides or less in Figure 5C. This figure shows that some read lengths (in this sample, those with a length divisible by three) are more informative than others concerning reading frame - a property which could be used for more accurate overlapping gene-ORF detection in future. Such a reading frame is also detectable in the ORF of many overlapping gene candidates. To illustrate that many candidates show a clear reading frame, in Figure 5C we show the ORF with the best ‘periodicity’ or reading frame bias from each antisense region with high coverage (1 RPM, 60% coverage). Further, within all of the OLG candidates meeting these thresholds, antisense embedded ORFs with many reads have a greater average periodicity than those with few reads.

**Figure 5.**
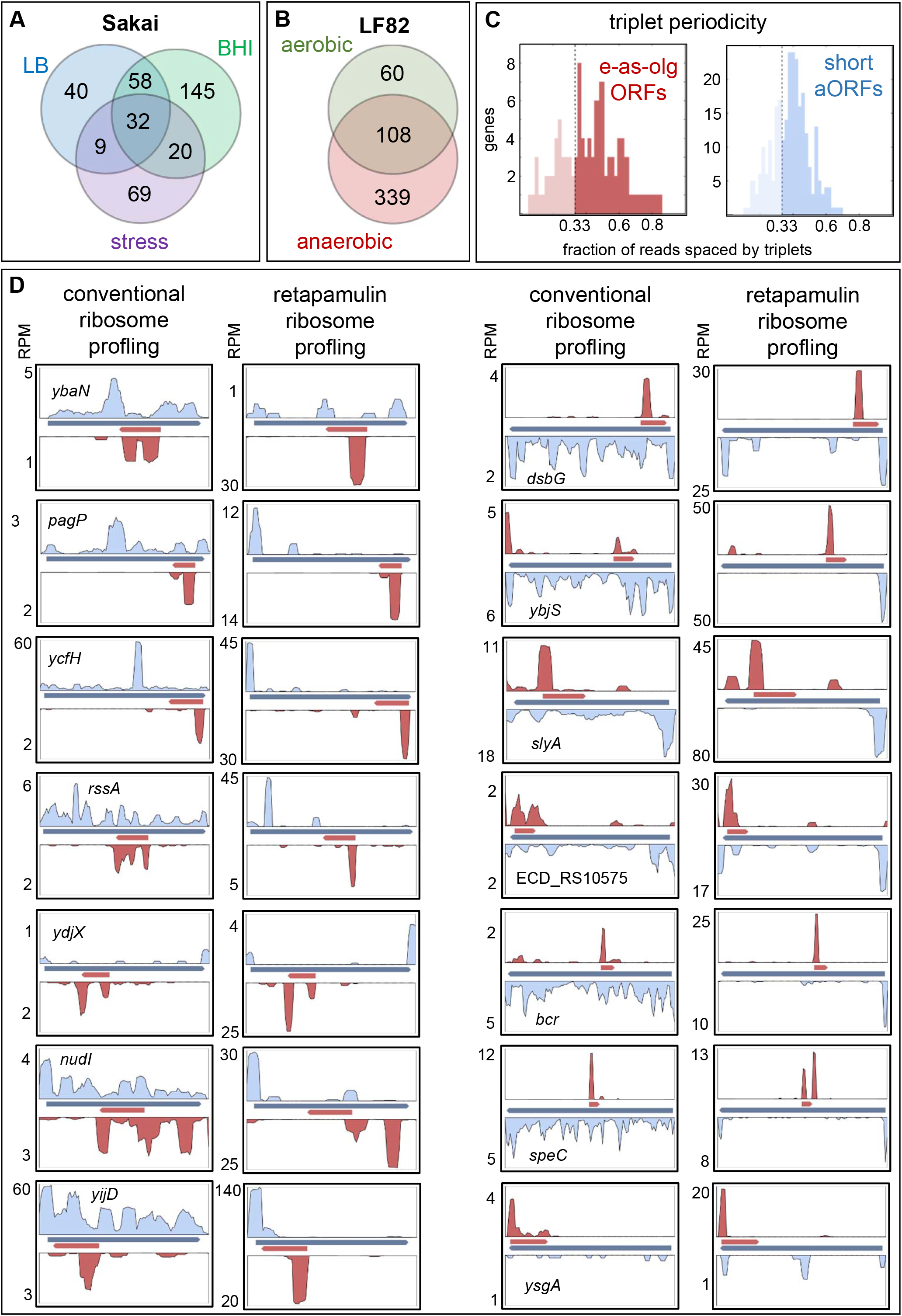
Further Analysis of Ribosome Profiling Data in Sakai, LF82, and BL21. (A) Differential expression of antisense embedded ORFs in EHEC Sakai across three media conditions. LB, Luria-Bertani broth; BHI, brain heart infusion; stress, 14°C with 4% NaCl in BHI. (B) Differential expression of antisense embedded ORFs in AIEC LF82 across two media conditions - aerobic and anaerobic growth in Schaedler broth. (C) Reading frames in ribosome profiling of strain BL21, for embedded antisense overlapping ORFs (e-as-olg ORFs) and annotated gene-ORFs (aORFs); the latter were length matched. For embedded antisense overlapping ORFs, the best candidate from each region of Table 1 is used. (D) Antisense embedded ORFs of strain BL21 found to be expressed either in conventional ribosome profiling (left panels), compared to data from retapamulin-assisted ribosome profiling (right panels). Retapamulin stalls the ribosomes specifically on the start codon, which increases the detectable signal (i.e., number of reads above the start codon). Left two columns, examples for antisense embedded ORFs in the revers DNA strand; right two columns, for the forward DNA strand.

Finally, retapamulin stalls the ribosome mainly at the start codon of mRNAs, such that the resulting bias in read depth is associated with translation initiation. Indeed, a few of the candidates reported here have been found in a previous study using retapamulin-stalled ribosome profiling (Table S11; Meydan et al., 2019). For these experiments, strain BL21 was used. We find 18 antisense overlapping ORFs with a high read depth of at least 2 RPM in both normal ribosome profiling data and 2 RPM in the putative start region (start of translation plus 30 codons) in retapamulin-stalled ribosome profiling data, and show 14 of these in Figure 5D, illustrating both the expected peak accentuation in the retapamulin treated sample and the simultaneous expression of genes with overlapping ORFs on both strands.

We searched for high sequence similarity with annotated proteins in the RefSeq bacterial database as detected with Diamond BLASTp (Table S12). All antisense embedded ORFs (≥ 93 bp) in the five *E. coli* strains were used as queries. Each *E. coli* strain contains roughly 30,000 embedded ORFs antisense to annotated gene-ORFs. In the BLASTp analysis, 87 to 326 embedded ORFs in each strain each received at least two hits to proteins in RefSeq, i.e., each of these queries had at least two subject matches (Table 1). The length distribution of these BLASTp hits is shown in Figure S4, the median length being 234 nt. High similarity (i.e., e-value ≤ 10^-10^) to a protein-coding gene in another organism is good evidence of a gene overlooked or discarded in annotation (e.g., Tatusova et al., 2016). Please note that we do not refer to any evolutionary relationship or conservation of the protein-matches found which is difficult to assess because of the overlapping nature, but only to the fact that a particular protein (giving rise to the match) had been annotated in another bacterium, which adds evidence for an inherent protein-coding ability of the ORF in question.

### Mis-annotation of the ‘Mother Genes’ of Overlapping Genes?

One may question the annotations of the ‘mother gene’ (i.e., by definition the annotated gene containing an ORF overlapping another expressed ORF, thus forming an overlapping gene pair), especially when this mother gene is a hypothetical gene. In response to this potential objection however, the utmost majority of the so-called hypothetical proteins have been detected by MS (e.g., Venter et al., 2011) and loci of embedded ORFs detected in *E. coli* by ribosome profiling typically show reads mapping to both DNA strands at the same time (Figure 1C-F, Figure 5D, Figure S1). Interestingly, a major proteogenomic study of 46 prokaryotes reported 245 “conflicts”, defined as overlapping ORF pairs in which both proteins encoded have strong signals in MS data. The naming “conflicts” exhibits how scientists have perceived OLGs (Venter et al., 2011); however, it is good and necessary that such ‘unfitting’ data are reported. Further examples of studies showing evidence for antisense genes with overlapping ORFs in prokaryotes have been recently reviewed (Ardern et al., 2020). In addition, we argue that the recognition of overlapping ORFs using recent annotation pipelines strongly indicates protein coding. However, OLGs are arbitrarily removed by these programs based on assumptions detailed above. For instance, in order to circumvent the automated overlapping gene-ORF removal, we masked the start codon of the annotated-ORF contained in the mother gene, i.e. *ompA*, in another study by changing the ATG to an NNN. This immediately resulted in the annotation of the overlapping gene-ORF candidate in question, i.e. *pop* (Zehentner et al., 2020). Furthermore, many overlapping-encoded proteins have a BLASTp hit in the RefSeq database (Table 1), which is a curated database and, thus, should be reported. There is therefore reason to suspect that many more OLGs await discovery and that we may enter a new paradigm regarding the coding capacity of the prokaryotic genome.

### Why Have Overlapping Genes Been Missed?

To date, only very few protein-coding embedded ORFs across all bacteria have been noted, and even fewer have been verified phenotypically. Why? Firstly, small and weakly expressed proteins are extremely difficult to detect for technical reasons (Pauli et al., 2015; Storz et al., 2014). Perhaps more importantly, OLGs are not permitted by annotation programs or GenBank (e.g., Delcher et al., 2007; NCBI, 2018). This appears to be based on the implicit assumption that their evolution and conservation are improbable due to an additional evolutionary constraint (Johnson and Chisholm, 2004), and a diminished optimality of adaptation processes (Rogozin et al., 2002), but early calculations that OLGs are possible so long as the sequence is not too highly specified (Yockey, 1979) should also be taken into account.

In addition to the above, overlapping protein-coding genes within phages have long been known (Sanger et al., 1977), and since bacteriophage contribute greatly to the evolution of bacterial genomes this raises again the question of why OLGs have been dismissed in bacteria. A common argument for viral genomes’ unique status in comparison to bacteria is an assumed selection pressure towards reduced genome sizes in viruses (Chirico et al., 2010). However, this assumption has been shown to have weak support, as the genome can retain function if OLG pairs are decompressed (Jaschke et al., 2012) and the capsid space is often not fully utilized (Brandes and Linial, 2016). However, it is unknown whether some overlapping viral genes may be repressed during the prophage state.

Another aspect of why long overlaps between coding ORFs might have been missed may be illustrated by the extremely low number of expressed OLG candidates in *E. coli* K-12 compared to any other *E. coli* examined here. In *E. coli* K-12, we only found 4 OLG loci expressed in ribosome profiling (Table 1) (a number which did not dramatically change dependent on read depth thresholds), while the other strains had at least 84 and up to 190 such loci. We compared our results with the OLGs reported in Meydan et al. (2019) and Weaver et al. (2019). Meydan et al. (2019) do not report any embedded antisense overlapping gene-ORFs at least 30 codons long based on their retapamulin ribosome profiling data in BL21 and K12. Weaver et al. (2019), using the ribosome stalling peptide Onc112, report three antisense embedded OLGs in the K-12 strain BW25113, of which one matches one of our predictions, namely co-ordinates NC_000913.3:3797772-3798014 on the minus strand. The other two were excluded from our analysis as one overlapped a pseudogene and the other was in close proximity to an annotated gene-ORF on the same strand. Furthermore, Meydan et al. (2019) also found 8 overlapping gene-ORFs at least 30 codons long, either same-strand or partially overlapping in their ORFs, which have a clear start site in both BL21 and the K-12 strain BW25113 (listed in their Table S7). They report many more shorter OLGs as well. Similar to our results of extremely reduced expression in K-12, they find with their retapamulin method and a different substrain (BW25113) that the K-12 derivate expresses less than one quarter of unique genes with overlapping ORFs of all lengths as the other strain does. Therefore, while their data confirm that some OLGs are expressed across divergent strains it is clear that these eight are only a very small subset of all expressed OLGs in the species of *E. coli.* Thus, some OLGs exist in K-12 but clearly fewer than in any of the other strains. Indeed, the first observations of multiple OLGs in bacterial genomes were made in *Pseudomonas fluorescens* and *Chlamydia trachomatis* (Jensen et al., 2006; Silby et al., 2004) rather than in *E. coli* K-12, despite this strain probably being the best-examined bacterium of all. We hypothesize that strain K-12 is rather unusual with respect to OLGs. This may be related to K-12 being a lab strain since 1922 and has been through many serial passages (Bachmann, 1972). The B-strain BL21 has also been a lab strain since 1918 (Daegelen et al., 2009), but may have undergone fewer or less dramatic changes compared to K-12-their differences in this regard will be worth investigating in future as we learn more about OLGs.

### Do Further Overlapping Genes Await Discovery?

Assessing a phenotype of an unknown gene is by no means an easy task, even more so if it is a gene with an overlapping, embedded ORF, where interfering with the ORF will also affect the mother gene. Therefore, it is not surprising that only very few bacterial OLGs have been verified by demonstrating a phenotype so far. Examples are, in EDL933, the pairs *mbiA/yaaW, nog1/citC, asa*/EDL933_1238, and *pop/ompA* and in Sakai, the pairs *laoB*/ECs5115 and *ano*/ECs2385, (Fellner et al., 2014; Fellner et al., 2015; Hücker et al., 2018a; Hücker et al., 2018b; Vanderhaeghen et al., 2018; Zehentner et al., 2020). Thus, in the best case, roughly 1 OLG-ORF per 1000 annotated *E. coli* gene-ORFs is currently known. We suggest that beside those OLGs we detected here, numerous further OLGs await discovery. For instance, in *E. coli* and other bacteria, improved ribosome-profiling protocols which predict translation initiation sites detect both sense and antisense oriented OLG pairs among these (Meydan et al., 2019; Smith et al., 2019; Weaver et al., 2019).

In addition, mass spectrometry (MS) has detected OLGs as well, for both antisense and same strand overlaps. For instance, Kim et al. (2009) reported a few antisense OLGs in *Pseudomonas.* In *E. coli* K-12, the antisense OLG *yghX* overlapping *modA* was found by MS (Kurata et al., 2013), and a samestrand OLG, *gndA*, has been found embedded in *gnd* (Yuan et al., 2018). However, neither of these has been phenotypically assessed. Using N-terminal proteomics, Ndah et al. (2017) detected six proteins of at least 30 amino acids which were fully embedded on the same-strand as well as eight embedded antisense overlaps in MG1655 (Figure S5). Finally, plasmids and bacteriophage were excluded in this study, but contain OLGs as well, and many conditions have yet to be tested and are likely to show expression of additional overlapping gene-ORFs (e.g., see Figure 5A & C).

### Do Overlapping ORFs Form a Hidden Coding Reserve?

Hundreds of overlapping ORFs show evidence of being actively translated in the limited set of ribosome profiles available and analyzed for *E. coli* strains so far (also compare to File S1). We collated evidence for a few hundred fully embedded OLGs in five *E. coli* genomes, with a particular focus on the pathogenic EHEC strains EDL933 and Sakai (e.g., Table 1). Therefore, we suggest the number of genes with overlapping ORFs is at least 25 per 1000 annotated genes and probably even higher. There are a number of lines of evidence supporting this claim. There are plausibly many more OLGs which are not strongly expressed under the few conditions examined here, are not completely overlapping their antisense gene-ORF, are overlapping another gene-ORF the same strand, or are shorter than 30 amino acids. Additionally, it has recently been found that many arbitrarily chosen ORFs, including overlapping ones, can be expressed as proteins (VanOrsdel et al., 2018). Computational analysis has recently suggested that many protein domains are able to encode overlapping sequences with high similarity to other protein domains (Opuu et al., 2017), suggesting that the informational constraint is much lower than often assumed.

Further, the bacterial proteins produced by OLGs are proposed to be functional since regulated protein coding elements would confer an energy cost and population sizes in *E. coli* are sufficient to ensure that they would soon be removed by selection in the absence of any function (Bobay and Ochman, 2018; Kirchberger et al., 2020; Lynch and Marinov, 2015). There are many opportunities for further investigation. For instance, we suggest that unsolved questions of where novel genes come from (2012) might in the case of bacteria be partly explained via overprinting, as has been suggested decades ago, although for theoretical reasons only (Grassé, 1977).

More generally we propose that non-trivially OLGs constitute a large hidden coding reserve in bacteria, indicating another, unforeseen level of coding complexity in typical *E. coli* genomes in addition to others recently discovered (Grainger, 2016). Therefore, we propose that studying bacterial overlapping ORFs has the potential to open up interesting fields of research spanning genetics, protein sciences and evolutionary biology.

## EXPERIMENTAL PROCEDURES

### Transcriptomes of Strain EDL933

Strand specific transcriptomes from the study of Landstorfer et al. (2014) were analyzed according to the bioinformatics methods used for the “Ribosome profiling analysis” section below, aside from the use of cushaw3 (Liu et al., 2014) for aligning colorspace format input with default settings, and a minimum sequence quality score of 10 for cutadapt trimming. The eleven conditions assessed were: 10% LB medium, M9 minimal medium, 10% LB at pH9, 10% LB with 0.2 mg/ml nitrite at pH6, spinach leaf juice, radish sprouts, 10% LB at 15°C, 10% LB at pH4, 10% LB with antibiotic (2 μg/ml sulfamethoxazol and 0.4 μg/ml trimethoprim), pasteurized cow dung, and 100% LB-medium agar plate. In addition, EDL933 was co-cultured with amoeba *(Acanthamoeba castellani);* Neuhaus et al. (2016). Cells were harvested in the transition from late exponential to early stationary phase in all conditions. Embedded antisense ORFs expressed in a sample with at least 5 reads, at least 10 RPKM (reads per kilobase ORF in a gene per million sequenced reads) and at least 50% coverage after removal of reads mapping to rRNA or tRNA are shown in Figure 1A. Where ORFs meeting these criteria overlapped within a particular sample, the ORF with the highest product of RPKM and coverage was picked to obtain a single representative ORF per region.

### Ribosome Profiling of Strain EDL933

Ribosome profiling data for *E. coli* O157:H7 EDL933 was taken from Neuhaus et al. (2017). Cells were grown in ten-fold diluted lysogeny broth (LB; 10 g/L peptone, 5 g/L yeast extract, 10 g/L NaCl). At the transition to early stationary phase, 170 μg/ml chloramphenicol was added. Ribosomal footprints were prepared following Ingolia (2010), but using RNase I. After clean-up and sequencing on a Illumina MiSeq, footprints were mapped to the reference genome existing at that time (NC_002655) from Perna et al. (2001) using Bowtie2 (Langmead and Salzberg, 2012). From the same samples, strand-specific mRNA sequencings were conducted. Note, only after our biological experiments (i.e., detection of OLG candidates, cloning, competitive growth) had begun, a new reference genome was published by Latif et al. (2014). The new reference genome was used in the subsequent re-analysis (see below). Evaluation of the ribosome profiling data was conducted as follows: The read count value above 20 counts / kbp was used as a first threshold for translation, which is above the 63^rd^ percentile. After this threshold-based search for significant antisense translation of embedded ORFs, false positives were removed by visual inspection. All embedded ORFs were inspected manually and likely false positives were excluded (e.g., in cases of uneven coverage or no clear gap to a neighboring annotated (a) ORF). For some embedded ORFs, the originally predicted start codon position was moved further downstream according to the ribosome profiling data. Combining ribosome profiling with RNAseq allows estimation of the translatability of an ORF, expressed by the ribosomal coverage value (RCV), which is the ratio of the RPKM value for the translatome over the RPKM value for the transcriptome for a given gene-ORF. The RCV can be used to distinguish RNA not currently translated from RNA in the process of translation (Hücker et al., 2017).

### Bioinformatics for the Re-analysis of *E. coli* EDL933 and Four Additional Strains

#### Ribosome Profiling Analysis

Ribosome profiling data were taken from the following studies: Data for *E. coli* strain Sakai (grown in LB) were taken from Hücker et al. (2017). Data for *E. coli* K-12 (grown in LB) were taken from Woolstenhulme et al. (2015), for *E. coli* strain BL21(DE3) grown in LB (abbreviated as BL21) from Meydan et al. (2019), for *E. coli* strain LF82, grown in Schaedler broth as described in Zehentner et al. (2020). For the latter, data was deposited in the NCBI Sequence Read Archive (accession number PRJNA654277). All other data were downloaded from the NCBI Sequence Read Archive. Information on strains, genomes and ribosome profiling data are in Table S1.

Fastq files were trimmed using Cutadapt (Martin, 2011) with a minimum sequence quality of 30 and adapter sequences predicted using DNApi (Tsuji and Weng, 2016). The output of cutadapt was then aligned against the appropriate genome using Bowtie2, with the “local” alignment option, a seed length of 19 (L 19) and zero mismatches (N 0). Reads overlapping regions of the genome corresponding to tRNA and rRNA were removed from the BAM files. Using only reads with MAPQ of at least 30, RPKM and coverage values were calculated for all embedded antisense ORFs at least 30 codons long using BEDTools intersect (Quinlan and Hall, 2010) after excluding ORFs within 100 bp of an annotated gene-ORF on the same strand, to avoid false positive inferences. ORF co-ordinates were calculated with an “ORFFinder” perl script written by Dr. Christopher Huptas. For strains EDL933, Sakai, and K-12 the two biological replicates available in LB from the studies cited above were merged. For strains BL21 and LF82, just one sample was available, but read depth was high (over 15 million and nearly 8 million mapped reads respectively, outside of tRNA and rRNA loci). All samples had at least 4 million mapped reads; K-12 had over 41 million. After the quality filtering described, ORFs meeting minimum thresholds of 1 read per million (RPM) and coverage of 60% were taken as putative OLG candidates in each strain, and regions with multiple overlapping candidates were determined with the BEDTools tools coverage, intersect, and merge. The number of distinct regions is reported in Table 1, and all candidates with their expression values are listed in Table S2.

#### BLAST

All ORFs fully embedded within and antisense to annotated gene-ORFs in the main chromosomes were used as queries in a Diamond (BLASTp option) protein search (Altschul et al., 1990; Buchfink et al., 2014) against bacterial proteins from the RefSeq database (version downloaded 29 June 2020 - files at https://ftp.ncbi.nlm.nih.gov/refseq/release/bacteria/*gz), and hits with e-value less than 10^-10^ were retained. Only those ORFs with more than one match in the RefSeq database are reported in Table 1.

#### Comparisons between Five Strains

A multi-locus tree of the five *E. coli* strains was inferred using loci extracted, aligned, and concatenated with the Genome Taxonomy Database Toolkit (GTDBTK) (Chaumeil et al., 2020; Parks et al., 2018). The genome of *Escherichia fergusonii* (GCF_000026225.1_ASM2622v1) was included as the outgroup for subsequent re-rooting, conducted with Newick Utilities (Junier and Zdobnov, 2010). A maximum likelihood phylogenetic tree was calculated from the multifasta protein alignment using IQTREE2 (Minh et al., 2020; Nguyen et al., 2015) with default settings, and with the best fitting protein evolution model inferred using the Bayesian information criterion (BIC) calculated by the IQTREE implementation of ModelFinder (Kalyaanamoorthy et al., 2017). The model chosen was “mtMet+F”, i.e. the mitochondrial metazoan exchange matrix with empirical amino acid frequencies, with a BIC of 28821.173. The bootstrap values of 1000 bootstraps were calculated with UFBoot (Minh et al., 2013).

Single copy orthologues of fully embedded overlapping open reading frames in EDL933 and each of the four comparison strains were inferred with “Orthofinder” (Emms and Kelly, 2015, 2019), in a pairwise manner comparing each strain to EDL933. Homologous ORFs are shown with a white background in the Circos (Krzywinski et al., 2009) tracks. Those expressed according to the different thresholds reported (1 RPM and 30% coverage, or 1 RPM and 60% coverage) were determined using BEDTools coverage.

#### Further Analysis of *E. coli* Sakai, LF82, and BL21 Ribosome Profiling Data

ORFs expressed with at least 1 read per million quality-filtered mapped reads and 60% coverage in ribosome profiling data for strain Sakai, derived from experiments in three different media conditions, were compared. Likewise, ORFs expressed above these thresholds in strain LF82 were compared for two conditions - aerobic and anaerobic in Schaedler broth media. The tendency for reads of different lengths to map at different codon positions in known gene-ORFs, indicating a reading frame, was also investigated using BEDTools intersect. It was observed that the BL21 data had the clearest reading frame. Due to difficulty in accurately predicting the translated ORF in any given region showing high expression, the sum of the reads for the ORFs with the best reading frame in each region is shown. Further, embedded antisense open reading frames in strain BL21, which had high expression of more than 2 RPM in both the normal ribosome profiling and retapamulin-treated sample within 30 nucleotides of the putative start codon were examined, with some of the clearest examples of expression displayed.

#### High Throughput Phenotyping of Overlapping Gene Candidates

ORFs of OGCs were cloned in pBAD/myc-HisC using standard cloning techniques without any tag. Each plasmid was transformed into *E. coli* O157:H7 EDL933. To create a bacteria pool, cells were cultivated in LB supplemented with ampicillin (100 μg/ml) in microtiter plates and optical density was measured. The cultures were diluted to OD_600_ = 0.5 and cells were combined in equal amounts. The cell count of the mixture was determined and LB medium, containing 100 μg/ml ampicillin, supplemented with different stressors was inoculated with approximately 1×10^5^ cells. The stressors and their final concentration are given in Table S3. The cultures were cultivated for 22 h at 37°C and 150 rpm. Expression of the OGCs was induced at time points t_0h_ and t_6.5h_ with 0.002% arabinose, since EHEC is able to degrade arabinose. After cultivation, plasmids were isolated, 1 μg was linearized with *NcoI* and further fragmented by ultrasonication to 350 bp (Covaris settings: Peak Incident Power 140W, Duty Factor 10%, Cycles Per Burst 200, treatment time 80 s). Plasmid fragments were processed with the TruSeq DNA PCR-Free Library Prep Kit according to the manufacturer’s protocol. Samples were pooled, quantified with the Perfecta NGS Library Quantification Kit for Illumina (Quanta Bioscience) and sequenced paired end on an Illumina MiSeq. Sequencing reads were processed with Fastq Groomer and mapped to an artificial sequence containing all OGCs with Bowtie2 using standard settings. Forward and reverse read samples were combined with MergeSamFiles. RPKM values for each candidate gene-ORF was determined using the Artemis browser (Rutherford et al., 2000) and a z-score was calculated (z_i,k_ = (x_i,k_ - x_i_) /σ_i_, with x_i.k_ the RPKM value of candidate i in condition k, x_i_ the mean RPKM and σ_i_ the standard deviation of the candidate in all conditions).

#### Competitive Growth Analysis of Single Genes

Competitive growth experiments were conducted with *E. coli* O157:H7 EDL933 containing pBAD/myc-HisC with either the intact OGC-ORFs or the translationally arrested mutated version of the OLG-ORF, which were cloned either using overlap extension PCR and standard cloning methods or QuikChange mutagenesis. Overnight cultures of EHEC EDL933 containing plasmids with either intact or mutated OGC were diluted to OD_600_ = 1 in LB and mixed in equal amounts. Four ml of this mixture was pelleted (12,000×g, 5 min, 4°C) and stored at −20°C until further processing of the samples. Those cells were used as time point zero reference (t0). A 10-ml LB culture containing 100 μg/ml ampicillin with appropriate stressors (see Table S3), was inoculated 1:30000 with the bacteria mix (100 μl of 1:300 dilution). For each growth experiment, a LB control condition was inoculated as well. Protein expression was induced with 0.002% arabinose at time points t0h and t6.5h. Plasmids of t0-samples and samples after cultivation (t = 22 h) were isolated and sequenced with Sanger sequencing. Peak heights of wild type and mutation site were measured and peak height ratios were determined. A two-tailed paired t-test was performed to determine significant differences of bacteria ratios between before and after cultivation at the significance level of α = 0.05.

#### Cappable-Seq for Transcriptional Start Site and Promoter Determination

The protocol of Ettwiller et al. (2016) was applied. Briefly, RNA was isolated with Trizol from early exponential and early stationary phase cultures of EHEC EDL933 in plain LB (OD_600_ = 0.3 and OD_600_ = 3.5), LB supplemented with L-malic acid (4 mM, OD_600_= 0.3 and OD_600_ = 4), LB supplemented with NaCl (500 mM, OD_600_ = 0.2 and OD_600_ = 1), and M9 minimal medium supplemented with 100 mM casamino acids and 5•10^−5^% thiamine; OD_600_ = 0.2 and OD_600_ = 1.5). Genomic DNA was digested with Turbo DNase (Invitrogen) and success of the digest was determined with a PCR reaction using primers binding to 16S ribosomal DNA. Sample preparation for Cappable-Seq and sequencing (75 bp, single end, and 10 million reads for each library) was performed by Vertis Biotechnologie AG (Freising).

Adapter sequences of sequencing reads were trimmed with Cutadapt (Martin, 2011) and reads were quality trimmed with Trimmomatic (Bolger et al., 2014). Remaining reads were mapped to the genome of EHEC (NZ_CP008957; Latif et al., 2014) using Bowtie2. Transcriptional start sites (TSS) were determined with two programs provided by Ettwiller et al. (2016), namely bam2firstbasegtf.pl (trims mapped reads to most 5’ base) and cluster_tss.pl (clusters nearby TSS to the best position). All analyses were performed with custom BASH and perl scripts (available on request). Gene associated TSS were searched in the region of 250 bp upstream of the assumed start codon. Only positions present in all three biological replicates were considered as reliable TSS. Automatic TSS identification was curated manually for OGCs. Differential TSS signals were determined with the *Bioconductor* package *edgeR* (v 3.28.0, McCarthy et al., 2012; Robinson et al., 2010) based on genome wide TSS positions identified in the analyzed samples. The fold change (significance cutoff log2FC > |1|) between stress and non-stress (LB) conditions in each growth phase and differences between growth phases were calculated and p-values were adjusted for multiple testing (FDR according to Benjamini and Hochberg, 1995, significance cutoff FDR < 0.05). Promoter motifs were created with the program weblogo3.0 (Crooks et al., 2004). Towards this end, 100 bp upstream of each gene associated TSS were used for logo creation with standard settings.

#### Western Blots of Overlapping Gene Candidates

OGCs were cloned in a pBAD derivative containing a SPA-tag, pBAD-C+SPA; the SPA was created according to Zeghouf et al. (2004), and cloned using standard techniques. Plasmids were transformed in *Escherichia coli* Top10 and recombinant proteins were expressed in 10 ml LB medium (induction with 0.002% L-arabinose at OD_600_ = 0.3). Controls (positive, protein Gst with 22 kDa in pBAD-C+SPA; negative, empty pBAD-C+SPA in *E. coli* Top10; mock, *E. coli* Top10 without plasmid) were treated in the same way. Samples of 1 ml were taken after 4 h of induction. Pellets were resolved in 50 μL sample buffer (2% SDS, 2%-mercaptoethanol, 40% glycerin, 0.04% Coomassie blue G250, 200 mM Tris/HCl at pH 6.8), heated at 95°C for 10 min and stored at −20°C. Prior to loading on the SDS gel, samples were centrifuged to collect cell debris. Two and a half micro liter of pBAD-C+SPA+gst sample as well as 5 μL of Spectra Multicolor Low Range Protein Ladder (Thermo Scientific) and 10 μl of samples or controls were loaded on tricine-SDS-gels (Schägger, 2006). Briefly, the separation gel had 16% and the stacking gel 4% polyacrylamide, and 3.3% crosslinking material was used for both. Protein separation was conducted at 25 mA per gel in 1× anode buffer and 1× cathode buffer. Gels were blotted on PVDF membrane (Merck, PSQ membrane, 0.2 μm) for 15 - 20 min at 12 V and subsequently blocked in nonfat dried milk at 4°C over night. The membrane was incubated after three washing steps with TBS-T in an appropriate amount of 1:1000 dilution of primary SPA alkaline phosphatase (AP) conjugated antibody in TBS-T for 1 h. Antibody detection was carried out with NBT and BCIP (100 μL NBT and 125 μL BCIP in 10 ml 0.1m Tris with 4 mM MgCl2 at pH 9.5). The reaction was stopped after removing BCIP/NBT with 3% trichloroacetic acid. Molecular weights were estimated using a linear standard curve generated with the protein marker (log[MW] vs. relative migration distance) in each blot (Dunker and Rueckert, 1969; Shapiro et al., 1967).

## SUPPLEMENTAL INFORMATION

Supplemental Information includes one file, five figures, and twelve tables.

## AUTHOR CONTRIBUTIONS

B.Z. conducted the analysis of overlapping ORFs in EDL933 for phenotypes, their expression, testing in competitive growth (global and single genes), and Cappable-Seq. Z.A. conducted the bioinformatics part of the study and helped in drafting the manuscript. M.K. conducted experiments with LF82. S.S. supervised the study and helped to draft the manuscript. K.N. supervised laboratory work, helped to design and perform all experimental analyses of this study and wrote the manuscript. All authors contributed to the final writing of the manuscript.

## ACKNOWLEDGMENTS

We would like to thank Romy Wecko for excellent technical assistance in conducting the molecular biology experiments and Dr. Christopher Huptas for providing the perl script ORFFinder. This work was partially funded by grants from the Deutsche Forschungsgemeinschaft (Sche316/3-1 and Sche316/3- 2) to S.S. under the direction of the SPP-1395 “Informations-und Kommunikationstheorie in der Molekularbiologie (InKoMBio)”.

## SUPPLEMENTAL INFORMATION

Document S1. **Additional analysis of BL21 data from Meydan et al. (2019)**

Figure S1. **Transcription and translation of 216 OGCs identified**

Figure S2. **Transcription start sites and their regulation for OGCs (if available)**

Figure S3. **Uncut Western blots of OGCs**

Figure S4. **Length distributions of BLASTp hits for 304 hits**

Figure S5. **N-terminal peptide fragments of overlapping genes in *E. coli* MG1655**

Table S1. **Accession numbers**

Table S2. **Footprint coverage values for all strains analyzed**

Table S3. **Supplements used in competitive growth experiments**

Table S4. **Overview of 864 expressed overlapping ORFs deduced from 12 growth conditions by RNAseq in EDL933**

Table S5. **Overview of experiments and results for all 216 OGCs**

Table S6. **Overview of 216 overlapping ORFs with footprint signals in EDL933**

Table S7. **Overview of 206 OGCs grown in the high-throughput overexpression phenotyping experiment and their z-scores**

Table S8. **Data for competitive growth experiments (shown in Figure 3D-F)**

Table S9. **Results for OGC51**

Table S10. **Additional OLG candidates, embedded and partial OLG in different strains**

Table S11. **Additional OLG candidates of strain BL21**

Table S12. **BLASTp hits in RefSeq for all strains analyzed**

