## Supplementary File S1 for "Evidence for Numerous Embedded Antisense Overlapping Genes in Diverse *E. coli* Strains"

**Additional analysis of BL21 data from Meydan et al. (2018) suggests even more expressed and translated OLGs**

We conducted further analyses of reading frame and start sites in the BL21 data, in each case, as previously, reporting numbers for ORF families. While the original report of the data did not include any antisense embedded genes of at least 30 codons long with their very strict threshold of at least 5 RPM at the peak, we find many good candidates. For instance, we find 103 unique embedded antisense ORFs of at least 30 codons with a start peak site with at least 1 RPM (within 3 nucleotides of a possible start codon), after filtering to the reads between 31 and 38 nucleotides long, which are the most informative for start sites in this data (i.e. at least six reads at a single position by a start site). In the 'non-drug' sample, we find 98 unique embedded antisense ORFs of at least 30 codons, with at least 0.75 RPM (equal to 12 reads) and 50% coverage, and with reads in the correct frame, as determined from annotated genes, predominating. Of these, 46 meet a binomial test for having significantly more than 33.33% of their reads in this frame ($p < 0.05$). Of the candidates with a start peak predicted in the retapamulin treated sample, 16 also were found in the "correct frame" set detected the no drug sample, of which 9 met the binomial test.
