## Supplementary Figure S1 for "Evidence for Numerous Embedded Antisense Overlapping Genes in Diverse *E. coli* Strains"

**Transcription and translation of 216 OGCs identified.** Sequencing reads shown in panels were created using Artemis v17 for selected genomic regions of *E. coli* O157:H7 strain EDL933. The expression signals are from two biological replicates for transcription (RNAseq, red / blue) and for translation (RIBOseq, green / black). In total, 216 overlapping gene candidates outside prophage regions have been found. Each OGC is numbered, and shown as pink / yellow arrows in each panel. The reads matching to the OGC in question have been highligthed in pink in the upper part of the panels. Annotated genes are shown in blue / white arrows. Numbers and gene locus tags correspond to the EDL933 genome (CP008957).

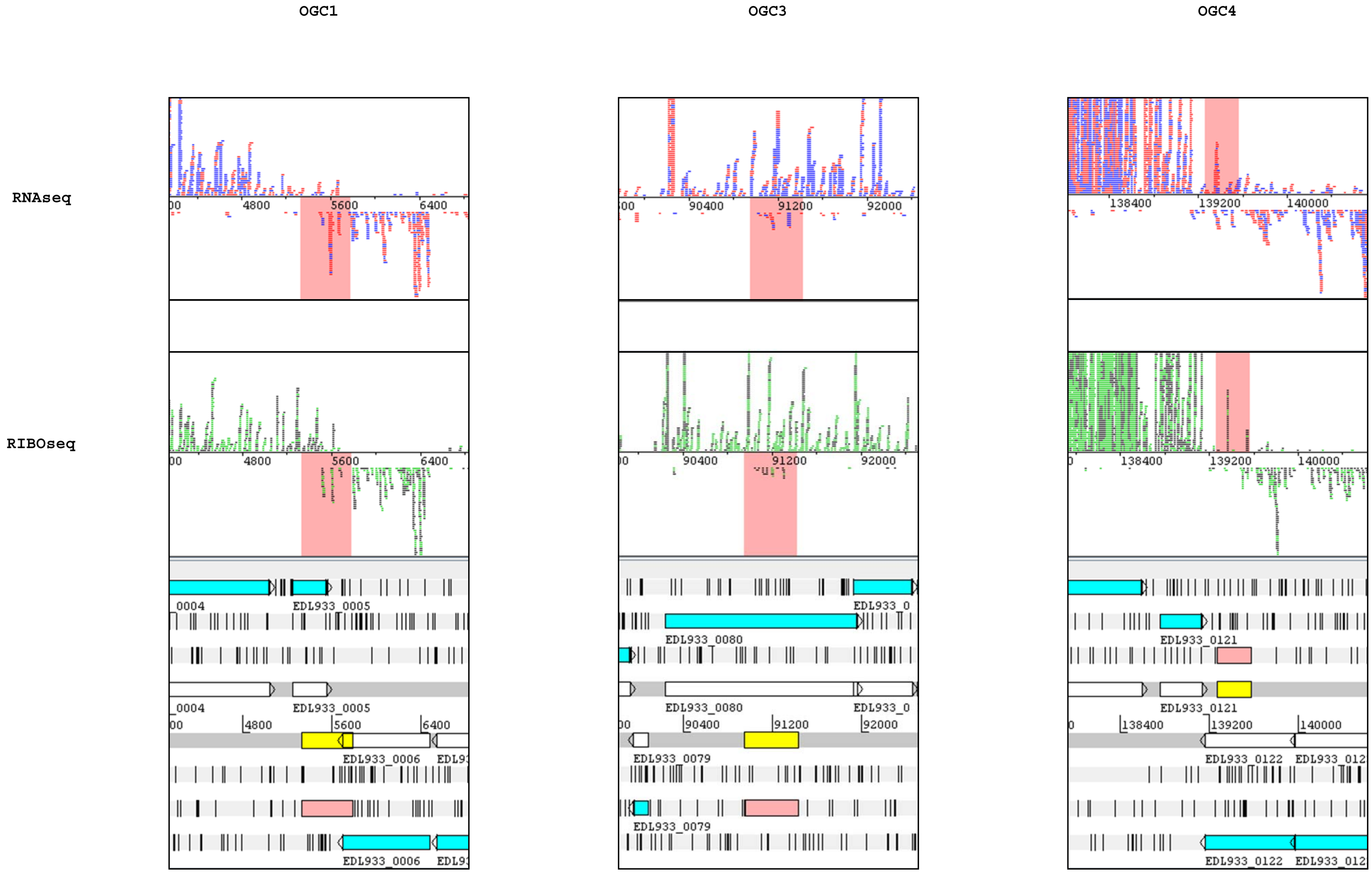
OGC1
OGC3
OGC4
RNAseq
RIBOseq
4800
5600
6400
90400
91200
92000
138400
139200
140000
EDL933_0005
EDL933_0080
EDL933_0121
EDL933_0006
EDL933_0079
EDL933_0122

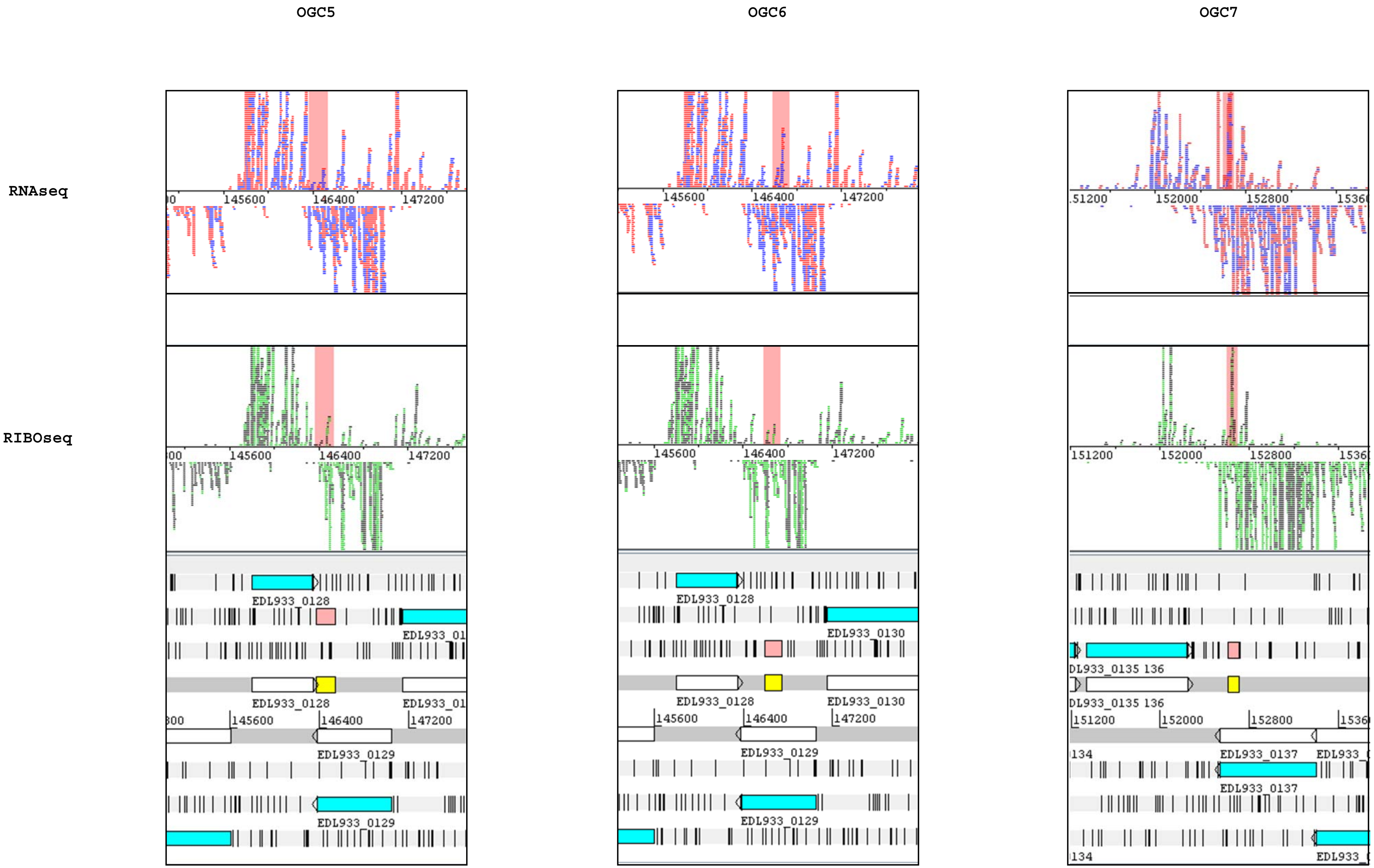
OGC5
OGC6
OGC7
RNAseq
RIBOseq
145600
146400
147200
151200
152000
152800
EDL933_0128
EDL933_0129
EDL933_0130
EDL933_0137
DL933_0135 136

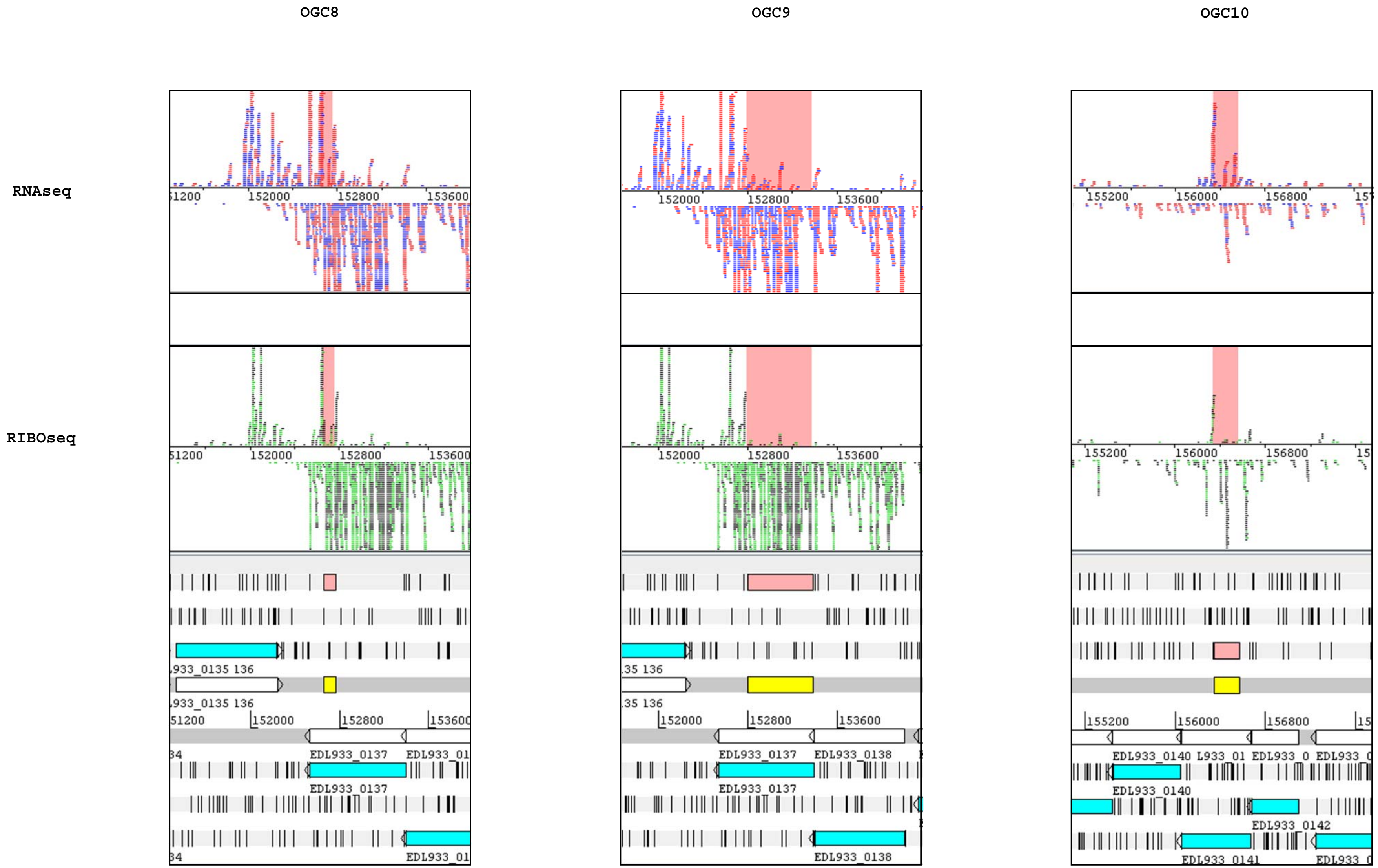
OGC8
OGC9
OGC10
RNAseq
RIBOseq
152000
152800
153600
155200
156000
156800
EDL933_0137
EDL933_0138
EDL933_0140
EDL933_0142
EDL933_0141

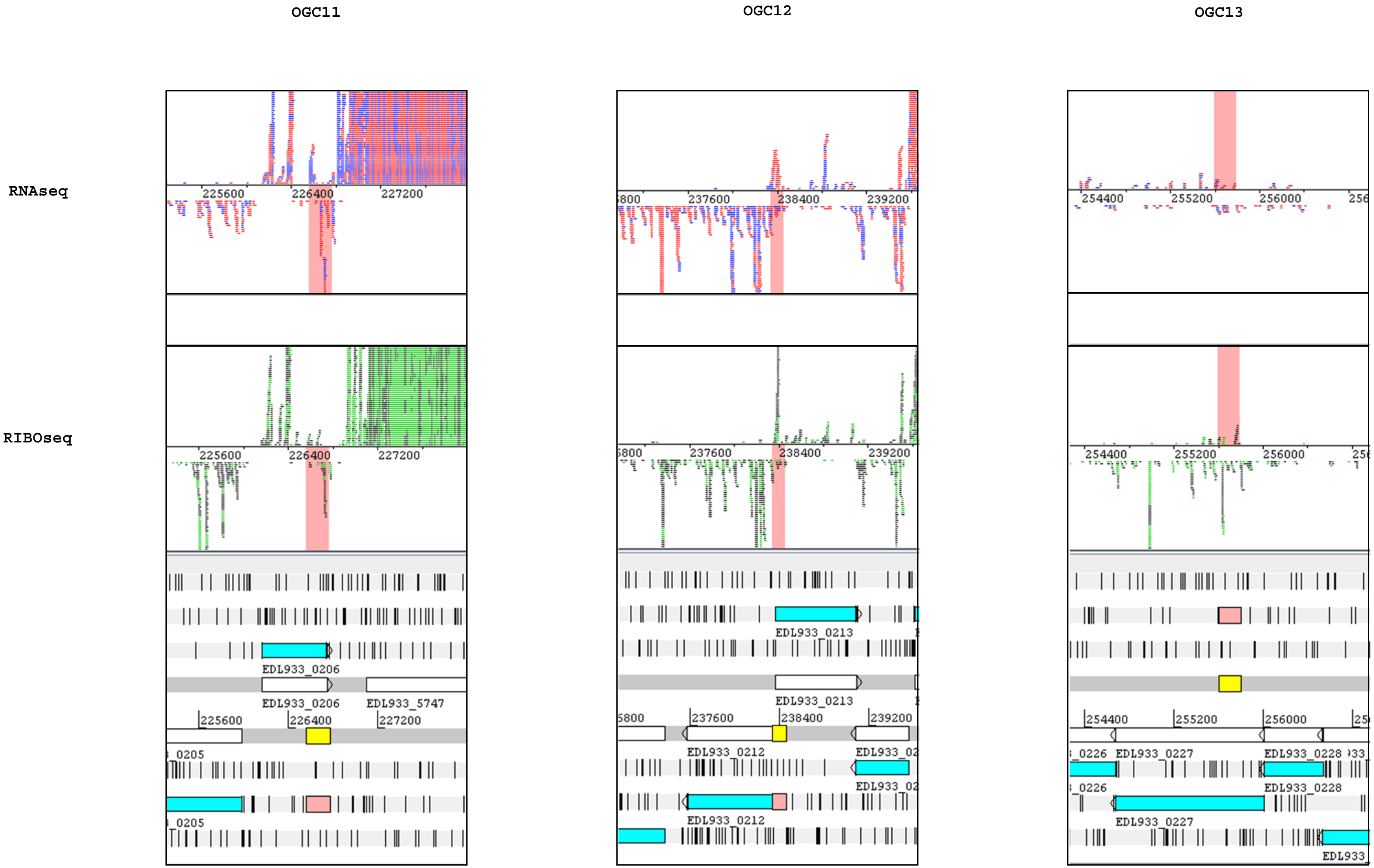
OGC11
OGC12
OGC13
RNAseq
RIBOseq
225600
226400
227200
237600
238400
239200
254400
255200
256000
EDL933_0206
EDL933_5747
EDL933_0213
EDL933_0212
EDL933_0227
EDL933_0228

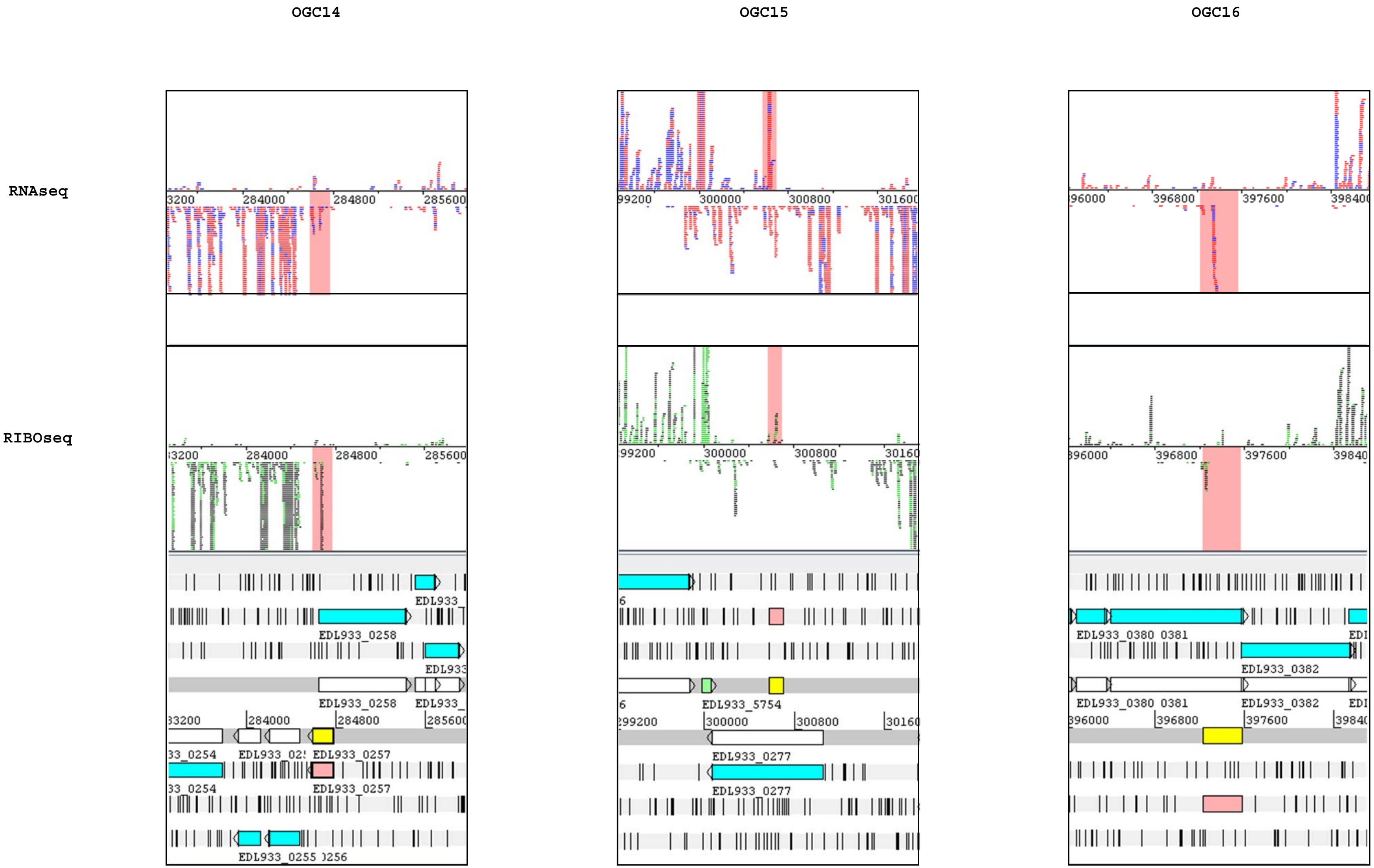
OGC14
OGC15
OGC16
RNAseq
RIBOseq
284000
284800
285600
300000
300800
396800
397600
EDL933_0258
EDL933_0257
EDL933_0255
EDL933_5754
EDL933_0277
EDL933_0380
0381
EDL933_0382

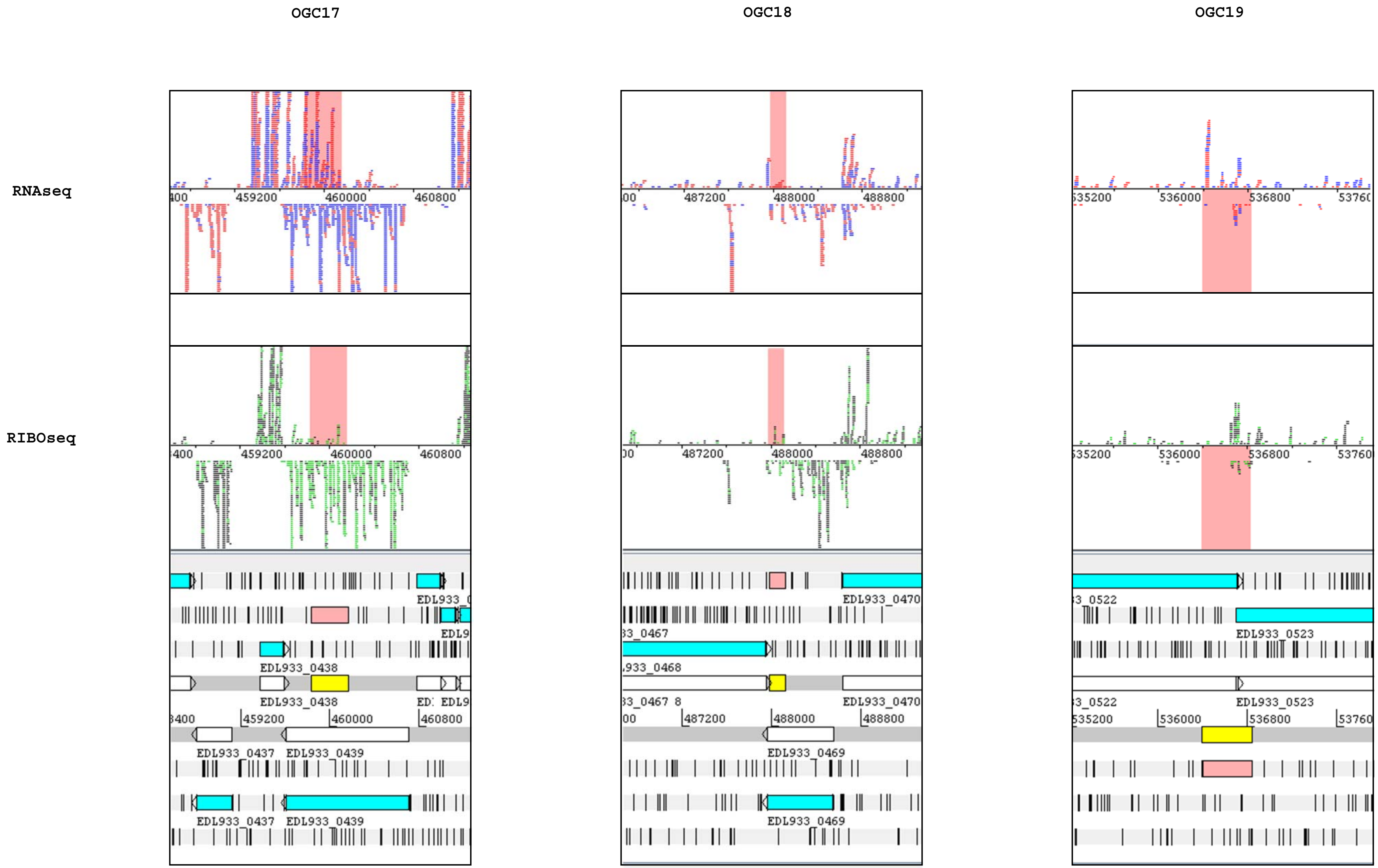
OGC17
OGC18
OGC19
RNAseq
RIBOseq
459200
460000
460800
487200
488000
488800
536000
536800
EDL933_0437
EDL933_0438
EDL933_0439
EDL933_0467
EDL933_0468
EDL933_0469
EDL933_0470
EDL933_0523

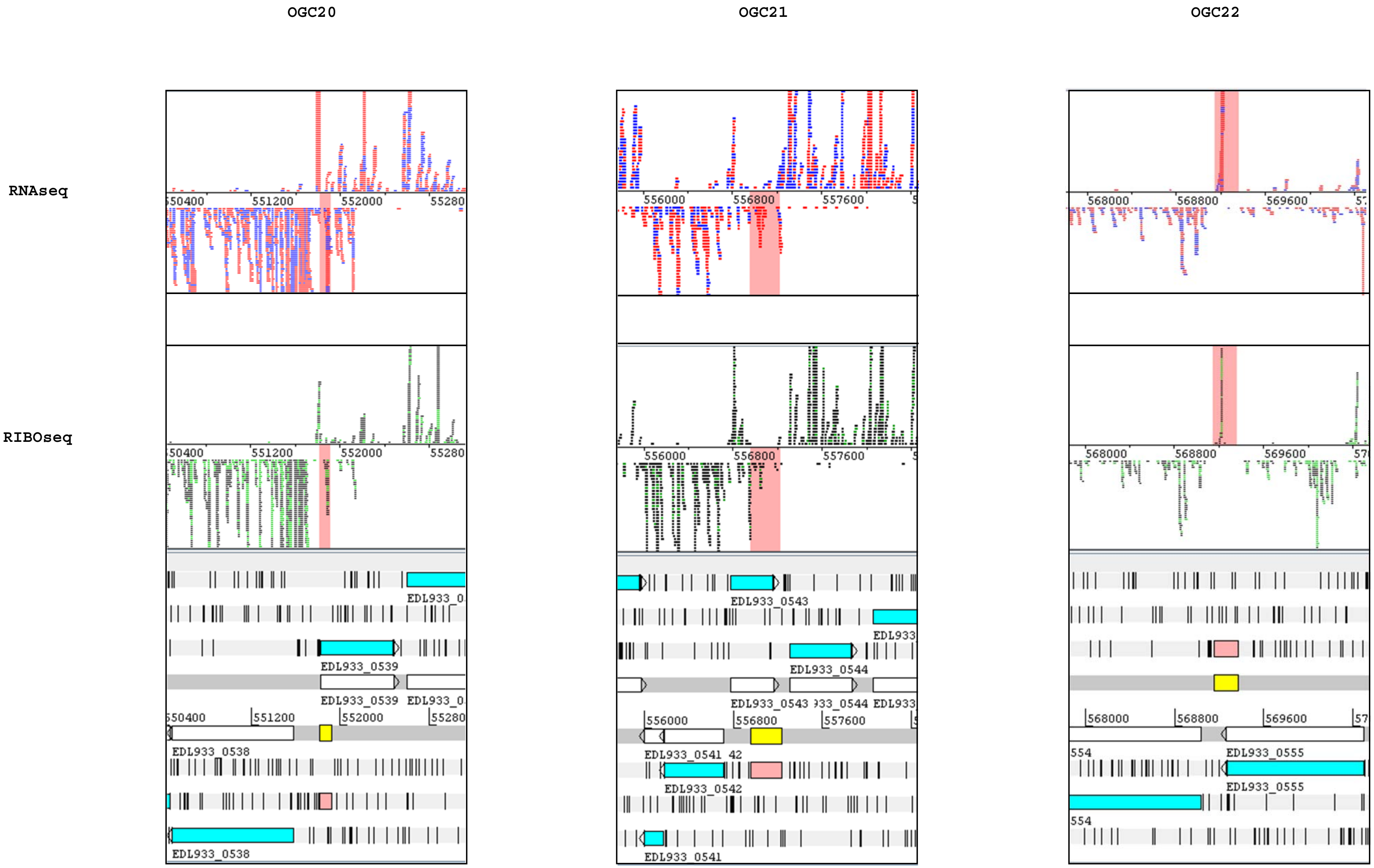

OGC20
OGC21
OGC22
RNAseq
RIBOseq
550400
551200
552000
55280
EDL933_0538
EDL933_0539
EDL933_0
556000
556800
557600
EDL933_0541
EDL933_0542
EDL933_0543
EDL933_0544
EDL933
568000
568800
569600
EDL933_0555
554

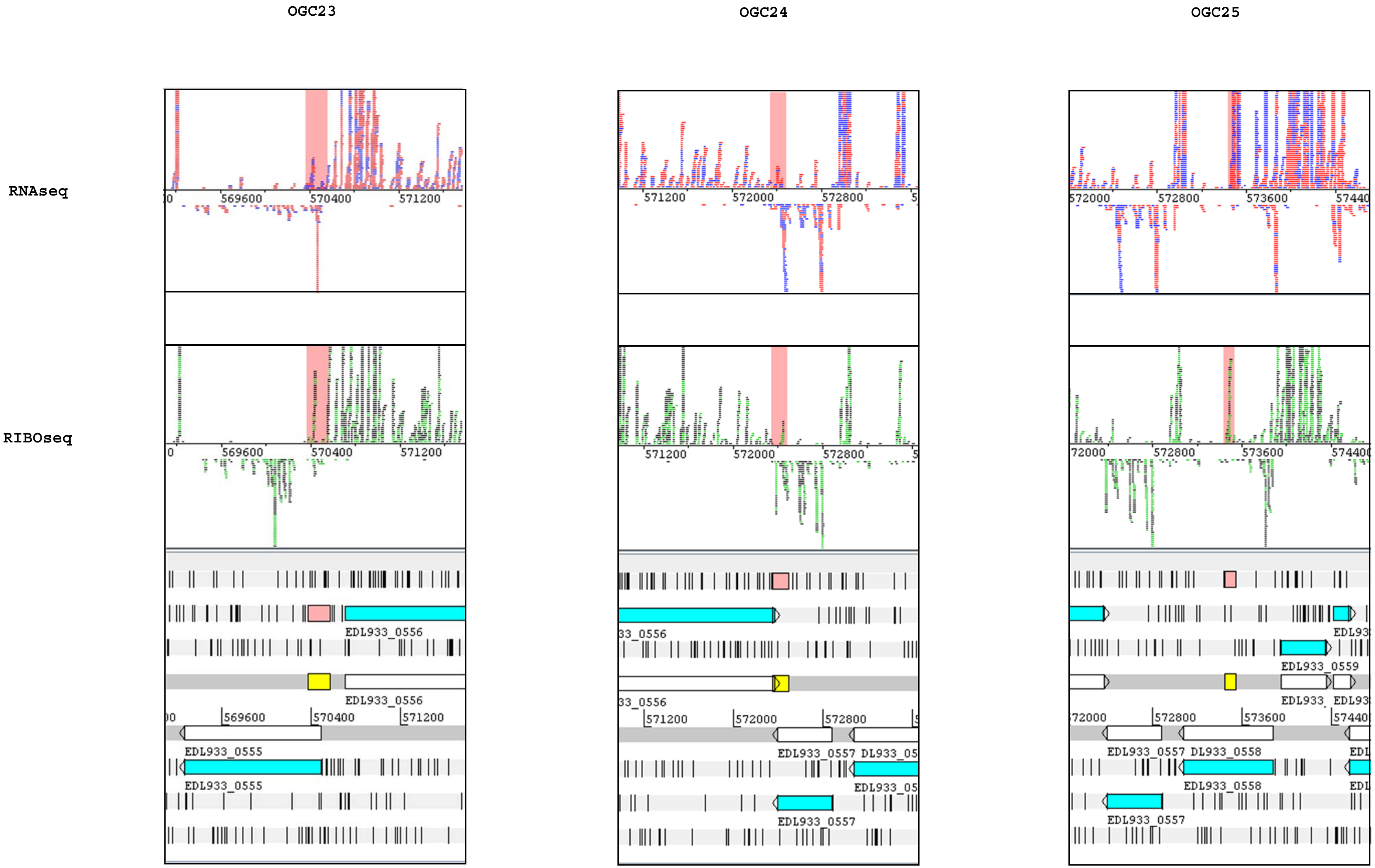
OGC23
OGC24
OGC25
RNAseq
RIBOseq
569600
570400
571200
EDL933_0555
EDL933_0556
571200
572000
572800
EDL933_0557
573600
EDL933_0558
EDL933_0559

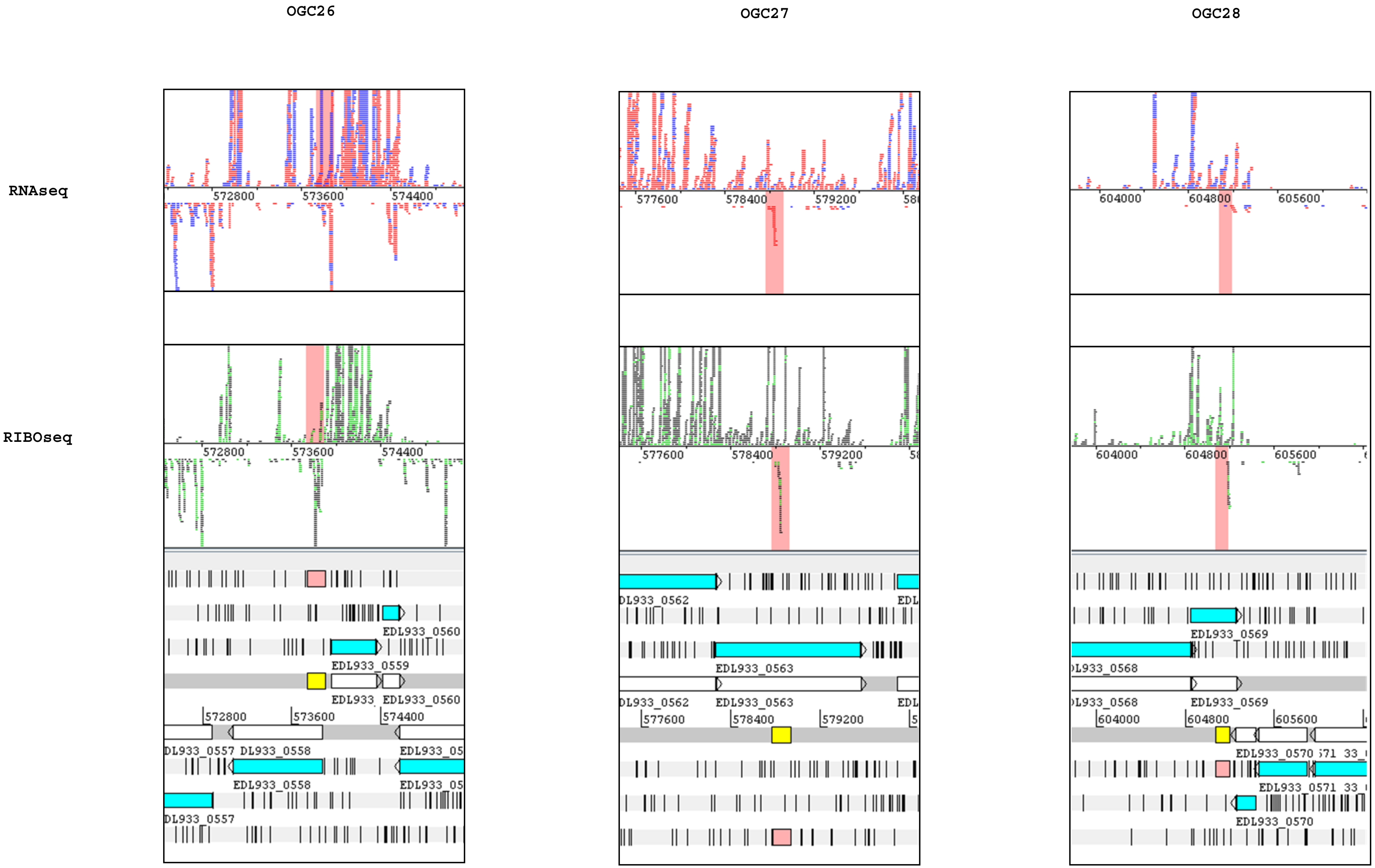
OGC26
OGC27
OGC28
RNAseq
RIBOseq
572800
573600
574400
577600
578400
579200
604000
604800
605600
EDL933_0560
EDL933_0559
EDL933_0558
EDL933_0562
EDL933_0563
EDL933_0569
EDL933_0570

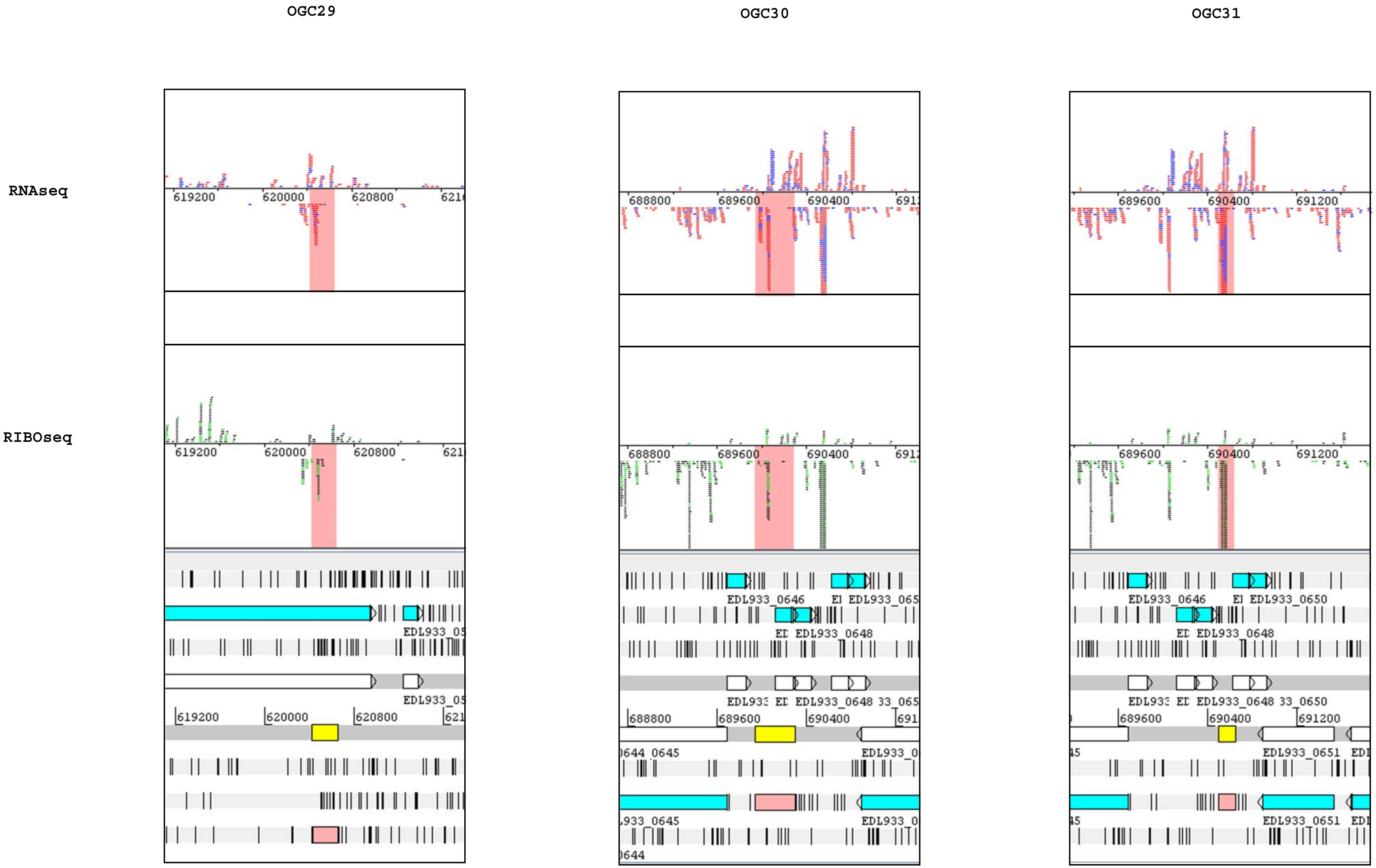
OGC29
OGC30
OGC31
RNAseq
RIBOseq
619200
620000
620800
688800
689600
690400
691200
EDL933_0646
EDL933_0648
EDL933_0650
EDL933_0651
EDL933_0645

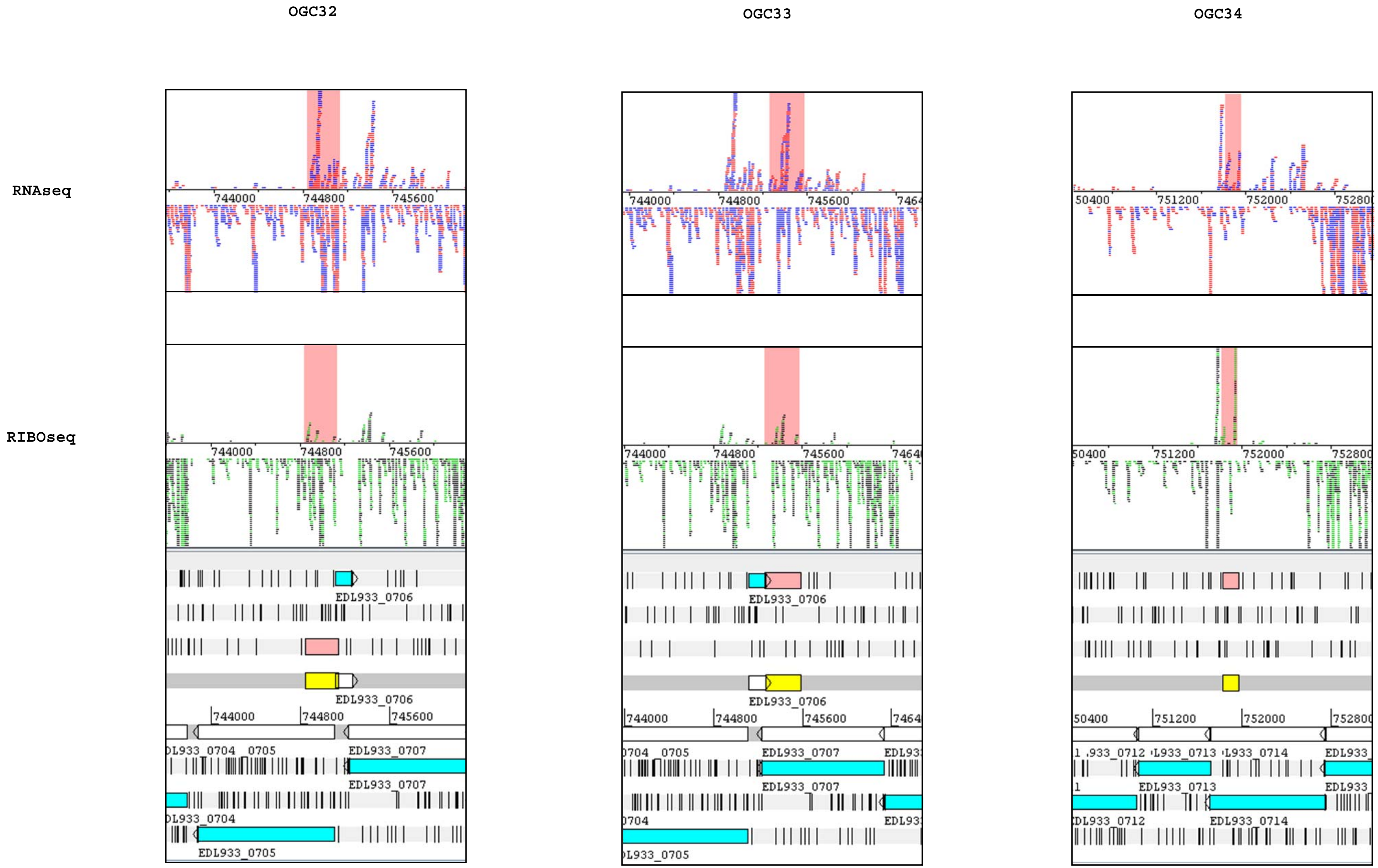
OGC32
OGC33
OGC34
RNAseq
RIBOseq
744000
744800
745600
751200
752000
EDL933_0706
EDL933_0707
EDL933_0705
EDL933_0713
EDL933_0714

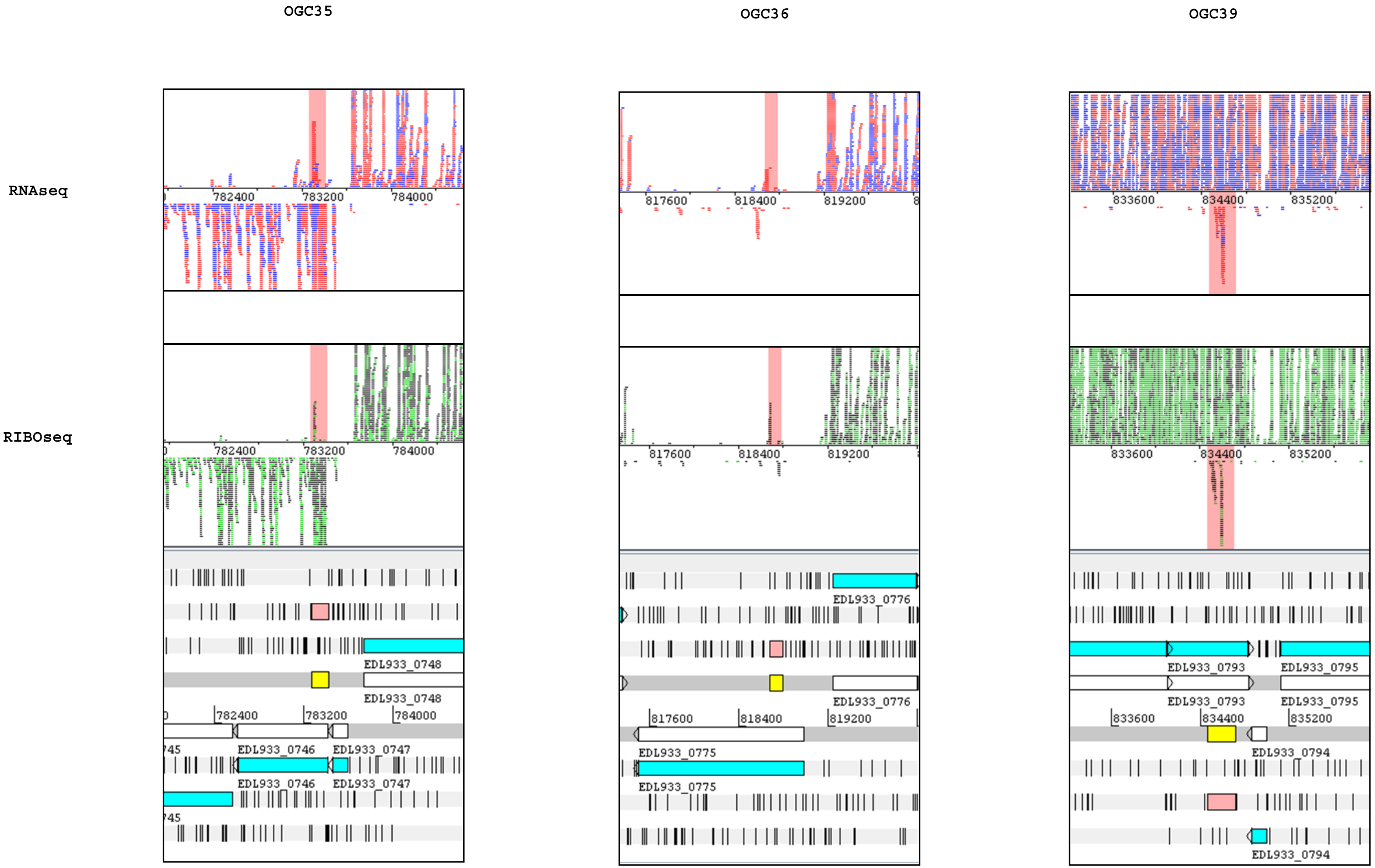
OGC35
OGC36
OGC39
RNAseq
RIBOseq
782400
783200
784000
817600
818400
819200
833600
834400
835200
EDL933_0746
EDL933_0747
EDL933_0748
EDL933_0775
EDL933_0776
EDL933_0793
EDL933_0794
EDL933_0795

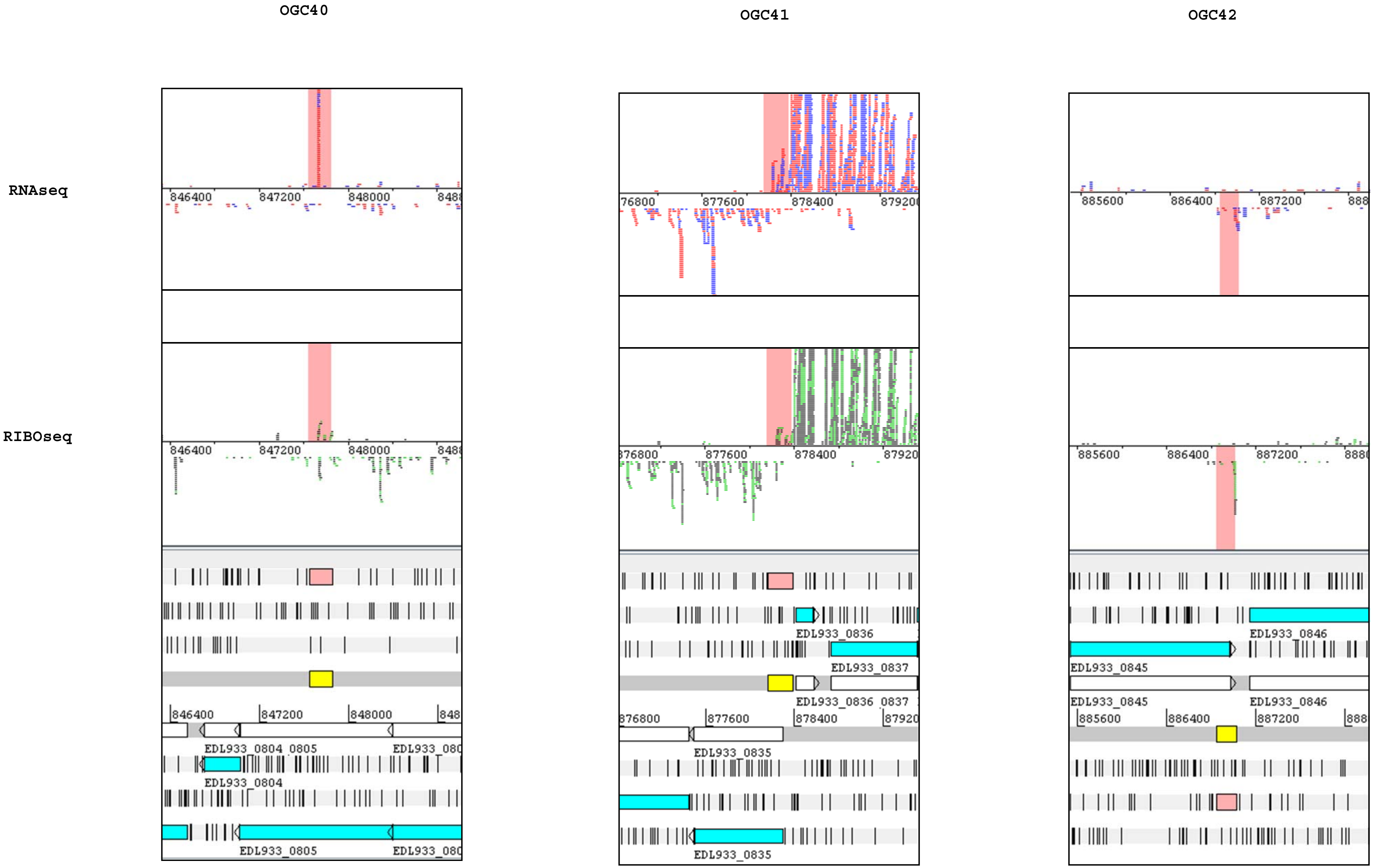
OGC40
OGC41
OGC42
RNAseq
RIBOseq
EDL933_0804_0805
EDL933_0804
EDL933_0805
EDL933_0836
EDL933_0837
EDL933_0836_0837
EDL933_0835
EDL933_0846
EDL933_0845

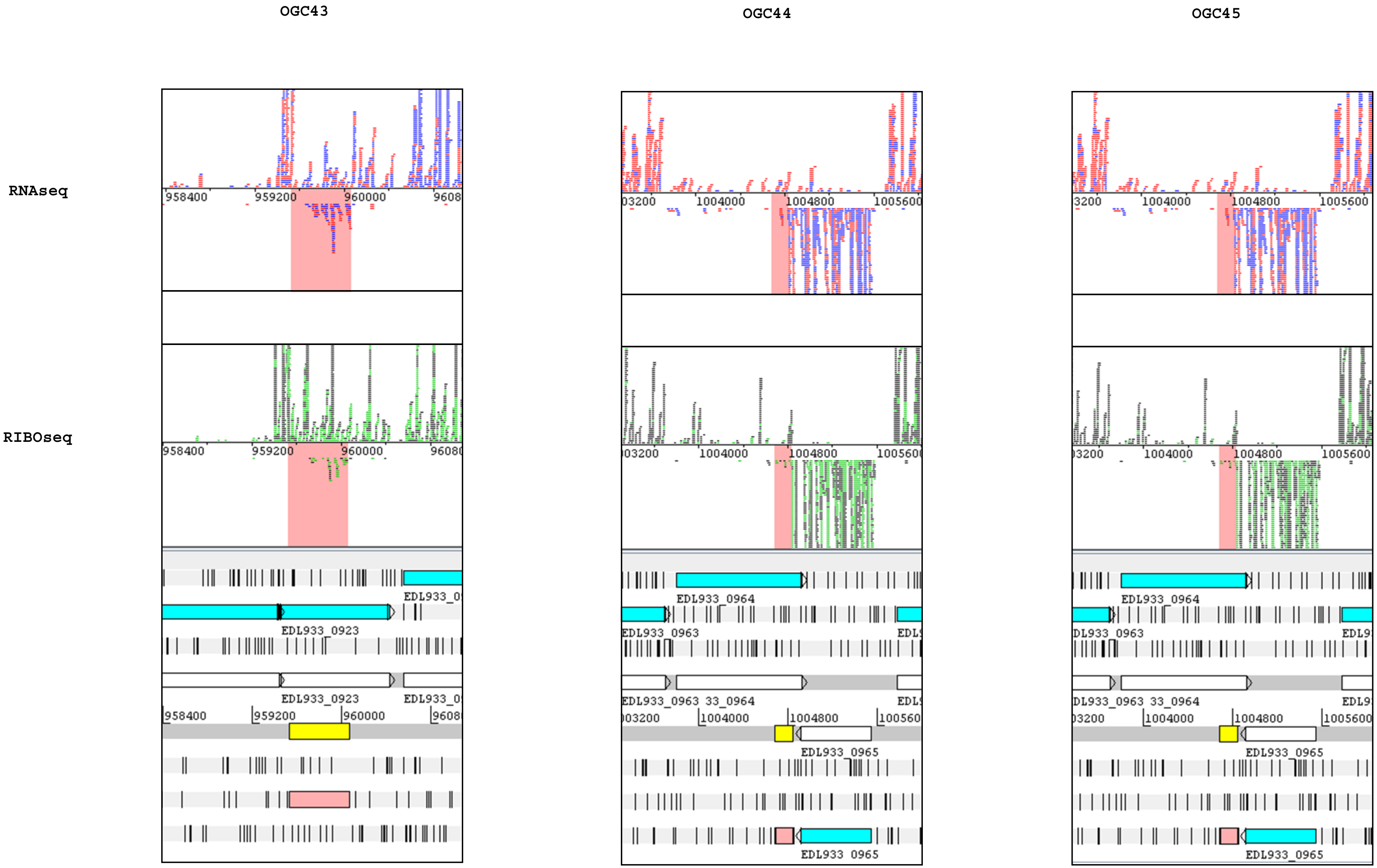

OGC43
OGC44
OGC45
RNAseq
RIBOseq
958400
959200
960000
1004000
1004800
1005600
EDL933_0923
EDL933_0963
EDL933_0964
EDL933_0965

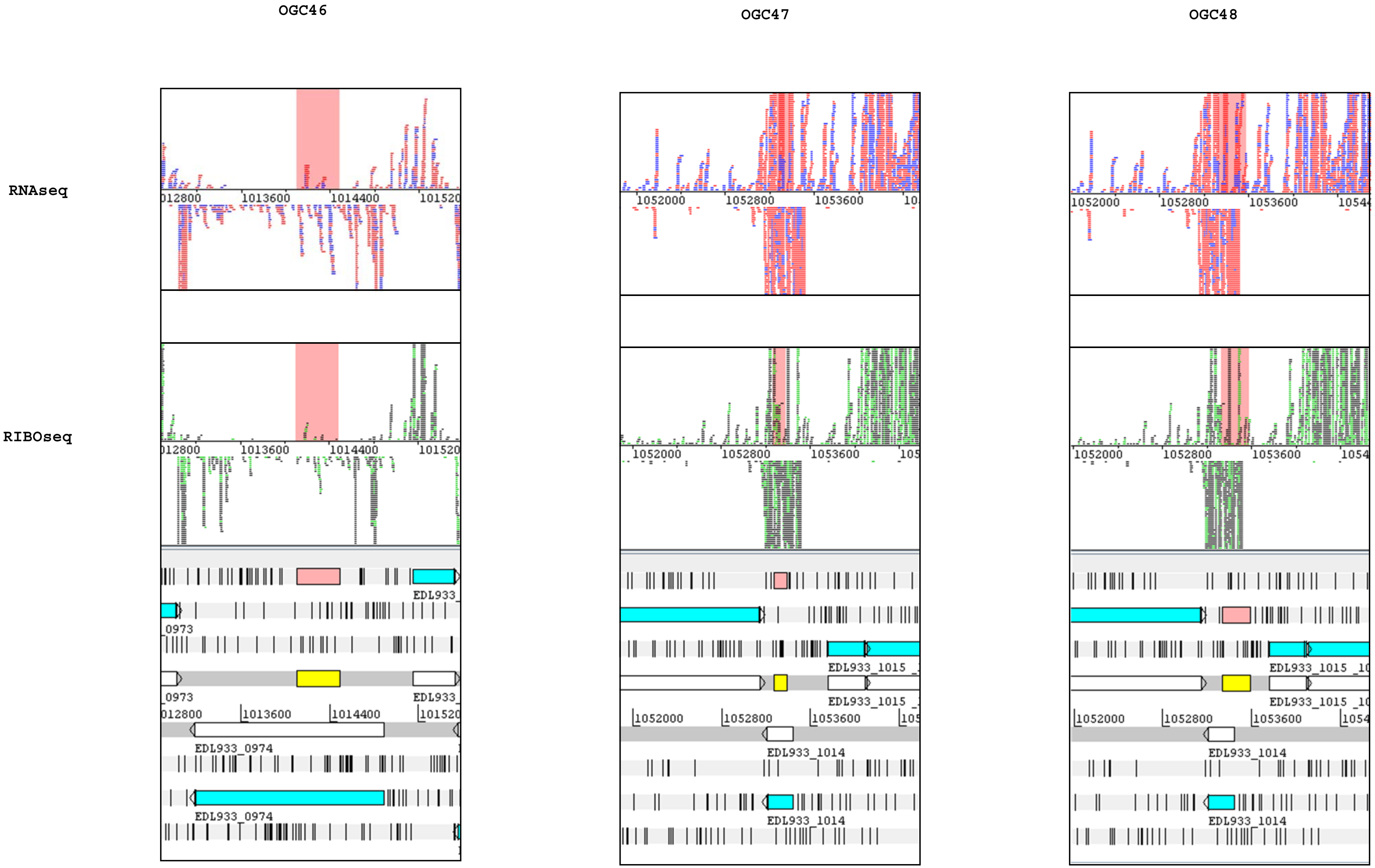

OGC46
OGC47
OGC48
RNAseq
RIBOseq
1013600
1014400
1052000
1052800
1053600
EDL933_0974
EDL933_1014
EDL933_1015

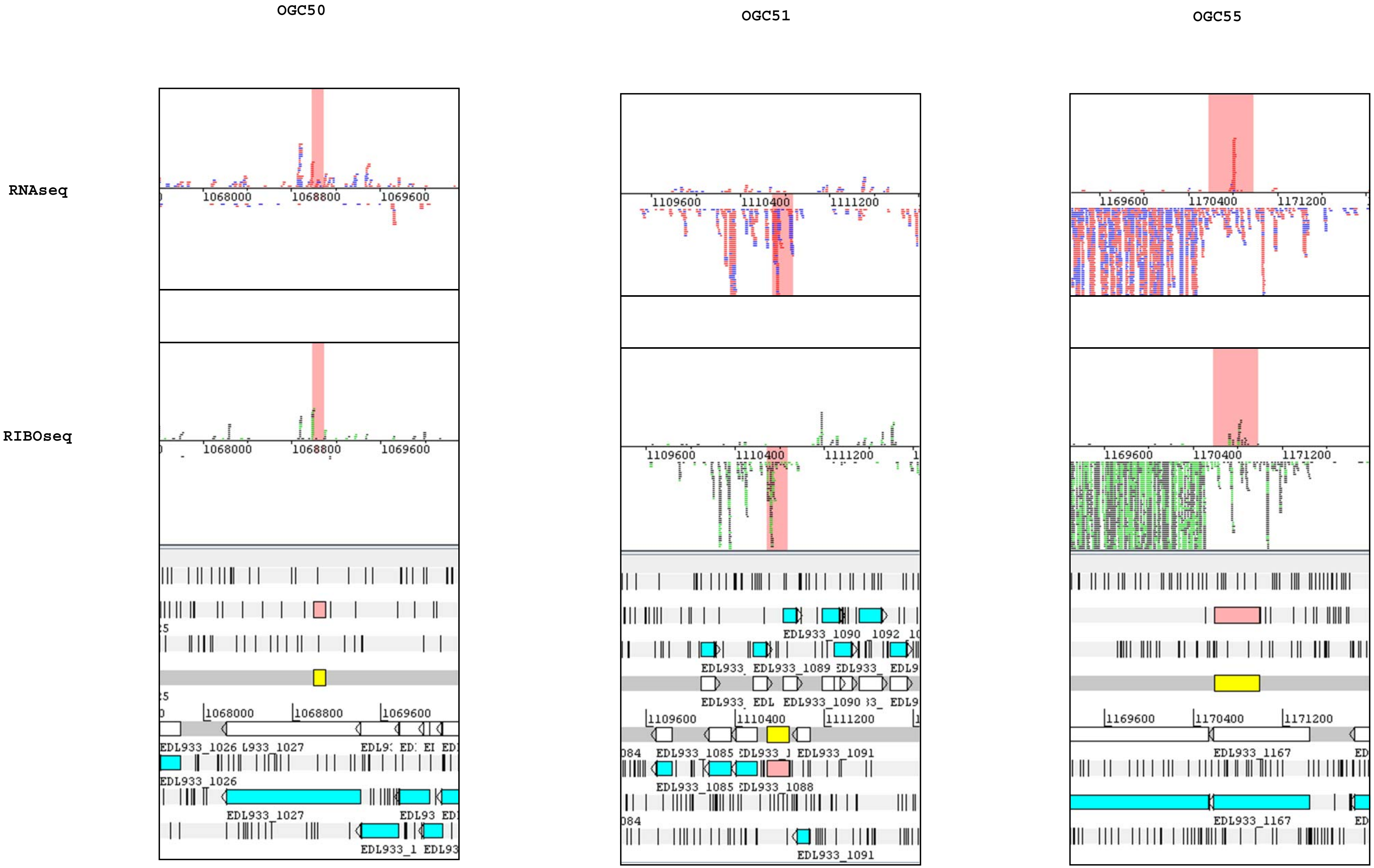
OGC50
OGC51
OGC55
RNAseq
RIBOseq
1068000
1068800
1069600
1109600
1110400
1111200
1169600
1170400
1171200
EDL933_1026
EDL933_1027
EDL933_1085
EDL933_1088
EDL933_1089
EDL933_1090
EDL933_1091
EDL933_1167

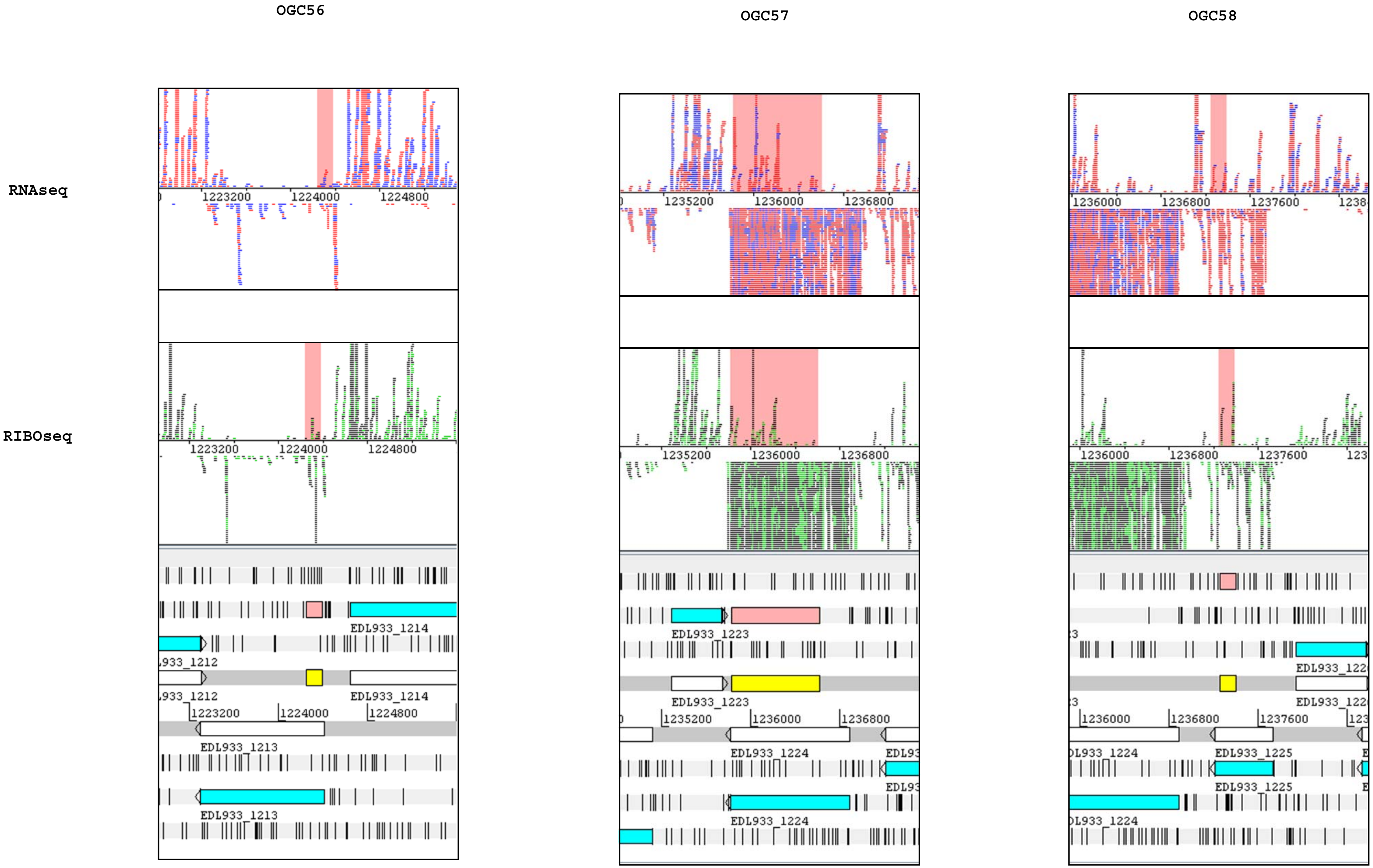
OGC56
OGC57
OGC58
RNAseq
RIBOseq
1223200
1224000
1224800
EDL933_1214
EDL933_1213
1235200
1236000
1236800
EDL933_1223
EDL933_1224
1237600
EDL933_1225

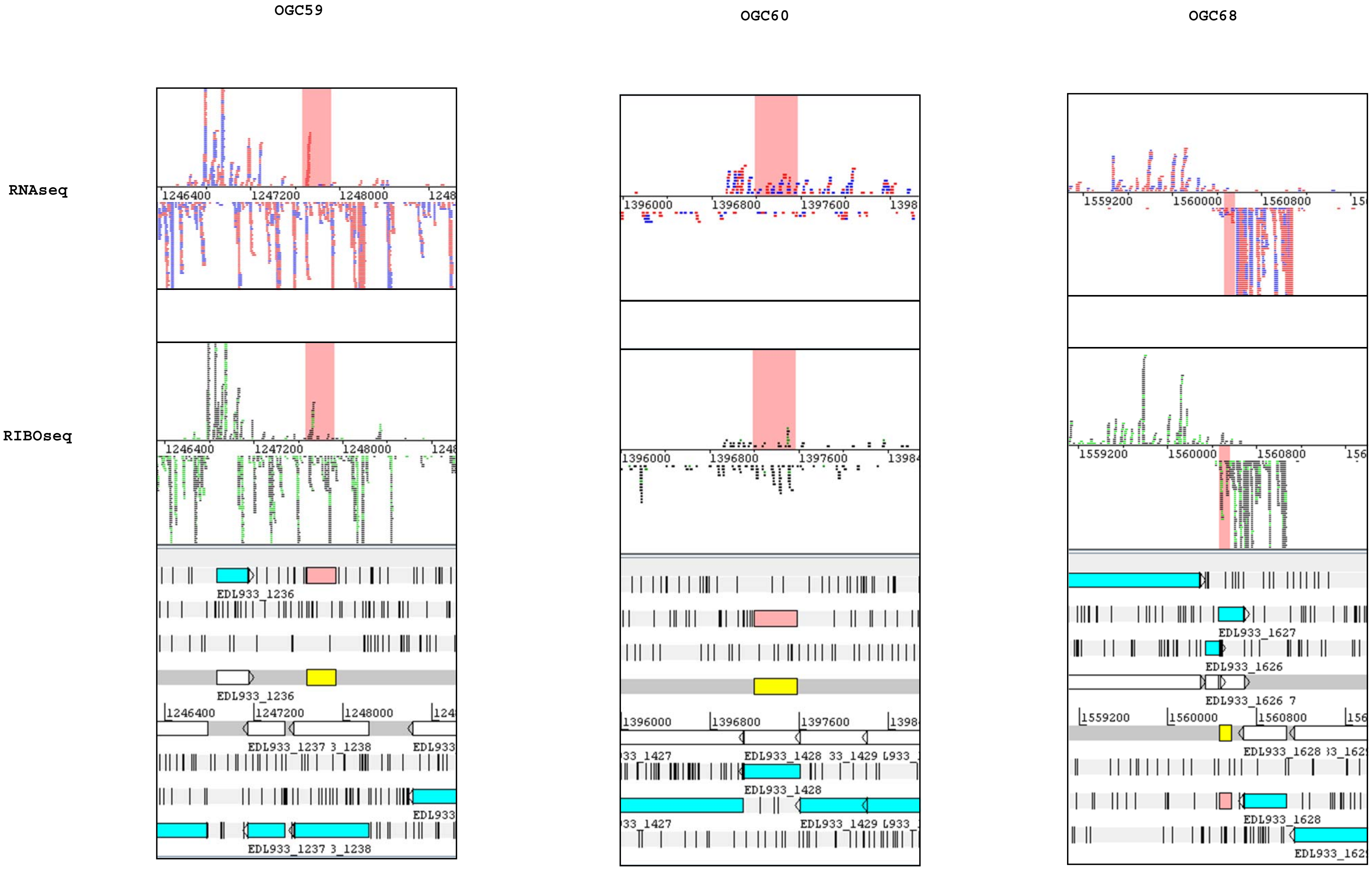
OGC59
OGC60
OGC68
RNAseq
RIBOseq
1246400
1247200
1248000
1396000
1396800
1397600
1559200
1560000
1560800
EDL933_1236
EDL933_1237
EDL933_1428
EDL933_1627
EDL933_1626
EDL933_1628

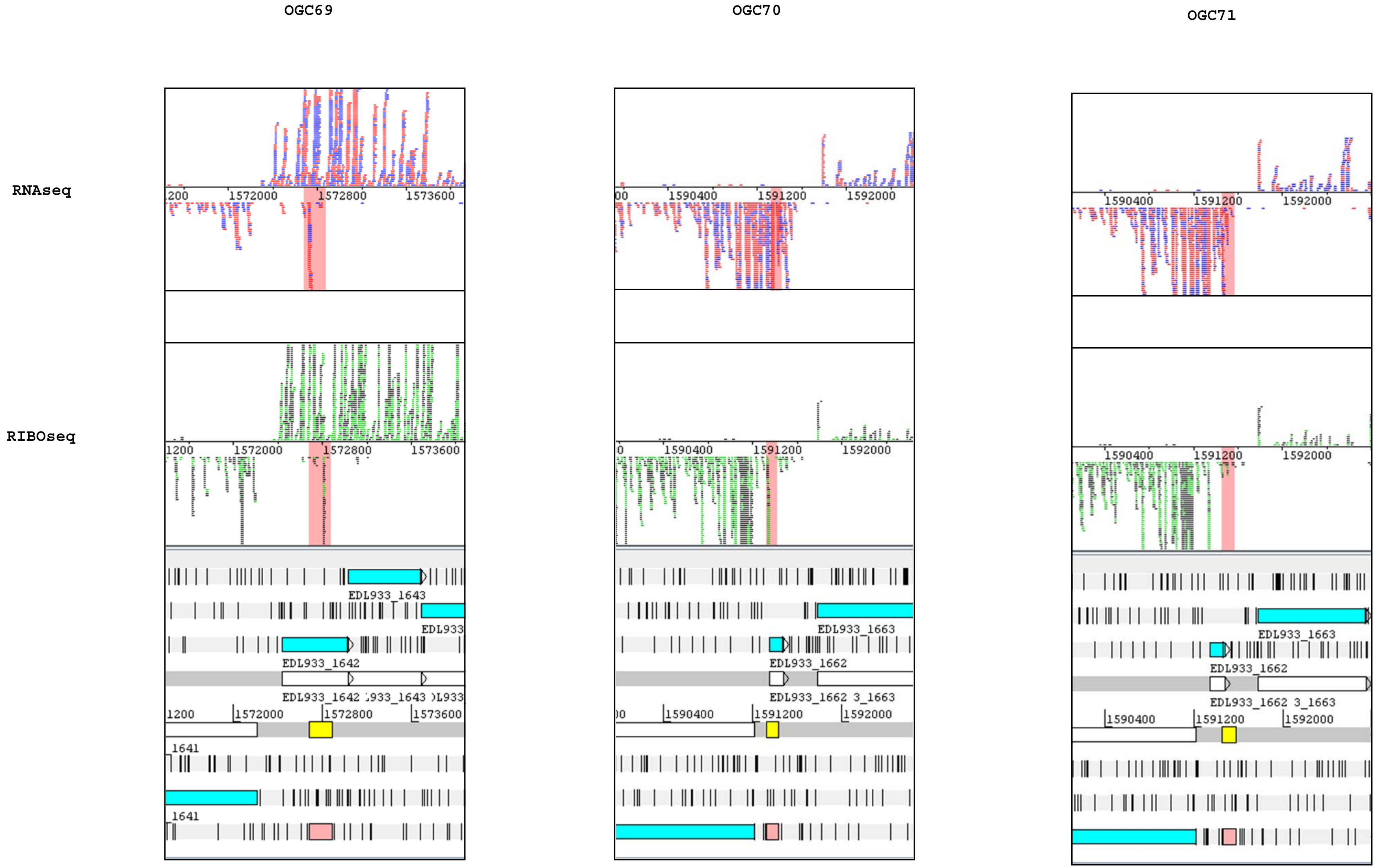
OGC69
OGC70
OGC71
RNAseq
RIBOseq
1572000
1572800
1573600
1590400
1591200
1592000
EDL933_1643
EDL933_1642
EDL933_1662
EDL933_1663

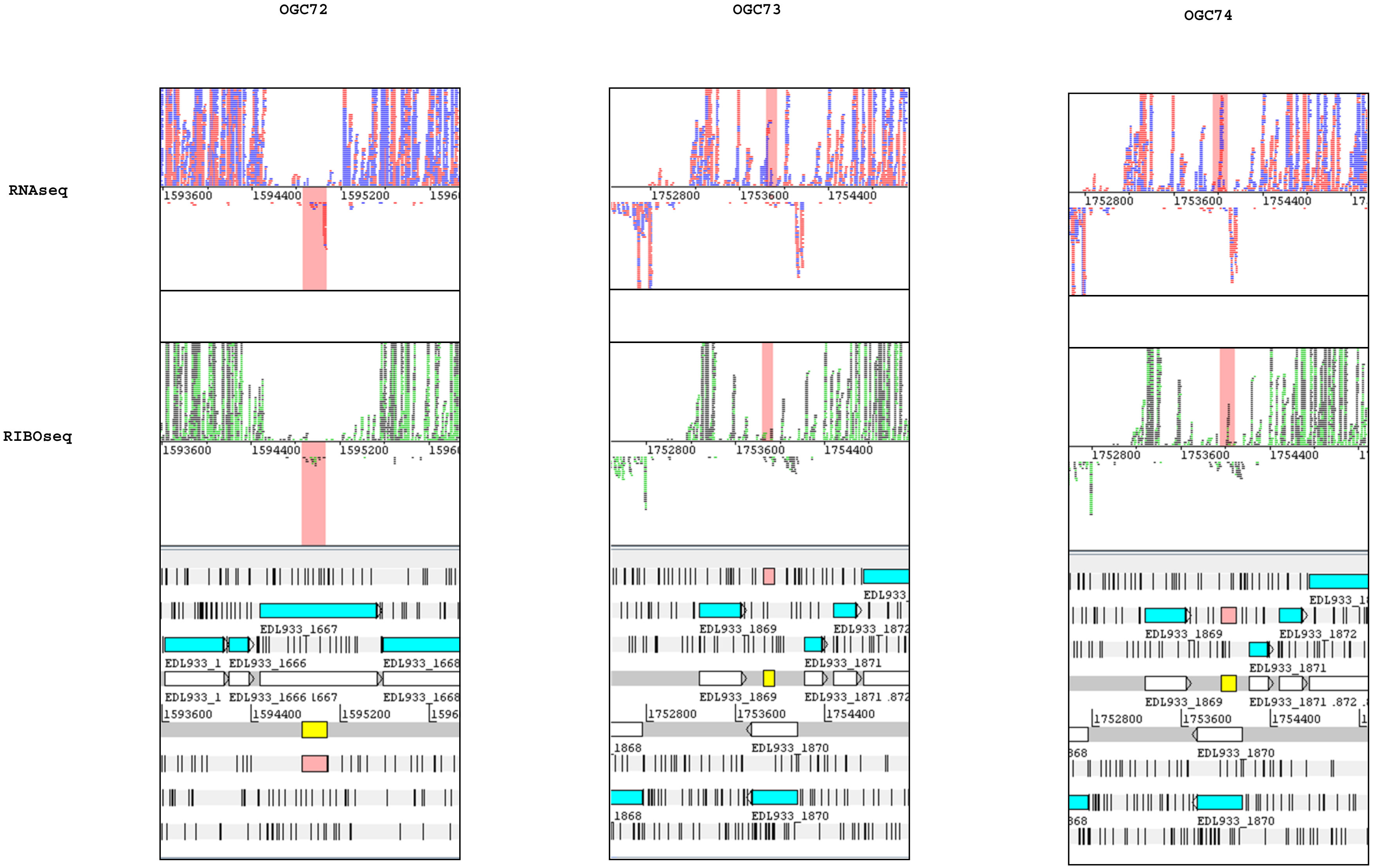
OGC72
OGC73
OGC74
RNAseq
RIBOseq
1593600
1594400
1595200
1752800
1753600
1754400
EDL933_1667
EDL933_1666
EDL933_1668
EDL933_1869
EDL933_1871
EDL933_1872
EDL933_1870

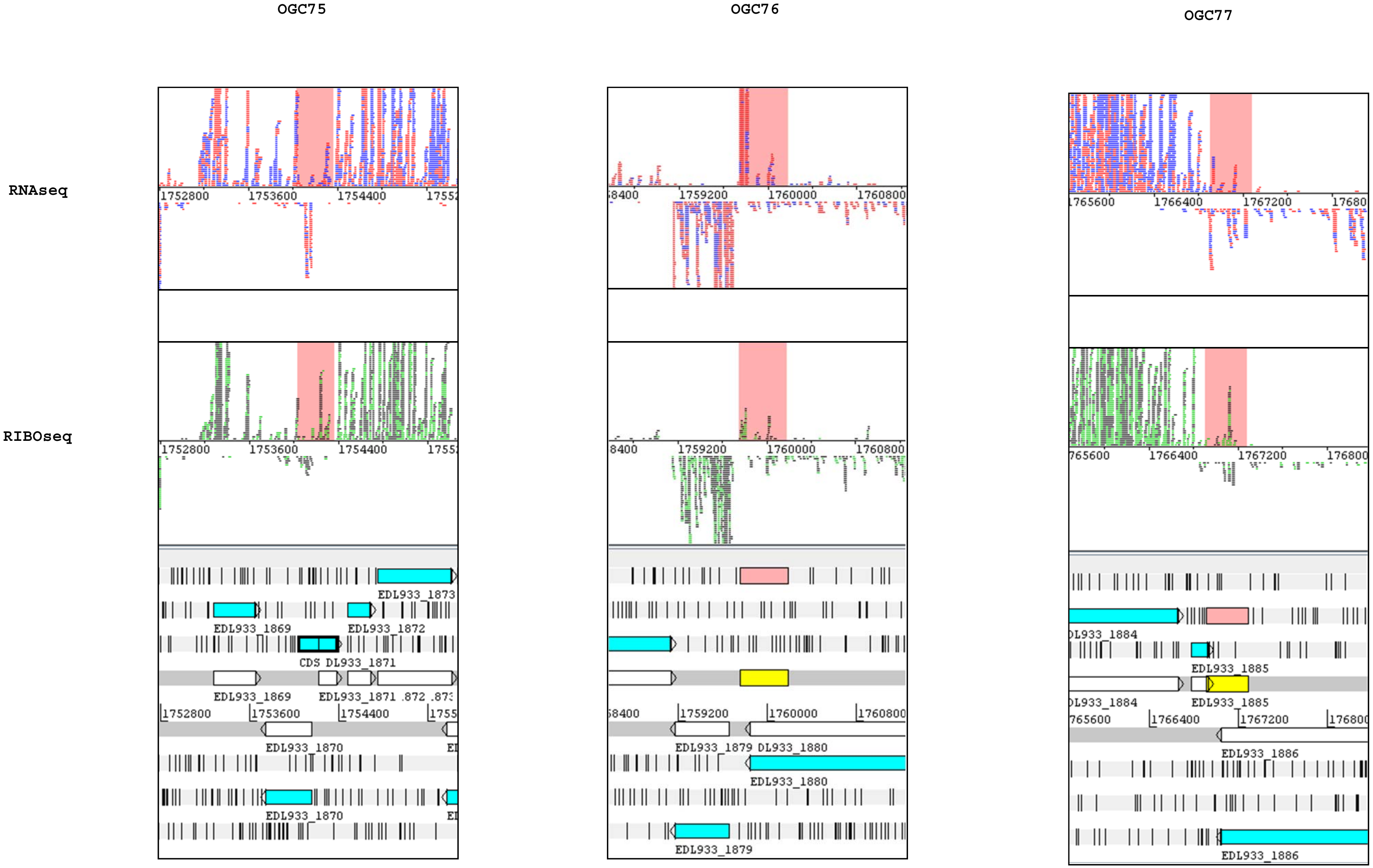
OGC75
OGC76
OGC77
RNAseq
RIBOseq
EDL933_1873
EDL933_1869
EDL933_1872
CDS DL933_1871
EDL933_1871
EDL933_1870
EDL933_1879
DL933_1880
EDL933_1880
EDL933_1885
EDL933_1886

OGC78 OGC79 OGC80

RNAseq

RIBOseq

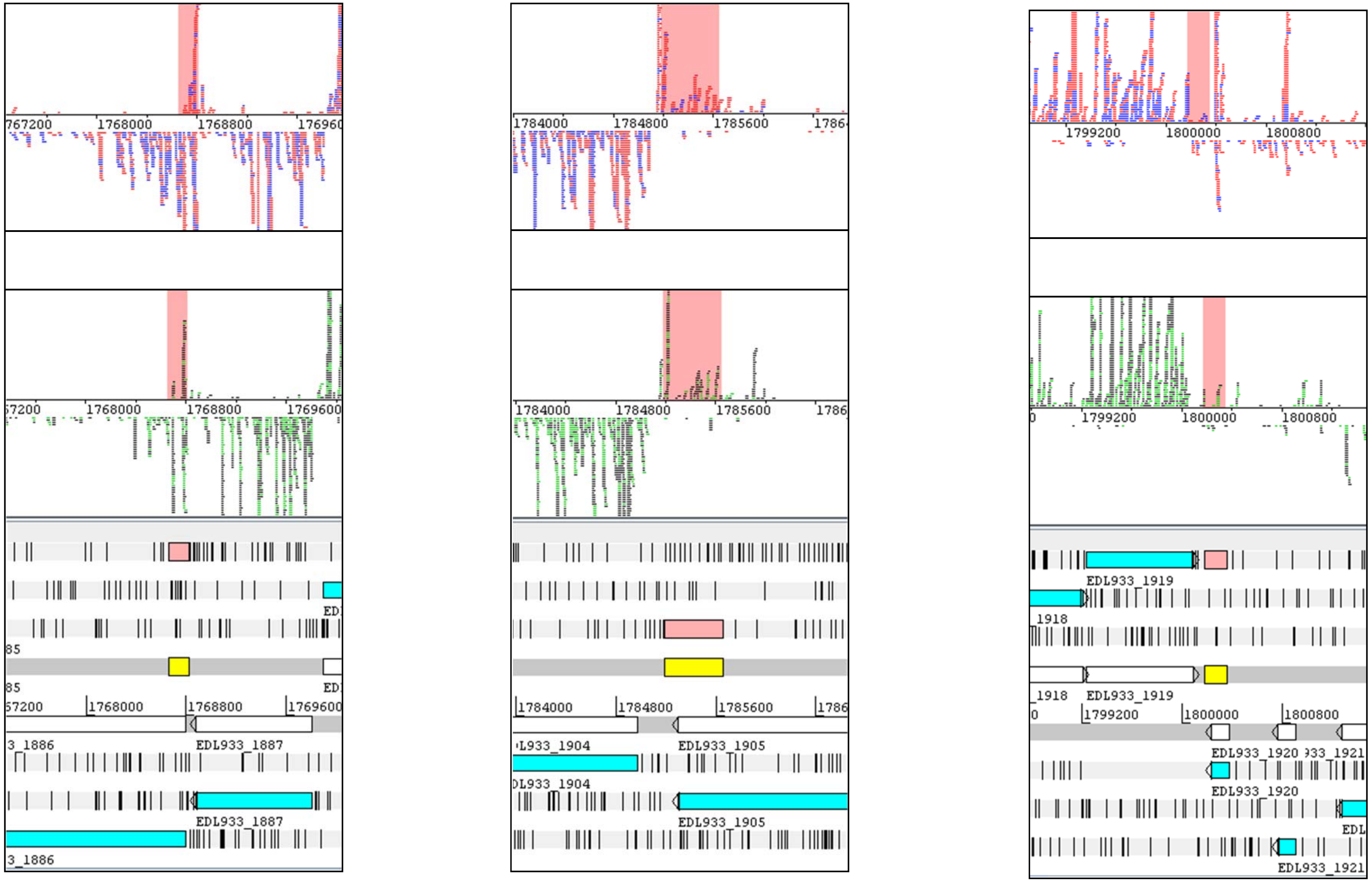

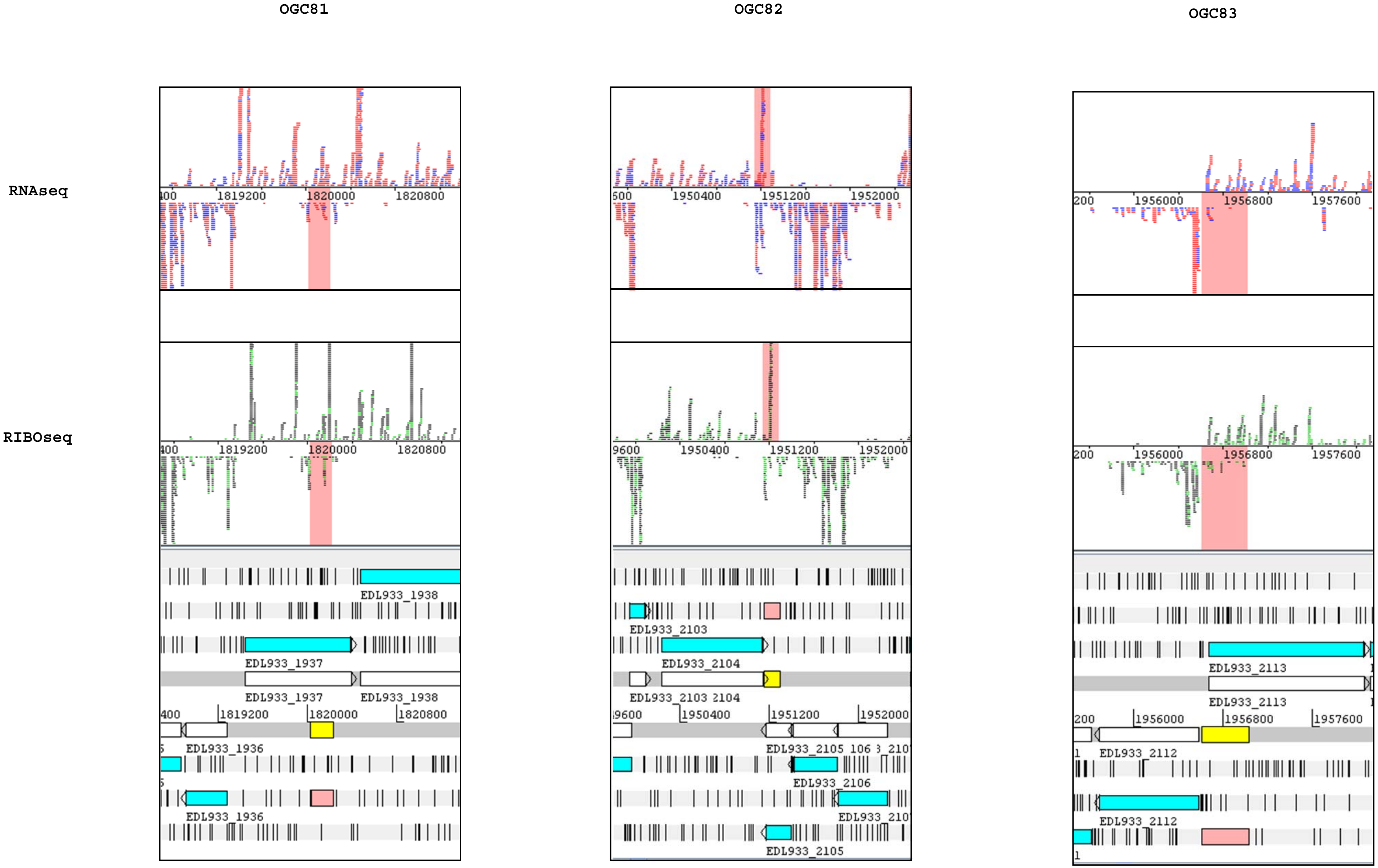
OGC81
OGC82
OGC83
RNAseq
RIBOseq
1819200
1820000
1820800
1950400
1951200
1952000
1956000
1956800
1957600
EDL933_1936
EDL933_1937
EDL933_1938
EDL933_2103
EDL933_2104
EDL933_2105
EDL933_2106
EDL933_2112
EDL933_2113

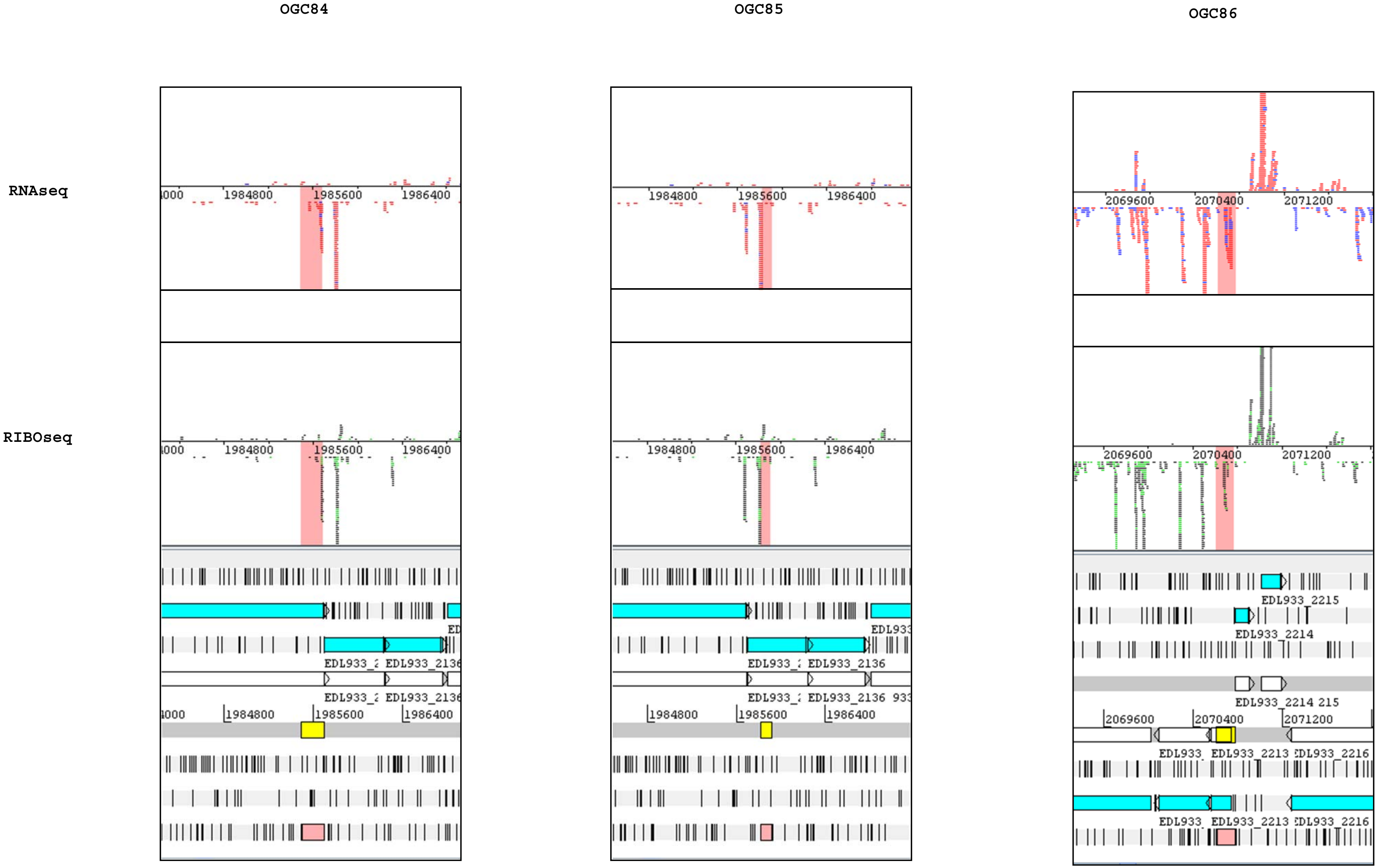
OGC84
OGC85
OGC86
RNAseq
RIBOseq
1984800
1985600
1986400
2069600
2070400
2071200
EDL933_2136
EDL933_2213
EDL933_2214
EDL933_2215
EDL933_2216

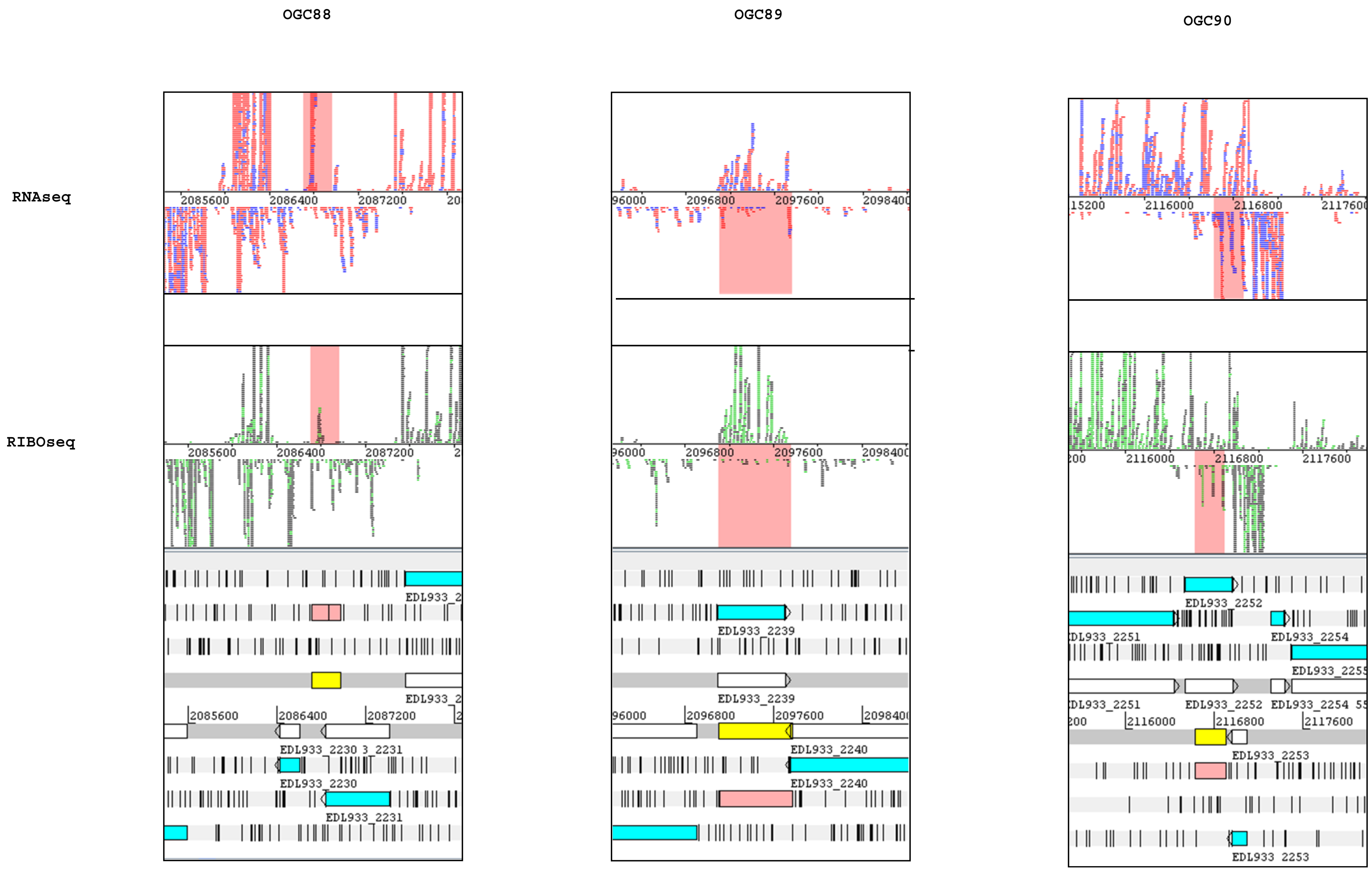
OGC88
OGC89
OGC90
RNAseq
RIBOseq
2085600
2086400
2087200
2096800
2097600
2098400
2116000
2116800
2117600
EDL933_2230
EDL933_2231
EDL933_2239
EDL933_2240
EDL933_2251
EDL933_2252
EDL933_2253
EDL933_2254

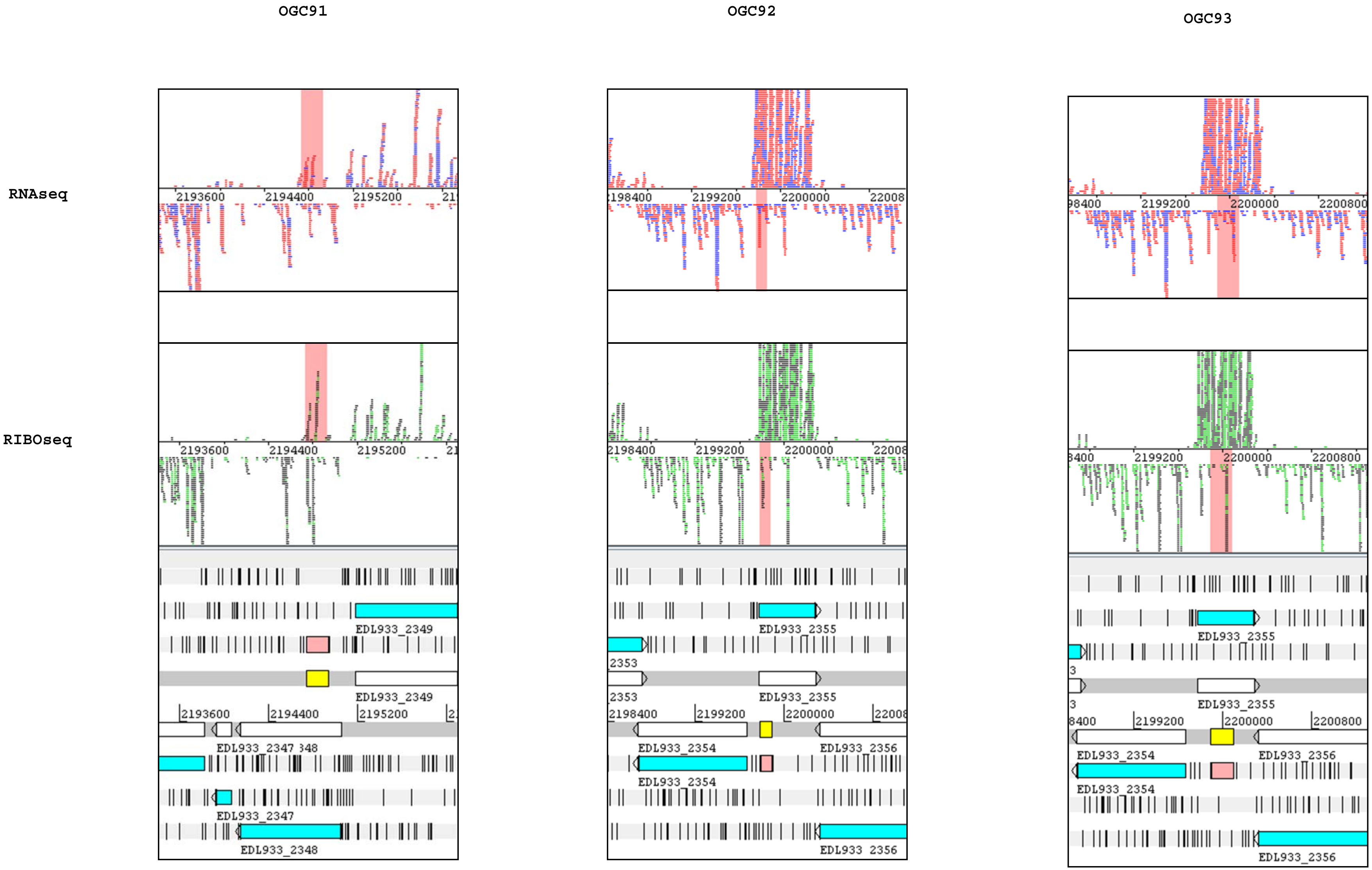
OGC91
OGC92
OGC93
RNAseq
RIBOseq
2193600
2194400
2195200
2198400
2199200
2200000
2200800
EDL933_2347
EDL933_2348
EDL933_2349
EDL933_2353
EDL933_2354
EDL933_2355
EDL933_2356

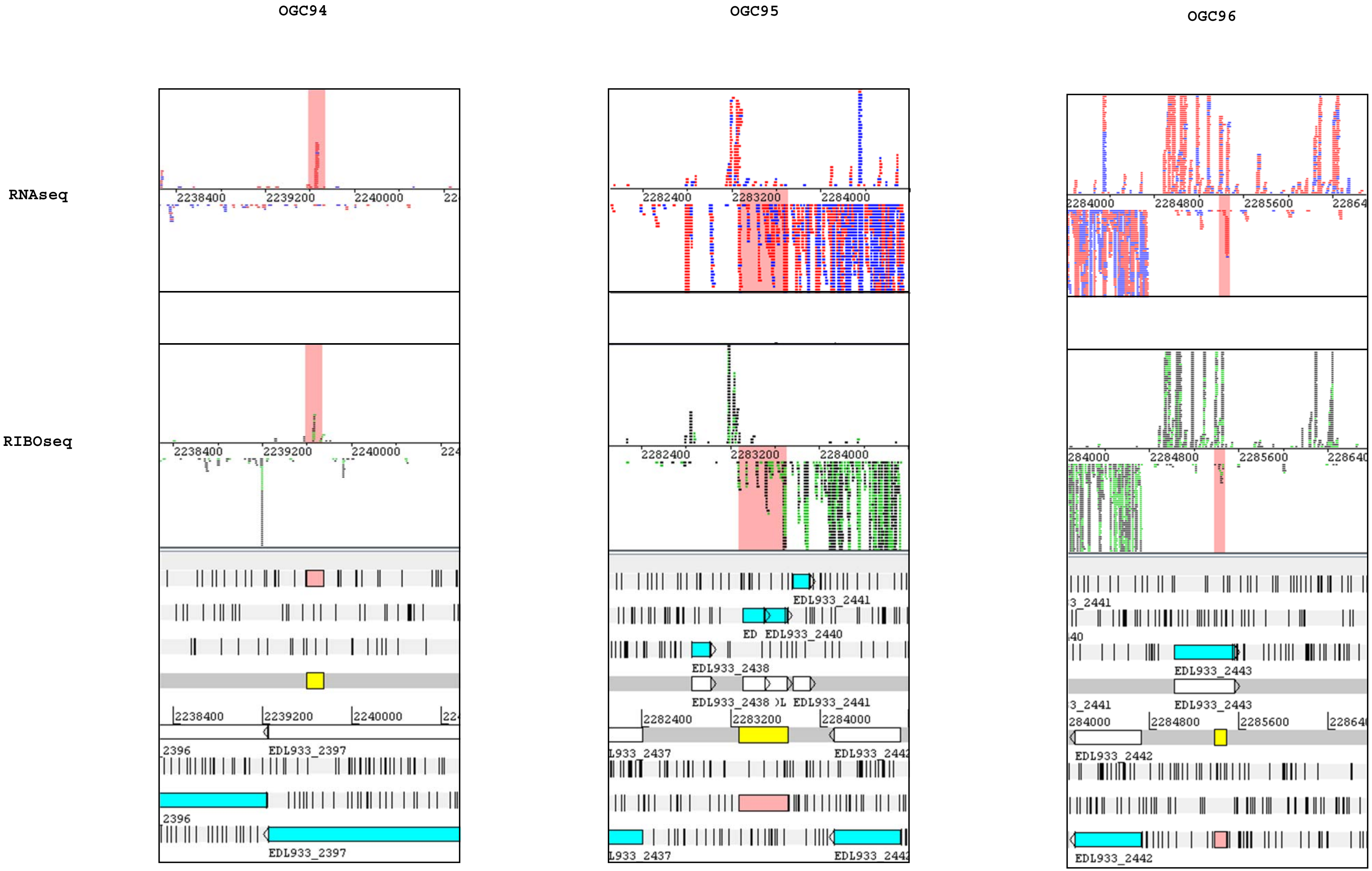
OGC94
OGC95
OGC96
RNAseq
RIBOseq
EDL933_2397
EDL933_2441
EDL933_2440
EDL933_2438
EDL933_2437
EDL933_2443
EDL933_2442

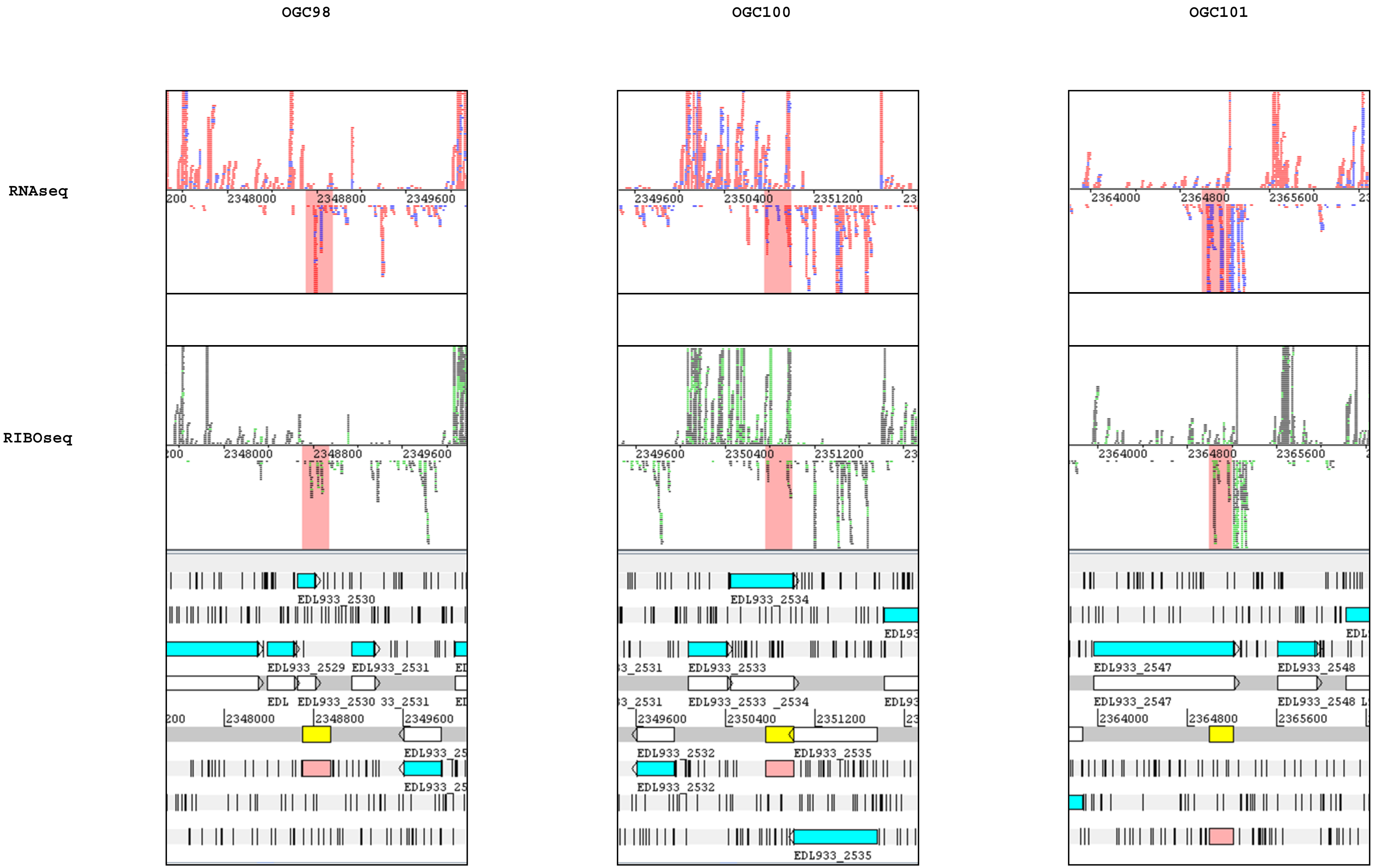
OGC98
OGC100
OGC101
RNAseq
RIBOseq
EDL933_2530
EDL933_2529
EDL933_2531
EDL933_2534
EDL933_2533
EDL933_2532
EDL933_2535
EDL933_2547
EDL933_2548

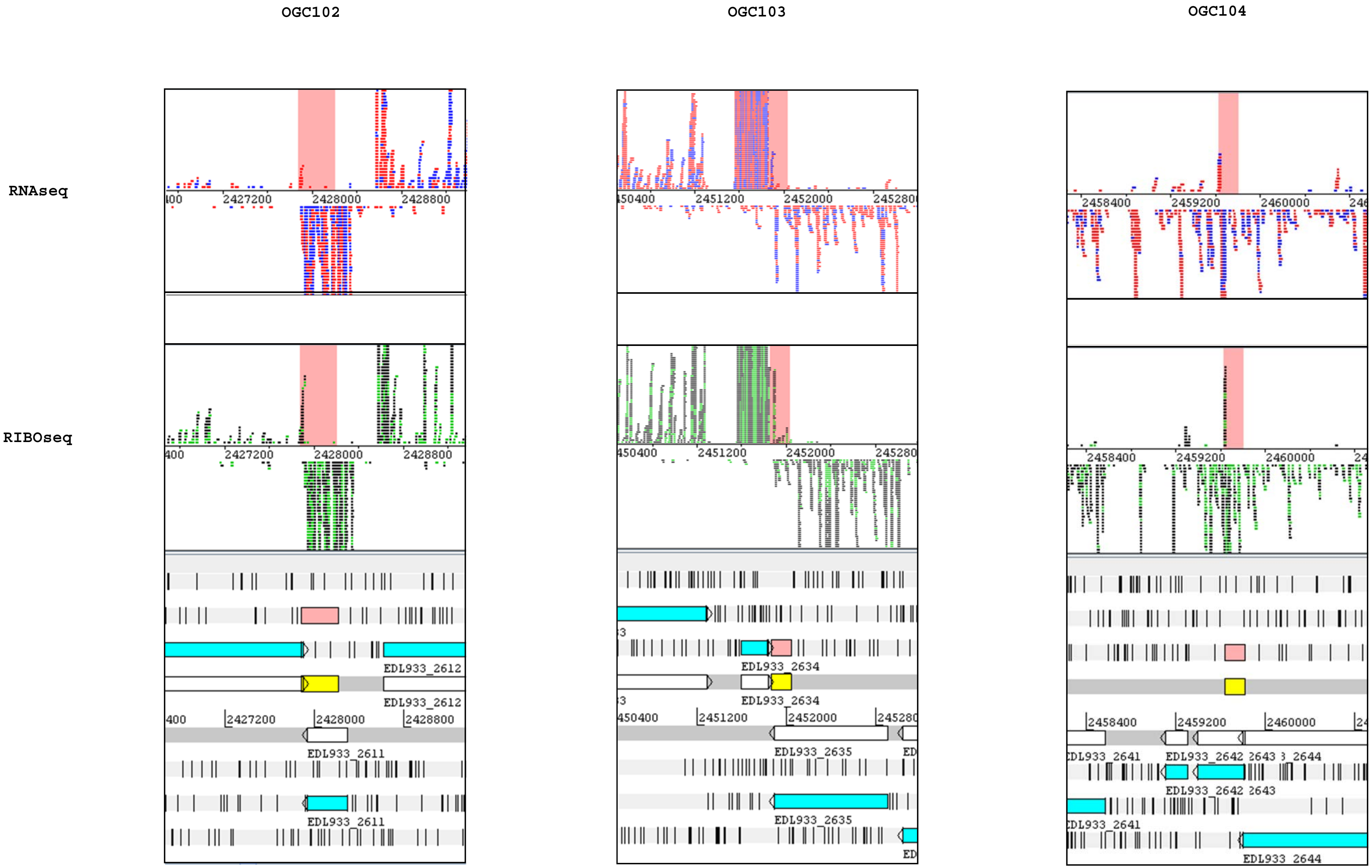

OGC102
OGC103
OGC104
RNAseq
RIBOseq
2427200
2428000
2428800
2451200
2452000
2458400
2459200
2460000
EDL933_2612
EDL933_2611
EDL933_2634
EDL933_2635
EDL933_2642
2643
EDL933_2644

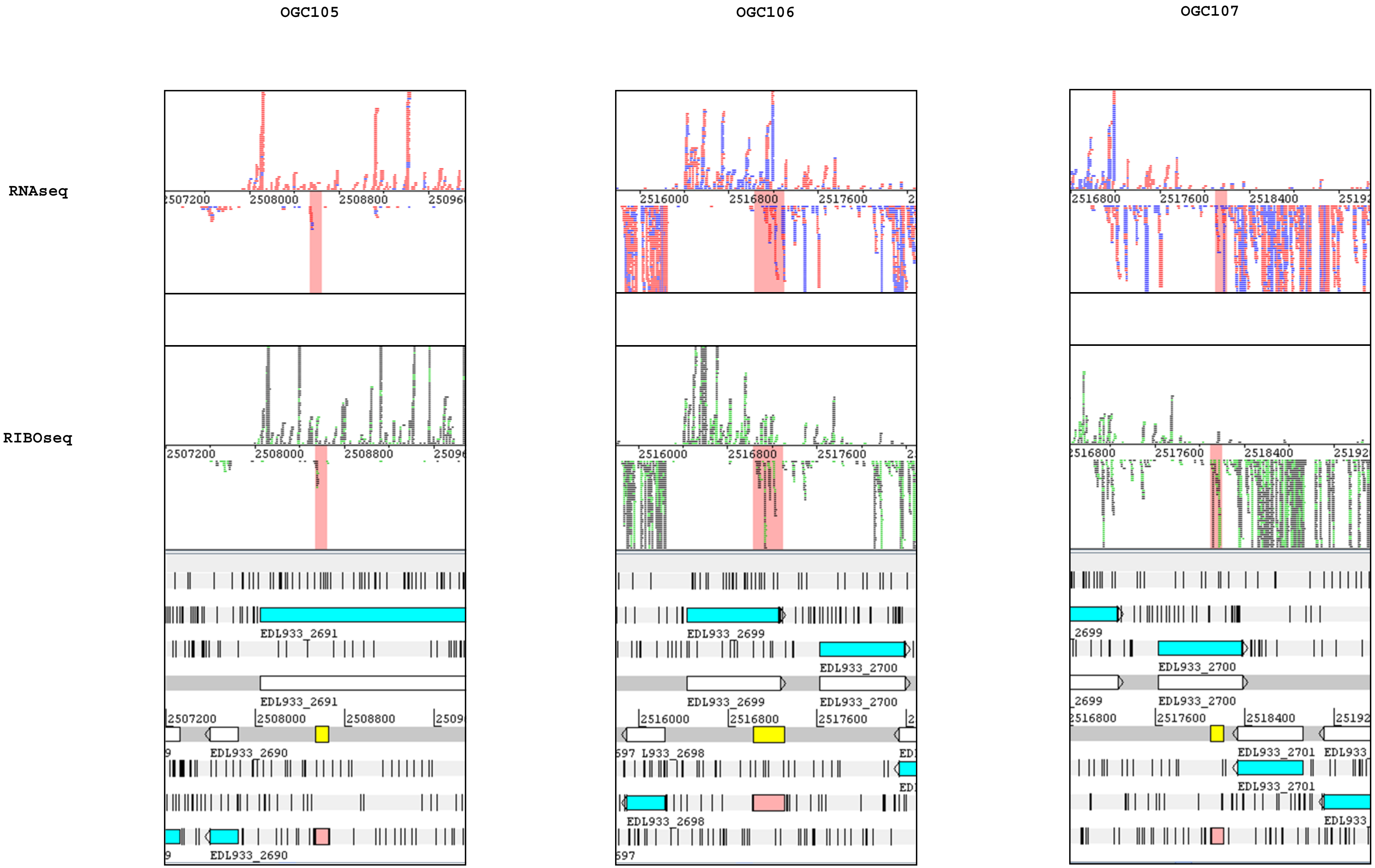
OGC105
OGC106
OGC107
RNAseq
RIBOseq
EDL933_2691
EDL933_2690
EDL933_2699
EDL933_2700
EDL933_2698
EDL933_2701

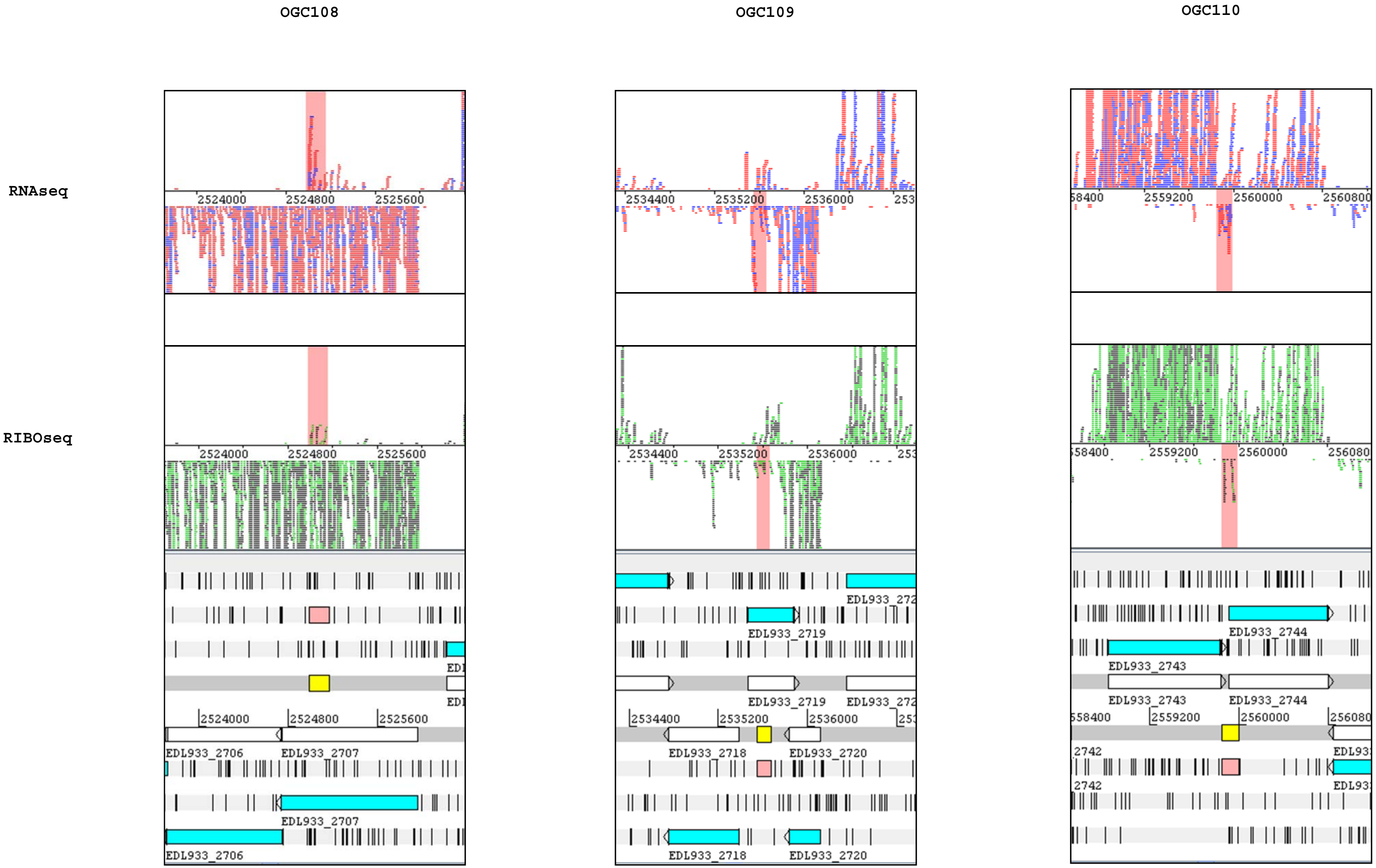
OGC108
OGC109
OGC110
RNAseq
RIBOseq
2524000
2524800
2525600
EDL933_2706
EDL933_2707
2534400
2535200
2536000
EDL933_2718
EDL933_2719
EDL933_2720
2559200
2560000
EDL933_2743
EDL933_2744

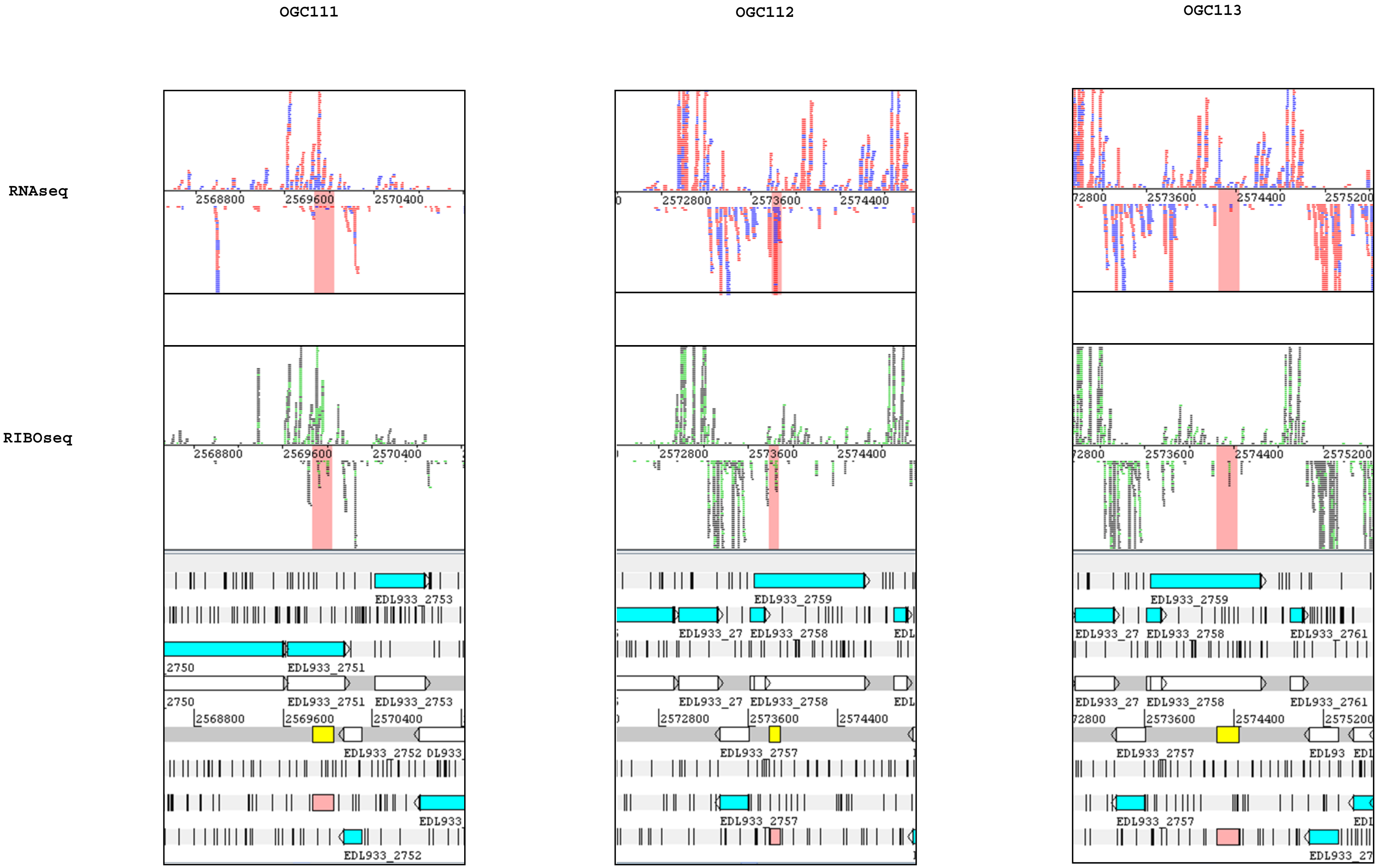
OGC111
OGC112
OGC113
RNAseq
RIBOseq
2568800
2569600
2570400
2572800
2573600
2574400
2575200
EDL933_2753
2750
EDL933_2751
EDL933_2752
EDL933_2759
EDL933_2758
EDL933_2757
EDL933_2761

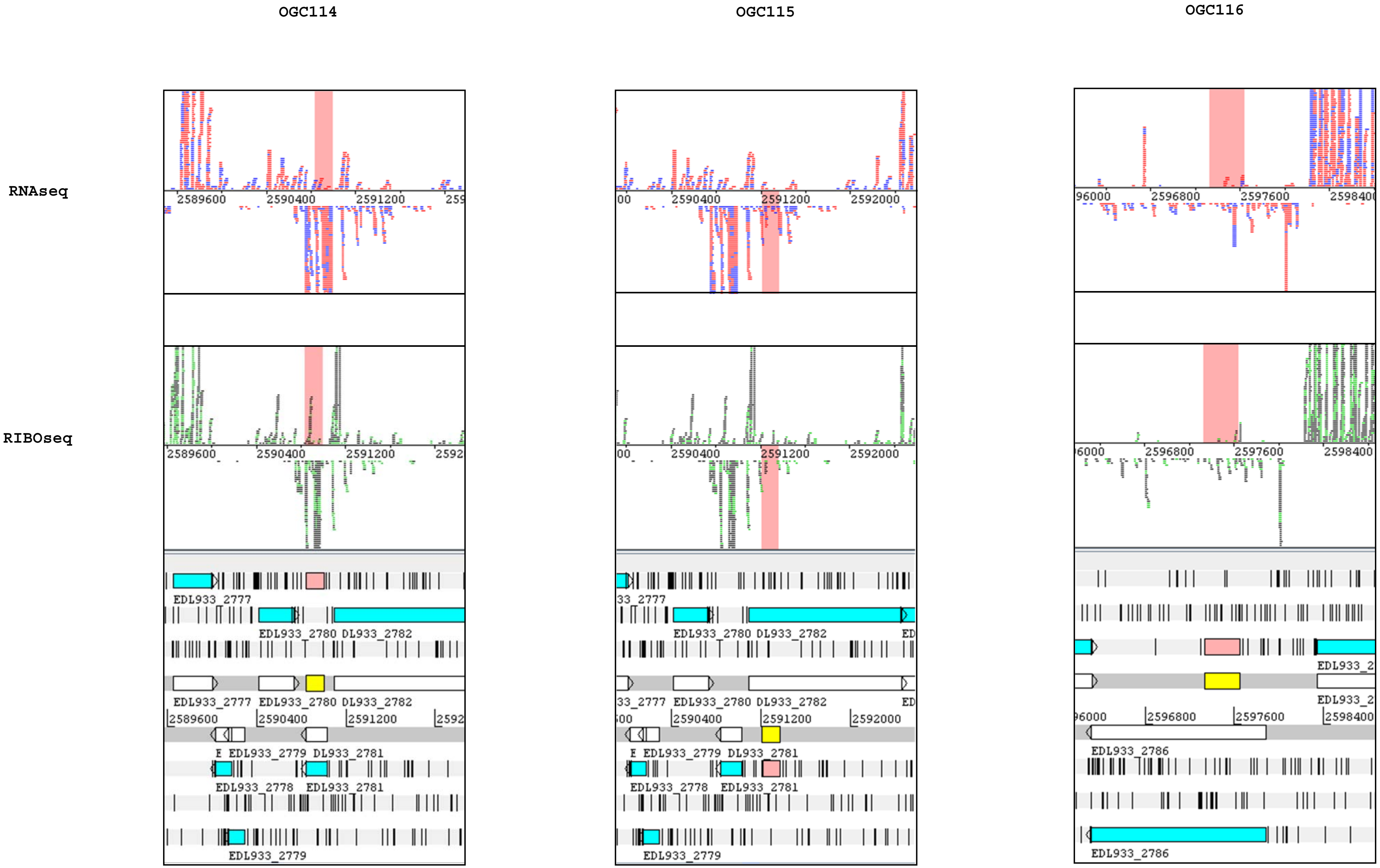
OGC114
OGC115
OGC116
RNAseq
RIBOseq
EDL933_2777
EDL933_2780
DL933_2782
EDL933_2779
DL933_2781
EDL933_2778
EDL933_2781
EDL933_2786
2589600
2590400
2591200
2592000
2596800
2597600
2598400

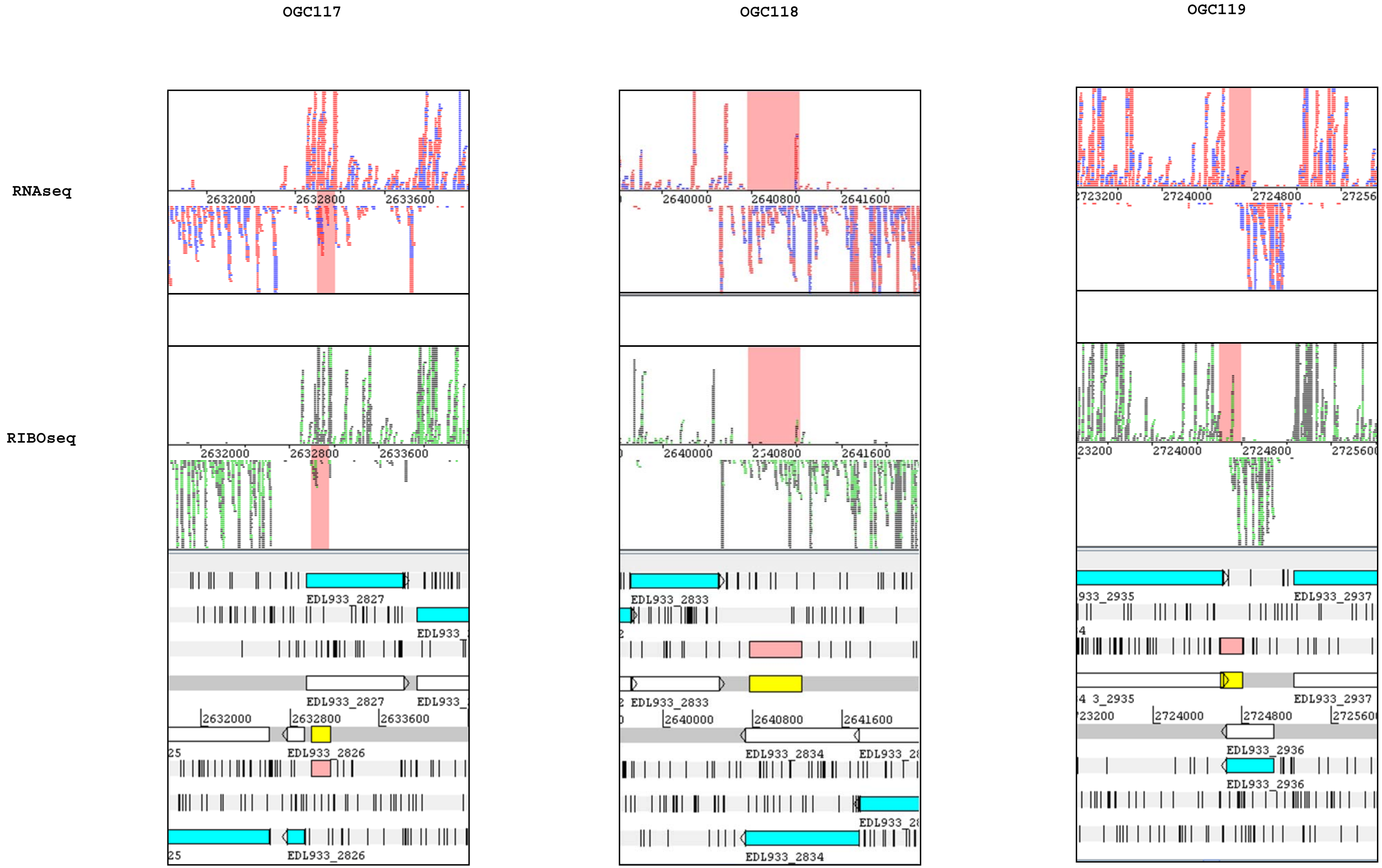
OGC117
OGC118
OGC119
RNAseq
RIBOseq
2632000
2632800
2633600
2640000
2640800
2641600
2724000
2724800
EDL933_2827
EDL933_2826
EDL933_2833
EDL933_2834
EDL933_2935
EDL933_2936
EDL933_2937

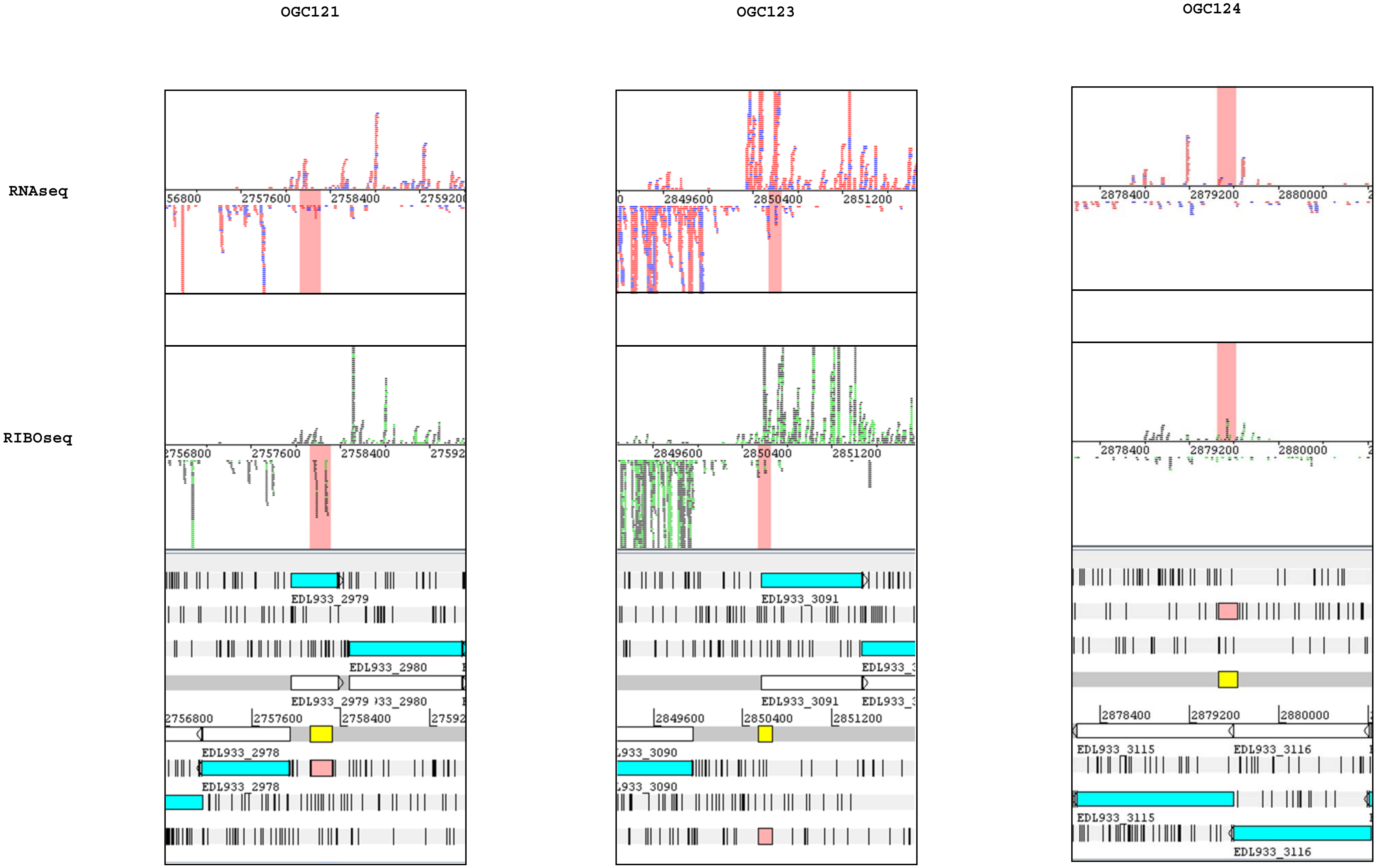
OGC121
OGC123
OGC124
RNAseq
RIBOseq
EDL933_2979
EDL933_2980
EDL933_2978
EDL933_3091
EDL933_3090
EDL933_3115
EDL933_3116
2757600
2758400
2849600
2850400
2851200
2878400
2879200
2880000

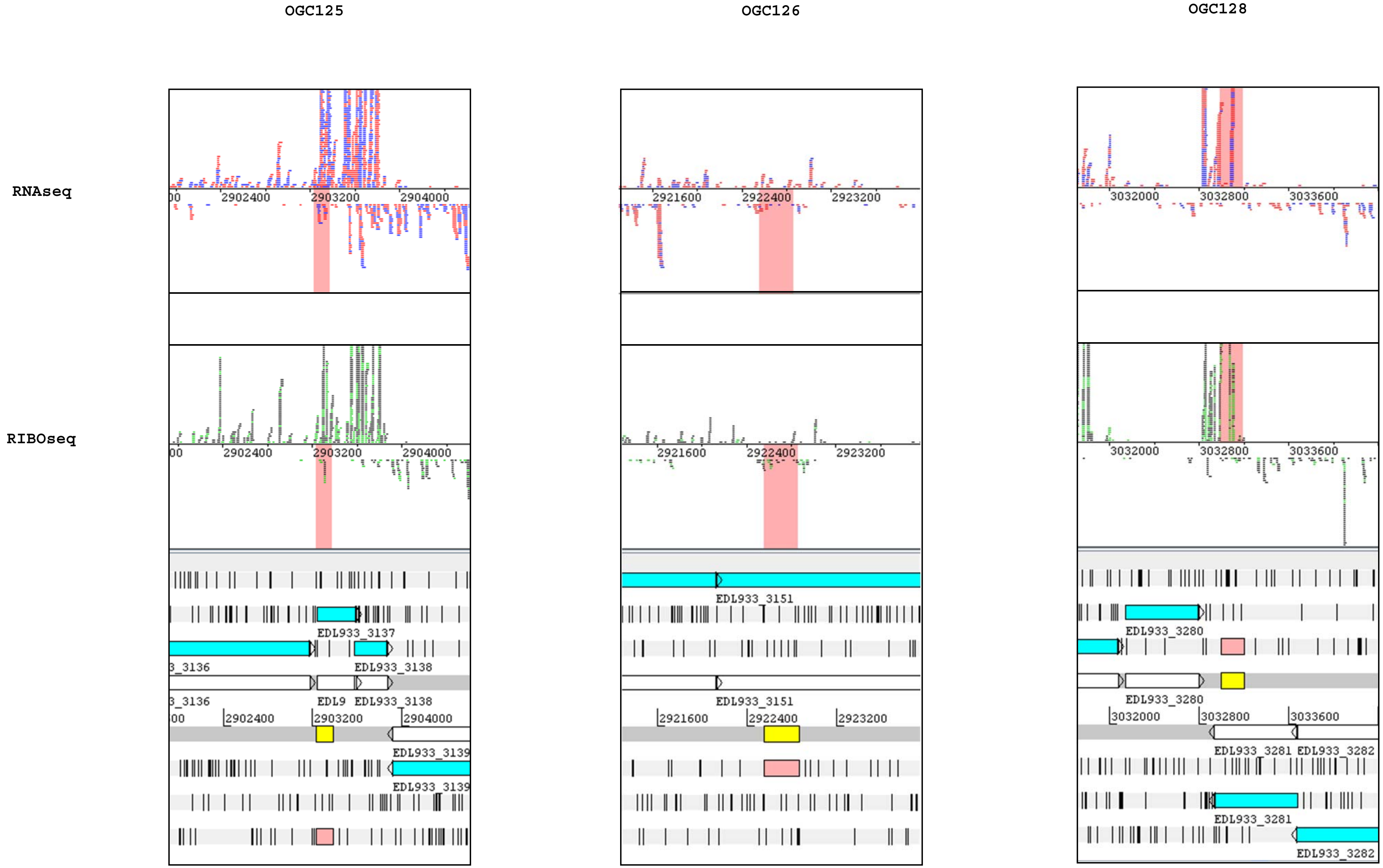
OGC125
OGC126
OGC128
RNAseq
RIBOseq
2902400
2903200
2904000
2921600
2922400
2923200
3032000
3032800
3033600
EDL933_3137
EDL933_3138
EDL933_3139
EDL933_3151
EDL933_3280
EDL933_3281
EDL933_3282

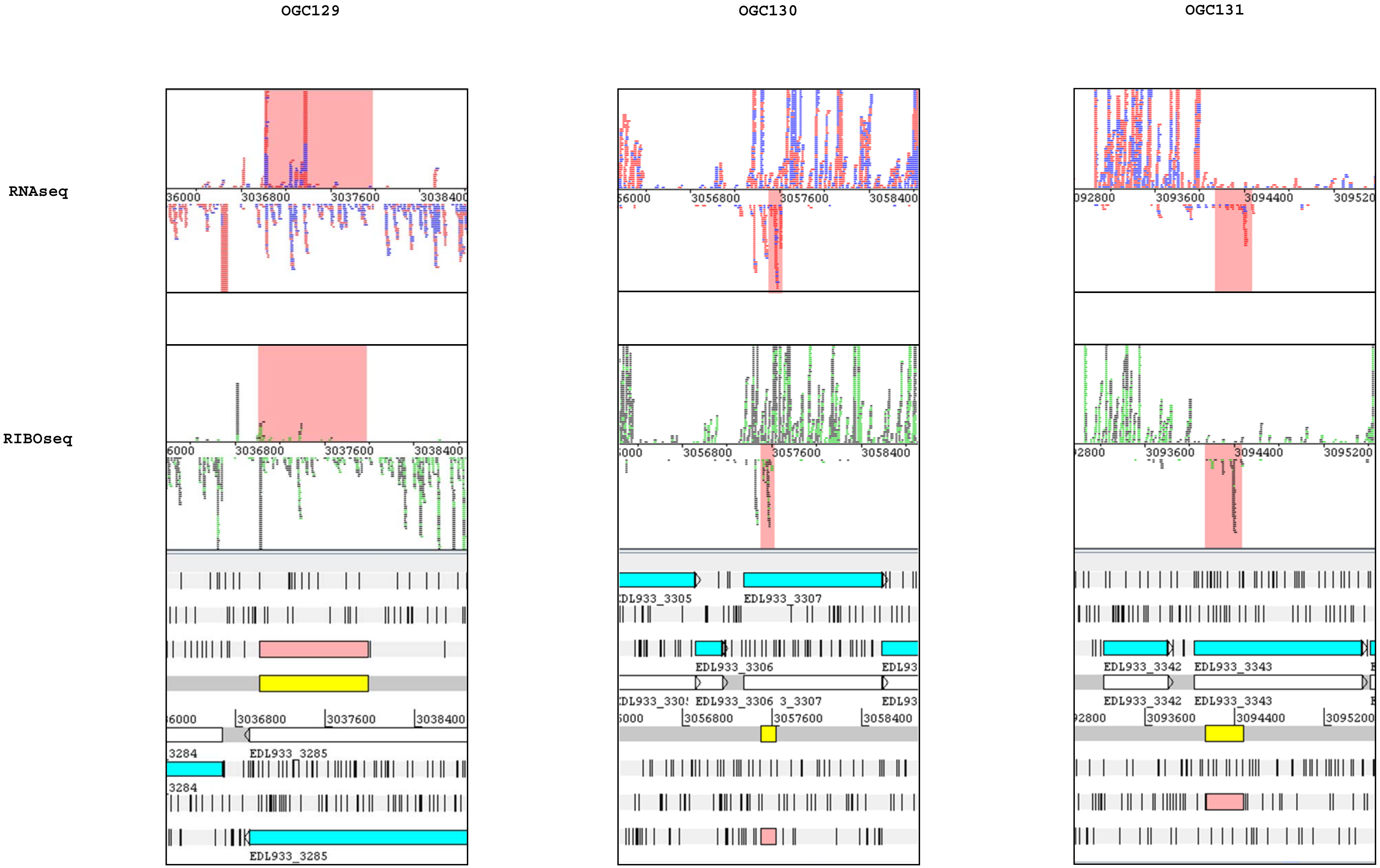
OGC129
OGC130
OGC131
RNAseq
RIBOseq
3036800
3037600
EDL933_3285
3056800
3057600
3058400
EDL933_3305
EDL933_3307
EDL933_3306
3093600
3094400
EDL933_3342
EDL933_3343

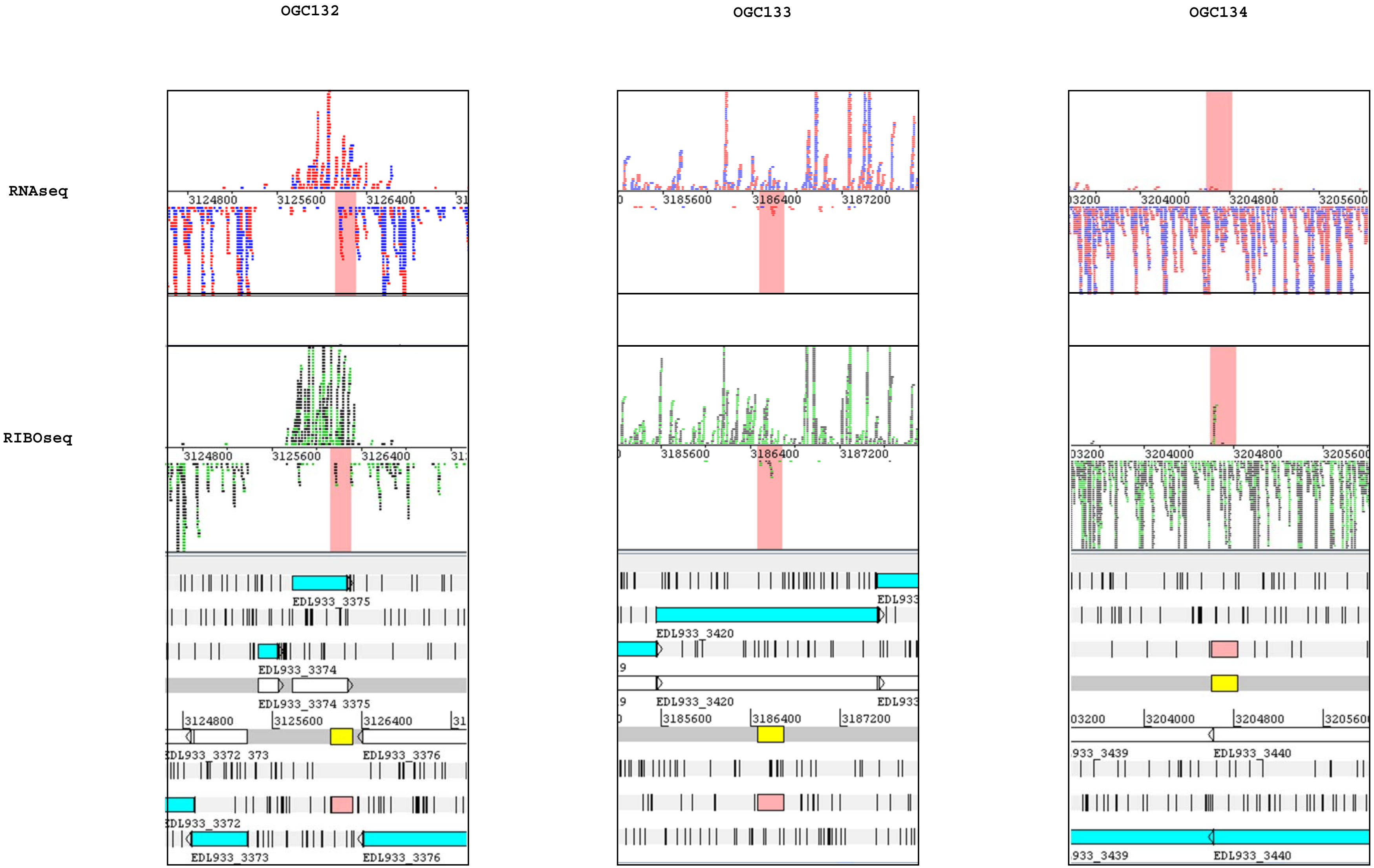
OGC132
OGC133
OGC134
RNAseq
RIBOseq
3124800
3125600
3126400
3185600
3186400
3187200
3204000
3204800
3205600
EDL933_3375
EDL933_3374
EDL933_3374 3375
EDL933_3376
EDL933_3373
EDL933_3420
EDL933_3440

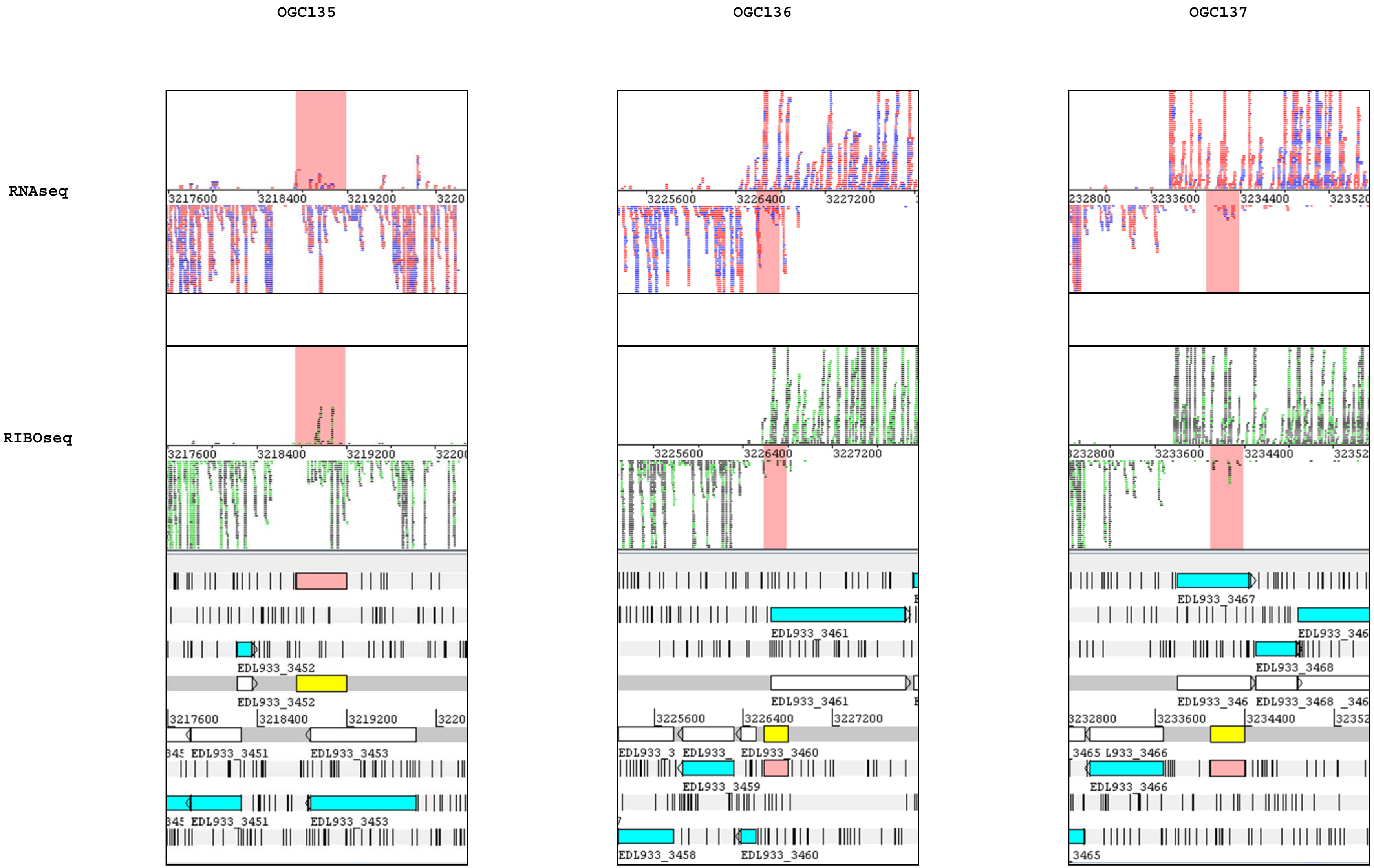
OGC135
OGC136
OGC137
RNAseq
RIBOseq
EDL933_3451
EDL933_3452
EDL933_3453
EDL933_3458
EDL933_3459
EDL933_3460
EDL933_3461
EDL933_3465
EDL933_3466
EDL933_3467
EDL933_3468

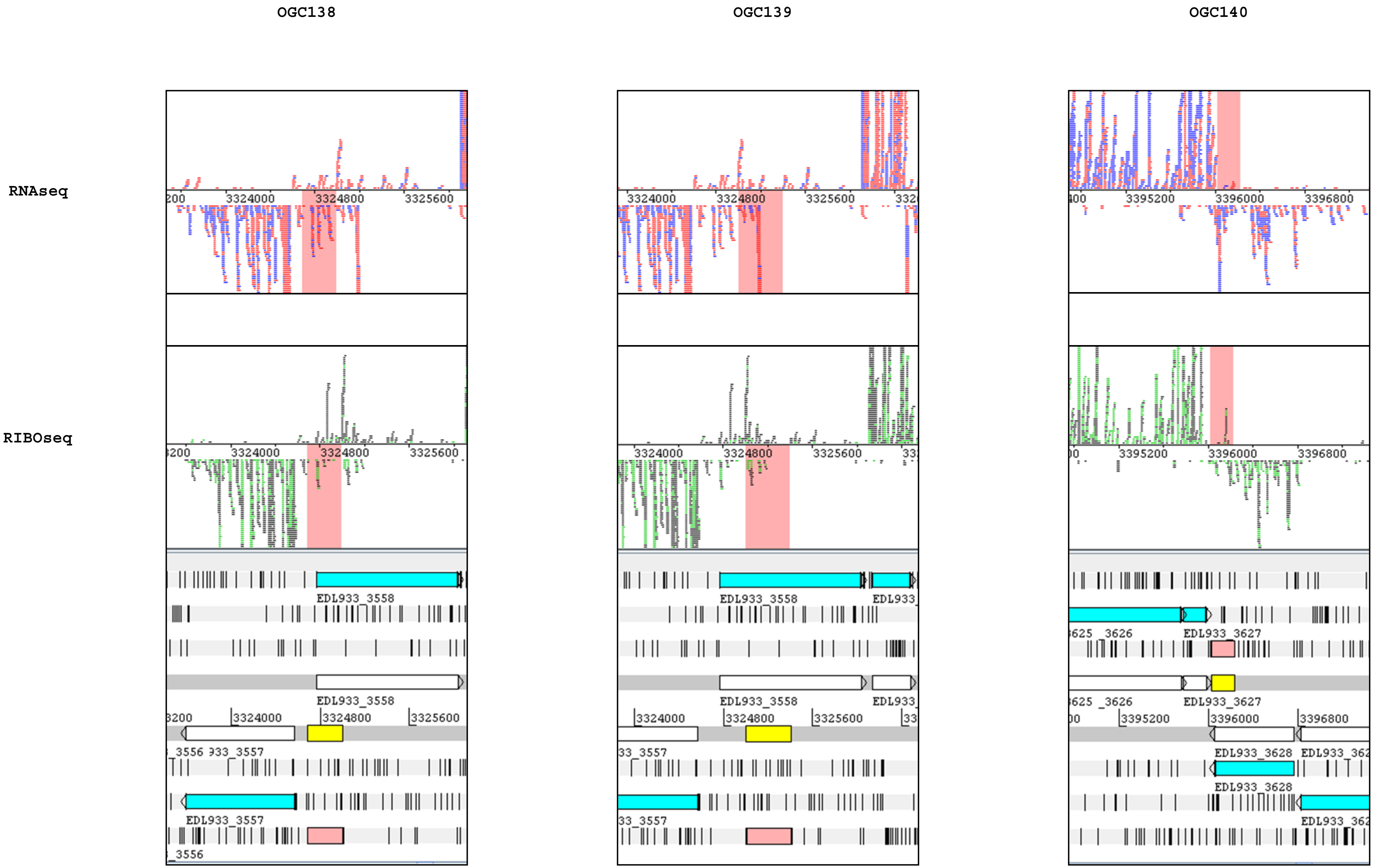

OGC138
OGC139
OGC140
RNAseq
RIBOseq
3324000
3324800
3325600
3395200
3396000
3396800
EDL933_3558
EDL933_3557
EDL933_3627
EDL933_3628

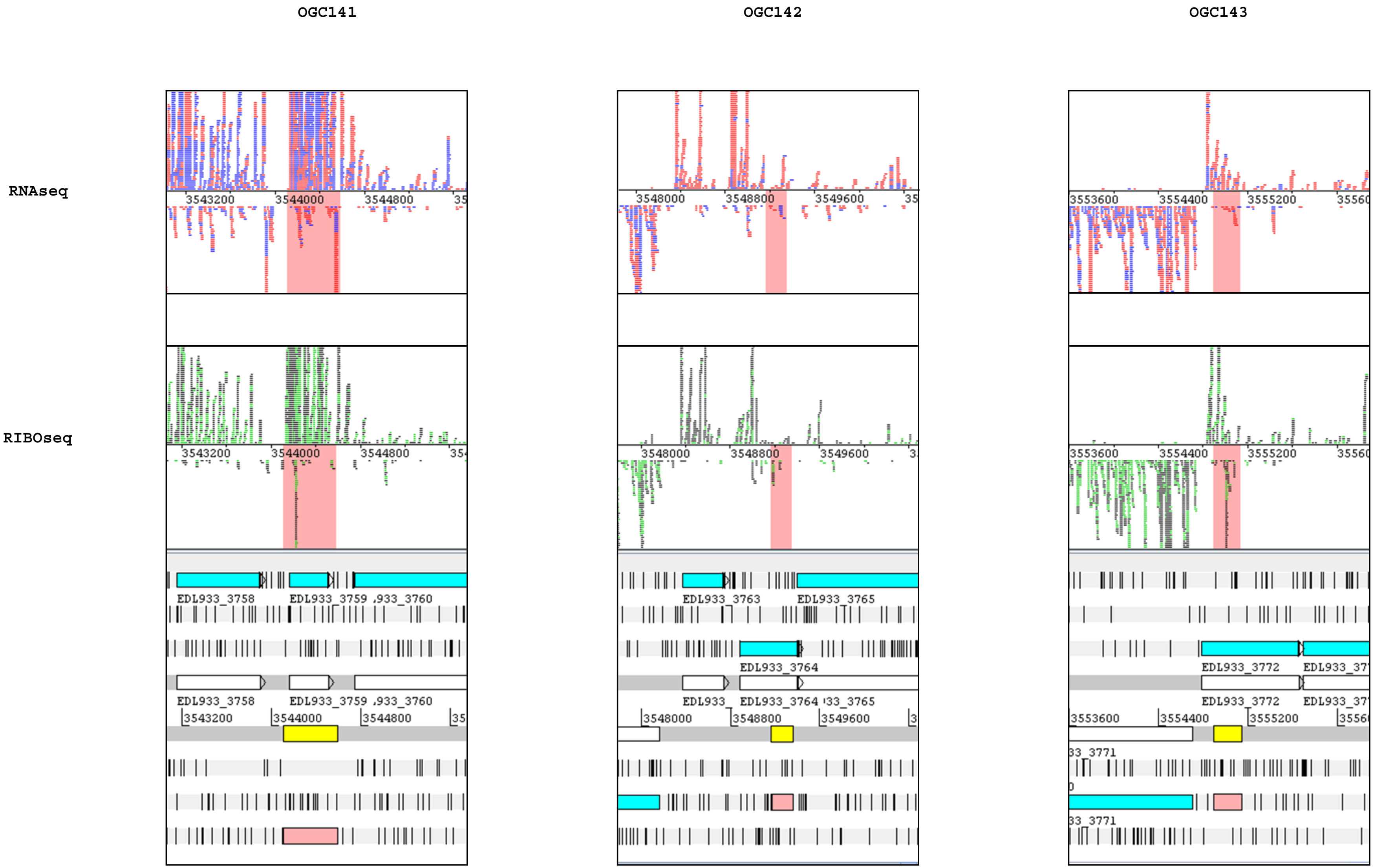
OGC141
OGC142
OGC143
RNAseq
RIBOseq
3543200
3544000
3544800
3548000
3548800
3549600
3553600
3554400
3555200
EDL933_3758
EDL933_3759
EDL933_3763
EDL933_3764
EDL933_3765
EDL933_3772

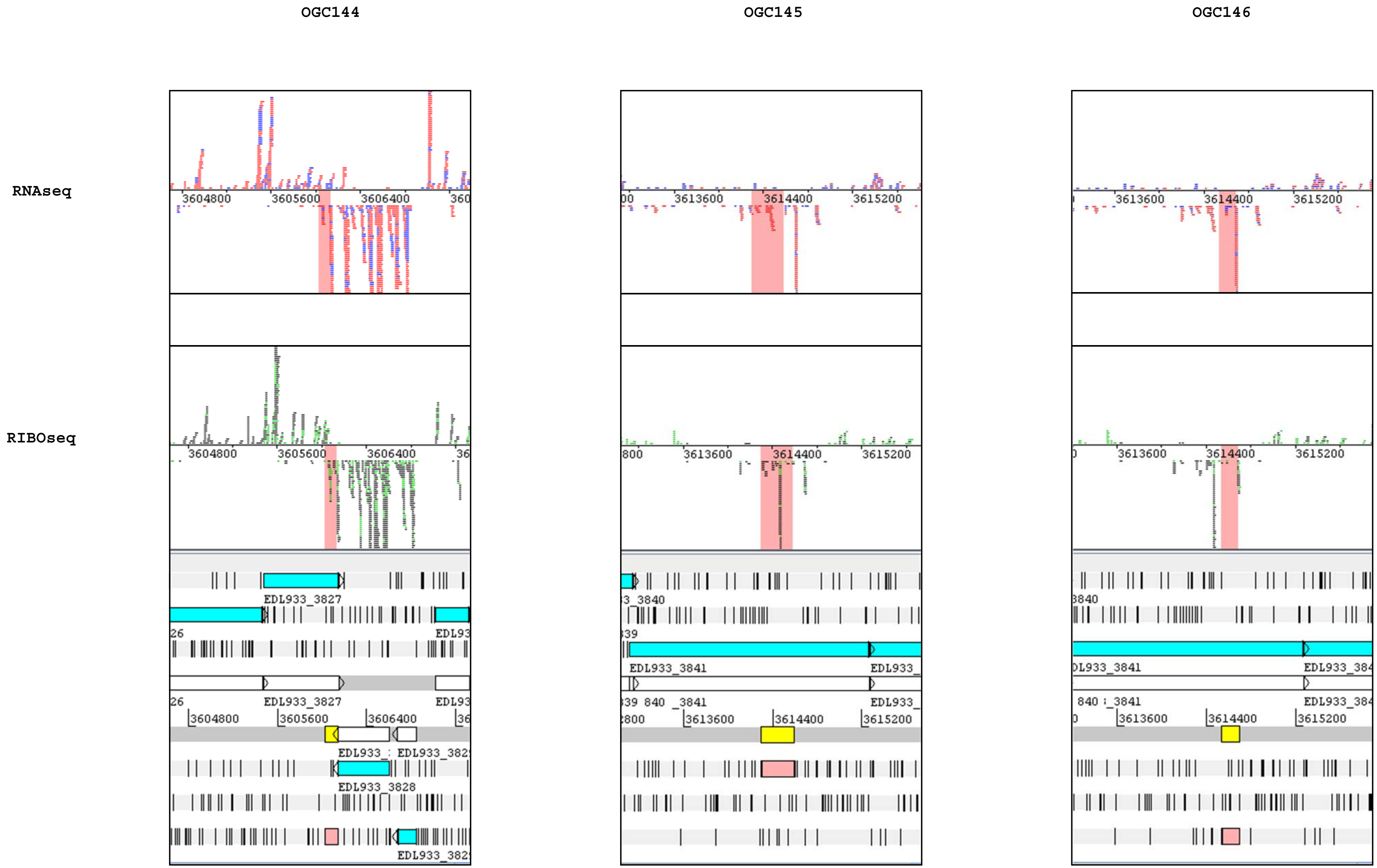
OGC144
OGC145
OGC146
RNAseq
RIBOseq
3604800
3605600
3606400
3613600
3614400
3615200
EDL933_3827
EDL933_3828
EDL933_3841

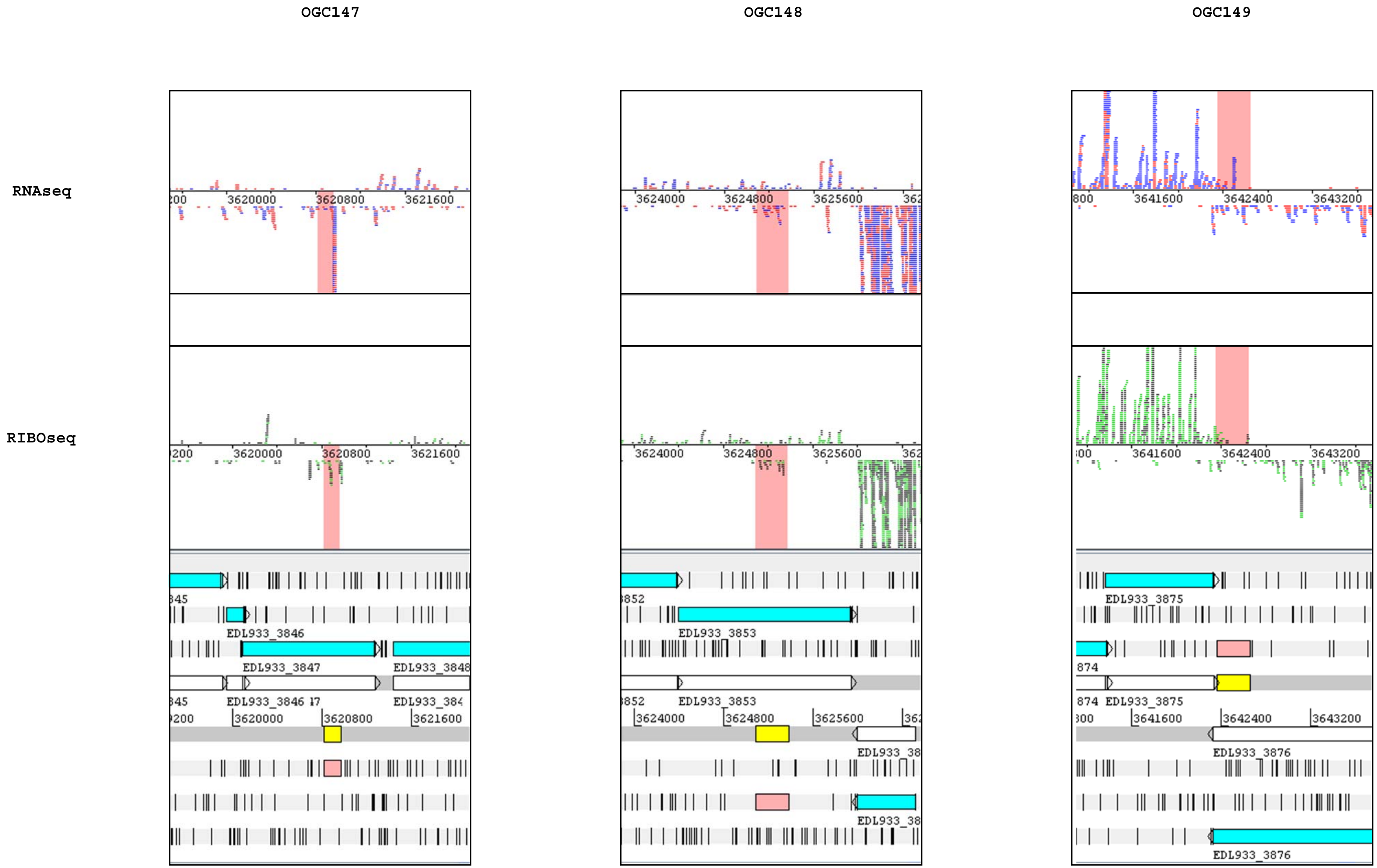
OGC147
OGC148
OGC149
RNAseq
RIBOseq
3620000
3620800
3621600
3624000
3624800
3625600
3641600
3642400
3643200
EDL933_3846
EDL933_3847
EDL933_3853
EDL933_3875
EDL933_3876

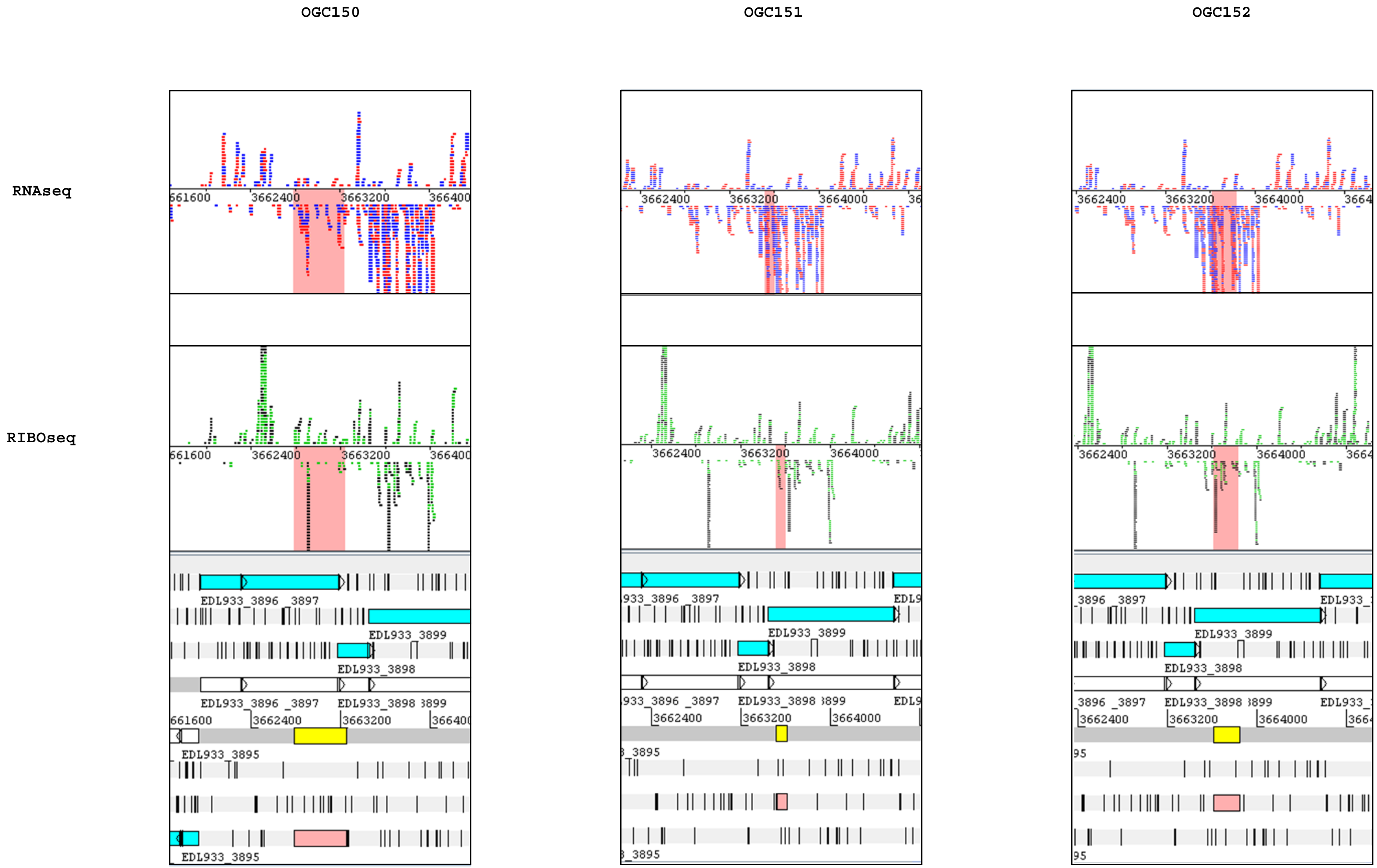
OGC150
OGC151
OGC152
RNAseq
RIBOseq
EDL933_3895
EDL933_3896
EDL933_3898
EDL933_3899
3662400
3663200
3664000

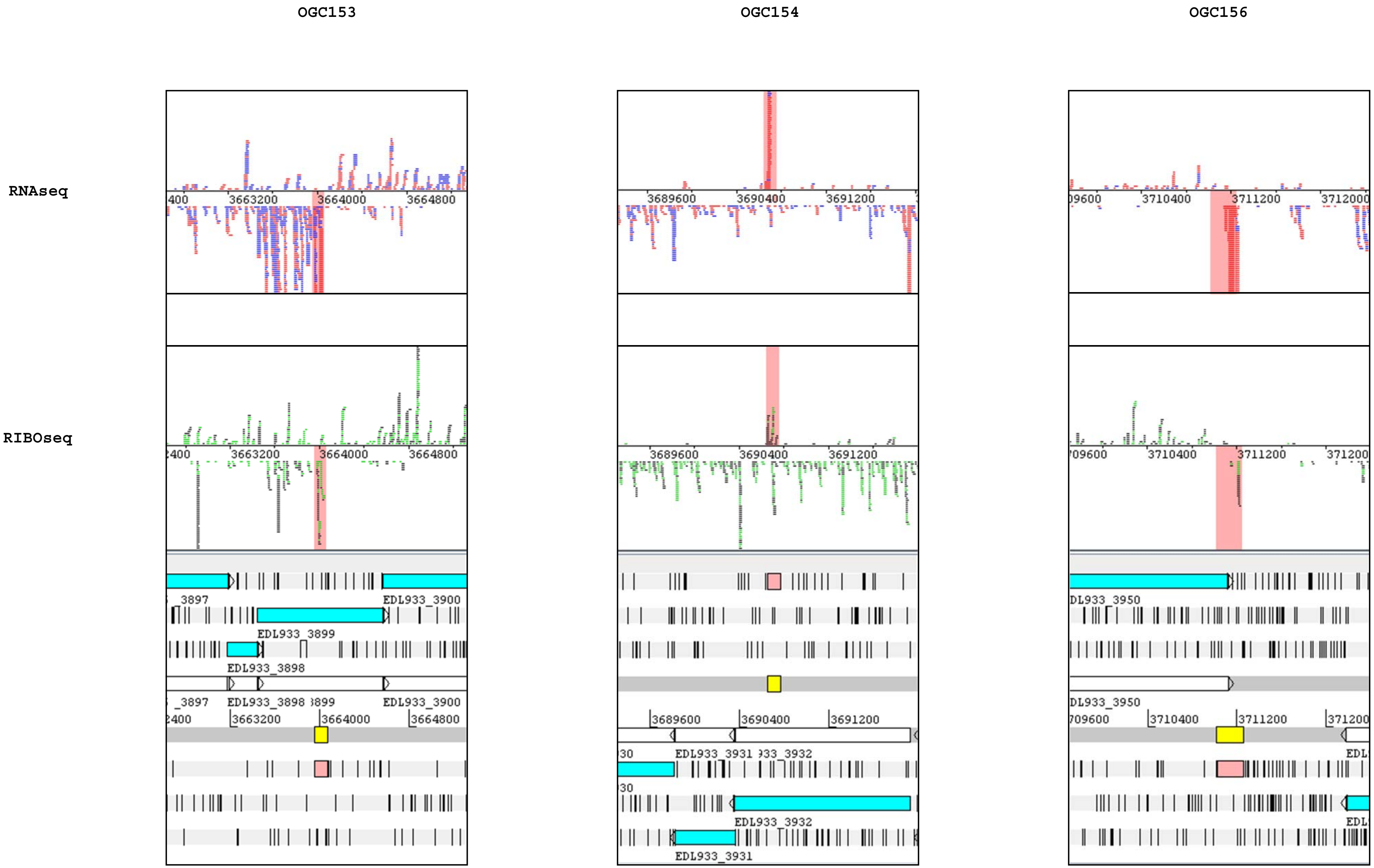
OGC153
OGC154
OGC156
RNAseq
RIBOseq
EDL933_3899
EDL933_3898
EDL933_3900
EDL933_3932
EDL933_3931
DL933_3950

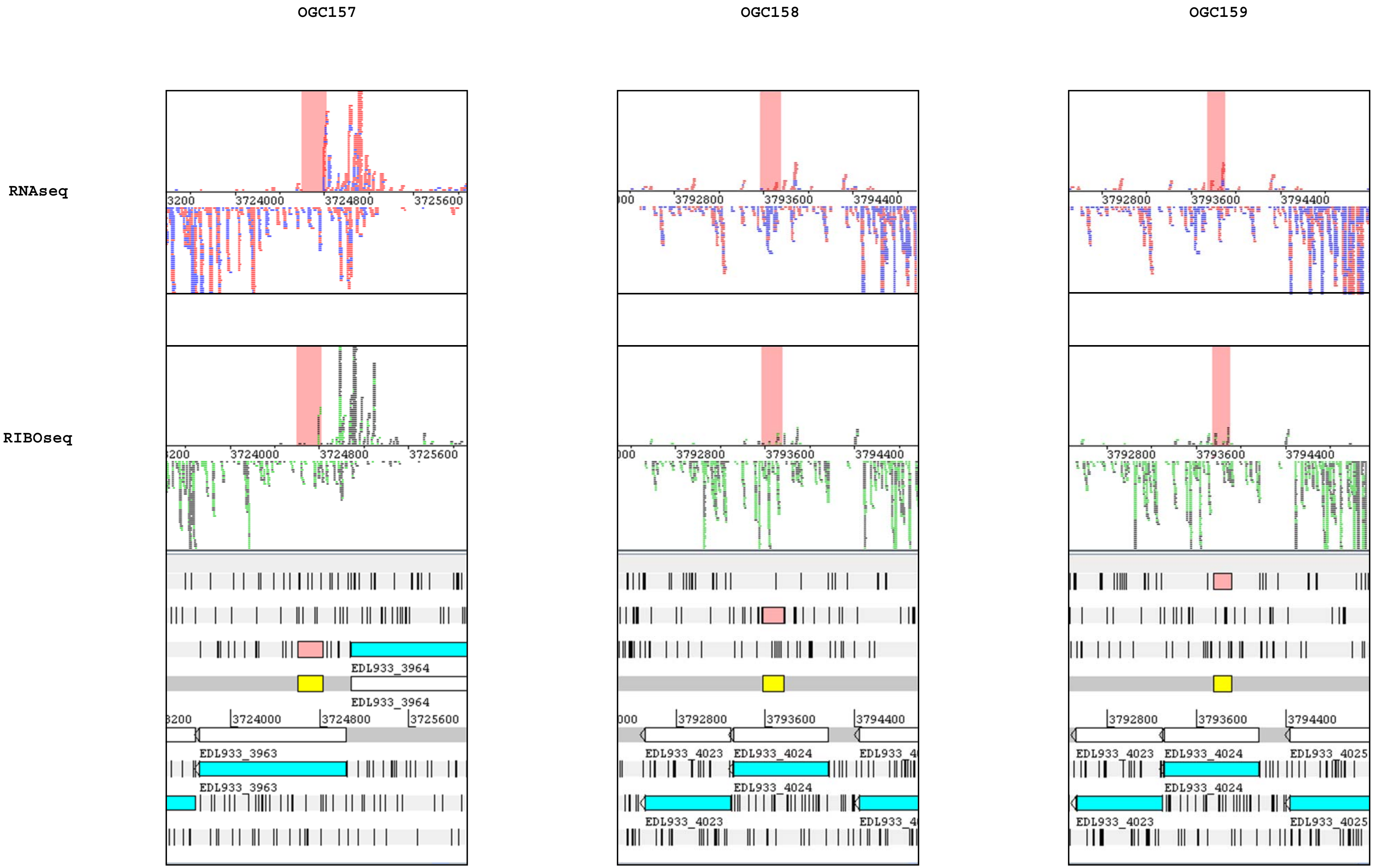
OGC157
OGC158
OGC159
RNAseq
RIBOseq
3724000
3724800
3725600
3792800
3793600
3794400
EDL933_3964
EDL933_3963
EDL933_4023
EDL933_4024
EDL933_4025

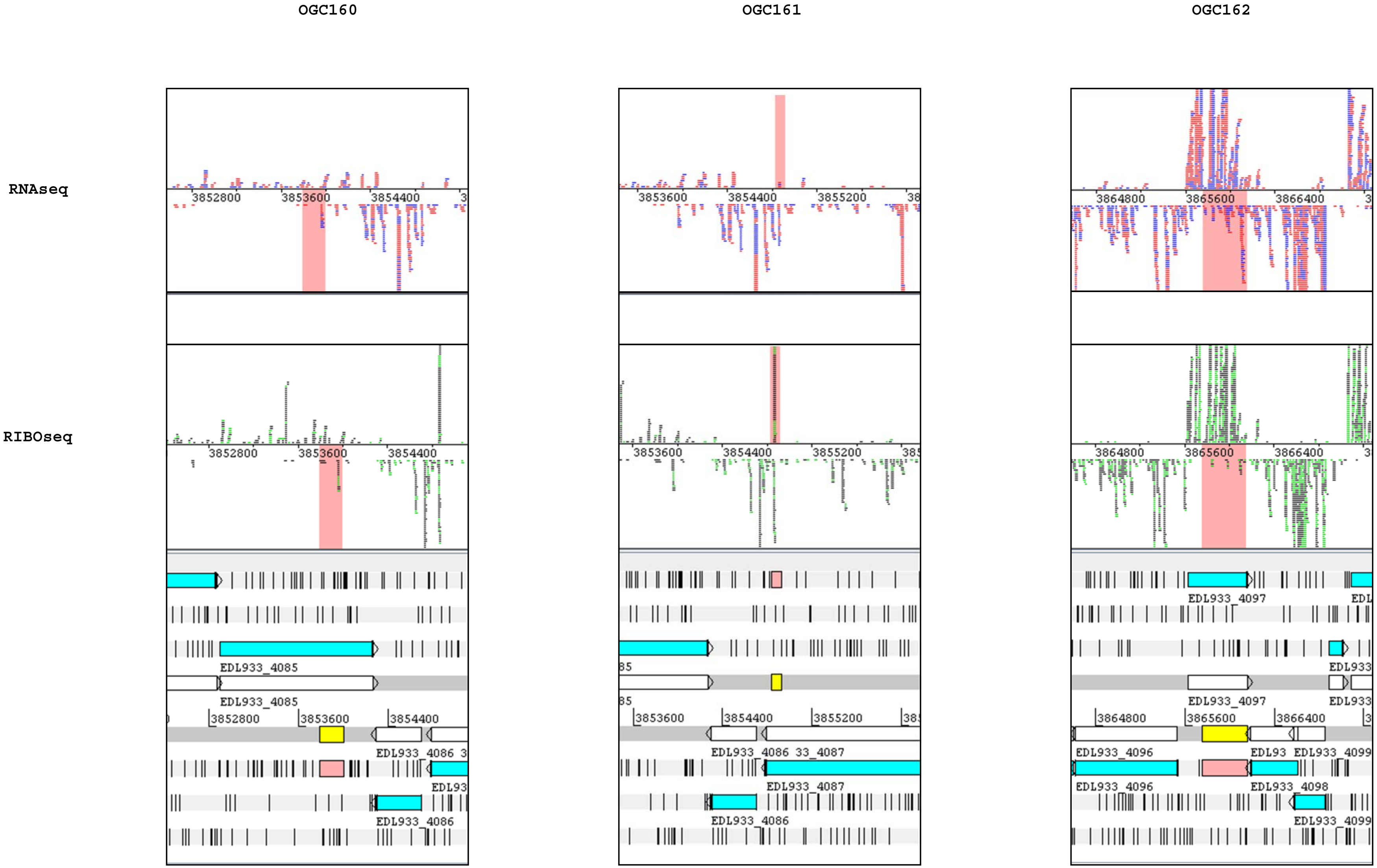

OGC160
OGC161
OGC162
RNAseq
RIBOseq
3852800
3853600
3854400
3855200
3864800
3865600
3866400
EDL933_4085
EDL933_4086
EDL933_4087
EDL933_4096
EDL933_4097
EDL933_4098
EDL933_4099

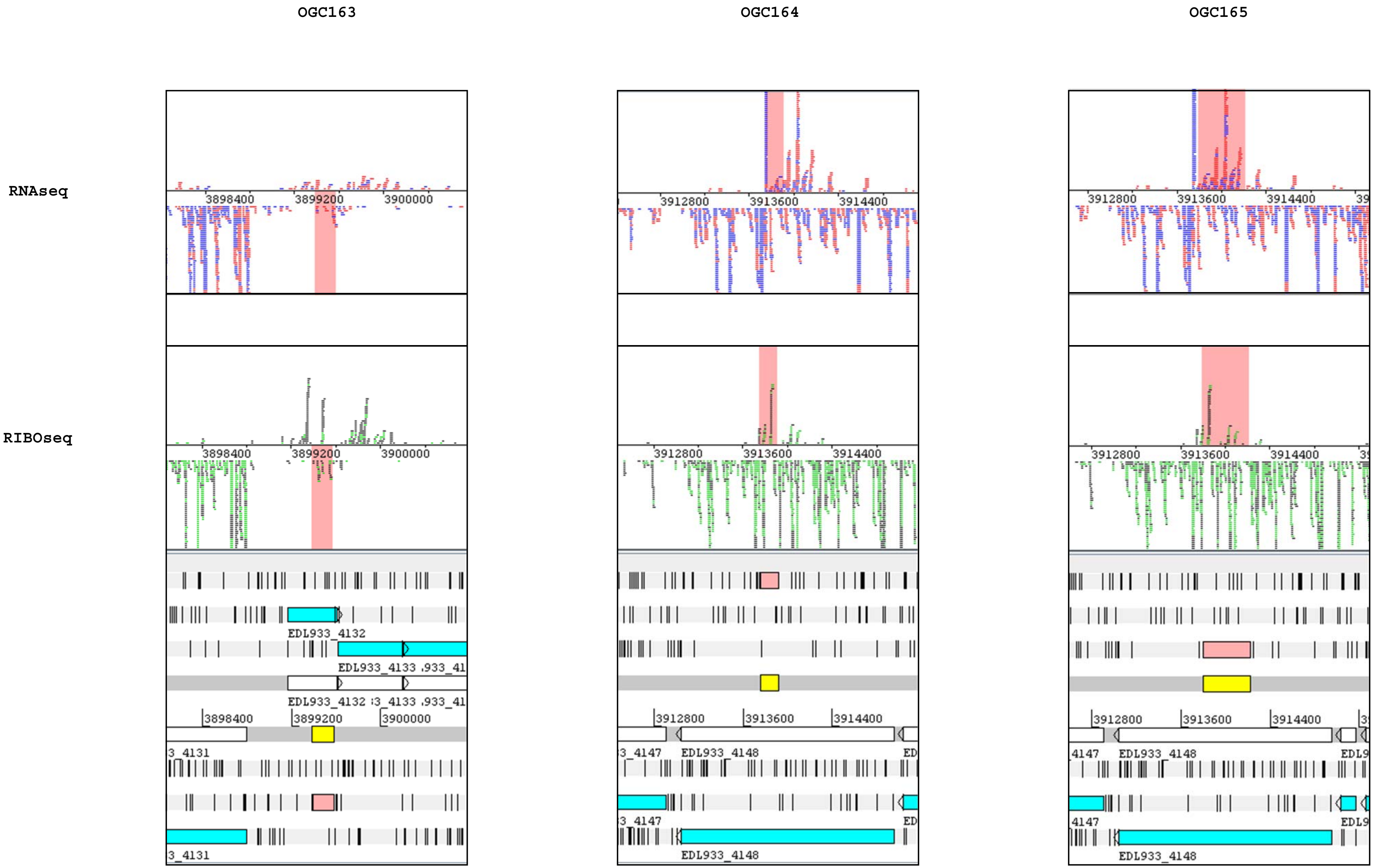
OGC163
OGC164
OGC165
RNAseq
RIBOseq
3898400
3899200
3900000
3912800
3913600
3914400
EDL933_4132
EDL933_4133
3_4131
3_4147
EDL933_4148
4147

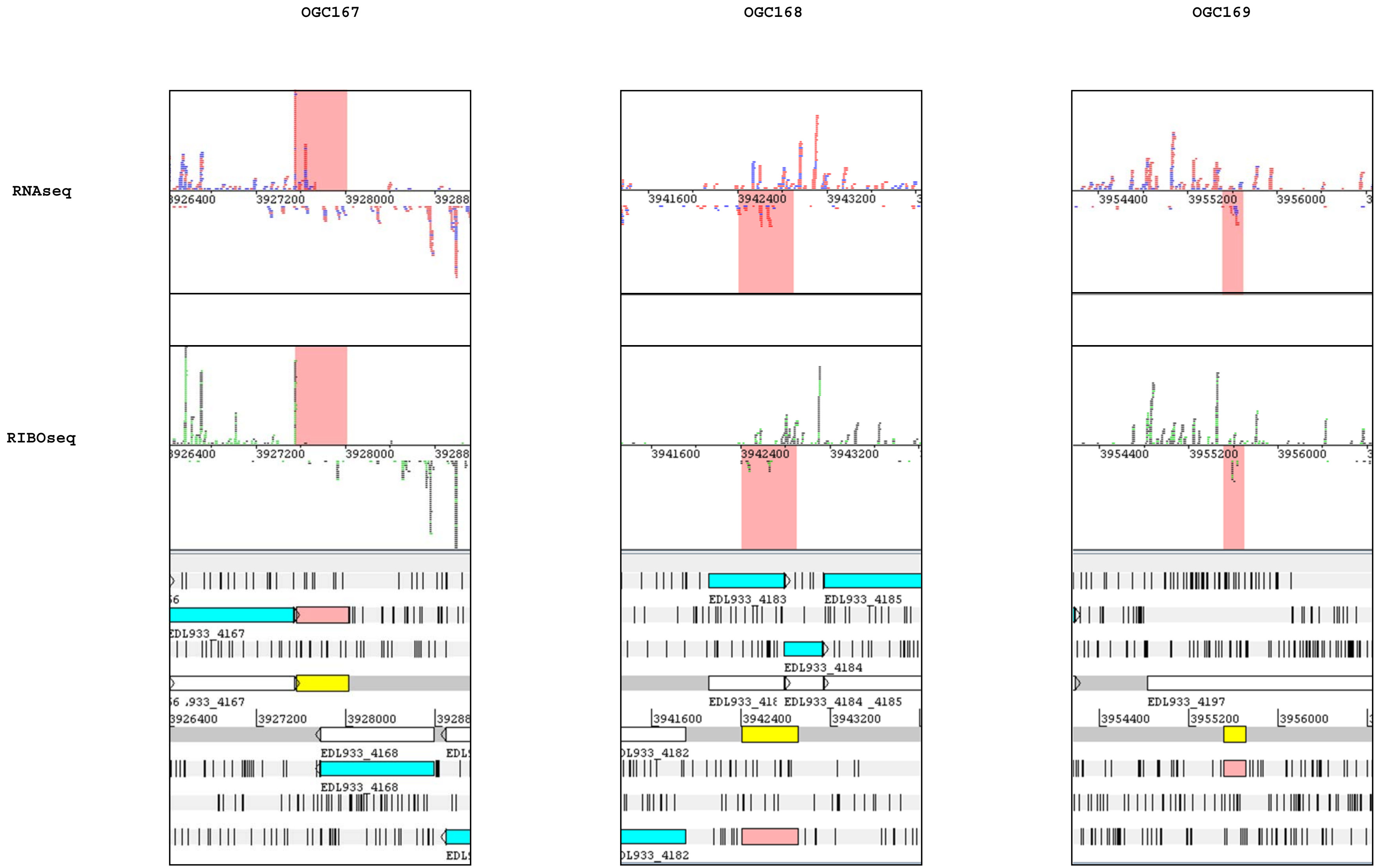
OGC167
OGC168
OGC169
RNAseq
RIBOseq
3927200
3928000
3941600
3942400
3943200
3954400
3955200
3956000
EDL933_4167
EDL933_4168
EDL933_4182
EDL933_4183
EDL933_4184
EDL933_4185
EDL933_4197

OGC171 OGC172 OGC173

RNAseq

RIBOseq

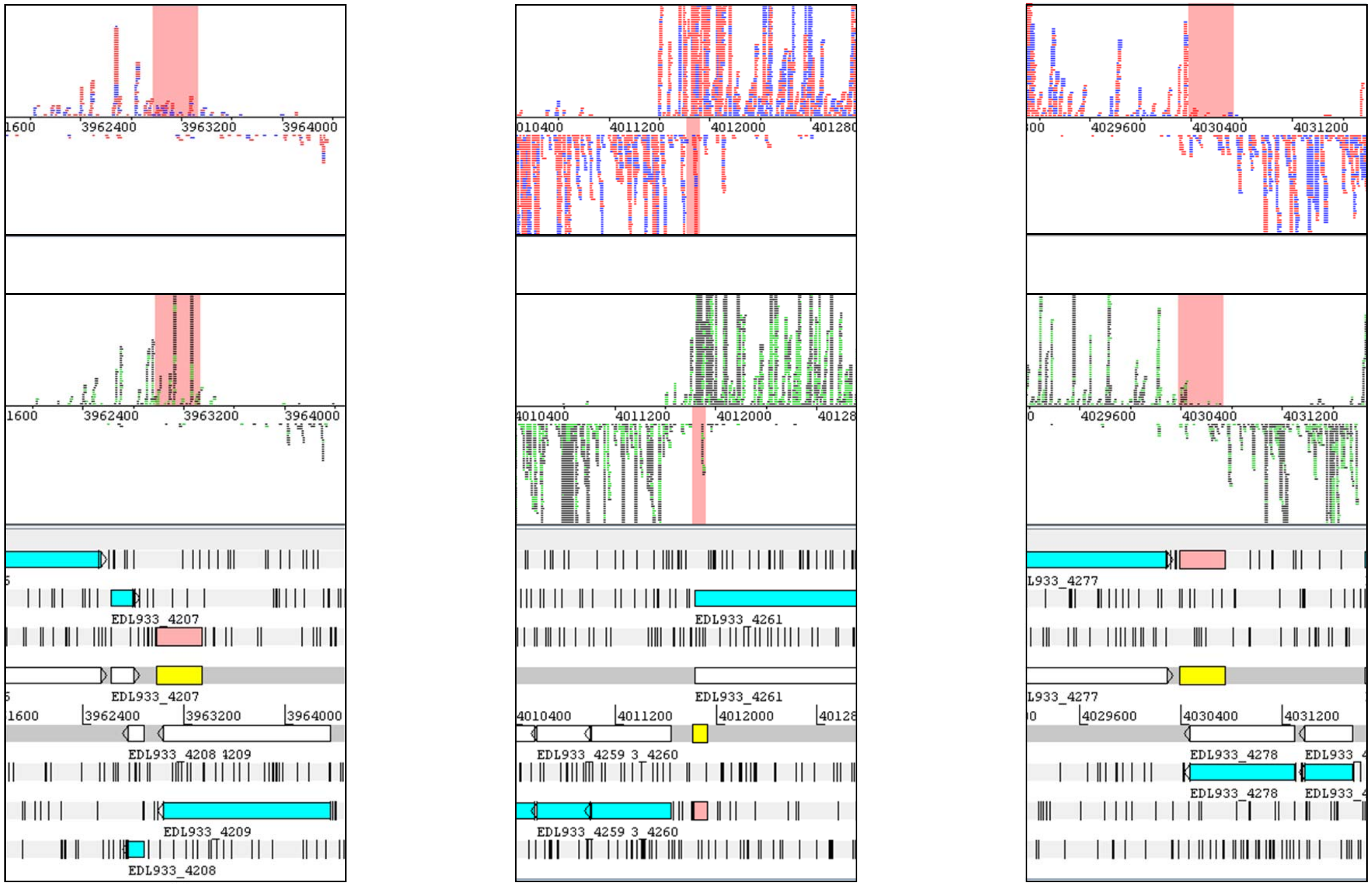

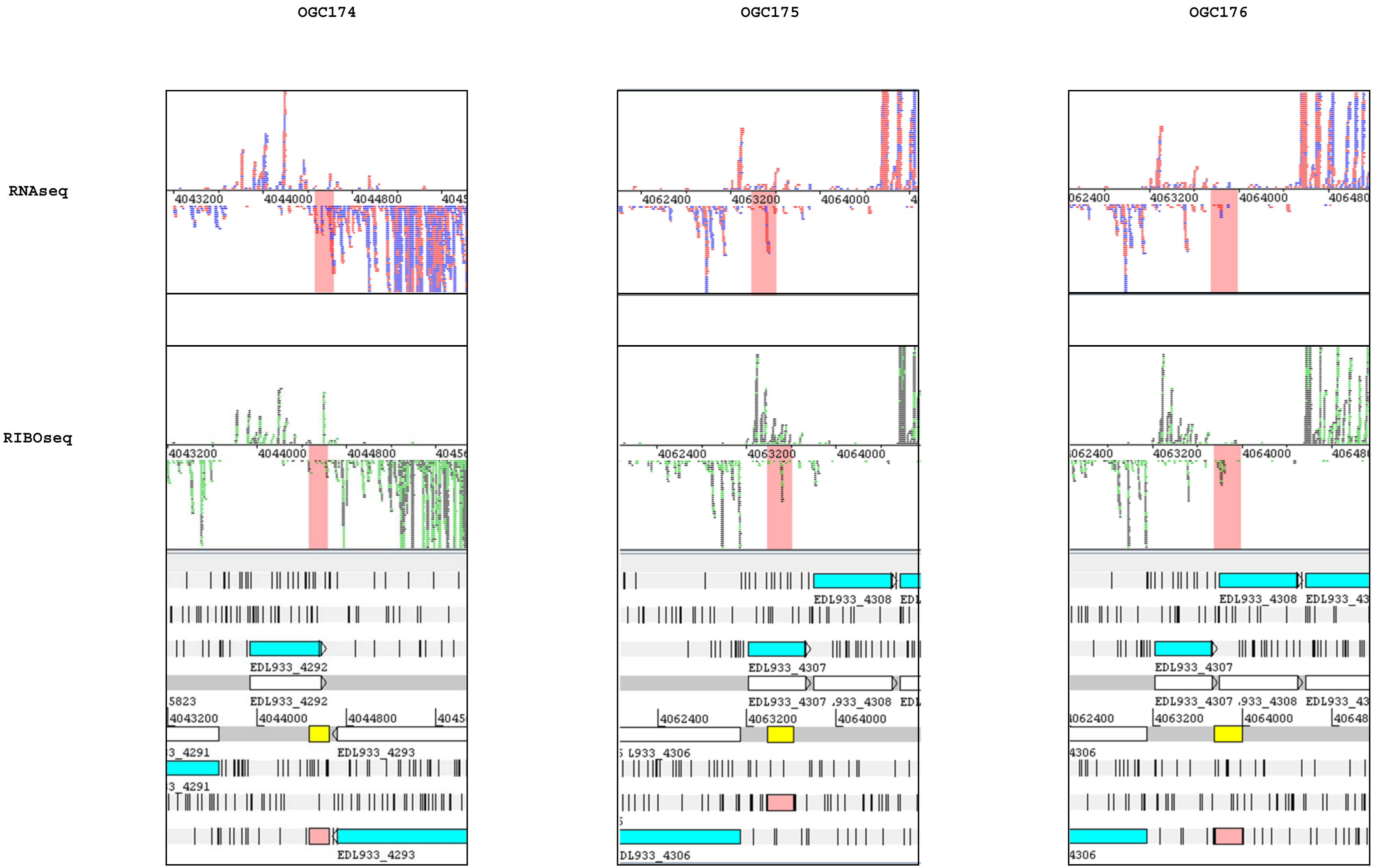
OGC174
OGC175
OGC176
RNAseq
RIBOseq
EDL933_4292
EDL933_4293
EDL933_4307
EDL933_4308
EDL933_4306

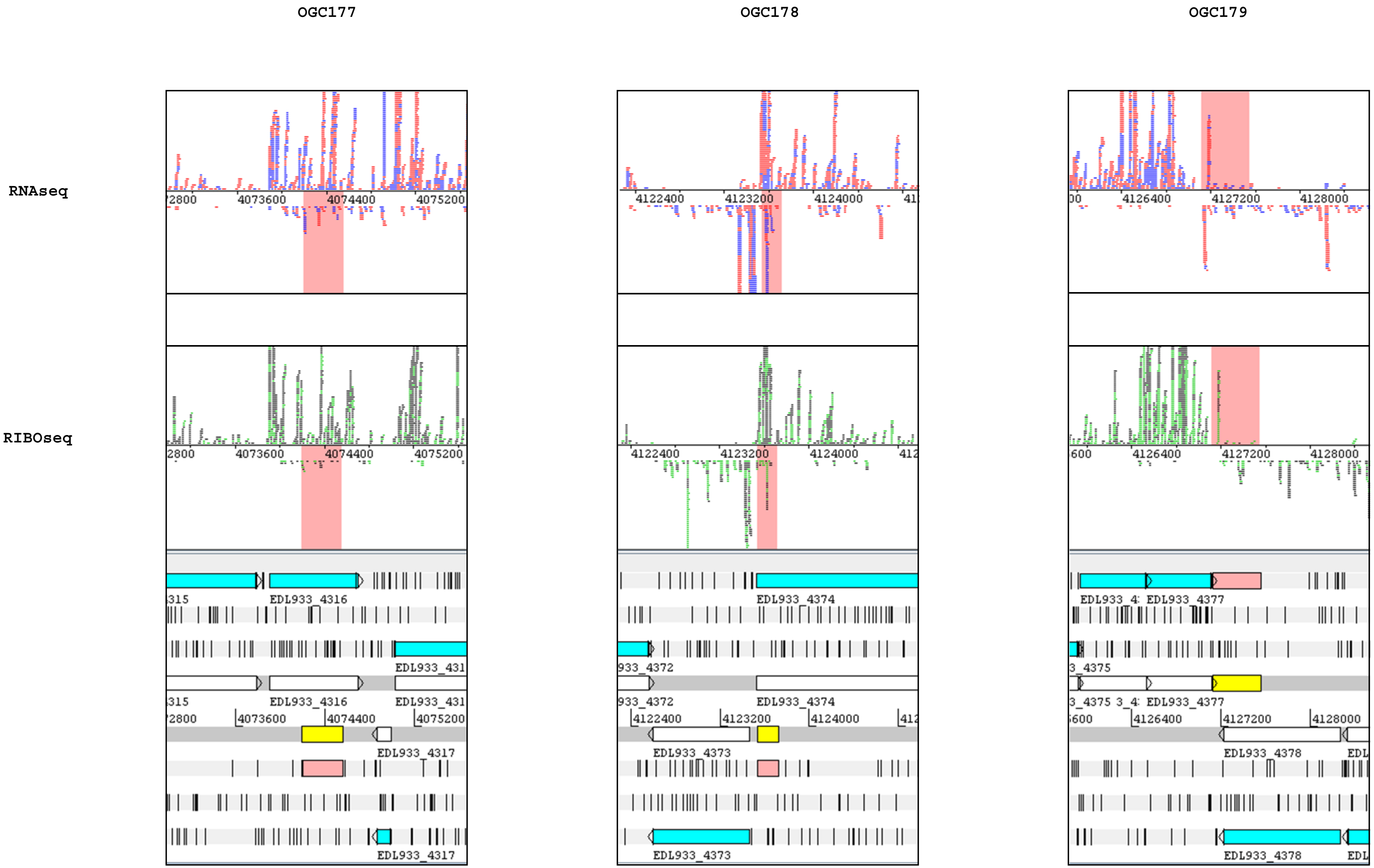
OGC177
OGC178
OGC179
RNAseq
RIBOseq
EDL933_4316
EDL933_4317
EDL933_4372
EDL933_4373
EDL933_4374
EDL933_4375
EDL933_4377
EDL933_4378

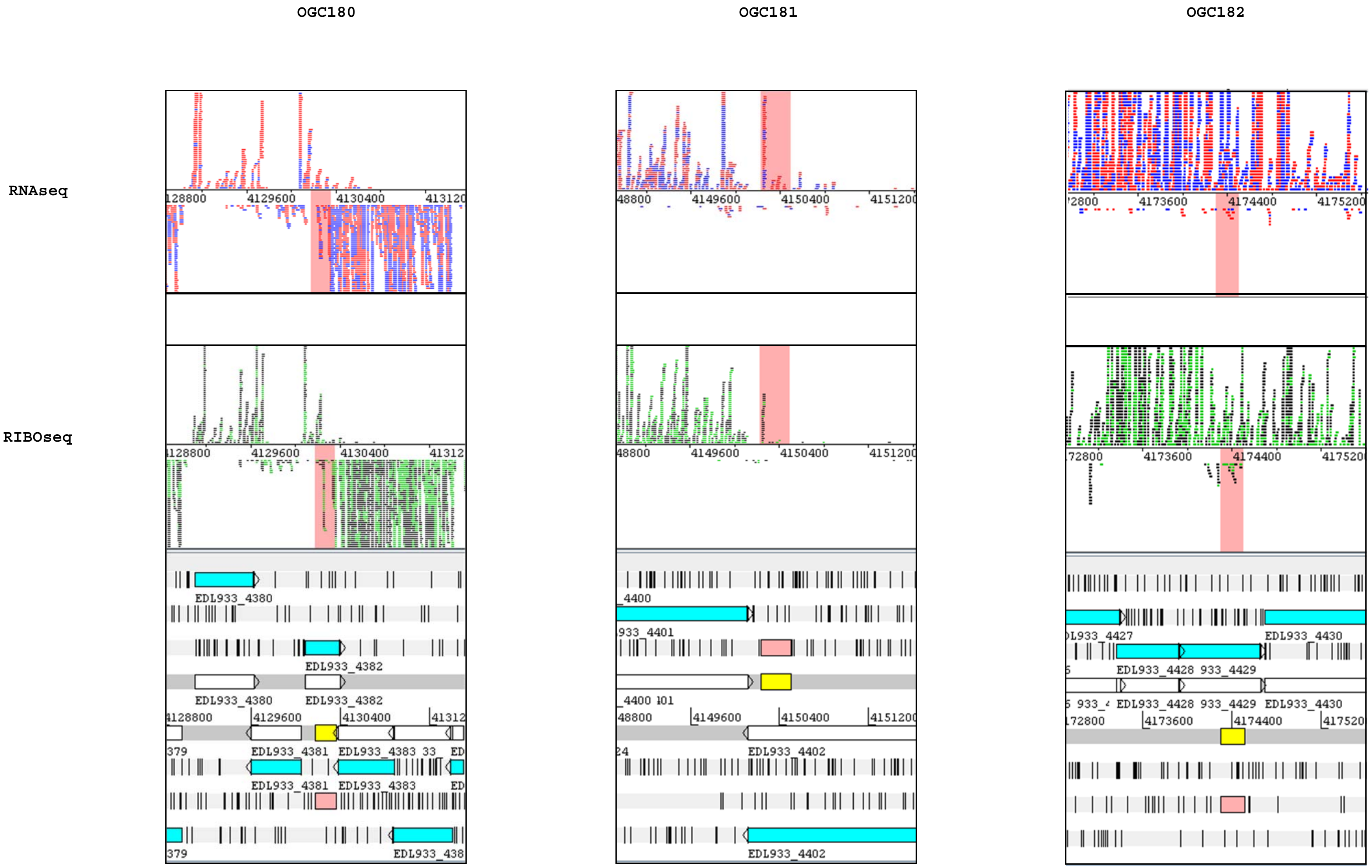
OGC180
OGC181
OGC182
RNAseq
RIBOseq
4129600
4130400
4149600
4150400
4173600
4174400
EDL933_4380
EDL933_4382
EDL933_4381
EDL933_4383
EDL933_4402
EDL933_4428
EDL933_4430

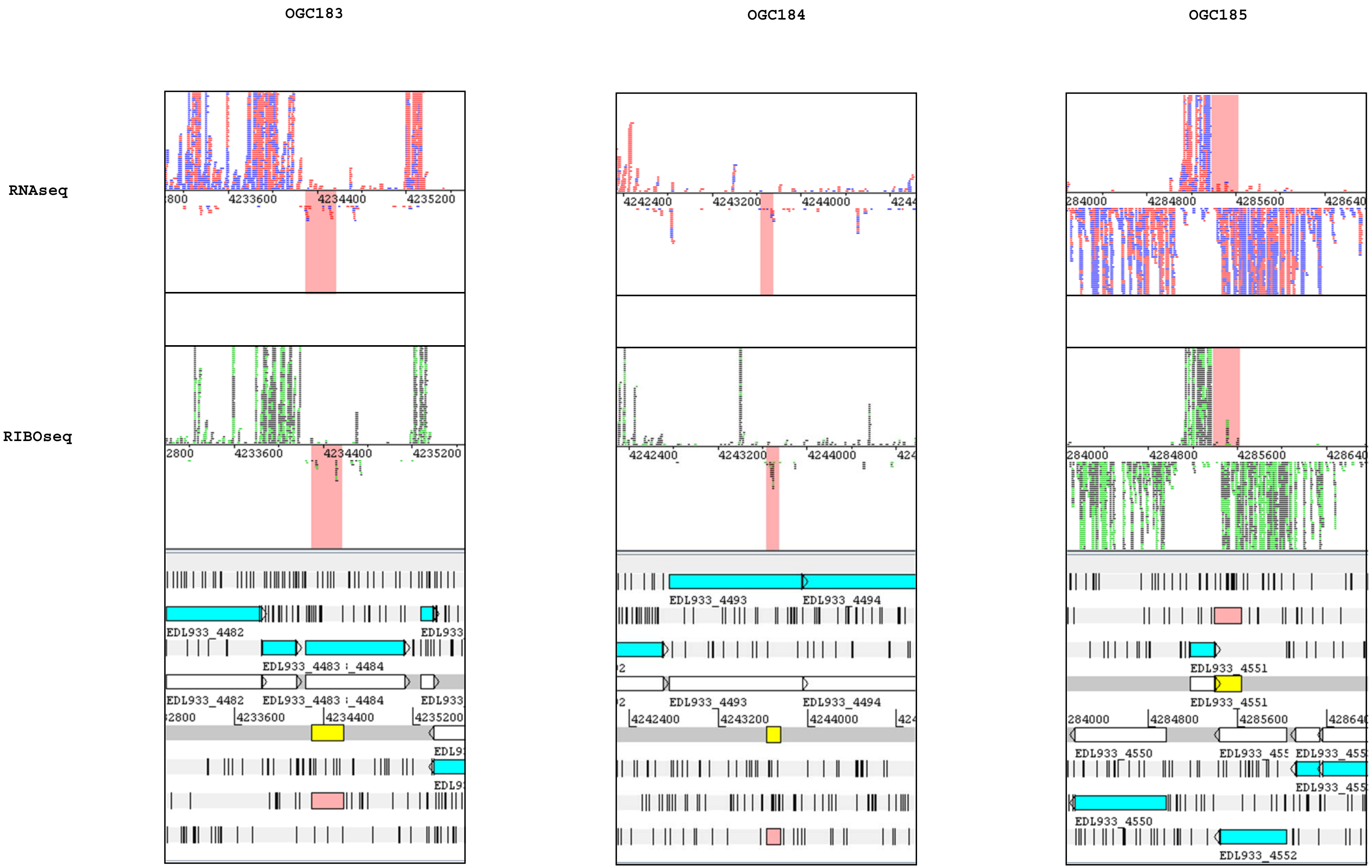
OGC183
OGC184
OGC185
RNAseq
RIBOseq
4233600
4234400
4235200
4242400
4243200
4244000
284000
4284800
4285600
EDL933_4482
EDL933_4483
_4484
EDL933_4493
EDL933_4494
EDL933_4550
EDL933_4551
EDL933_4552

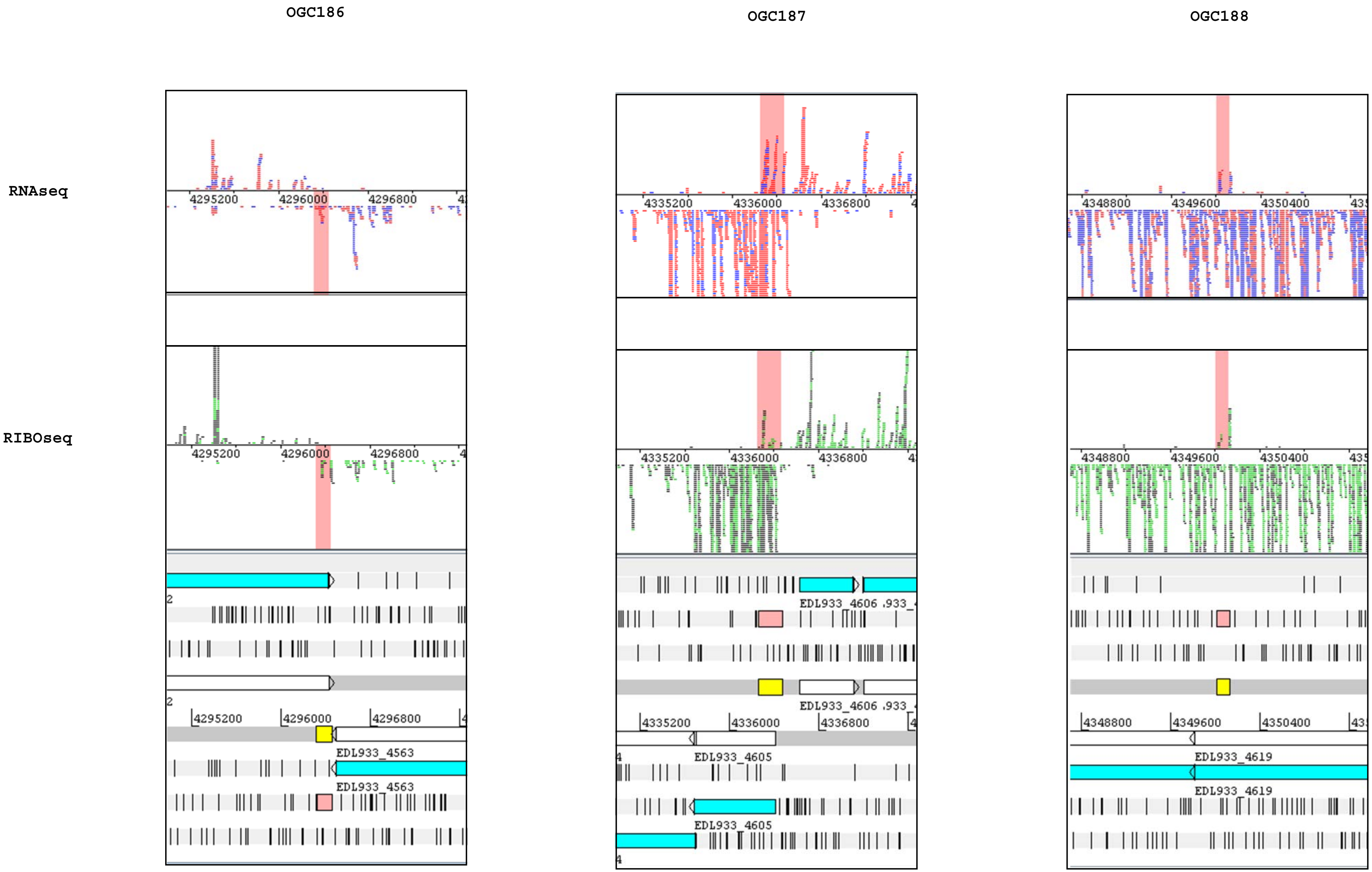

OGC186
OGC187
OGC188
RNAseq
RIBOseq
4295200
4296000
4296800
4335200
4336000
4336800
4348800
4349600
4350400
EDL933_4563
EDL933_4605
EDL933_4606
EDL933_4619

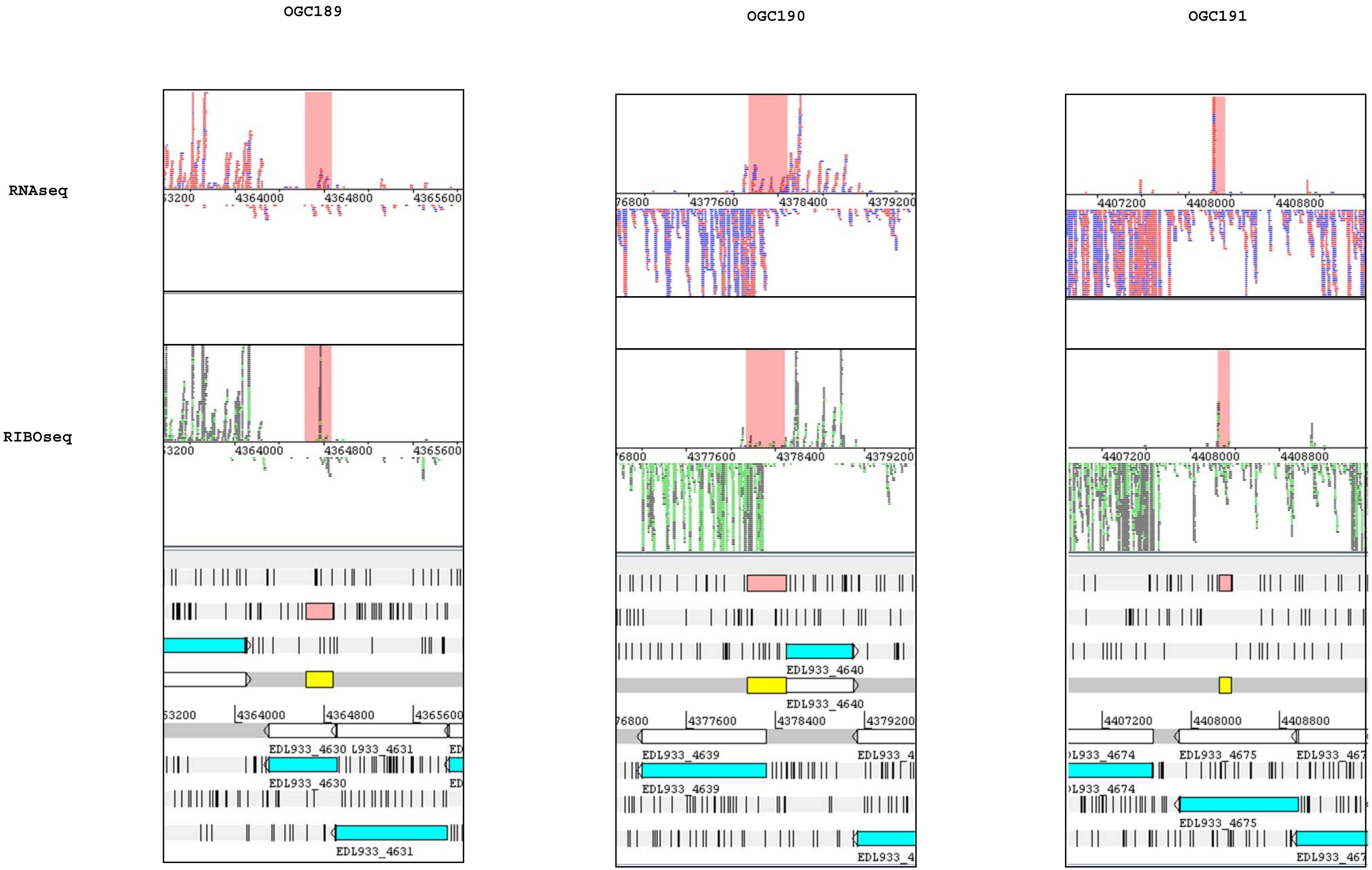

OGC189
OGC190
OGC191
RNAseq
RIBOseq
4364000
4364800
4365600
4377600
4378400
4379200
4407200
4408000
4408800
EDL933_4630
EDL933_4631
EDL933_4639
EDL933_4640
EDL933_4675

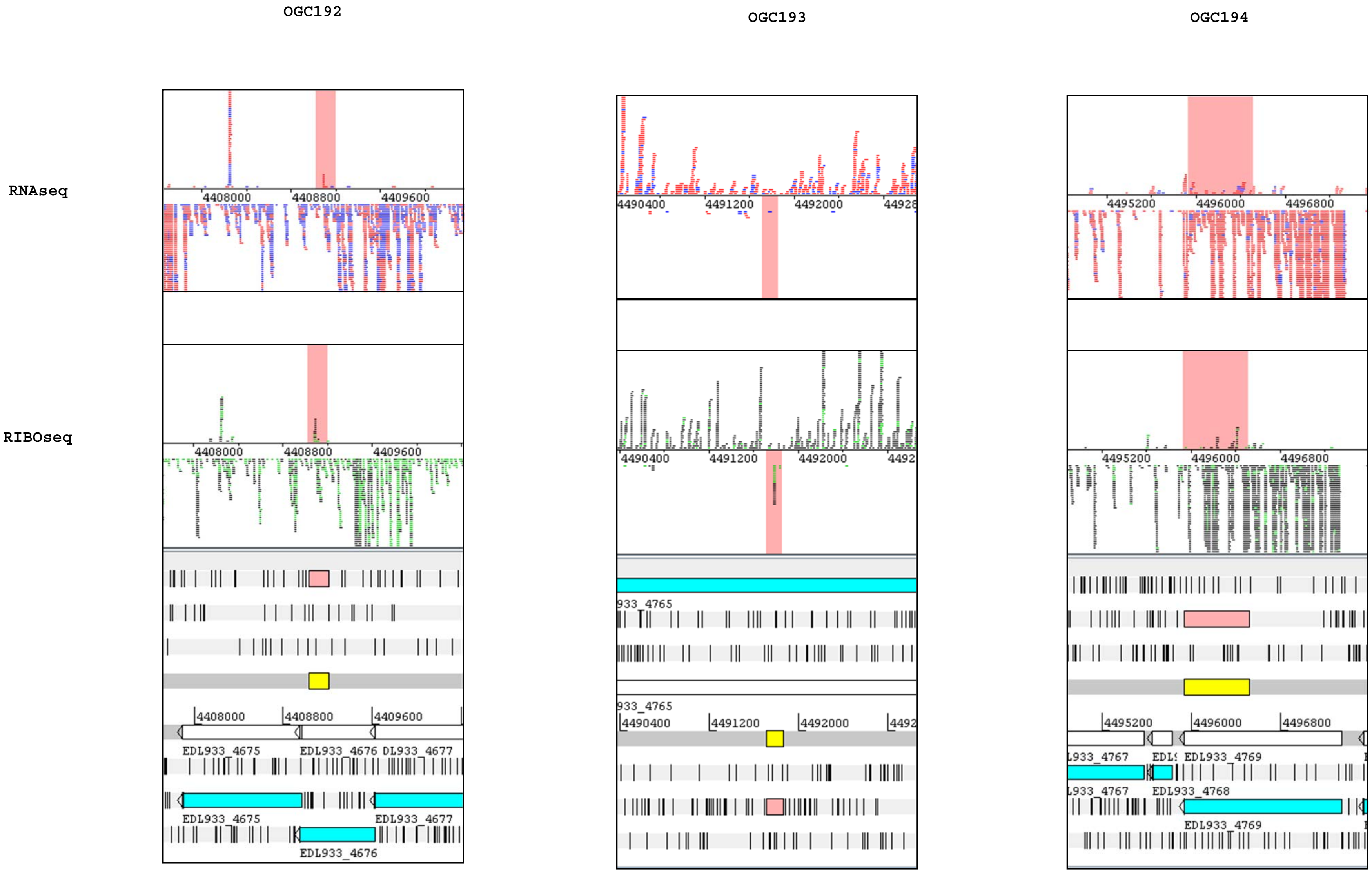
OGC192
OGC193
OGC194
RNAseq
RIBOseq
4408000
4408800
4409600
EDL933_4675
EDL933_4676
DL933_4677
EDL933_4677
4490400
4491200
4492000
933_4765
4495200
4496000
4496800
EDL933_4768
EDL933_4769

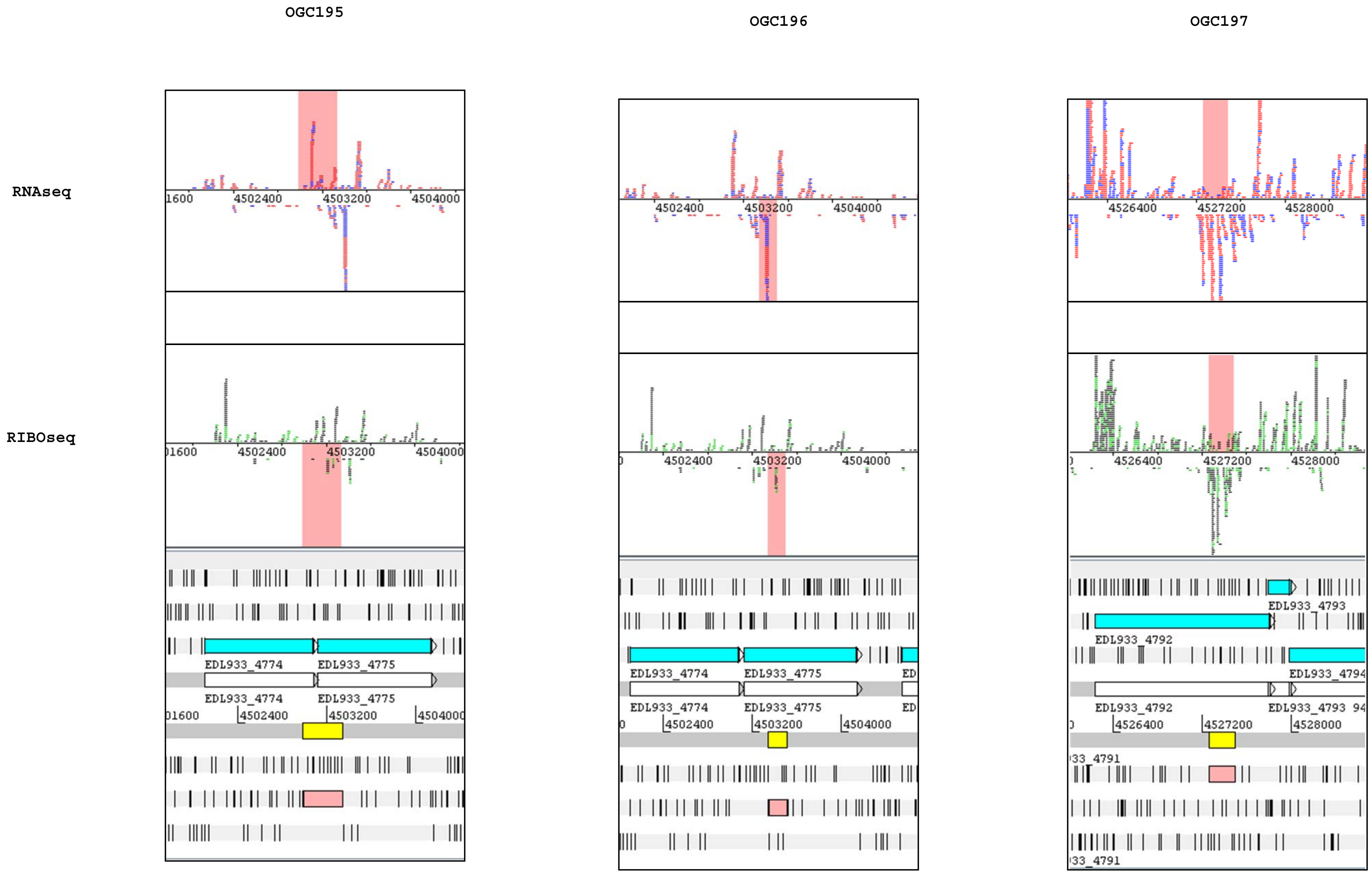
OGC195
OGC196
OGC197
RNAseq
RIBOseq
4502400
4503200
4504000
4526400
4527200
4528000
EDL933_4774
EDL933_4775
EDL933_4792
EDL933_4793
EDL933_4794

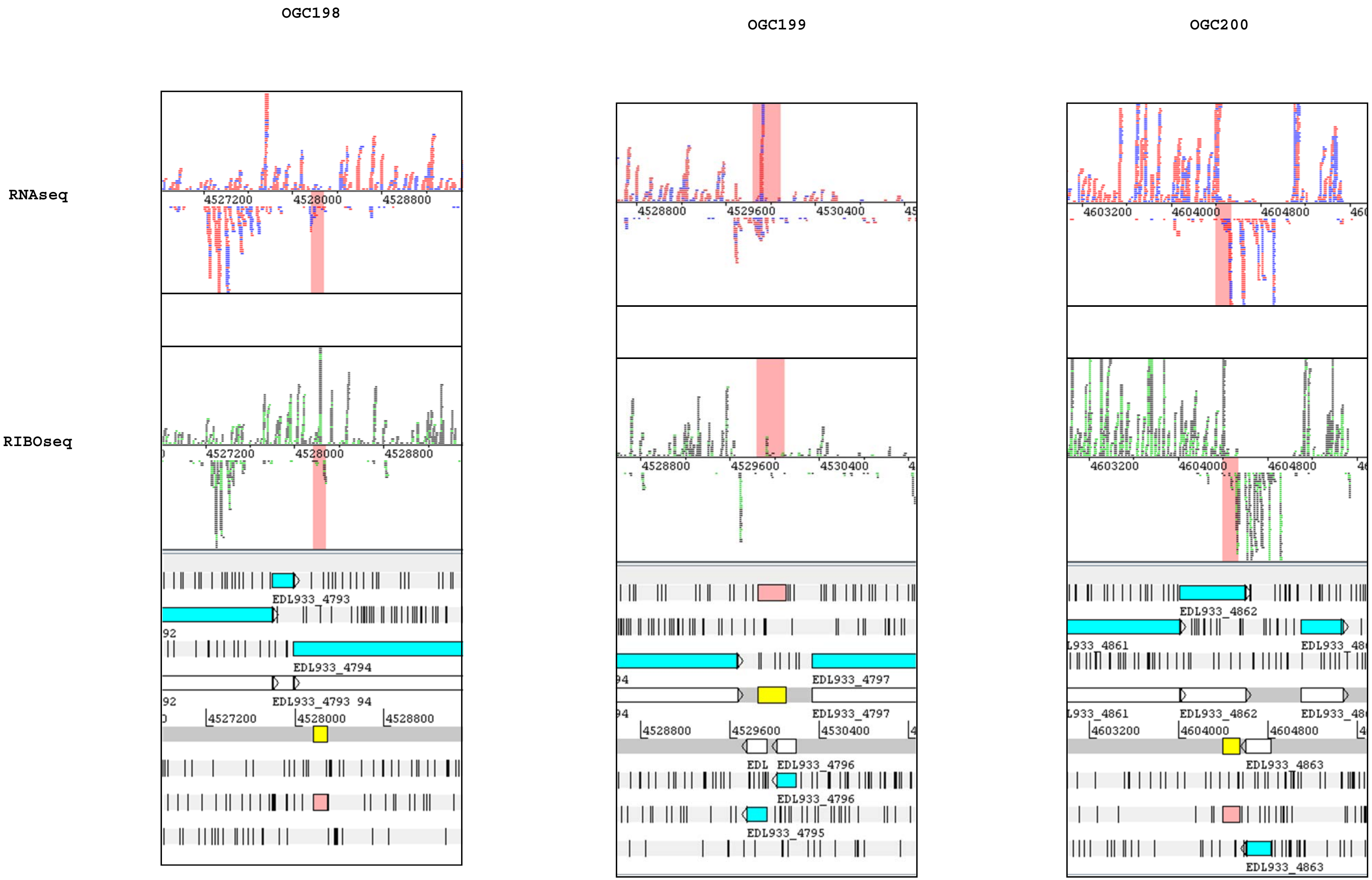
OGC198
OGC199
OGC200
RNAseq
RIBOseq
4527200
4528000
4528800
EDL933_4793
EDL933_4794
4528800
4529600
4530400
EDL933_4797
EDL933_4796
EDL933_4795
4603200
4604000
4604800
EDL933_4862
EDL933_4863

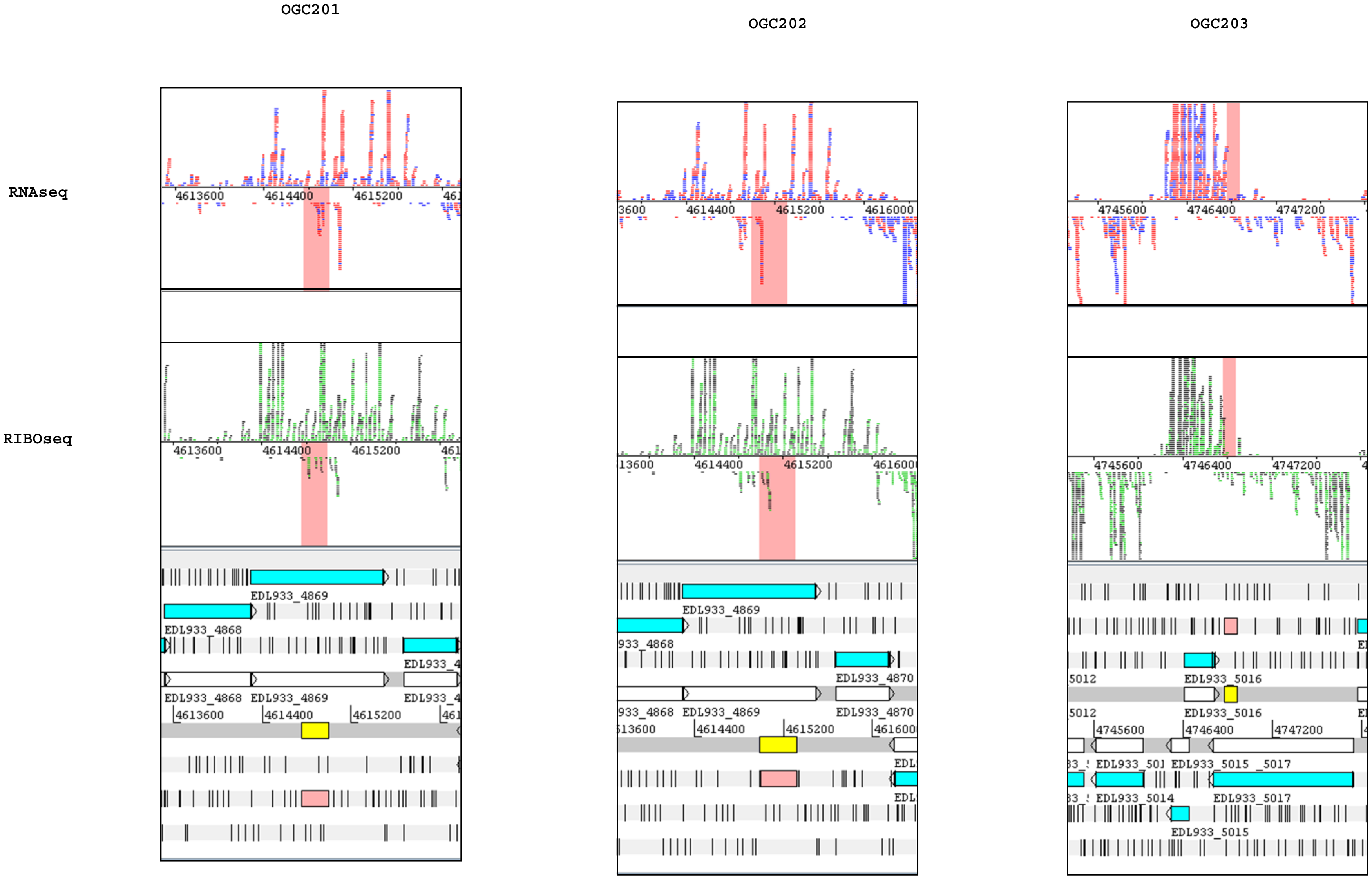
OGC201
OGC202
OGC203
RNAseq
RIBOseq
4613600
4614400
4615200
4616000
4745600
4746400
4747200
EDL933_4868
EDL933_4869
EDL933_4870
EDL933_5014
EDL933_5015
EDL933_5016
EDL933_5017

OGC204 OGC205 OGC206

RNAseq

RIBOseq

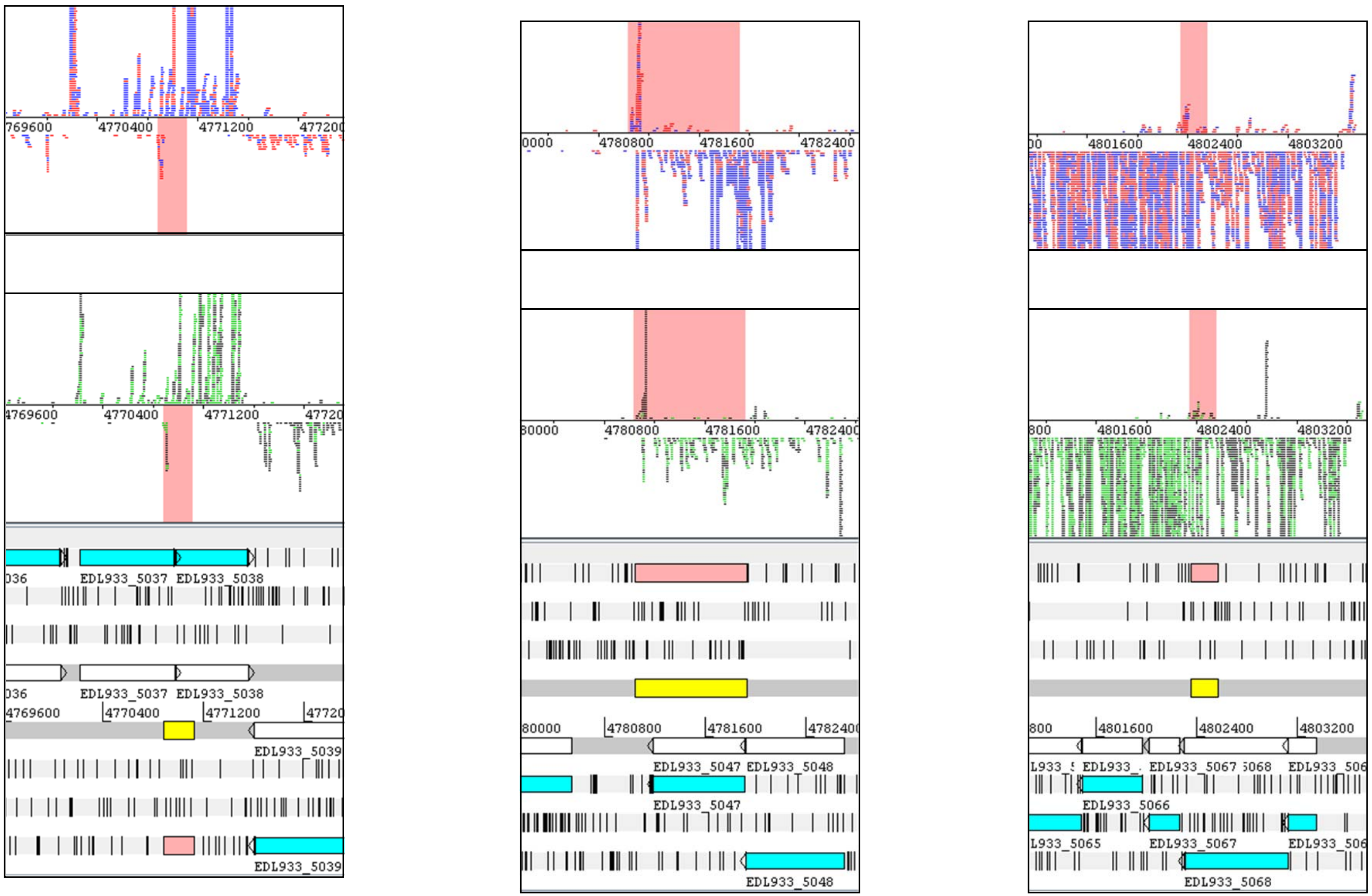

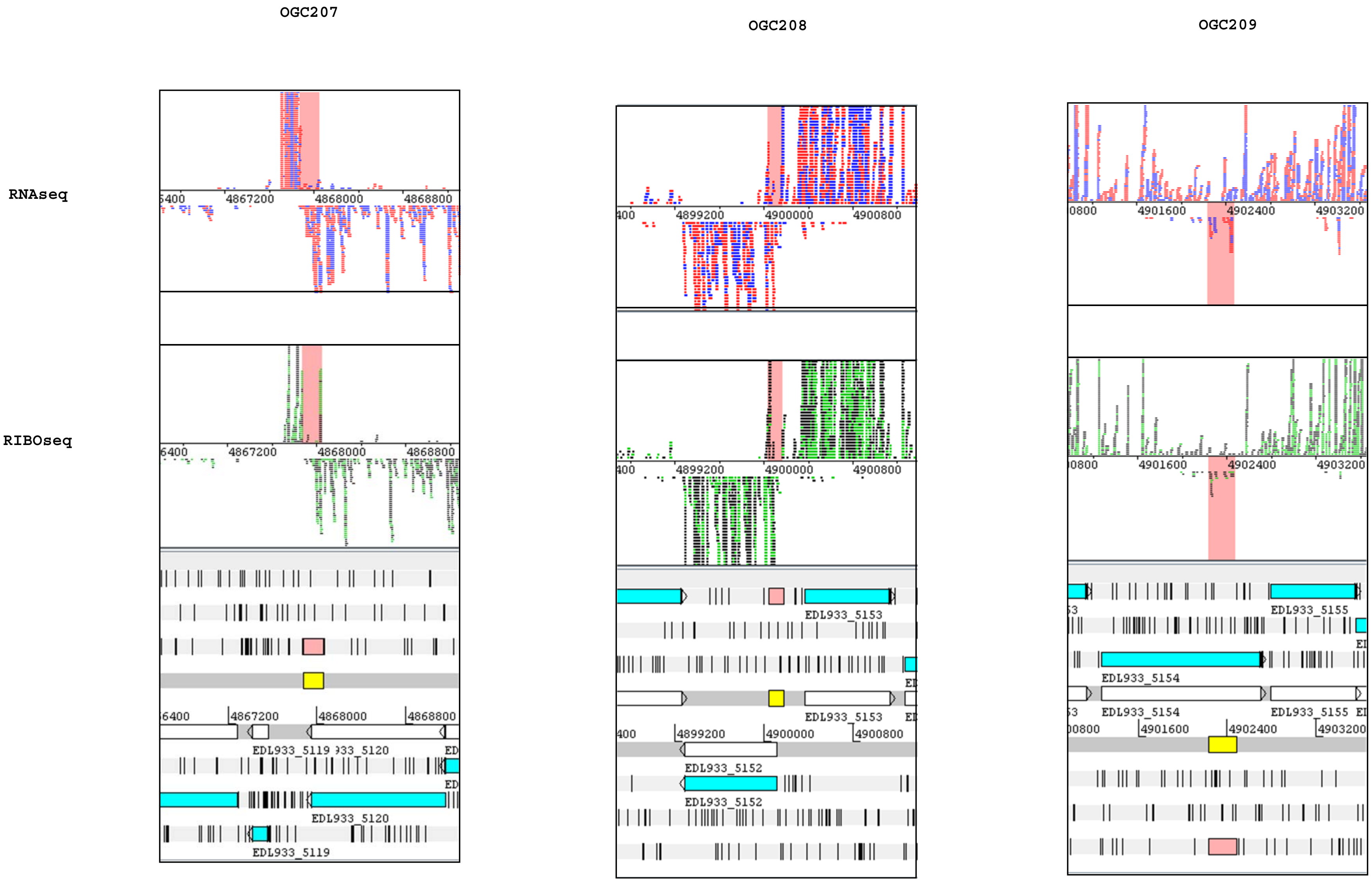
OGC207
OGC208
OGC209
RNAseq
RIBOseq
4867200
4868000
4868800
4899200
4900000
4900800
4901600
4902400
4903200
EDL933_5119
EDL933_5120
EDL933_5152
EDL933_5153
EDL933_5154
EDL933_5155

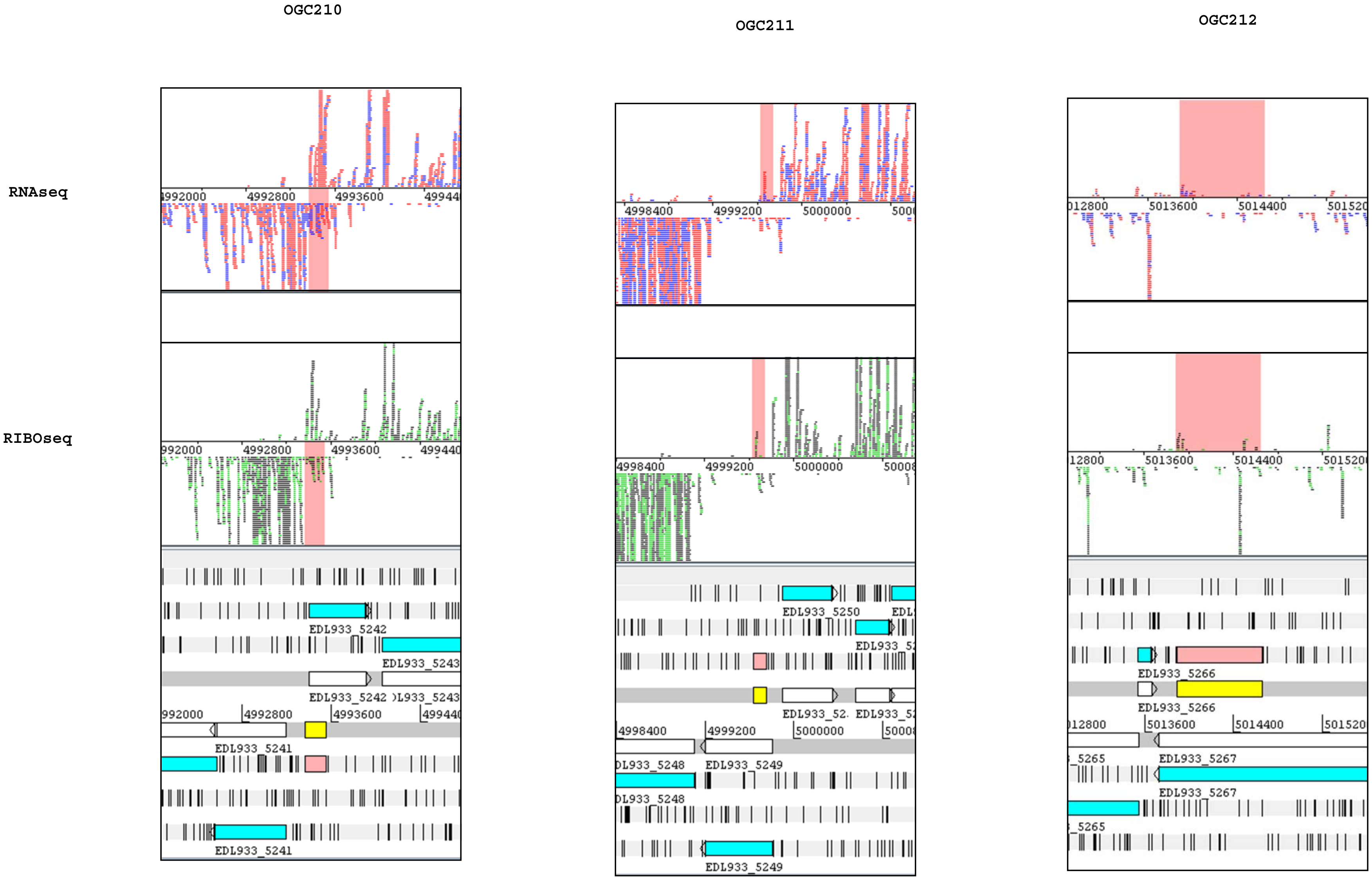
OGC210
OGC211
OGC212
RNAseq
RIBOseq
EDL933_5241
EDL933_5242
EDL933_5243
EDL933_5248
EDL933_5249
EDL933_5250
EDL933_5265
EDL933_5266
EDL933_5267

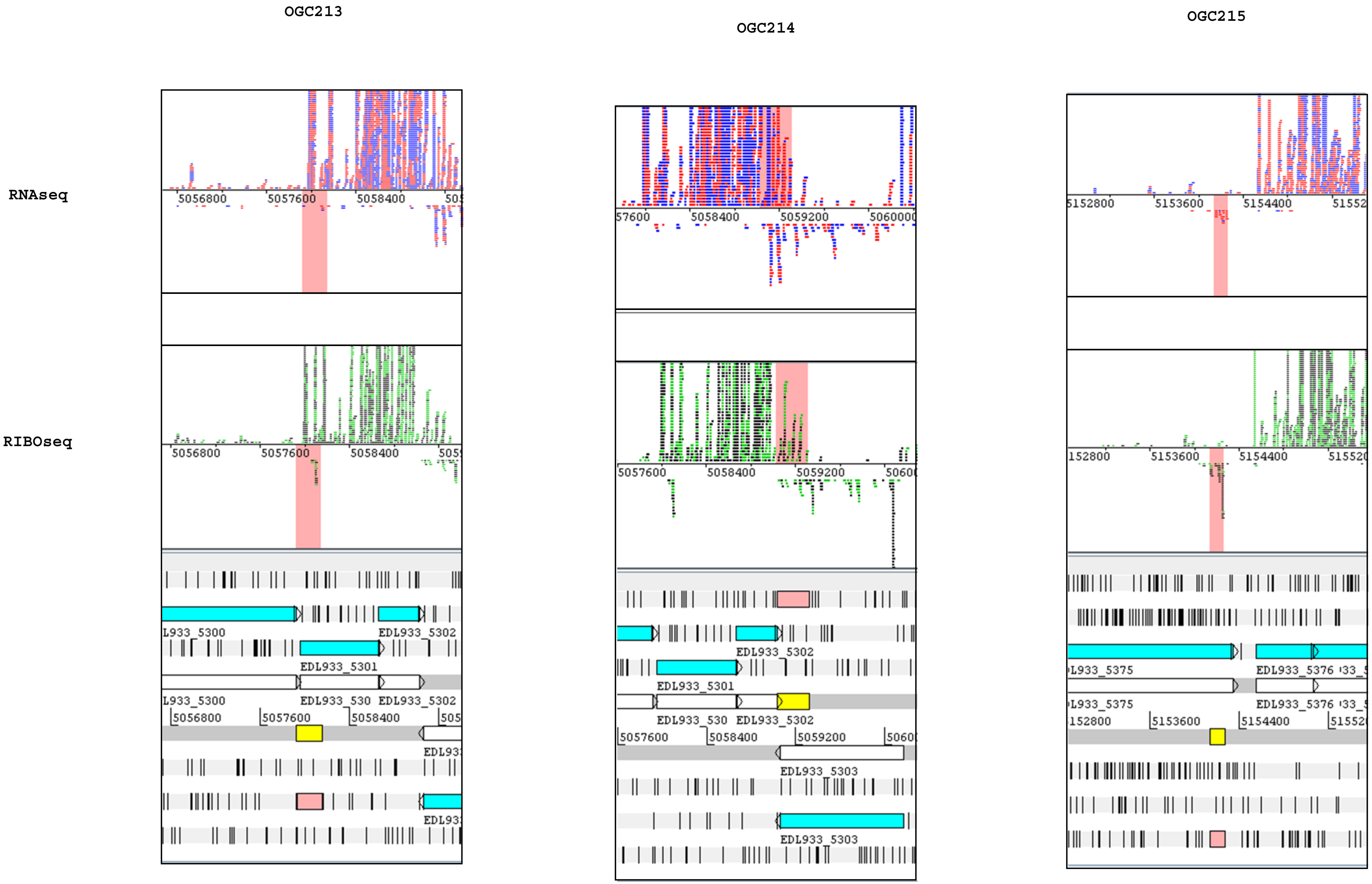
OGC213
OGC214
OGC215
RNAseq
RIBOseq
EDL933_5301
EDL933_5302
EDL933_5303
EDL933_5375
EDL933_5376

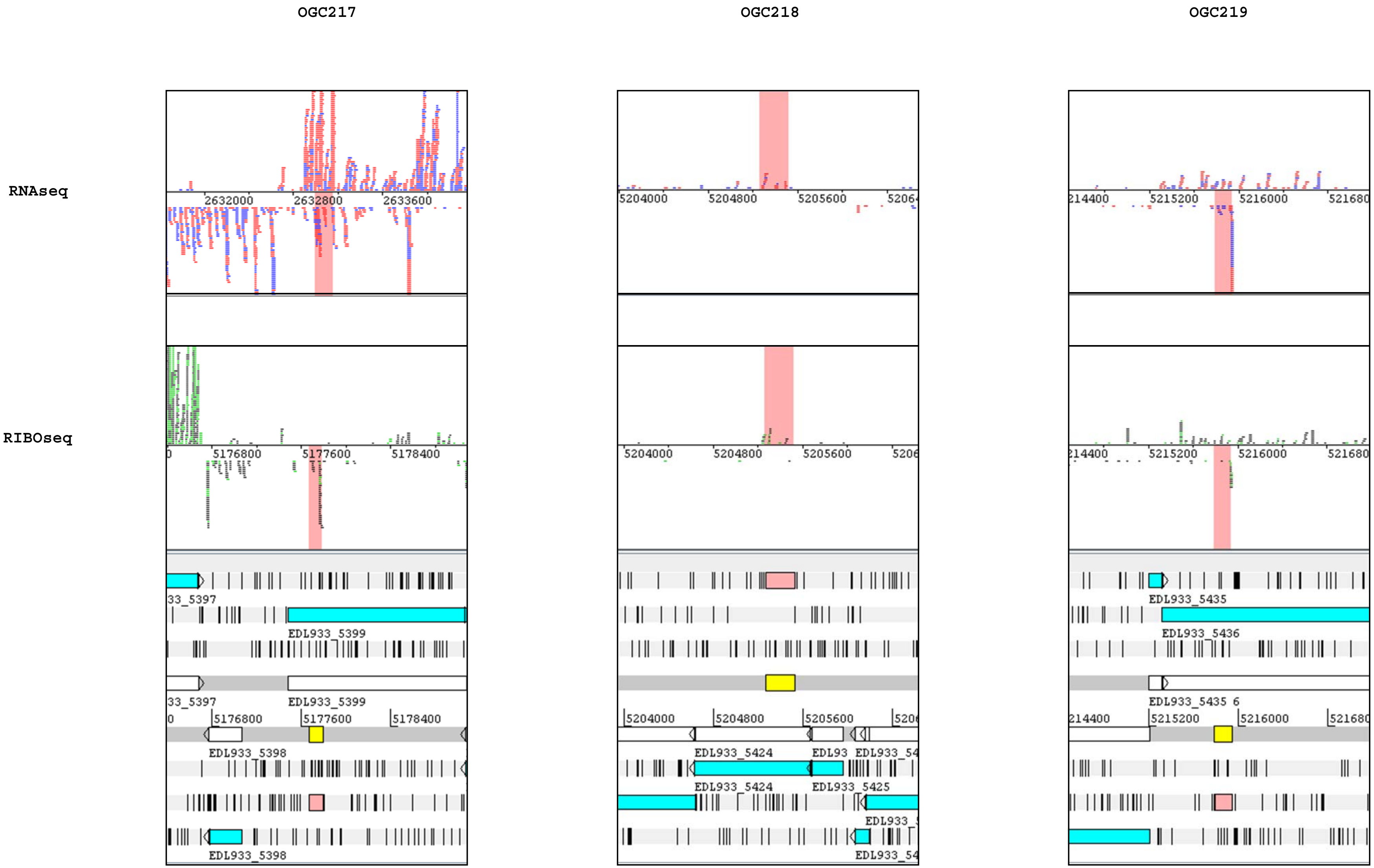
OGC217
OGC218
OGC219
RNAseq
RIBOseq
2632000
2632800
2633600
5204000
5204800
5205600
5215200
5216000
5176800
5177600
5178400
EDL933_5399
EDL933_5398
EDL933_5424
EDL933_5425
EDL933_5435
EDL933_5436

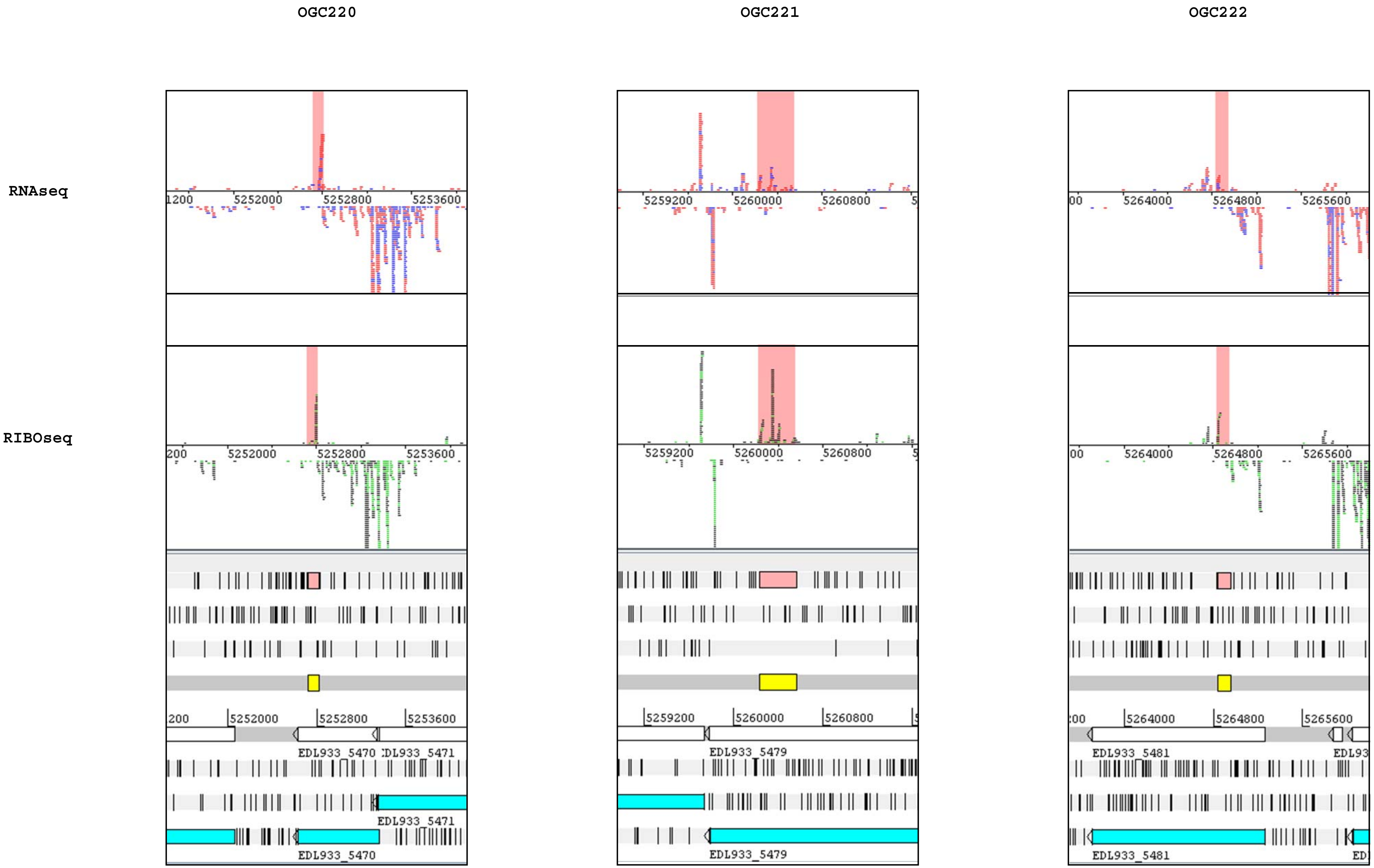

OGC220
OGC221
OGC222
RNAseq
RIBOseq
5252000
5252800
5253600
5259200
5260000
5260800
5264000
5264800
5265600
EDL933_5470
EDL933_5471
EDL933_5479
EDL933_5481

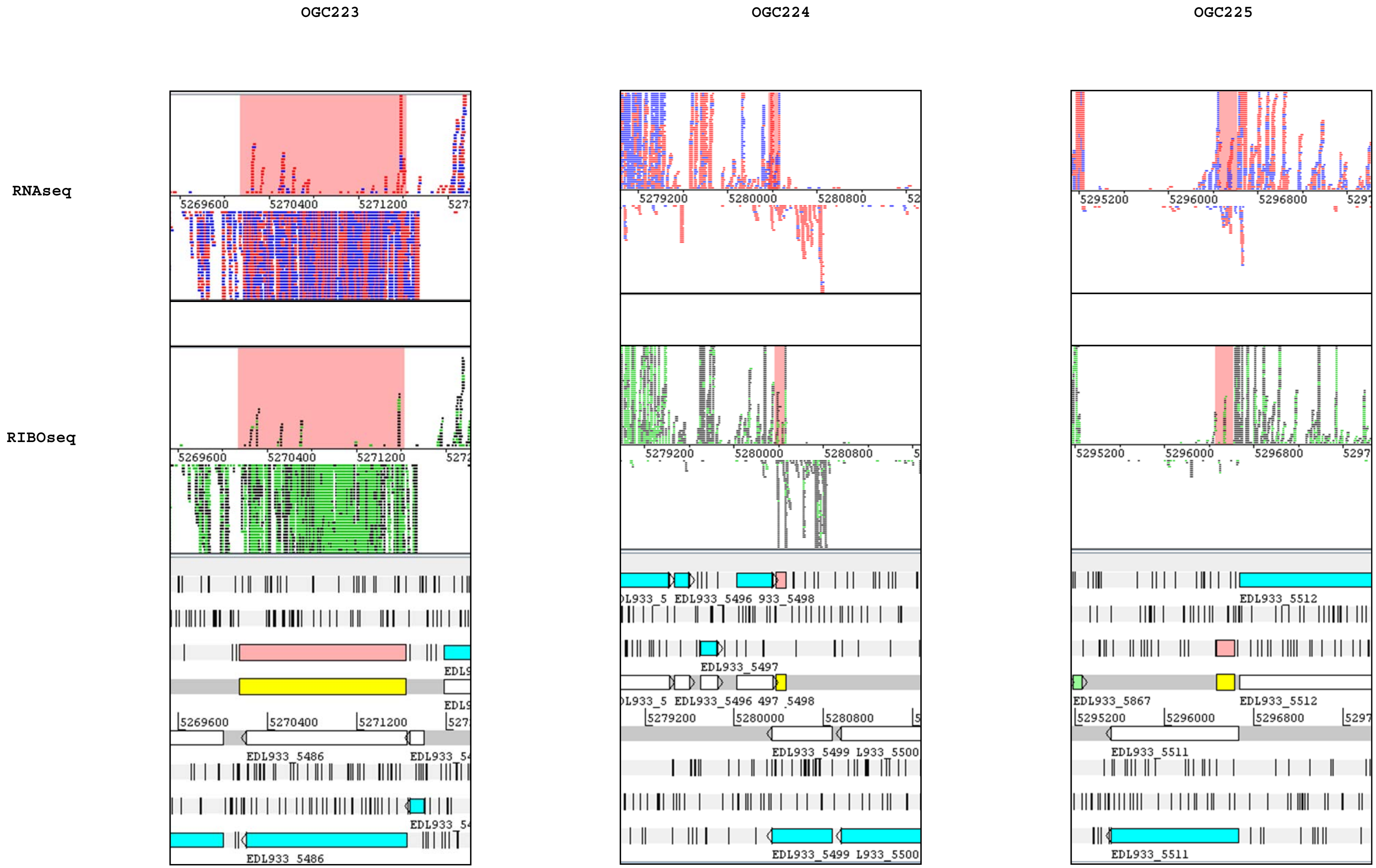
OGC223
OGC224
OGC225
RNAseq
RIBOseq
5269600
5270400
5271200
5279200
5280000
5280800
5295200
5296000
5296800
EDL933_5486
EDL933_5496
EDL933_5497
EDL933_5498
EDL933_5499
EDL933_5867
EDL933_5511
EDL933_5512

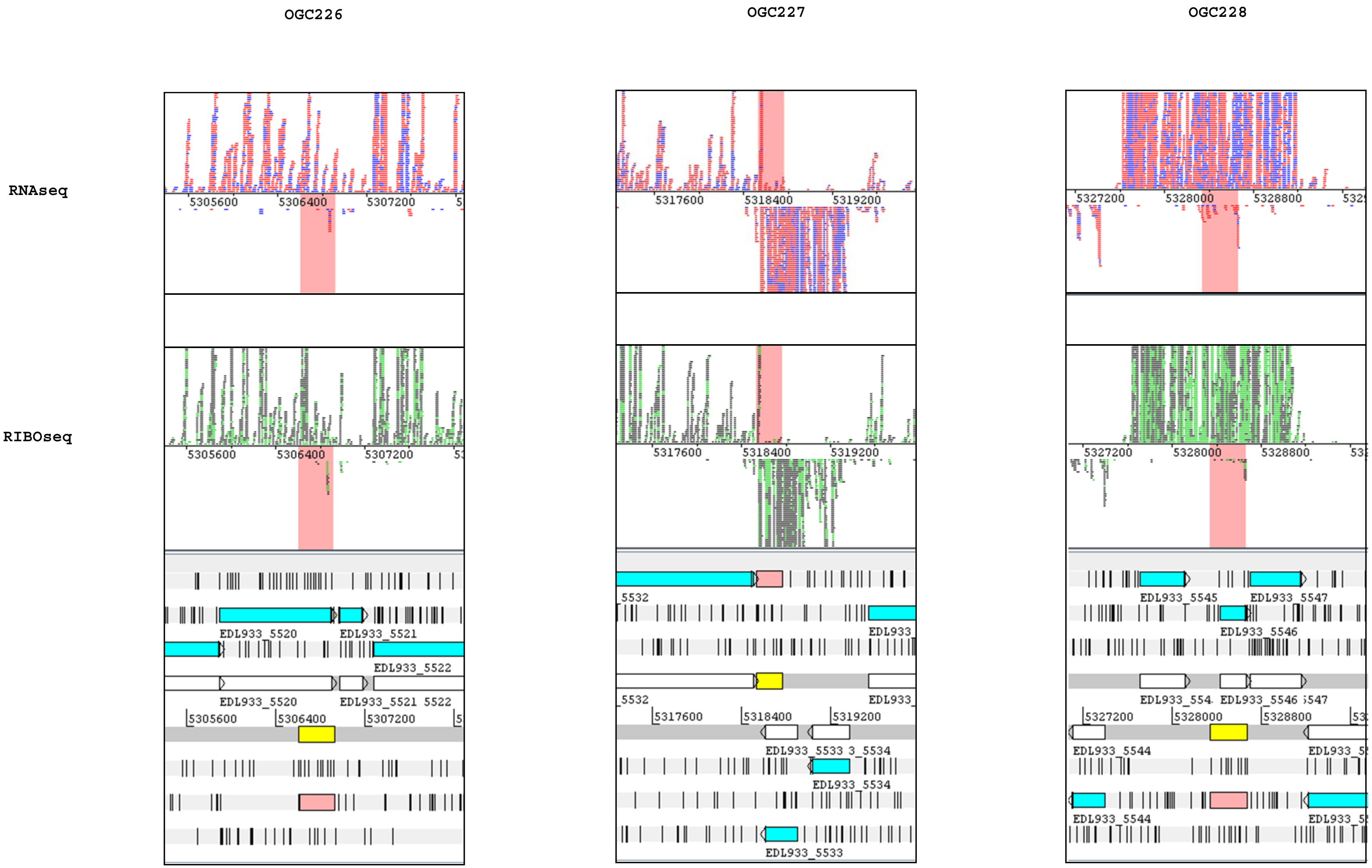
OGC226
OGC227
OGC228
RNAseq
RIBOseq
5305600
5306400
5307200
5317600
5318400
5319200
5327200
5328000
5328800
EDL933_5520
EDL933_5521
EDL933_5522
5522
EDL933_5533
EDL933_5534
EDL933_5544
EDL933_5545
EDL933_5546
EDL933_5547

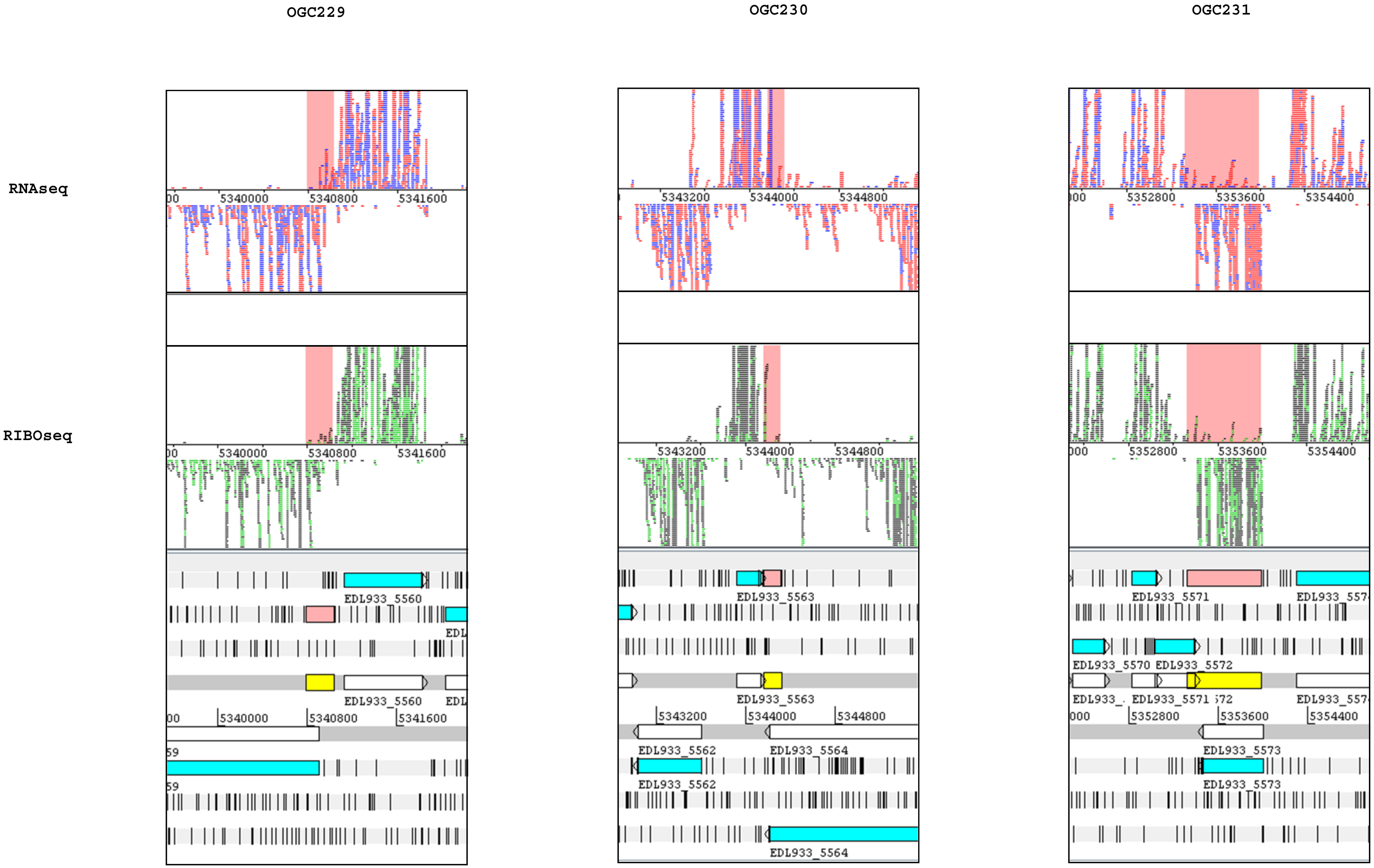
OGC229
OGC230
OGC231
RNAseq
RIBOseq
5340000
5340800
5341600
5343200
5344000
5344800
5352800
5353600
5354400
EDL933_5560
EDL933_5562
EDL933_5563
EDL933_5564
EDL933_5570
EDL933_5571
EDL933_5572
EDL933_5573

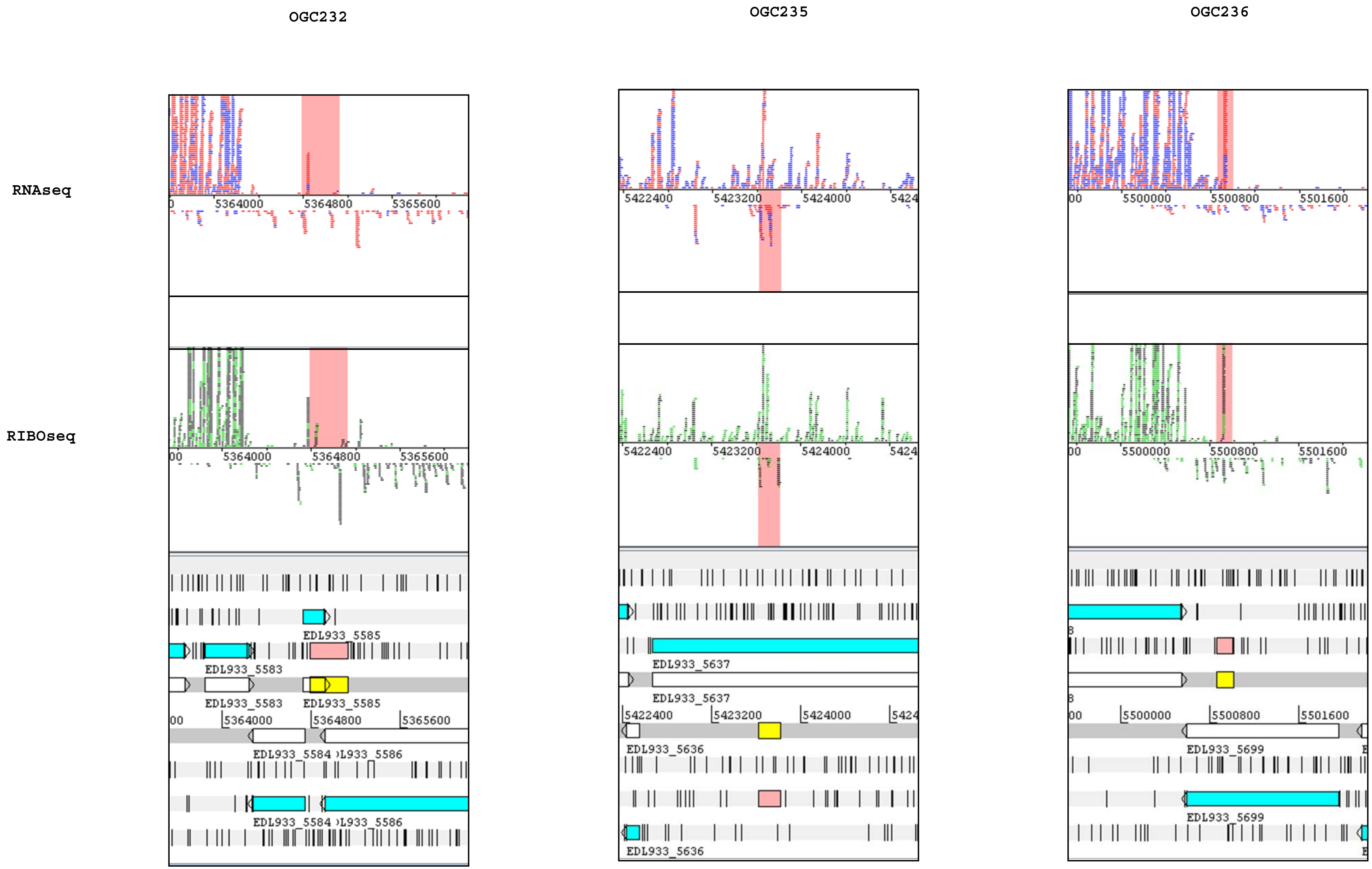
OGC232
OGC235
OGC236
RNAseq
RIBOseq
5364000
5364800
5365600
5422400
5423200
5424000
5500000
5500800
5501600
EDL933_5585
EDL933_5583
EDL933_5584
EDL933_5637
EDL933_5636
EDL933_5699

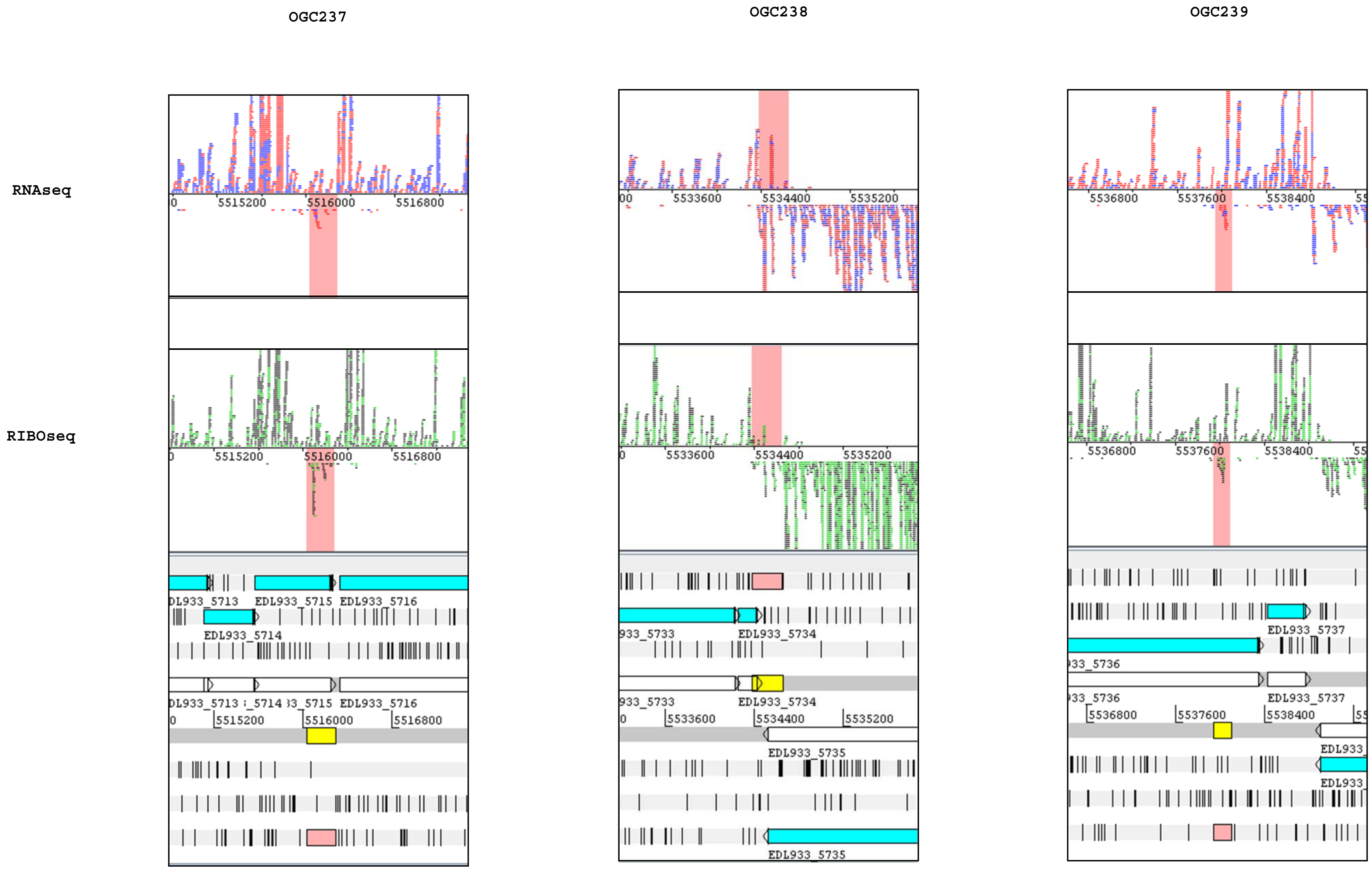
OGC237
OGC238
OGC239
RNAseq
RIBOseq
5515200
5516000
5516800
EDL933_5715
EDL933_5716
EDL933_5714
EDL933_5716
5533600
5534400
5535200
EDL933_5734
EDL933_5735
5536800
5537600
5538400
EDL933_5737

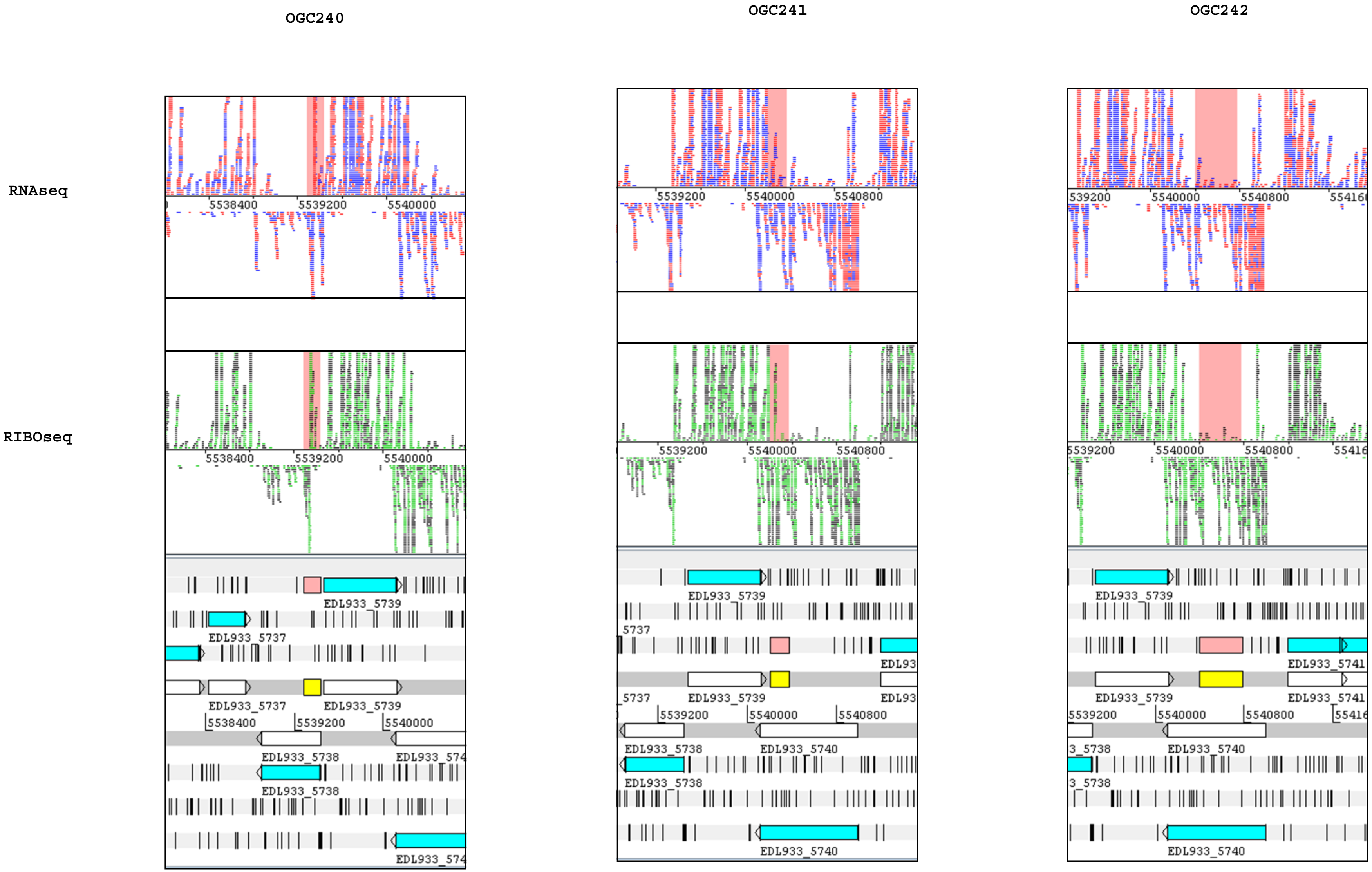
OGC240
OGC241
OGC242
RNAseq
RIBOseq
5538400
5539200
5540000
5540800
EDL933_5737
EDL933_5738
EDL933_5739
EDL933_5740
EDL933_5741
