## Supplementary Figure S2 for "Evidence for Numerous Embedded Antisense Overlapping Genes in Diverse *E. coli* Strains"

**Transcription start sites for the 216 OGCs.** Transcription start sites (TSS) were determined using Cappable-seq. For each OGC, the possible region for a TSS, 250 bp upstream of the start codon, was analyzed. This resulted in 148 TSS of 105 OCGs. Thus, for some OGCs no, one or multiple TSS were identified.

TSS regulation is shown as mean relative read score (RRS, see formula) for three biological replicates each and their standard deviations. The position $X$ is given in each panel for the respective TSS and corresponds to the genome annotation file CP008957.

$$\text{RSS} = \frac{\Sigma(reads\ mapping\ at\ strand{-}specific\ position\ X)}{\Sigma(mapped\ reads\ per\ condition)} \times 1{,}000{,}000$$

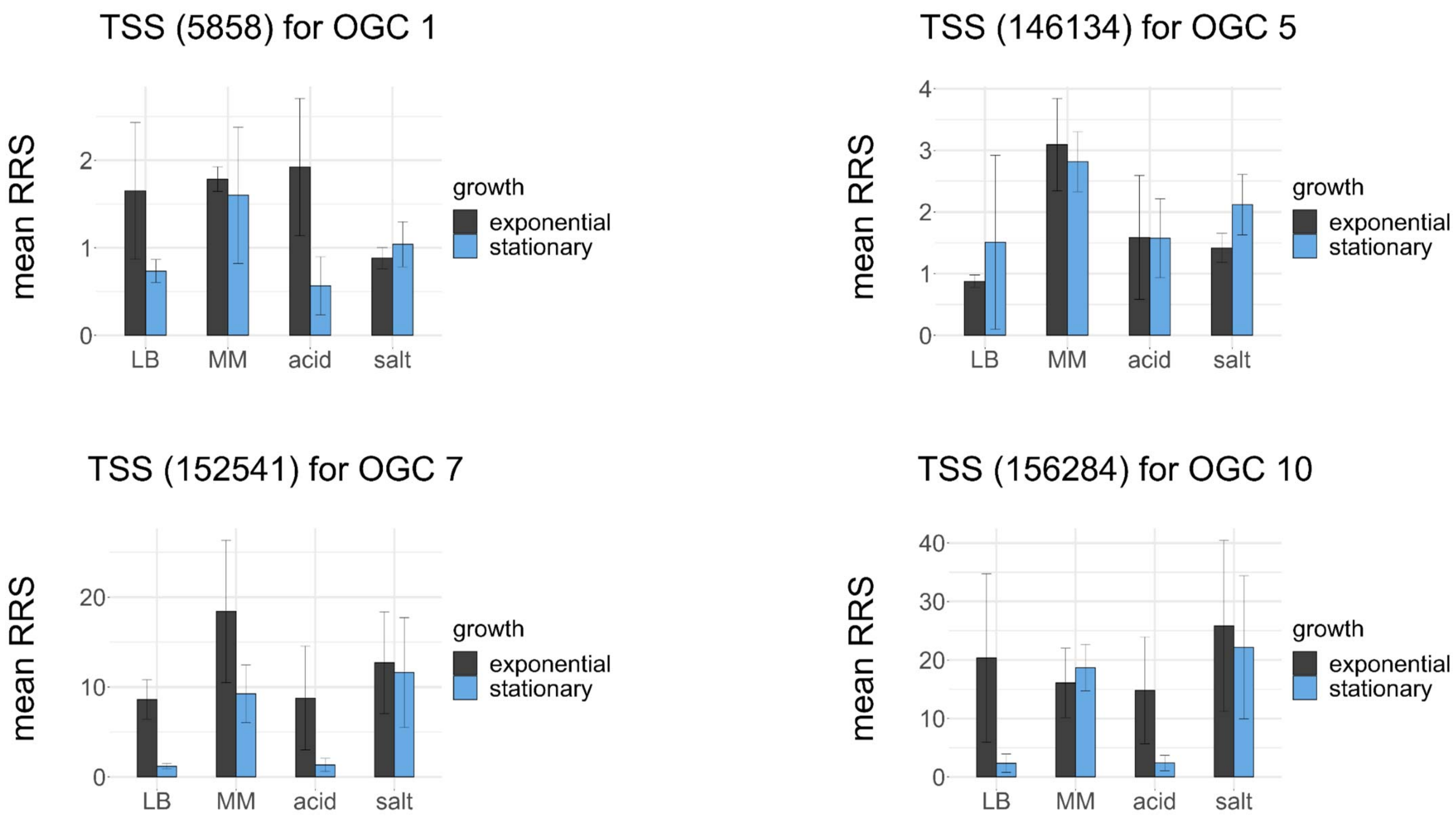
TSS (5858) for OGC 1
mean RRS
0
1
2
LB
MM
acid
salt
growth
exponential
stationary
TSS (146134) for OGC 5
mean RRS
0
1
2
3
4
LB
MM
acid
salt
growth
exponential
stationary
TSS (152541) for OGC 7
mean RRS
0
10
20
LB
MM
acid
salt
growth
exponential
stationary
TSS (156284) for OGC 10
mean RRS
0
10
20
30
40
LB
MM
acid
salt
growth
exponential
stationary

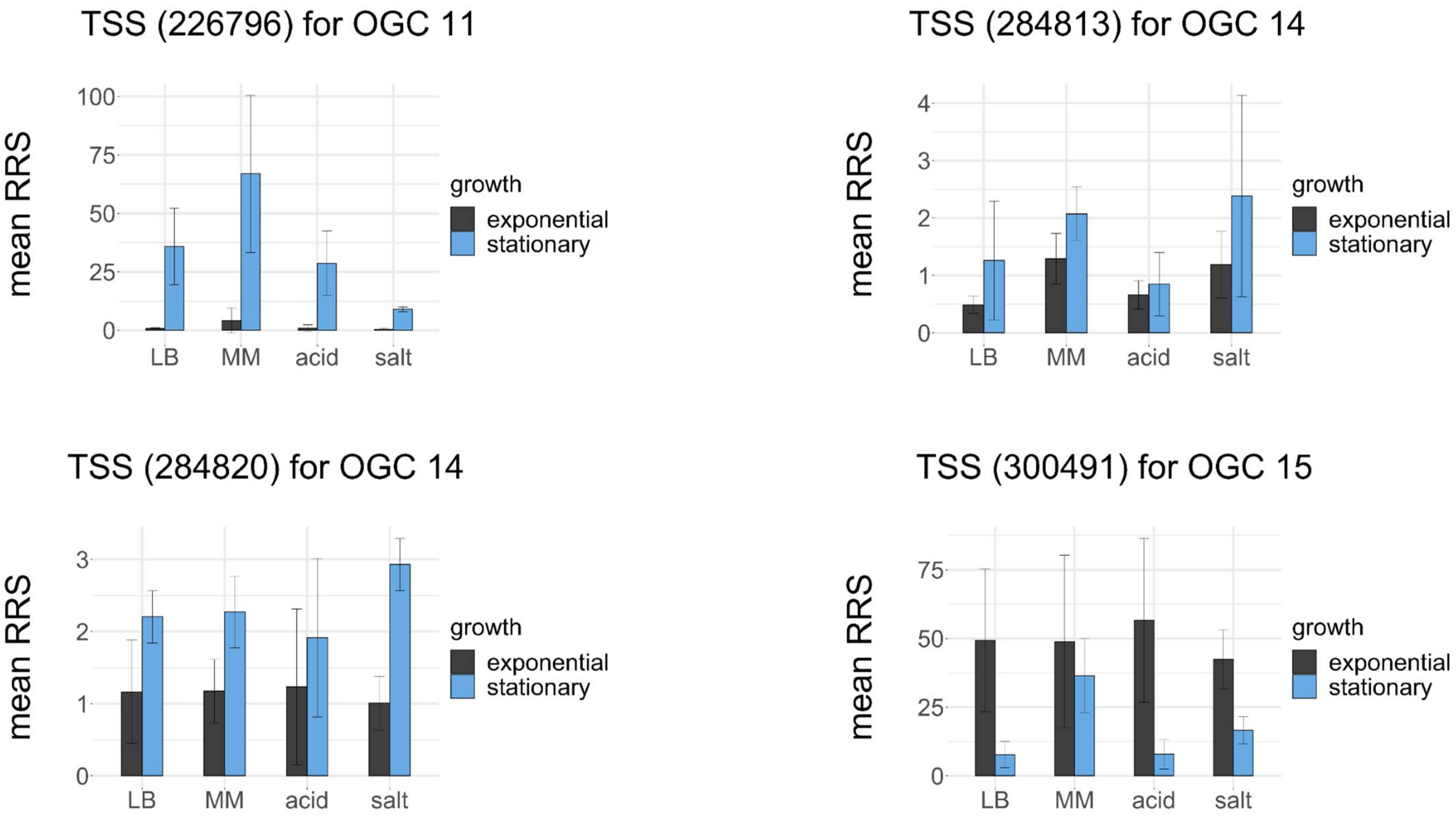
TSS (226796) for OGC 11
TSS (284813) for OGC 14
TSS (284820) for OGC 14
TSS (300491) for OGC 15
mean RRS
growth
exponential
stationary
LB
MM
acid
salt
0
25
50
75
100
1
2
3
4

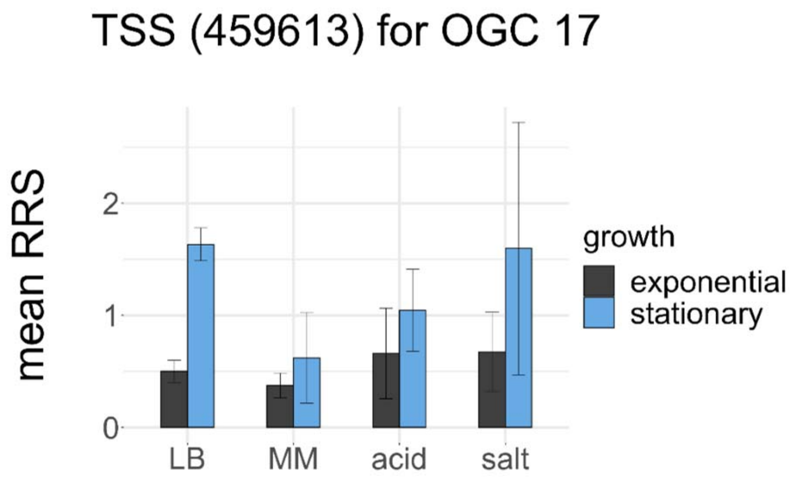
TSS (459613) for OGC 17
mean RRS
0
1
2
LB
MM
acid
salt
growth
exponential
stationary

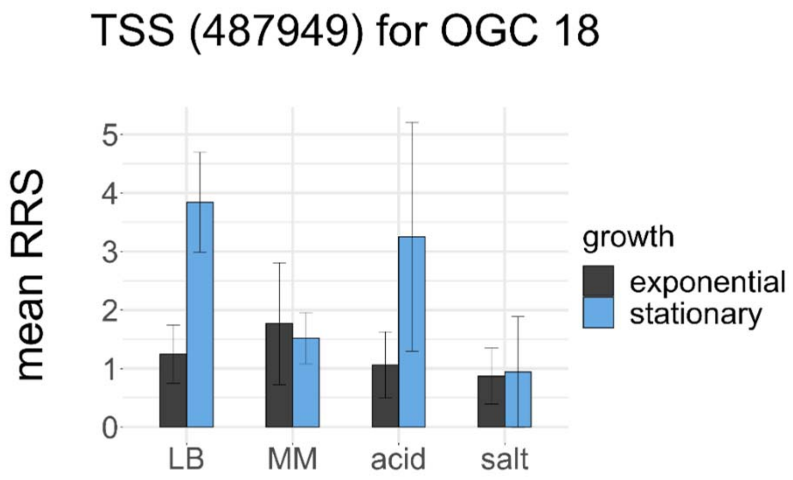
TSS (487949) for OGC 18
mean RRS
0
1
2
3
4
5
LB
MM
acid
salt
growth
exponential
stationary

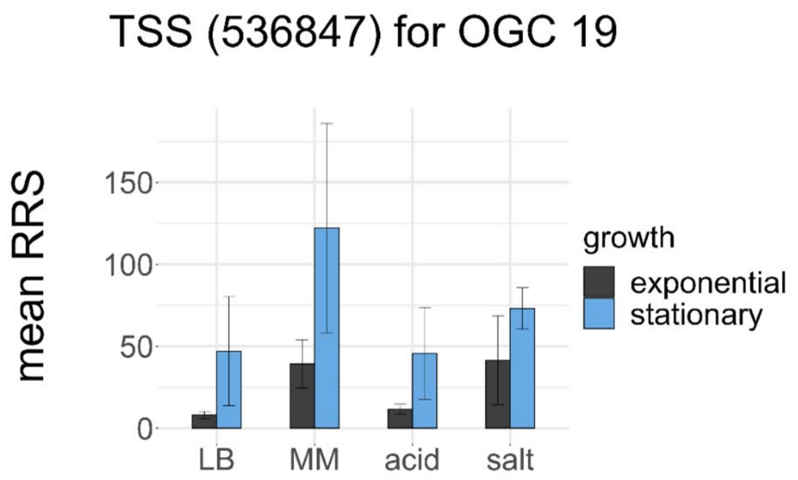
TSS (536847) for OGC 19
mean RRS
0
50
100
150
LB
MM
acid
salt
growth
exponential
stationary

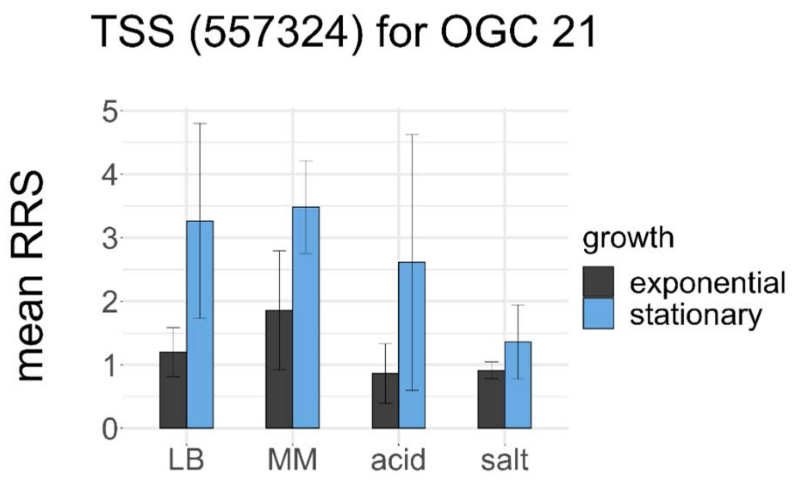
TSS (557324) for OGC 21
mean RRS
0
1
2
3
4
5
LB
MM
acid
salt
growth
exponential
stationary

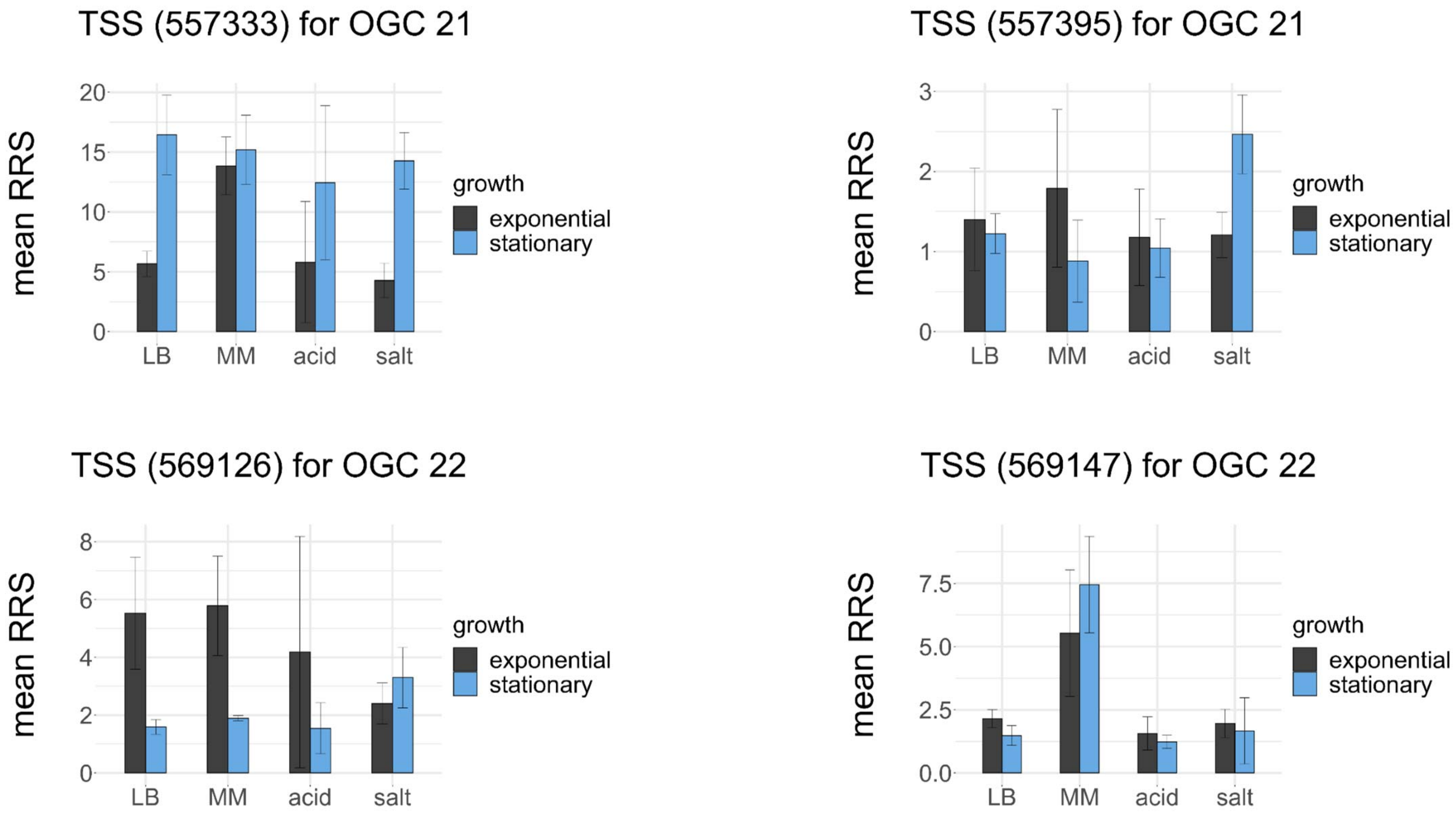

TSS (557333) for OGC 21
TSS (557395) for OGC 21
TSS (569126) for OGC 22
TSS (569147) for OGC 22
mean RRS
growth
exponential
stationary
LB
MM
acid
salt
0
5
10
15
20
1
2
3
4
6
8
0.0
2.5
5.0
7.5

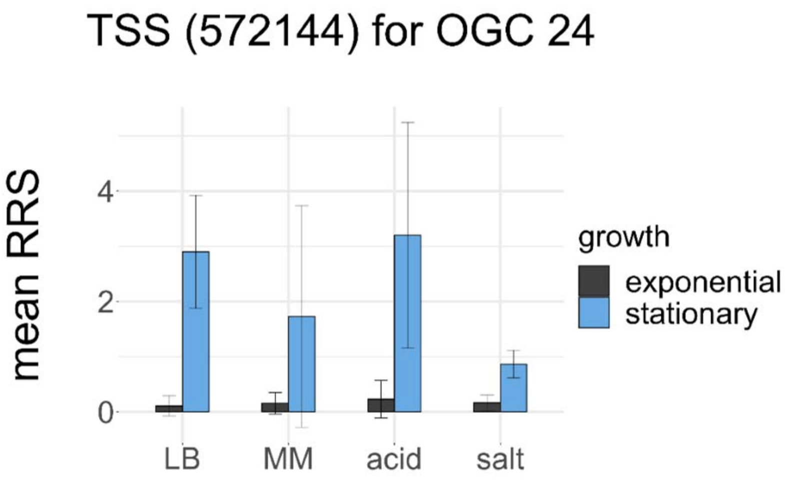
TSS (572144) for OGC 24
mean RRS
0
2
4
LB
MM
acid
salt
growth
exponential
stationary

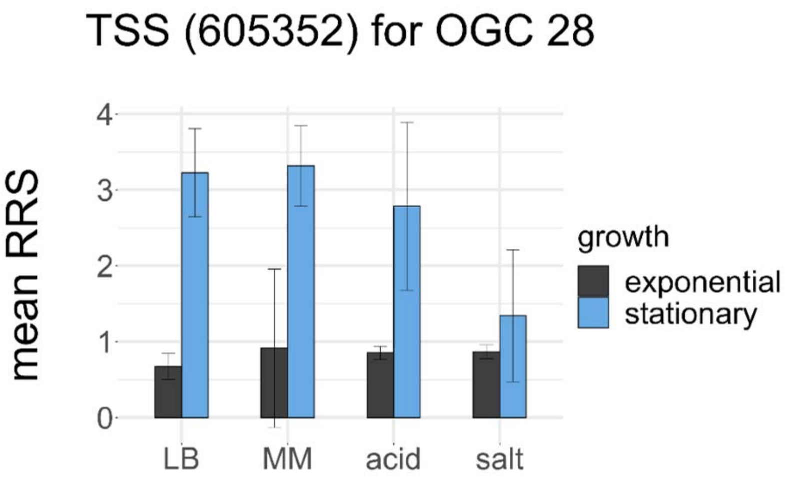
TSS (605352) for OGC 28
mean RRS
0
1
2
3
4
LB
MM
acid
salt
growth
exponential
stationary

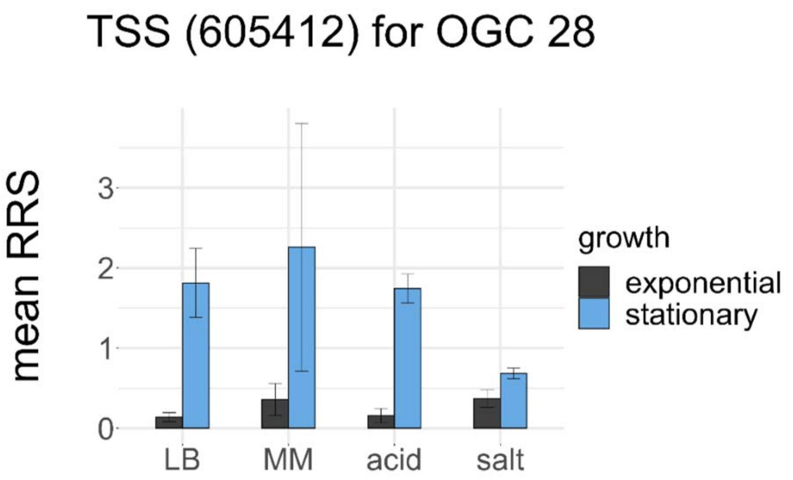
TSS (605412) for OGC 28
mean RRS
0
1
2
3
LB
MM
acid
salt
growth
exponential
stationary

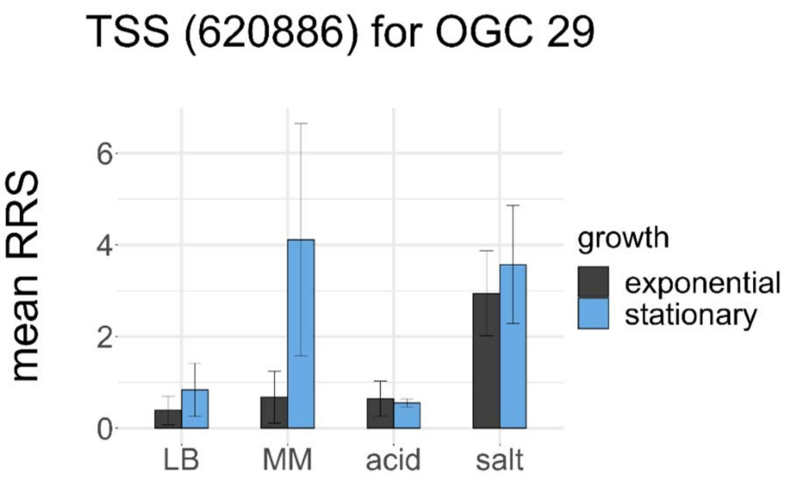
TSS (620886) for OGC 29
mean RRS
0
2
4
6
LB
MM
acid
salt
growth
exponential
stationary

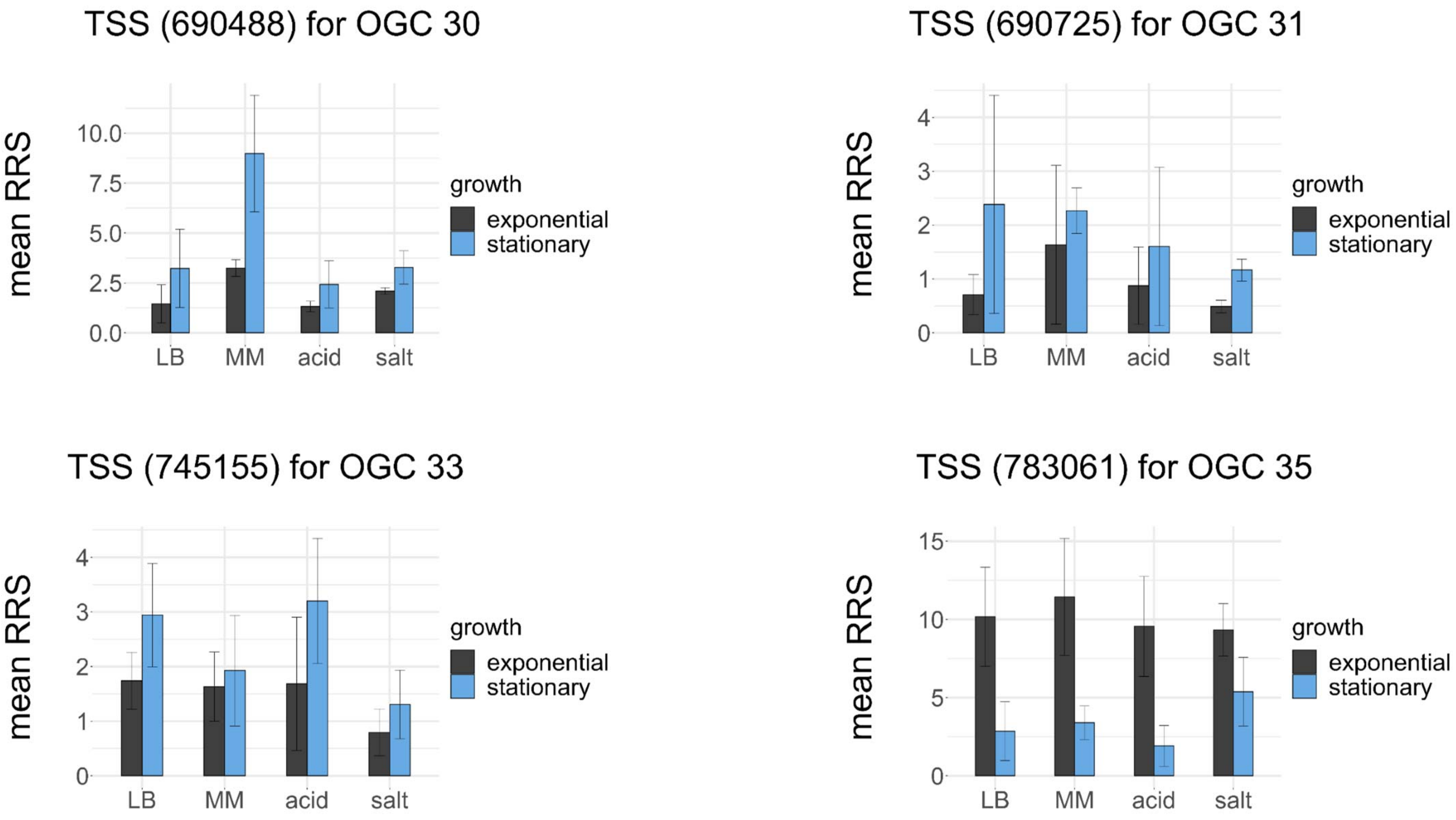

TSS (690488) for OGC 30
mean RRS
10.0
7.5
5.0
2.5
0.0
LB
MM
acid
salt
growth
exponential
stationary
TSS (690725) for OGC 31
mean RRS
4
3
2
1
0
LB
MM
acid
salt
growth
exponential
stationary
TSS (745155) for OGC 33
mean RRS
4
3
2
1
0
LB
MM
acid
salt
growth
exponential
stationary
TSS (783061) for OGC 35
mean RRS
15
10
5
0
LB
MM
acid
salt
growth
exponential
stationary

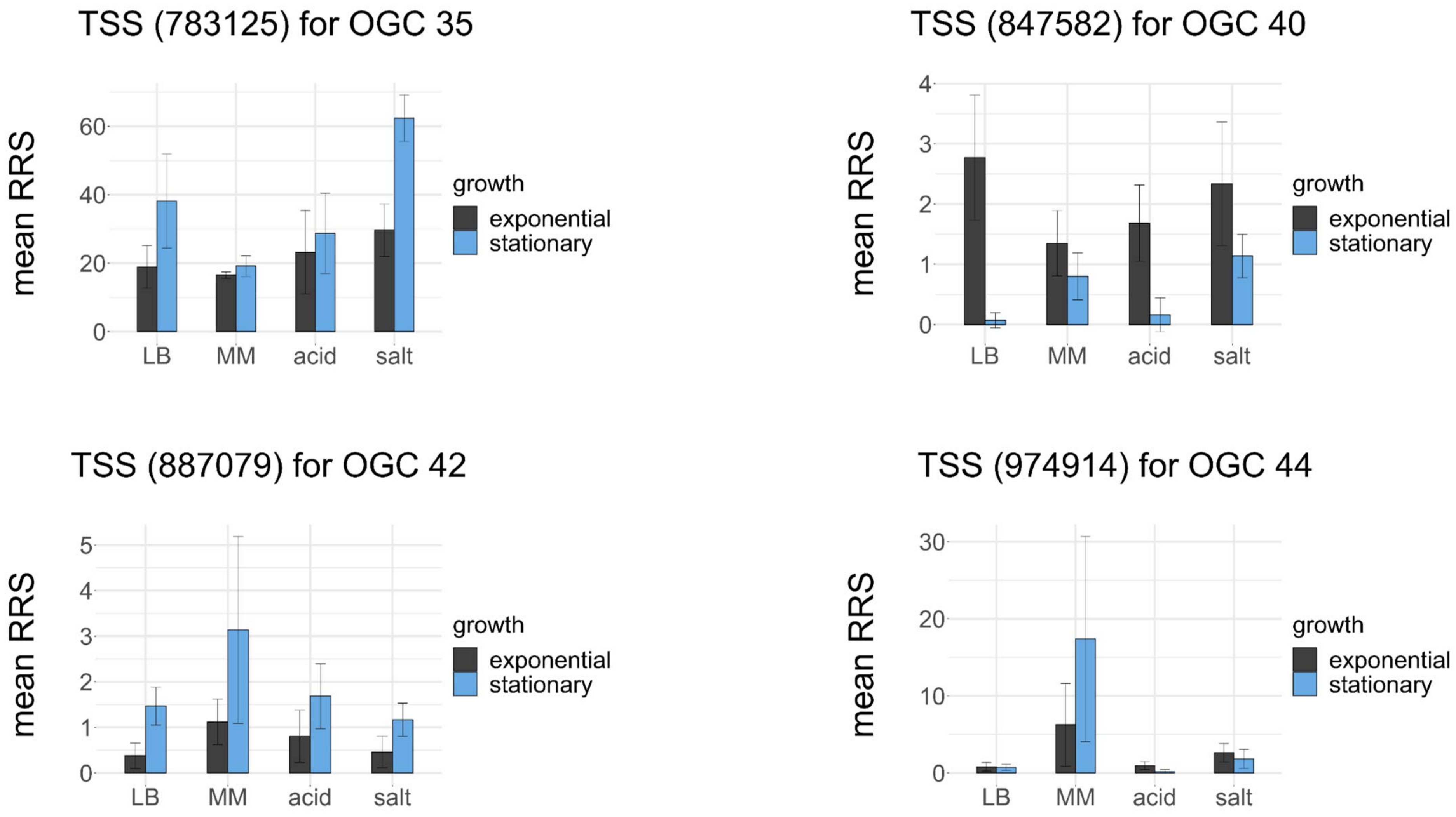

TSS (783125) for OGC 35
mean RRS
0
20
40
60
LB
MM
acid
salt
growth
exponential
stationary
TSS (847582) for OGC 40
mean RRS
0
1
2
3
4
LB
MM
acid
salt
growth
exponential
stationary
TSS (887079) for OGC 42
mean RRS
0
1
2
3
4
5
LB
MM
acid
salt
growth
exponential
stationary
TSS (974914) for OGC 44
mean RRS
0
10
20
30
LB
MM
acid
salt
growth
exponential
stationary

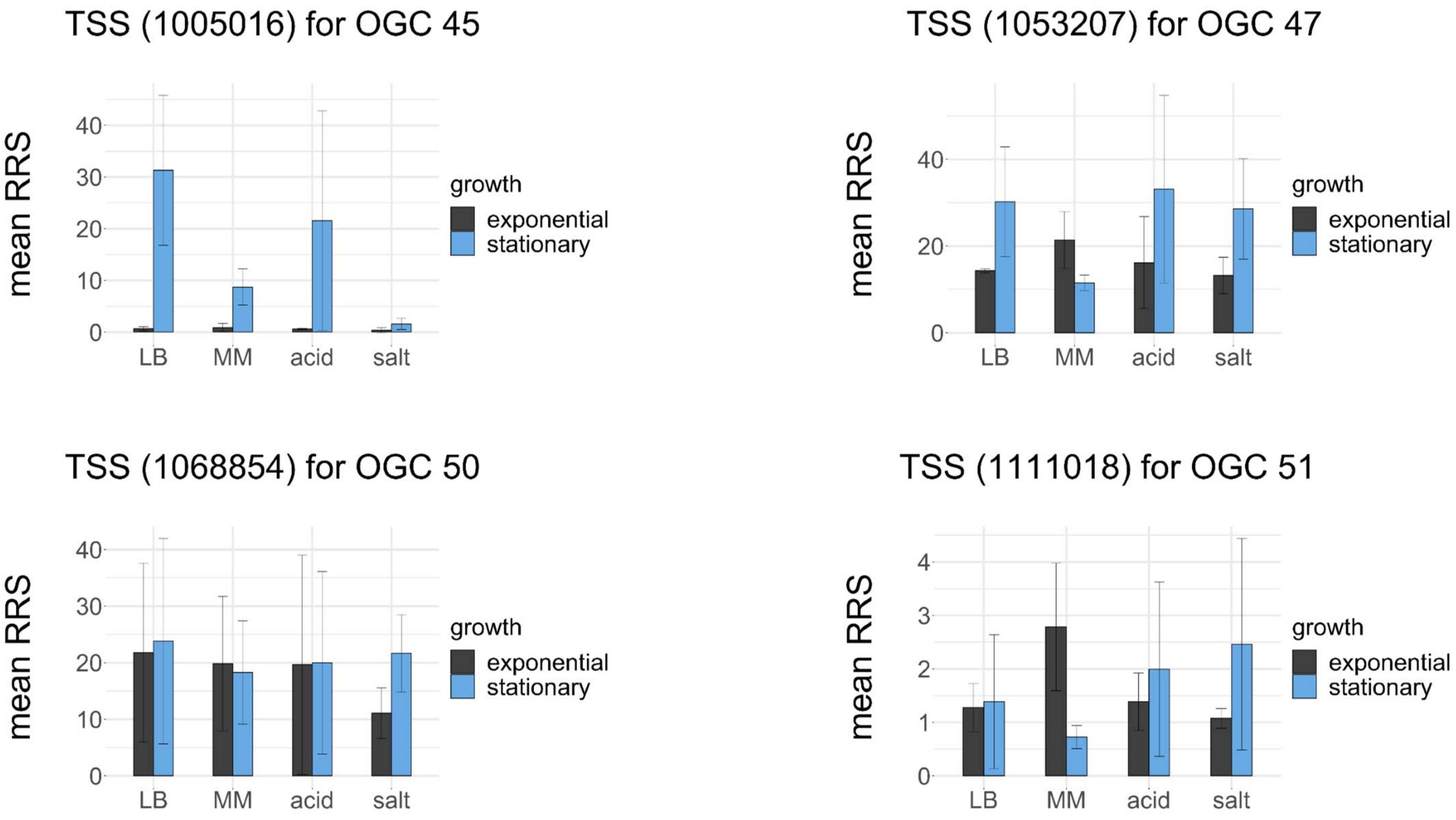
TSS (1005016) for OGC 45
TSS (1053207) for OGC 47
TSS (1068854) for OGC 50
TSS (1111018) for OGC 51
mean RRS
LB
MM
acid
salt
growth
exponential
stationary

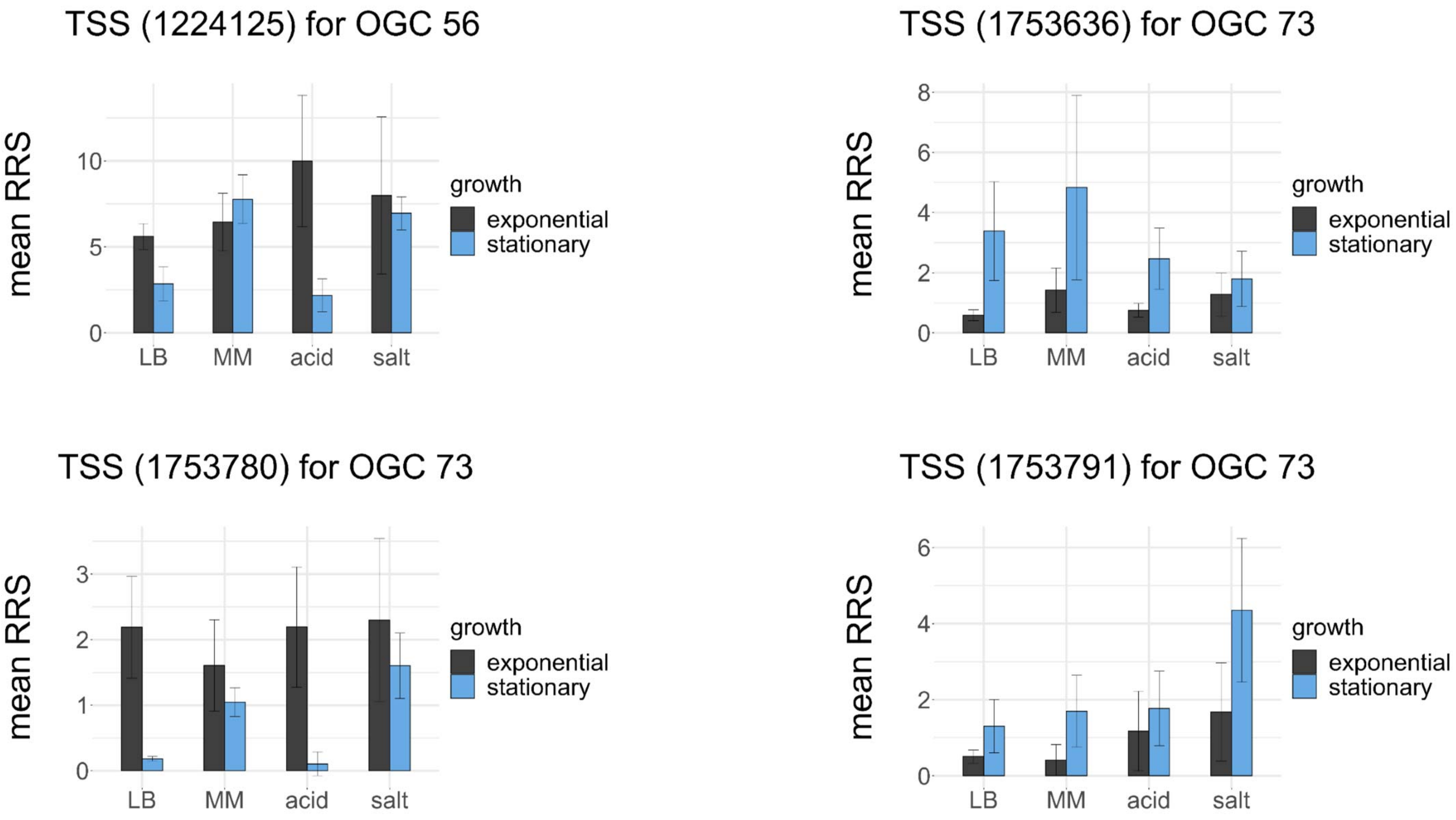
TSS (1224125) for OGC 56
mean RRS
0
5
10
LB
MM
acid
salt
growth
exponential
stationary
TSS (1753636) for OGC 73
mean RRS
0
2
4
6
8
LB
MM
acid
salt
growth
exponential
stationary
TSS (1753780) for OGC 73
mean RRS
0
1
2
3
LB
MM
acid
salt
growth
exponential
stationary
TSS (1753791) for OGC 73
mean RRS
0
2
4
6
LB
MM
acid
salt
growth
exponential
stationary

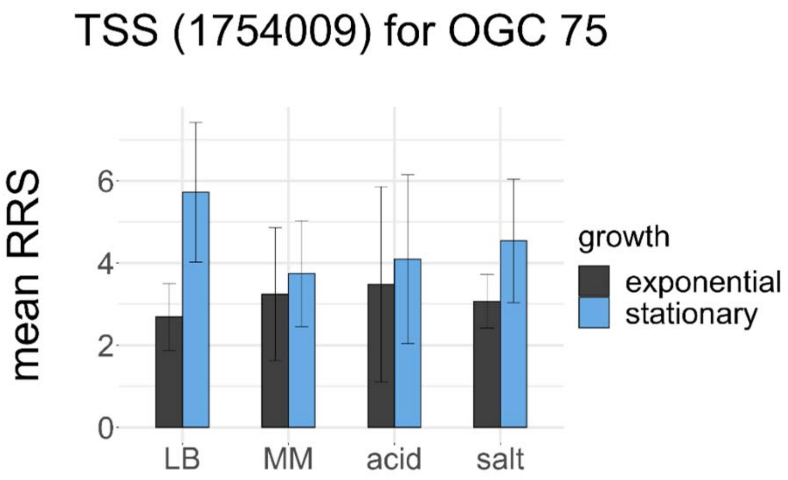
TSS (1754009) for OGC 75
mean RRS
0
2
4
6
LB
MM
acid
salt
growth
exponential
stationary

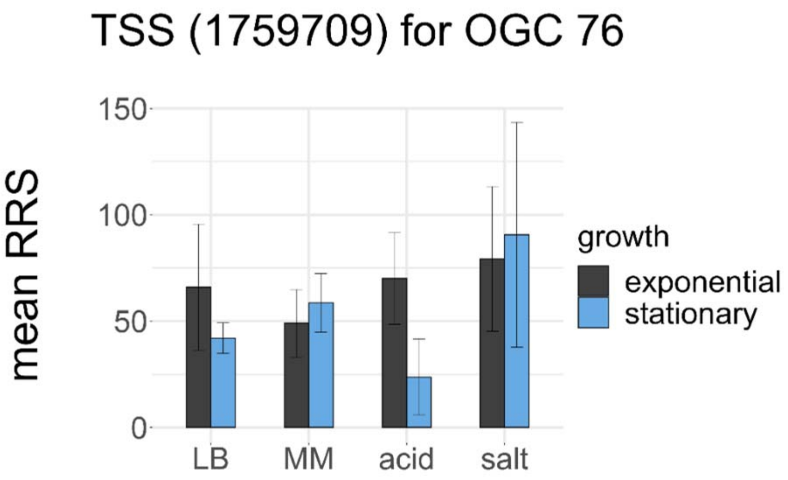
TSS (1759709) for OGC 76
mean RRS
0
50
100
150
LB
MM
acid
salt
growth
exponential
stationary

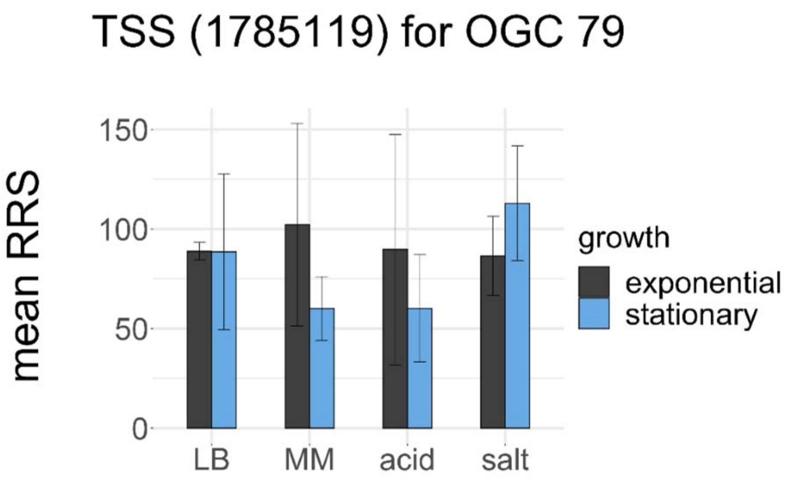
TSS (1785119) for OGC 79
mean RRS
0
50
100
150
LB
MM
acid
salt
growth
exponential
stationary

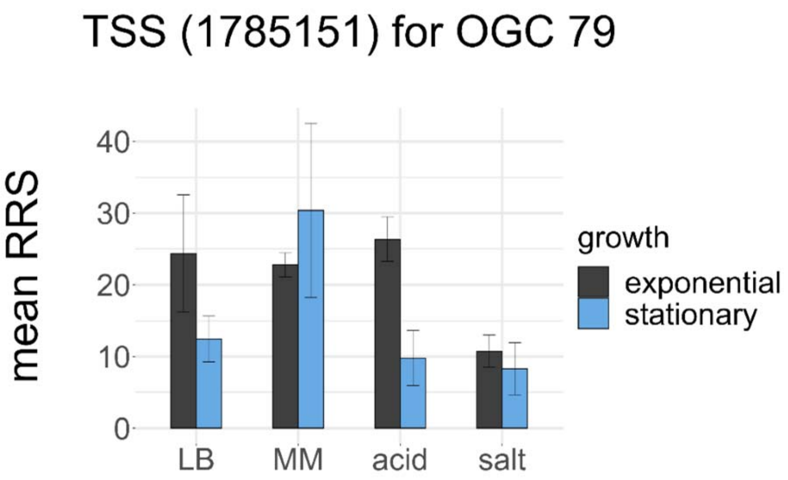
TSS (1785151) for OGC 79
mean RRS
0
10
20
30
40
LB
MM
acid
salt
growth
exponential
stationary

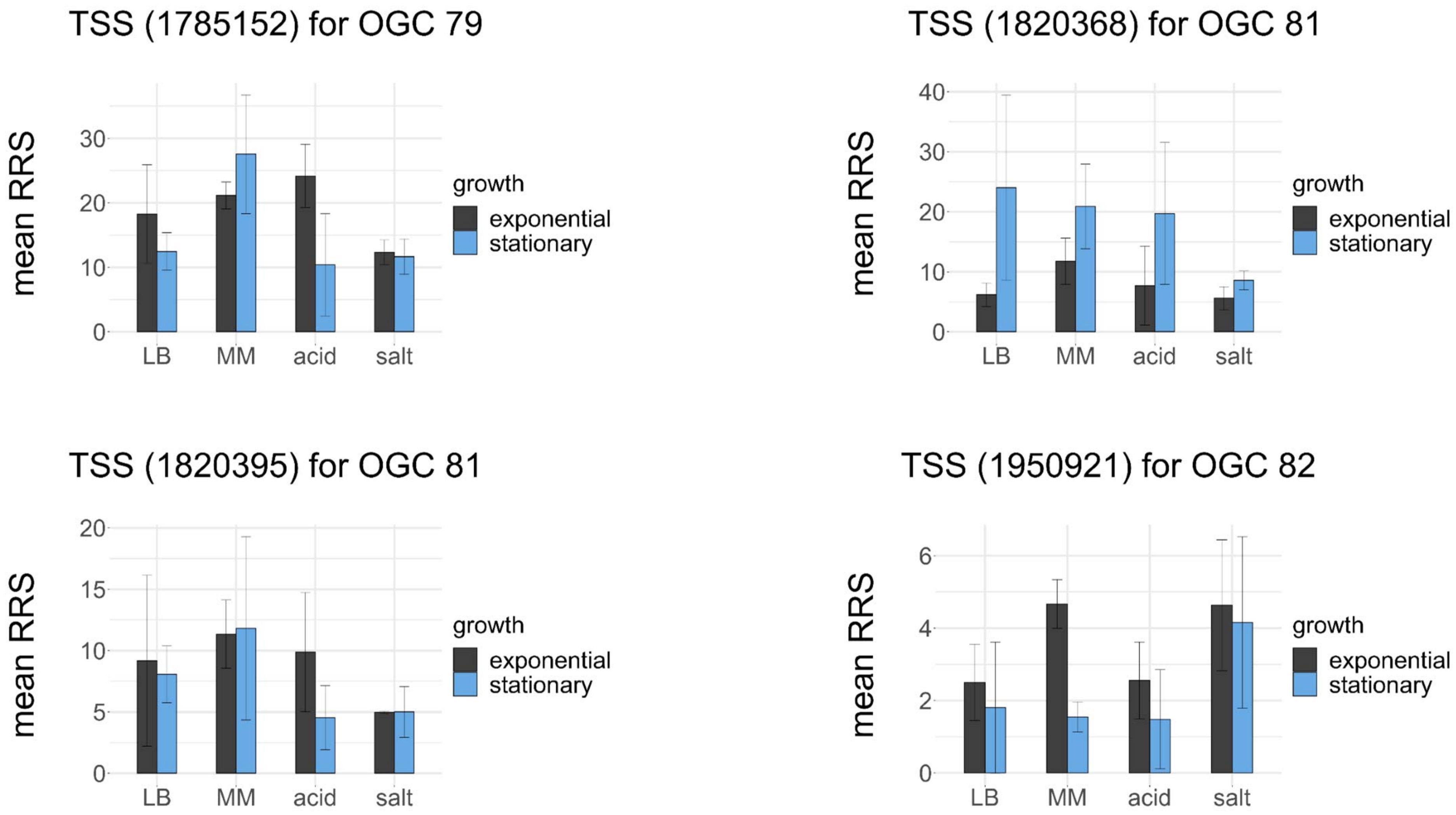
TSS (1785152) for OGC 79
TSS (1820368) for OGC 81
TSS (1820395) for OGC 81
TSS (1950921) for OGC 82
mean RRS
growth
exponential
stationary
LB
MM
acid
salt

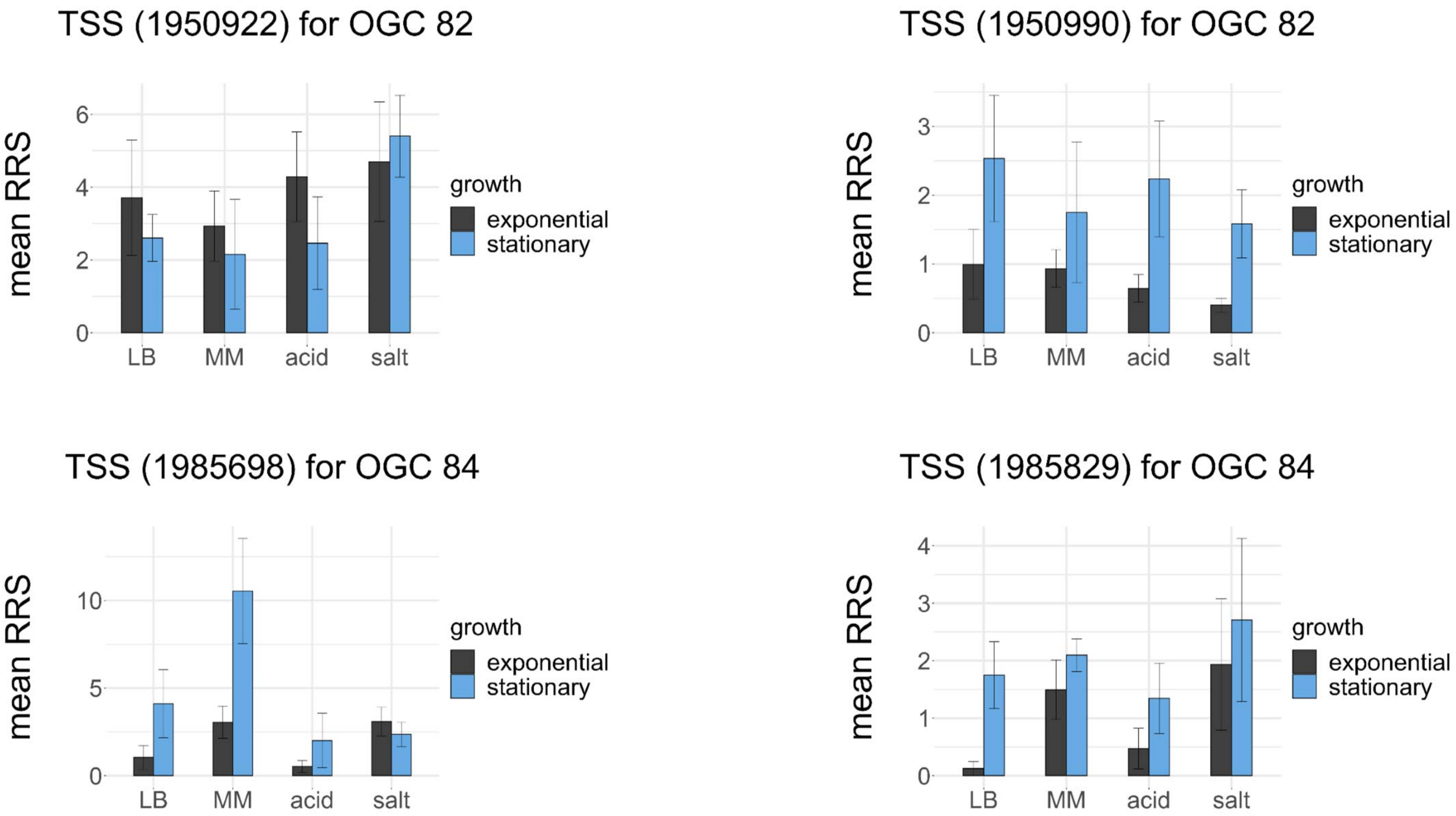

TSS (1950922) for OGC 82
mean RRS
0
2
4
6
LB
MM
acid
salt
growth
exponential
stationary
TSS (1950990) for OGC 82
mean RRS
0
1
2
3
LB
MM
acid
salt
growth
exponential
stationary
TSS (1985698) for OGC 84
mean RRS
0
5
10
LB
MM
acid
salt
growth
exponential
stationary
TSS (1985829) for OGC 84
mean RRS
0
1
2
3
4
LB
MM
acid
salt
growth
exponential
stationary

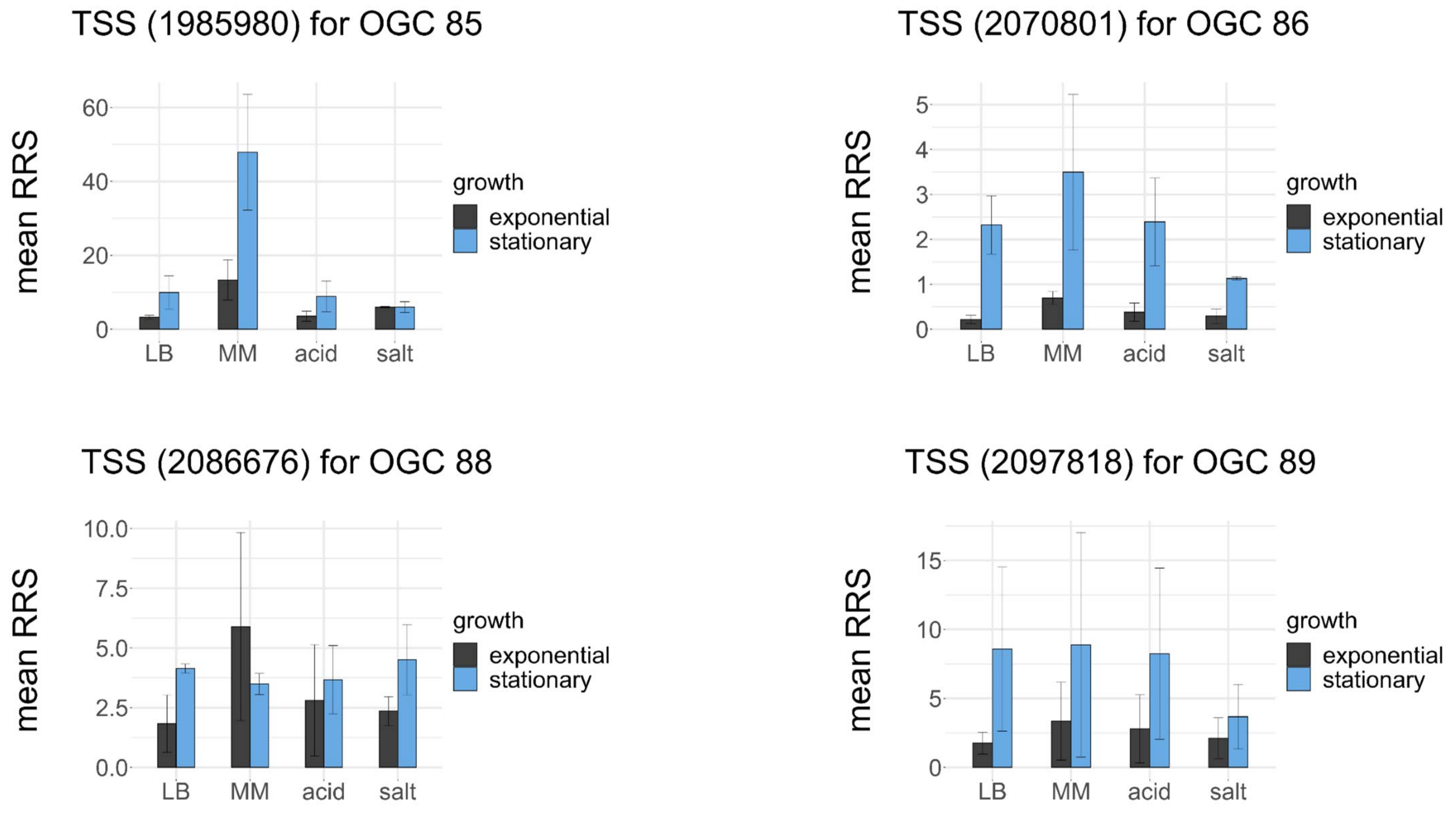

TSS (1985980) for OGC 85
TSS (2070801) for OGC 86
TSS (2086676) for OGC 88
TSS (2097818) for OGC 89
mean RRS
growth
exponential
stationary
LB
MM
acid
salt

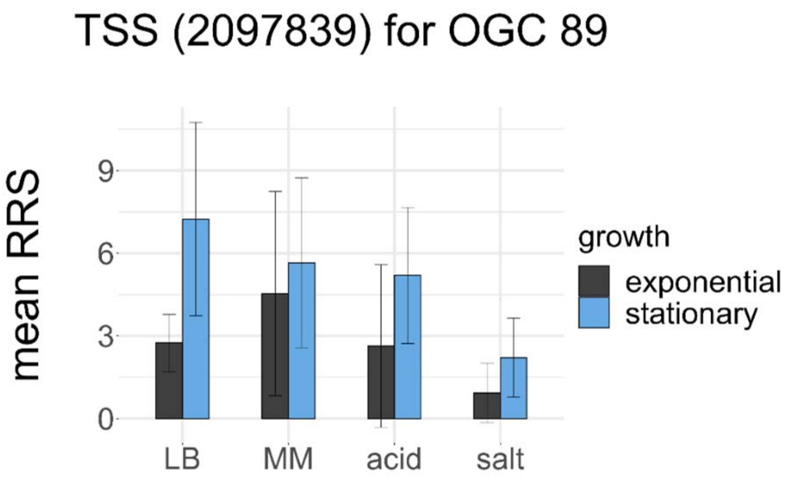
TSS (2097839) for OGC 89
mean RRS
0
3
6
9
LB
MM
acid
salt
growth
exponential
stationary

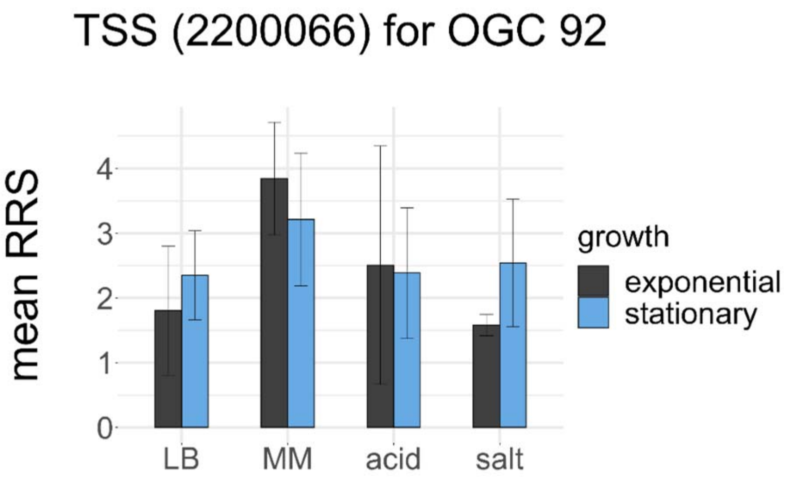
TSS (2200066) for OGC 92
mean RRS
0
1
2
3
4
LB
MM
acid
salt
growth
exponential
stationary

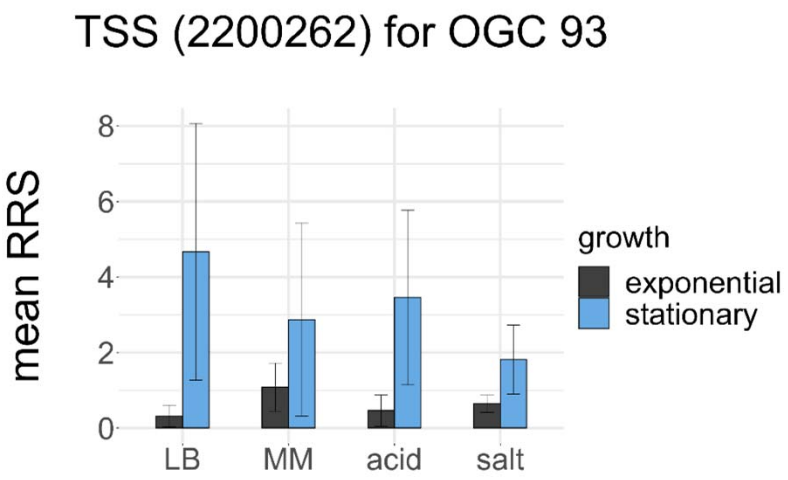
TSS (2200262) for OGC 93
mean RRS
0
2
4
6
8
LB
MM
acid
salt
growth
exponential
stationary

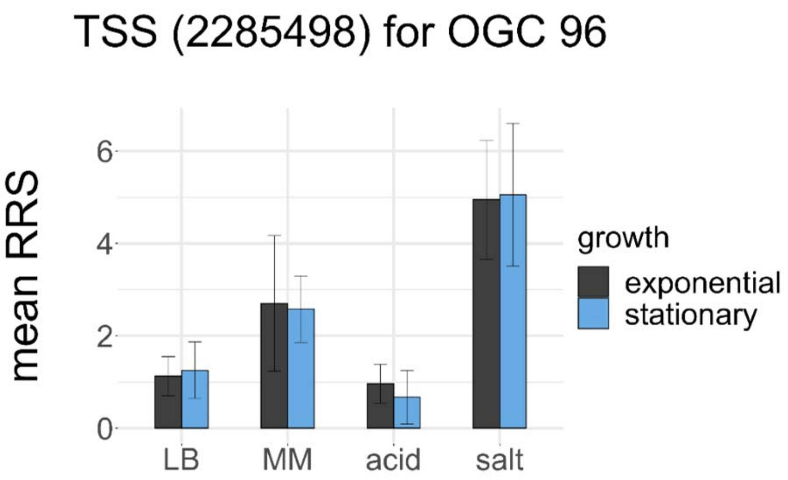
TSS (2285498) for OGC 96
mean RRS
0
2
4
6
LB
MM
acid
salt
growth
exponential
stationary

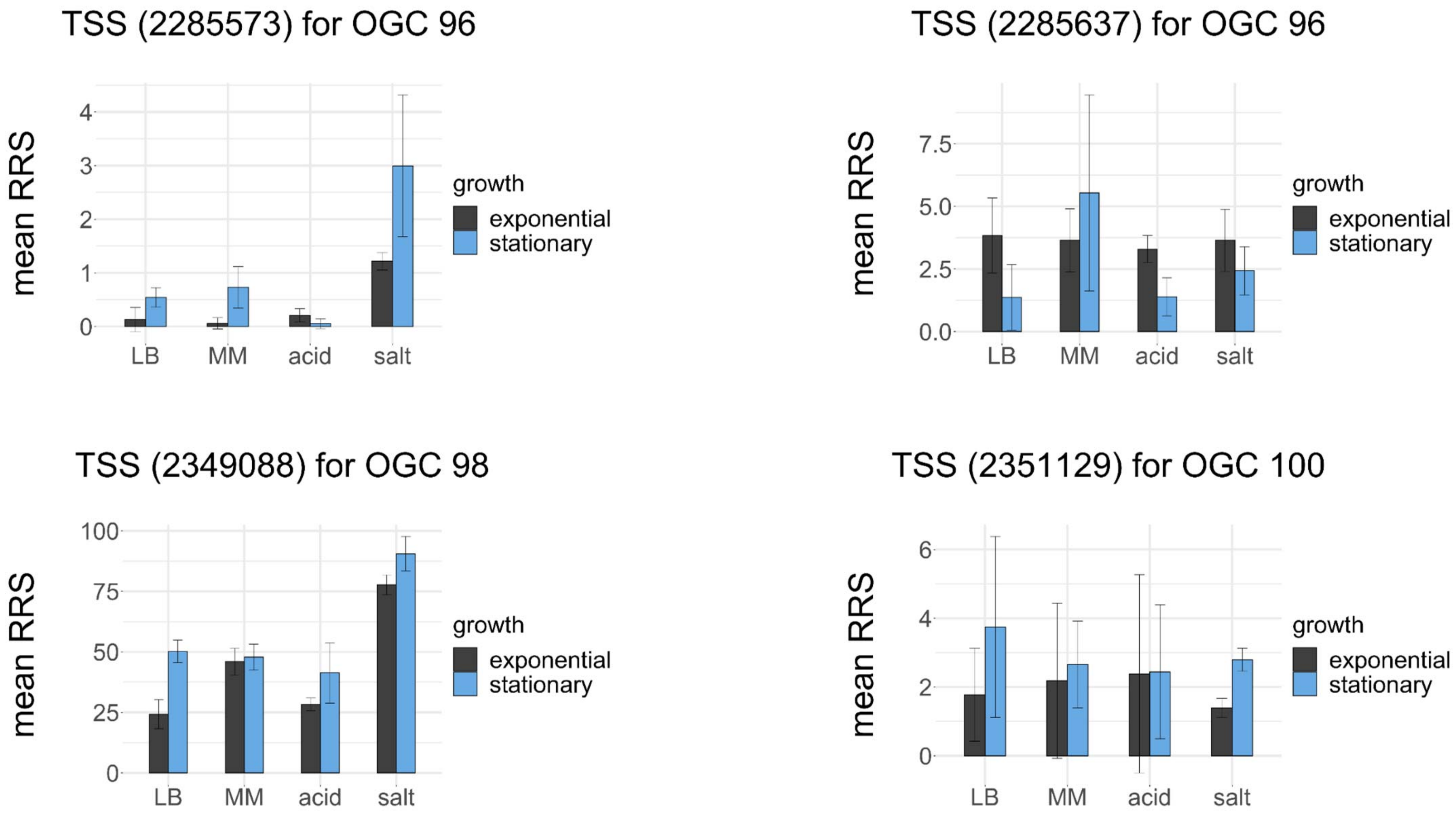

TSS (2285573) for OGC 96
TSS (2285637) for OGC 96
TSS (2349088) for OGC 98
TSS (2351129) for OGC 100
mean RRS
growth
exponential
stationary
LB
MM
acid
salt

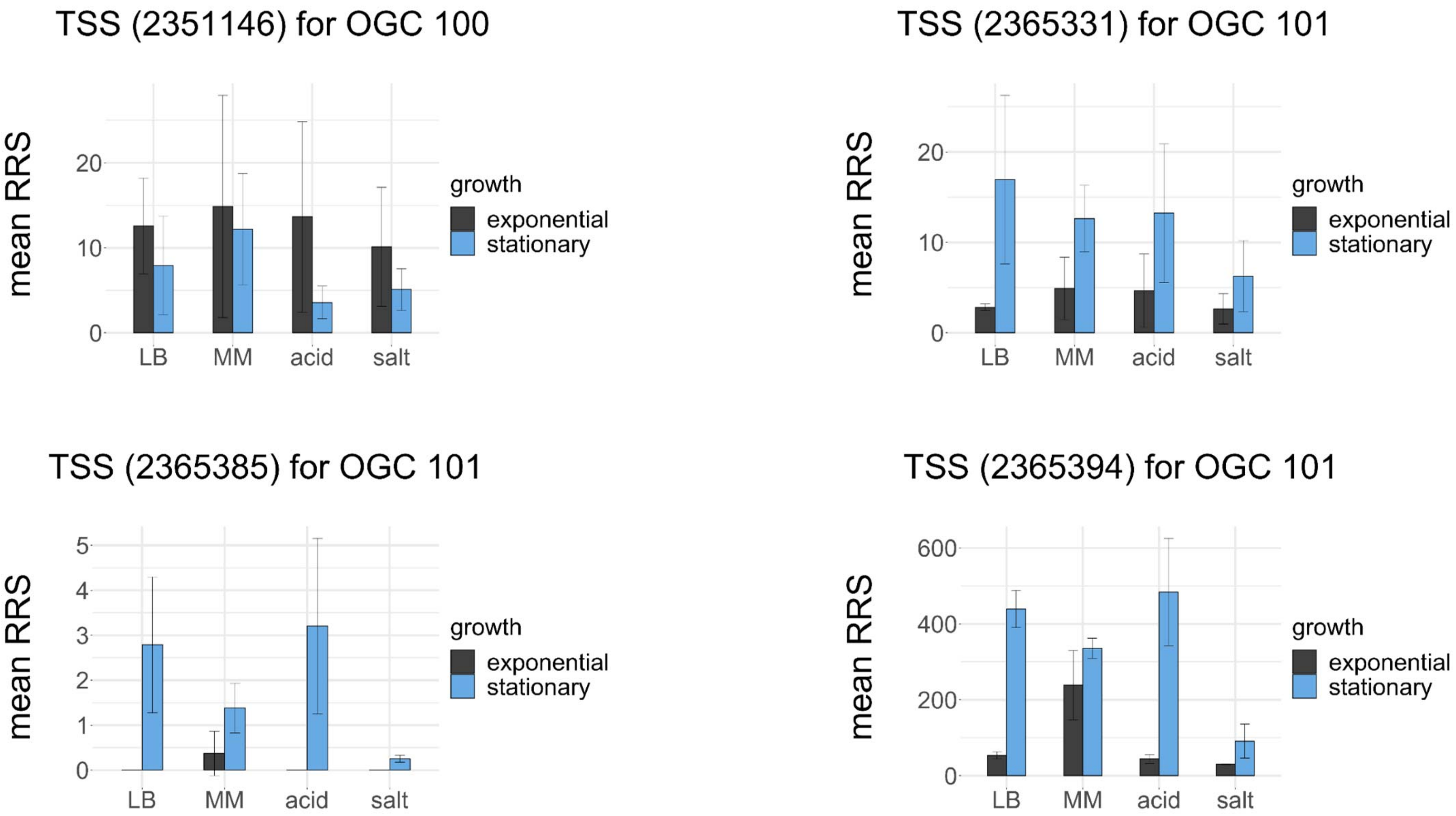

TSS (2351146) for OGC 100
TSS (2365331) for OGC 101
TSS (2365385) for OGC 101
TSS (2365394) for OGC 101
mean RRS
LB
MM
acid
salt
growth
exponential
stationary
0
10
20
1
2
3
4
5
200
400
600

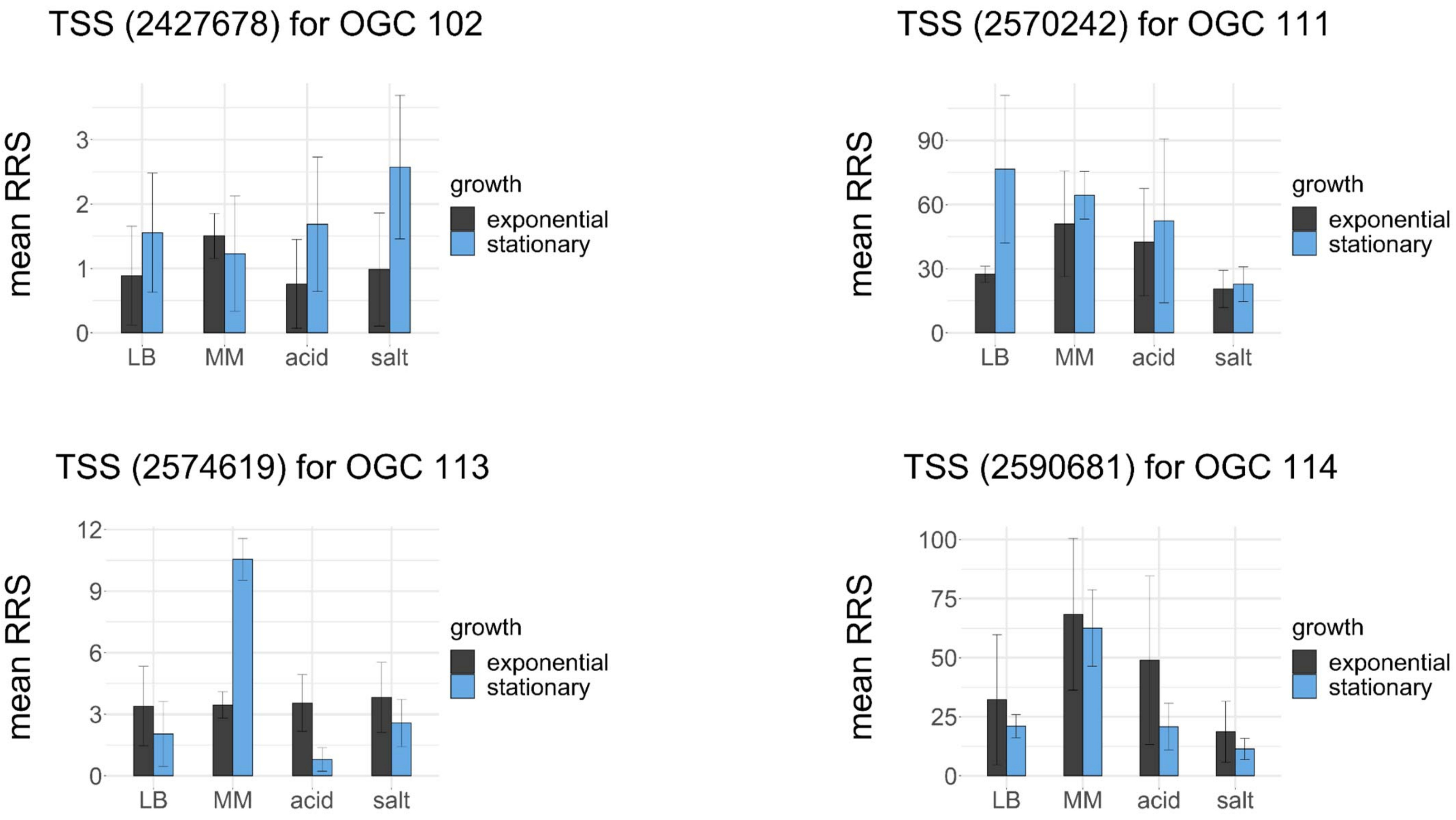

TSS (2427678) for OGC 102
mean RRS
0
1
2
3
LB
MM
acid
salt
growth
exponential
stationary
TSS (2570242) for OGC 111
mean RRS
0
30
60
90
LB
MM
acid
salt
growth
exponential
stationary
TSS (2574619) for OGC 113
mean RRS
0
3
6
9
12
LB
MM
acid
salt
growth
exponential
stationary
TSS (2590681) for OGC 114
mean RRS
0
25
50
75
100
LB
MM
acid
salt
growth
exponential
stationary

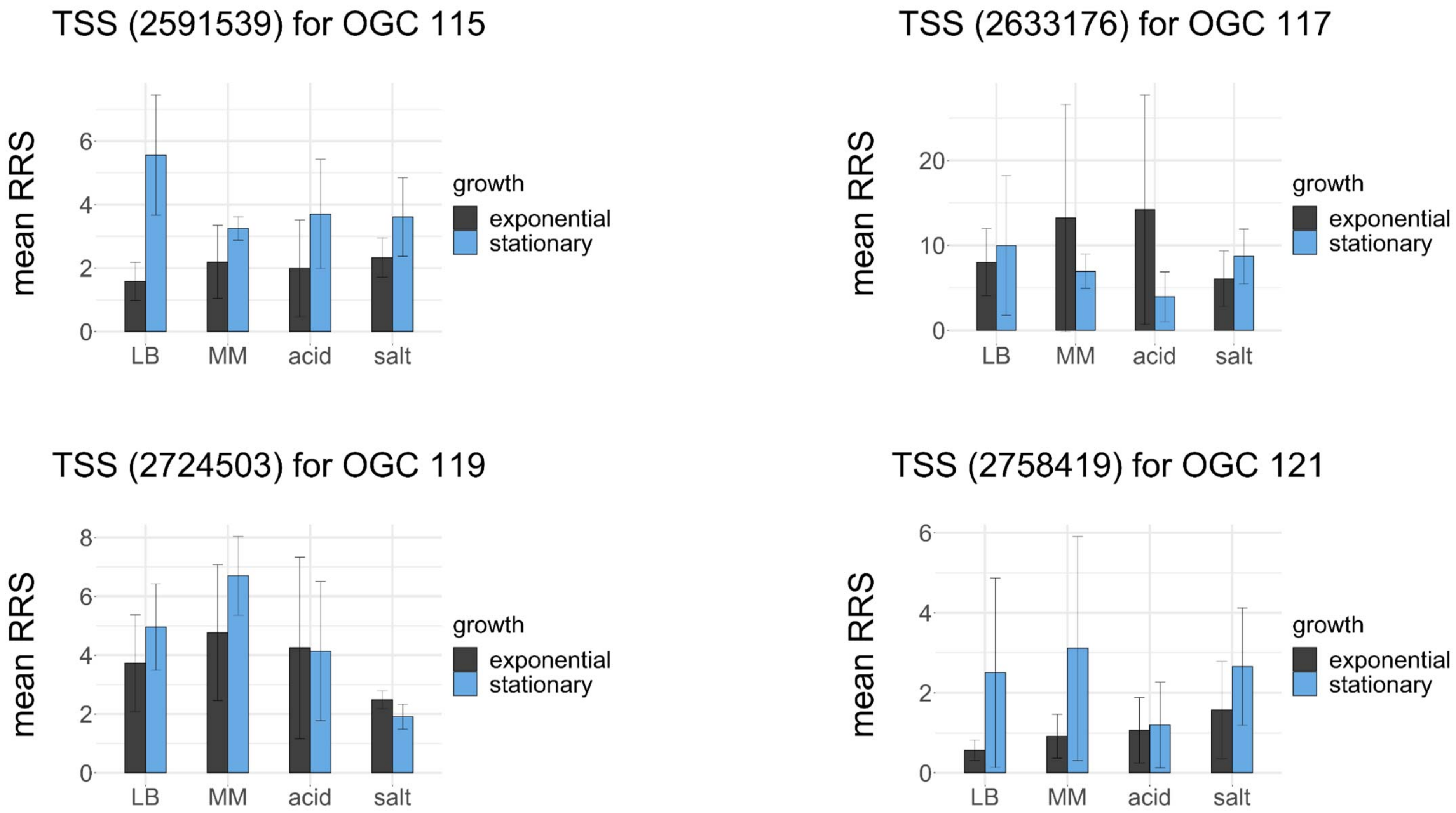
TSS (2591539) for OGC 115
TSS (2633176) for OGC 117
TSS (2724503) for OGC 119
TSS (2758419) for OGC 121
mean RRS
growth
exponential
stationary
LB
MM
acid
salt
0
2
4
6
8
10
20

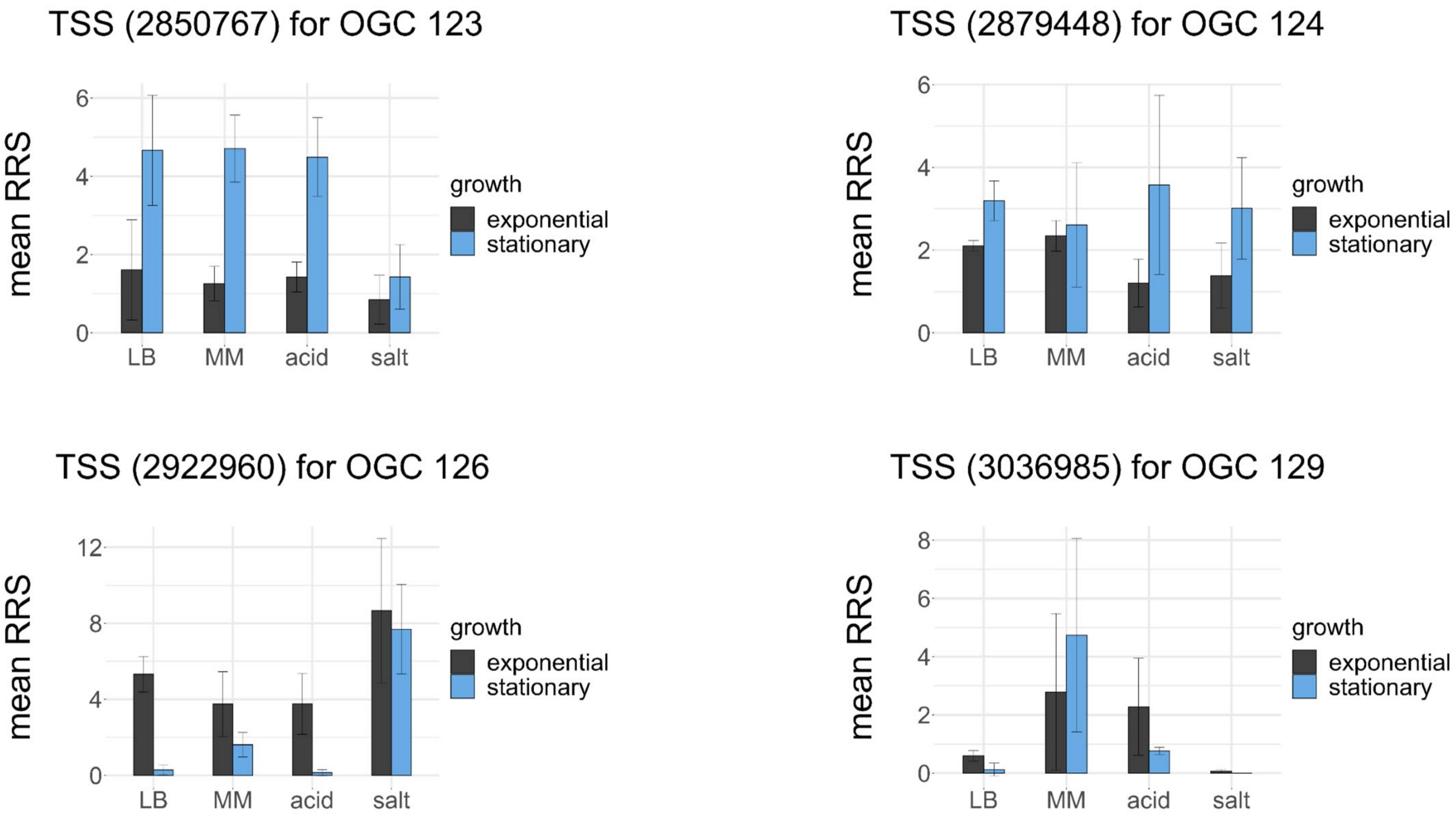
TSS (2850767) for OGC 123
TSS (2879448) for OGC 124
TSS (2922960) for OGC 126
TSS (3036985) for OGC 129
mean RRS
growth
exponential
stationary
LB
MM
acid
salt
0
2
4
6
8
12

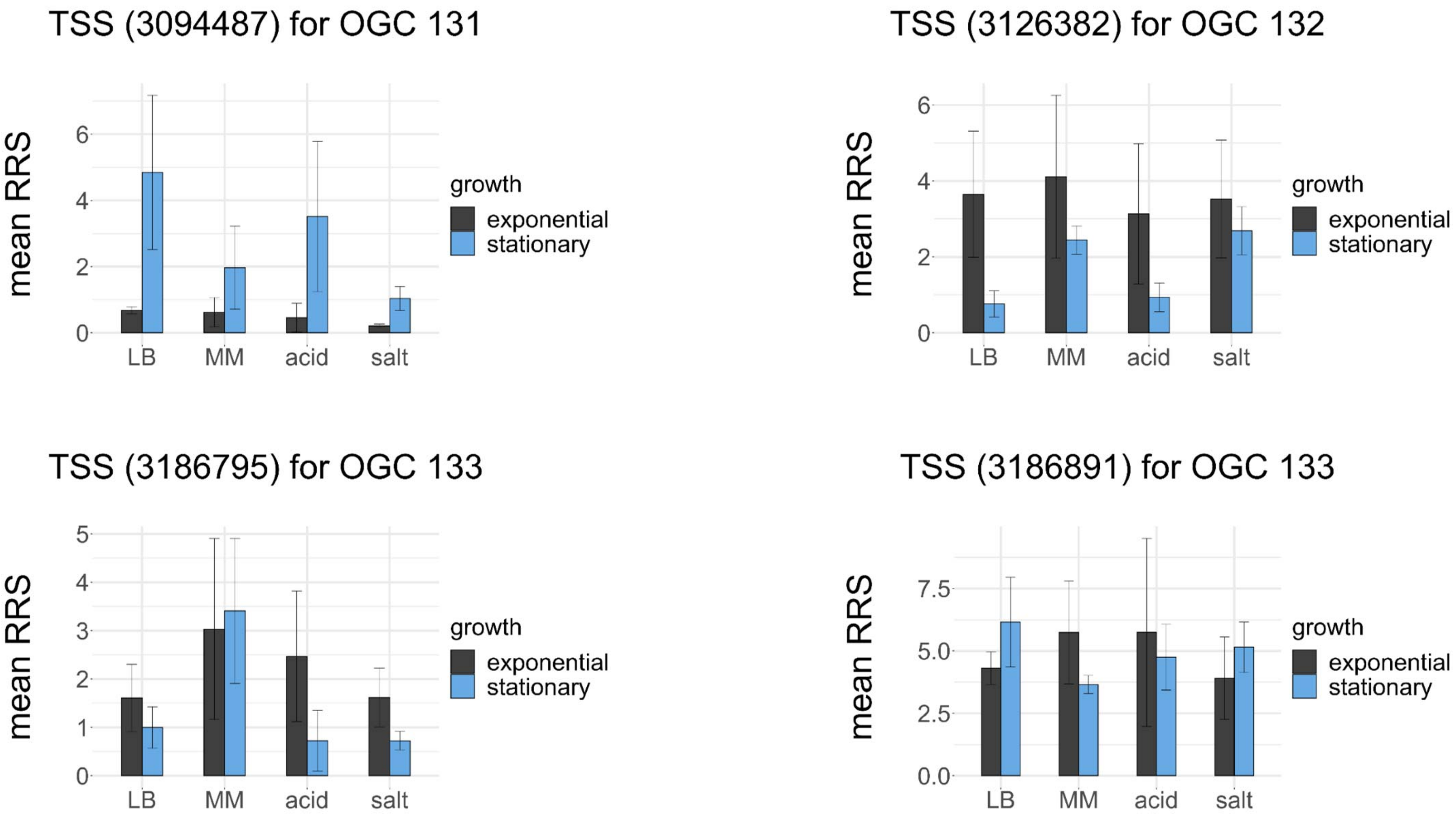
TSS (3094487) for OGC 131
TSS (3126382) for OGC 132
TSS (3186795) for OGC 133
TSS (3186891) for OGC 133
mean RRS
growth
exponential
stationary
LB
MM
acid
salt
0
1
2
2.5
3
4
5
5.0
6
7.5
0.0

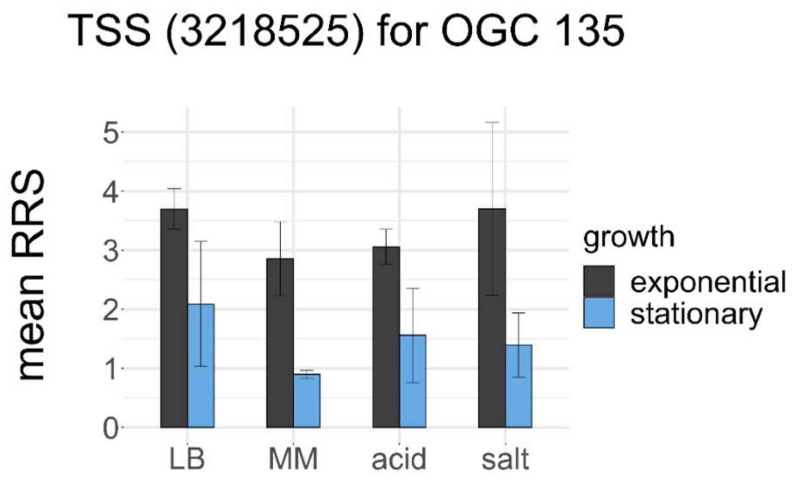
TSS (3218525) for OGC 135
mean RRS
0
1
2
3
4
5
LB
MM
acid
salt
growth
exponential
stationary

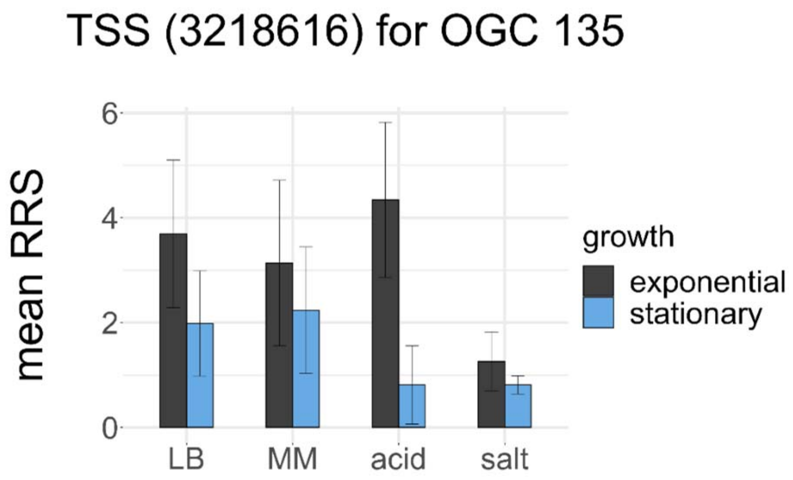
TSS (3218616) for OGC 135
mean RRS
0
2
4
6
LB
MM
acid
salt
growth
exponential
stationary

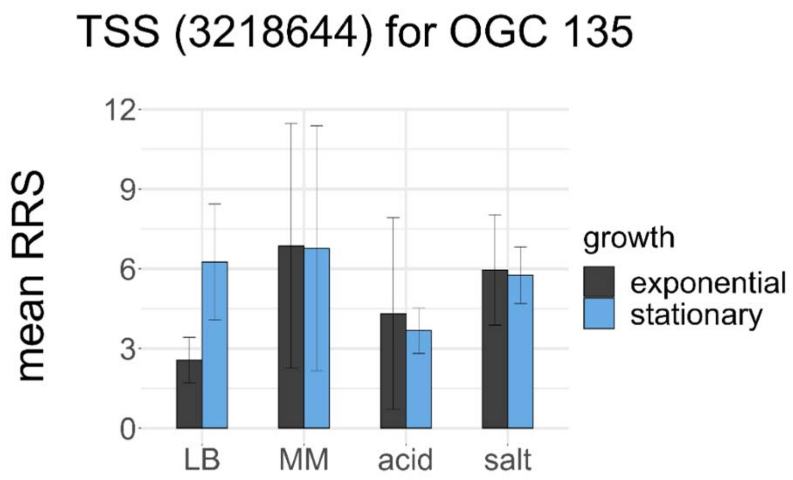
TSS (3218644) for OGC 135
mean RRS
0
3
6
9
12
LB
MM
acid
salt
growth
exponential
stationary

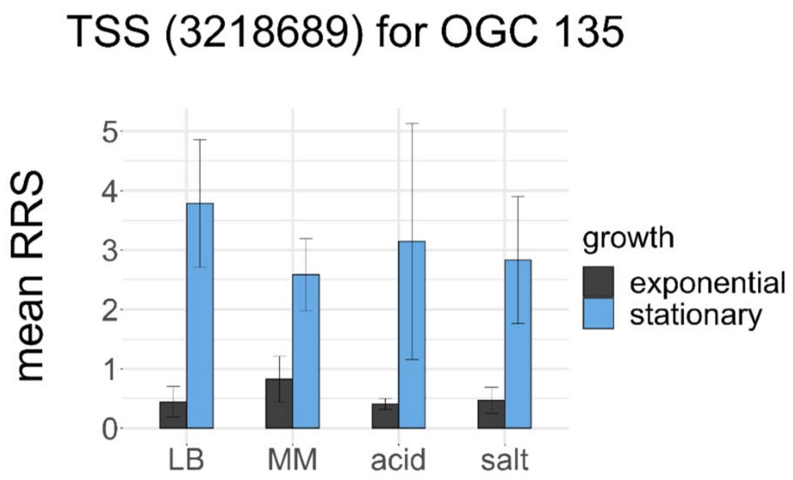
TSS (3218689) for OGC 135
mean RRS
0
1
2
3
4
5
LB
MM
acid
salt
growth
exponential
stationary

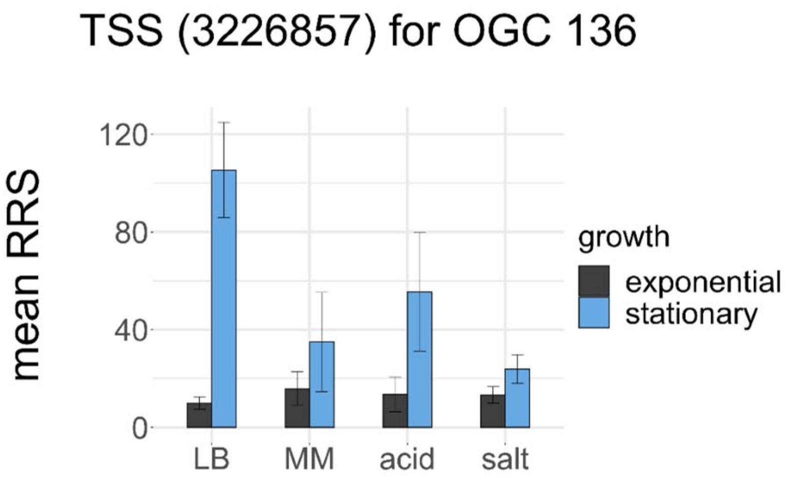
TSS (3226857) for OGC 136
mean RRS
120
80
40
0
LB
MM
acid
salt
growth
exponential
stationary

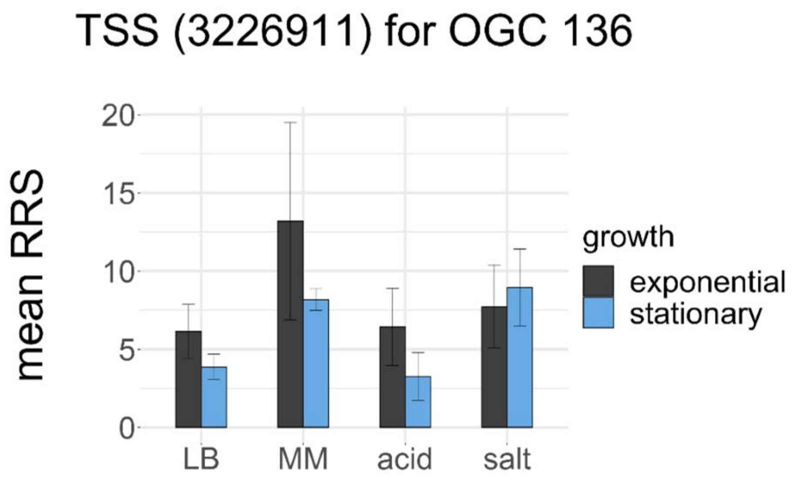
TSS (3226911) for OGC 136
mean RRS
20
15
10
5
0
LB
MM
acid
salt
growth
exponential
stationary

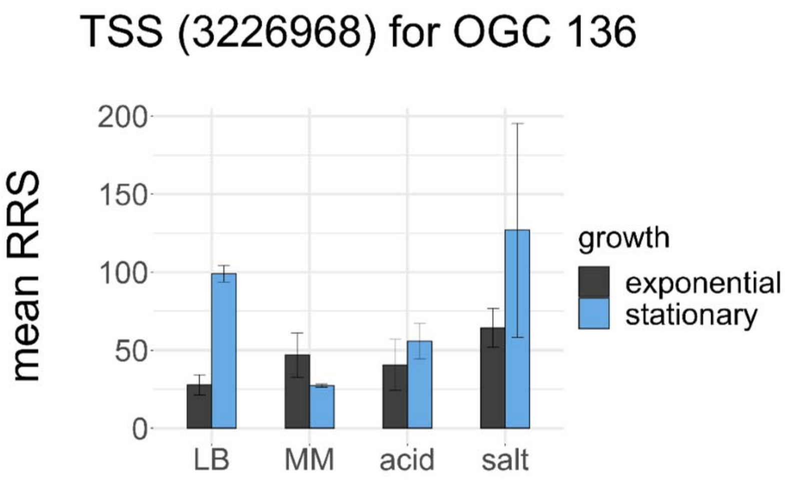
TSS (3226968) for OGC 136
mean RRS
200
150
100
50
0
LB
MM
acid
salt
growth
exponential
stationary

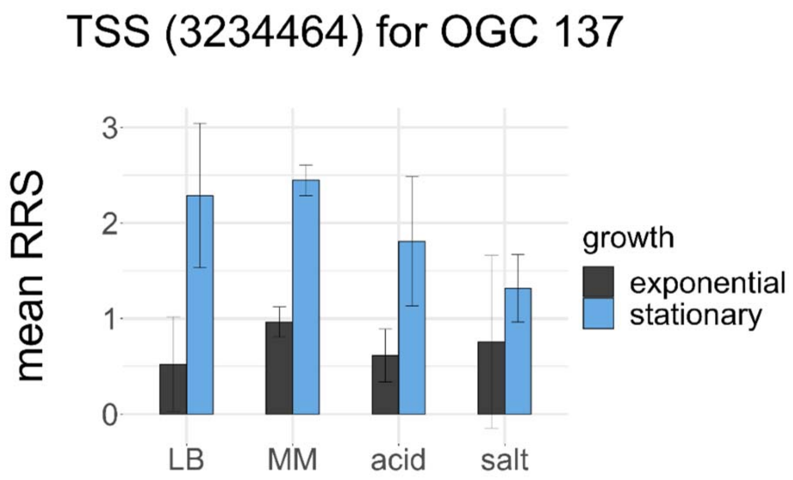
TSS (3234464) for OGC 137
mean RRS
3
2
1
0
LB
MM
acid
salt
growth
exponential
stationary

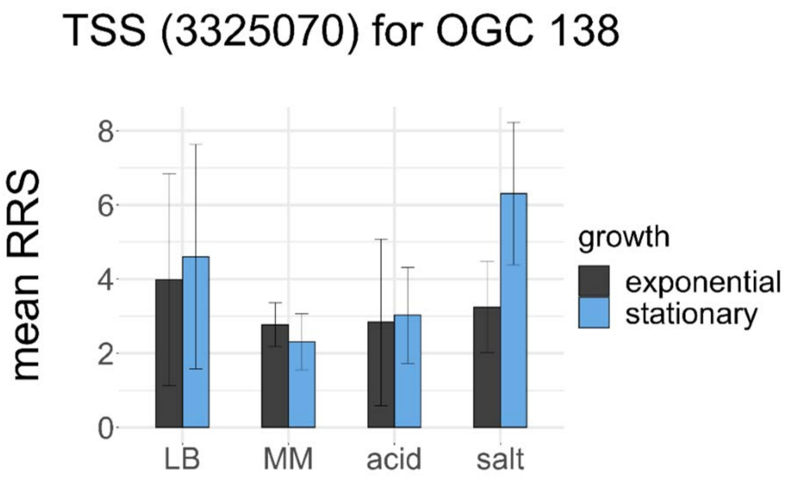
TSS (3325070) for OGC 138
mean RRS
0
2
4
6
8
LB
MM
acid
salt
growth
exponential
stationary

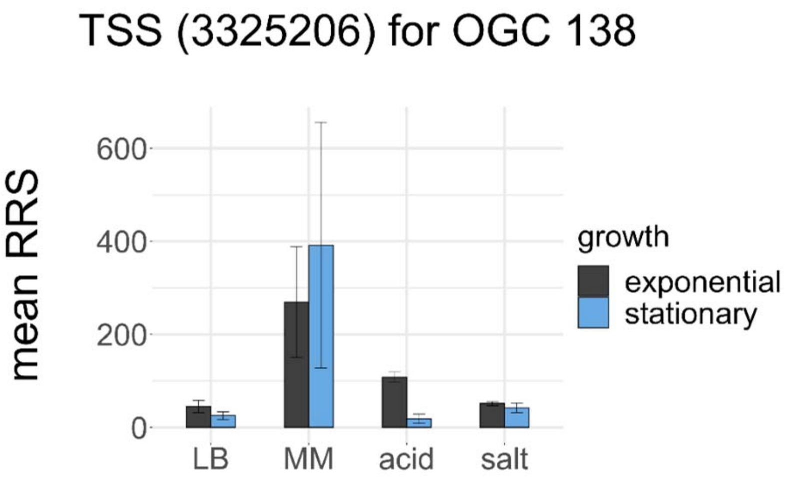
TSS (3325206) for OGC 138
mean RRS
0
200
400
600
LB
MM
acid
salt
growth
exponential
stationary

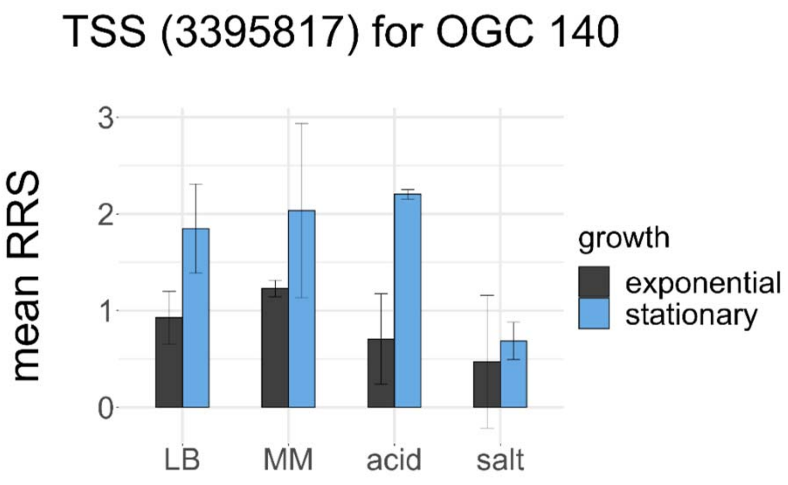
TSS (3395817) for OGC 140
mean RRS
0
1
2
3
LB
MM
acid
salt
growth
exponential
stationary

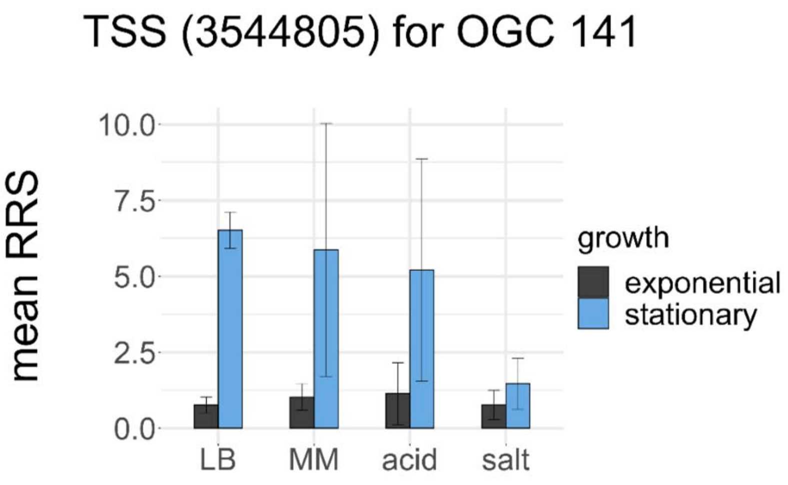
TSS (3544805) for OGC 141
mean RRS
0.0
2.5
5.0
7.5
10.0
LB
MM
acid
salt
growth
exponential
stationary

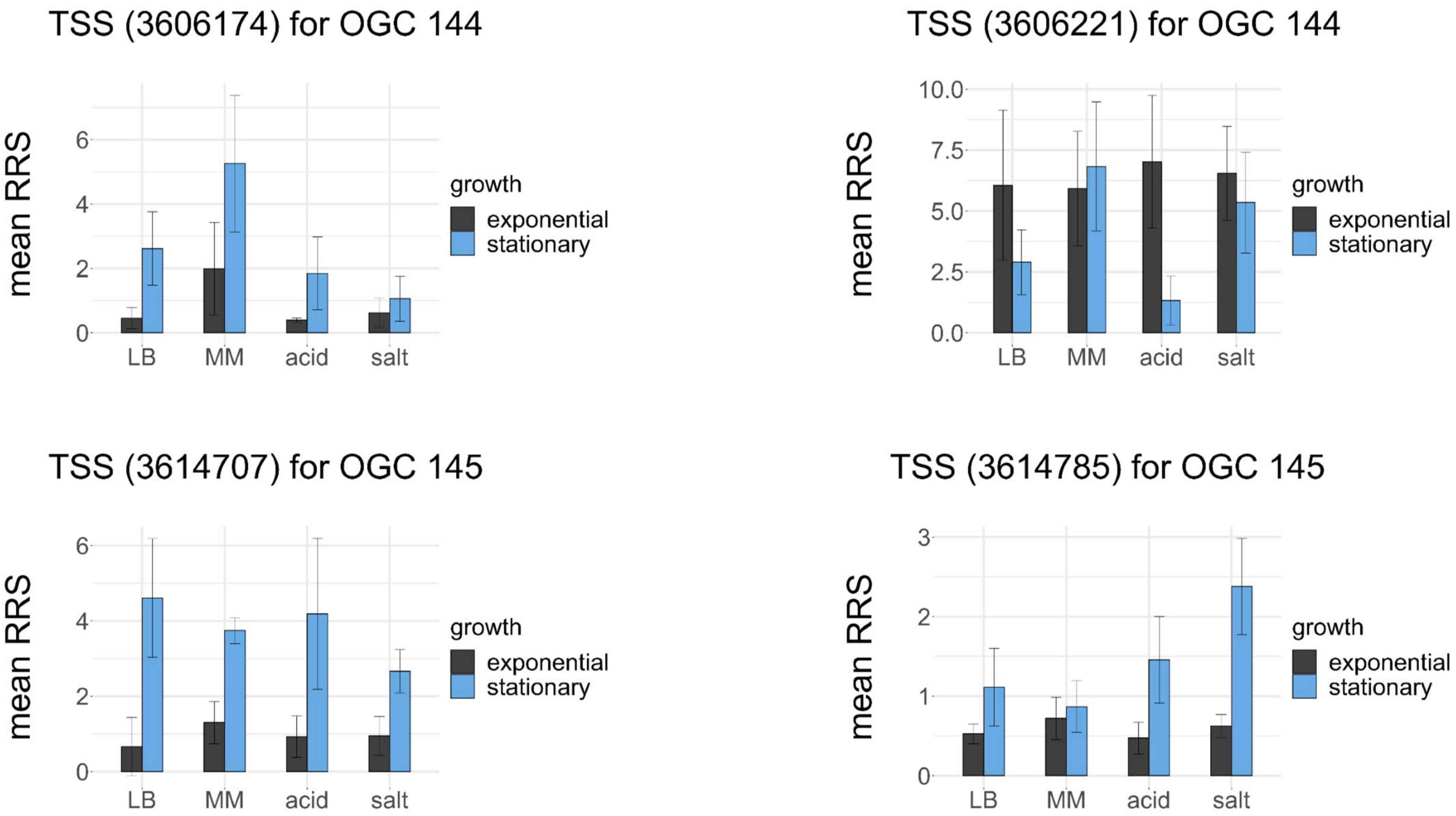
TSS (3606174) for OGC 144
mean RRS
0
2
4
6
LB
MM
acid
salt
growth
exponential
stationary
TSS (3606221) for OGC 144
mean RRS
0.0
2.5
5.0
7.5
10.0
LB
MM
acid
salt
growth
exponential
stationary
TSS (3614707) for OGC 145
mean RRS
0
2
4
6
LB
MM
acid
salt
growth
exponential
stationary
TSS (3614785) for OGC 145
mean RRS
0
1
2
3
LB
MM
acid
salt
growth
exponential
stationary

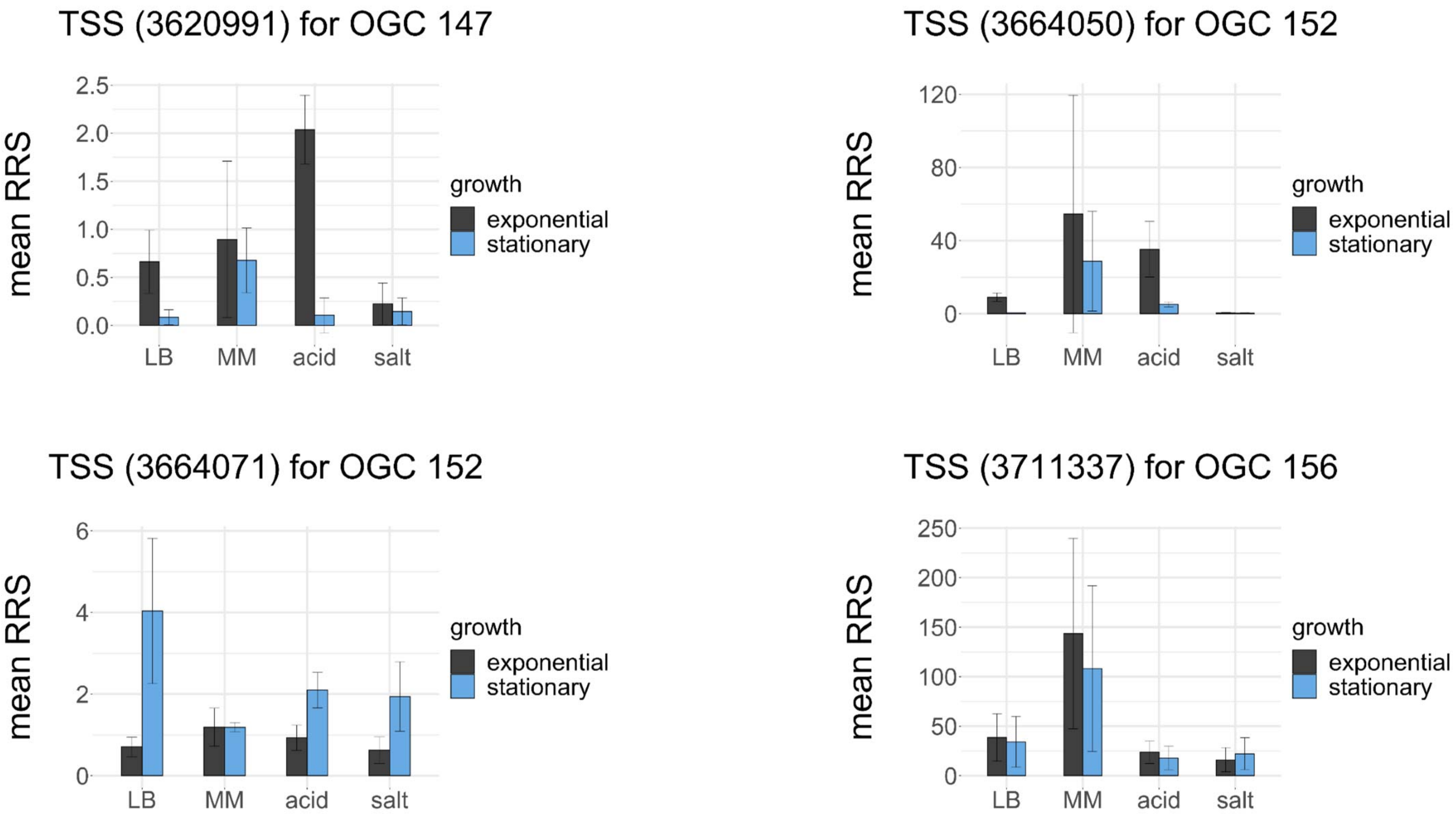

TSS (3620991) for OGC 147
mean RRS
2.5
2.0
1.5
1.0
0.5
0.0
LB
MM
acid
salt
growth
exponential
stationary
TSS (3664050) for OGC 152
mean RRS
120
80
40
0
LB
MM
acid
salt
growth
exponential
stationary
TSS (3664071) for OGC 152
mean RRS
6
4
2
0
LB
MM
acid
salt
growth
exponential
stationary
TSS (3711337) for OGC 156
mean RRS
250
200
150
100
50
0
LB
MM
acid
salt
growth
exponential
stationary

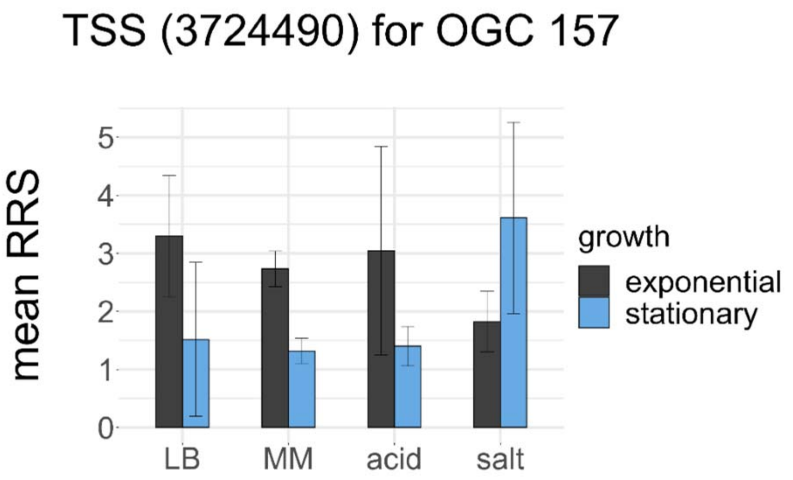
TSS (3724490) for OGC 157
mean RRS
0
1
2
3
4
5
LB
MM
acid
salt
growth
exponential
stationary

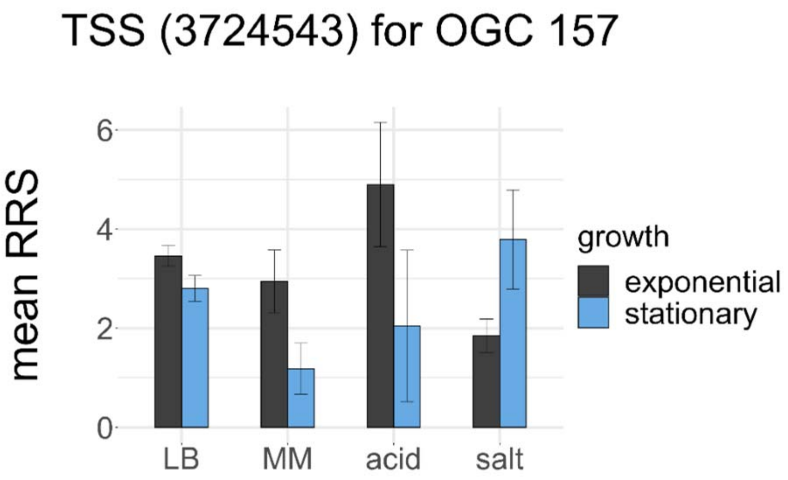
TSS (3724543) for OGC 157
mean RRS
0
2
4
6
LB
MM
acid
salt
growth
exponential
stationary

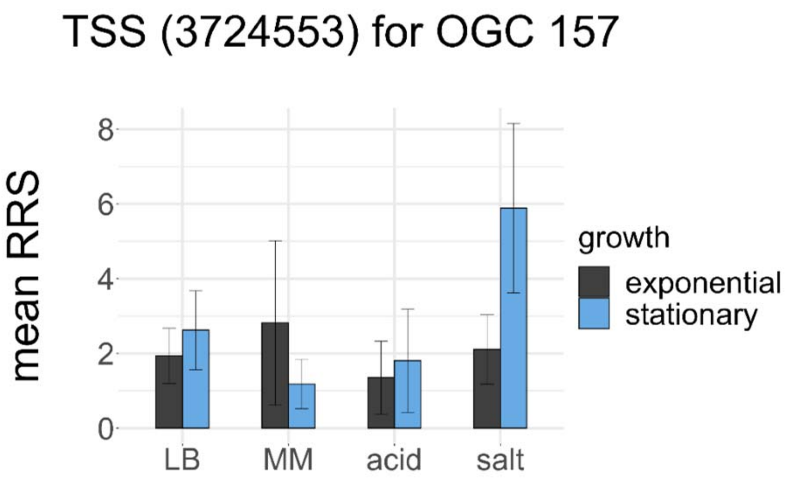
TSS (3724553) for OGC 157
mean RRS
0
2
4
6
8
LB
MM
acid
salt
growth
exponential
stationary

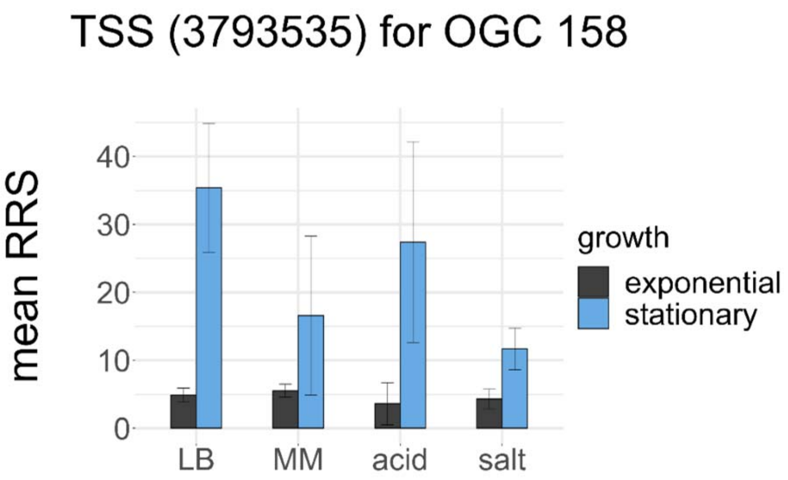
TSS (3793535) for OGC 158
mean RRS
0
10
20
30
40
LB
MM
acid
salt
growth
exponential
stationary

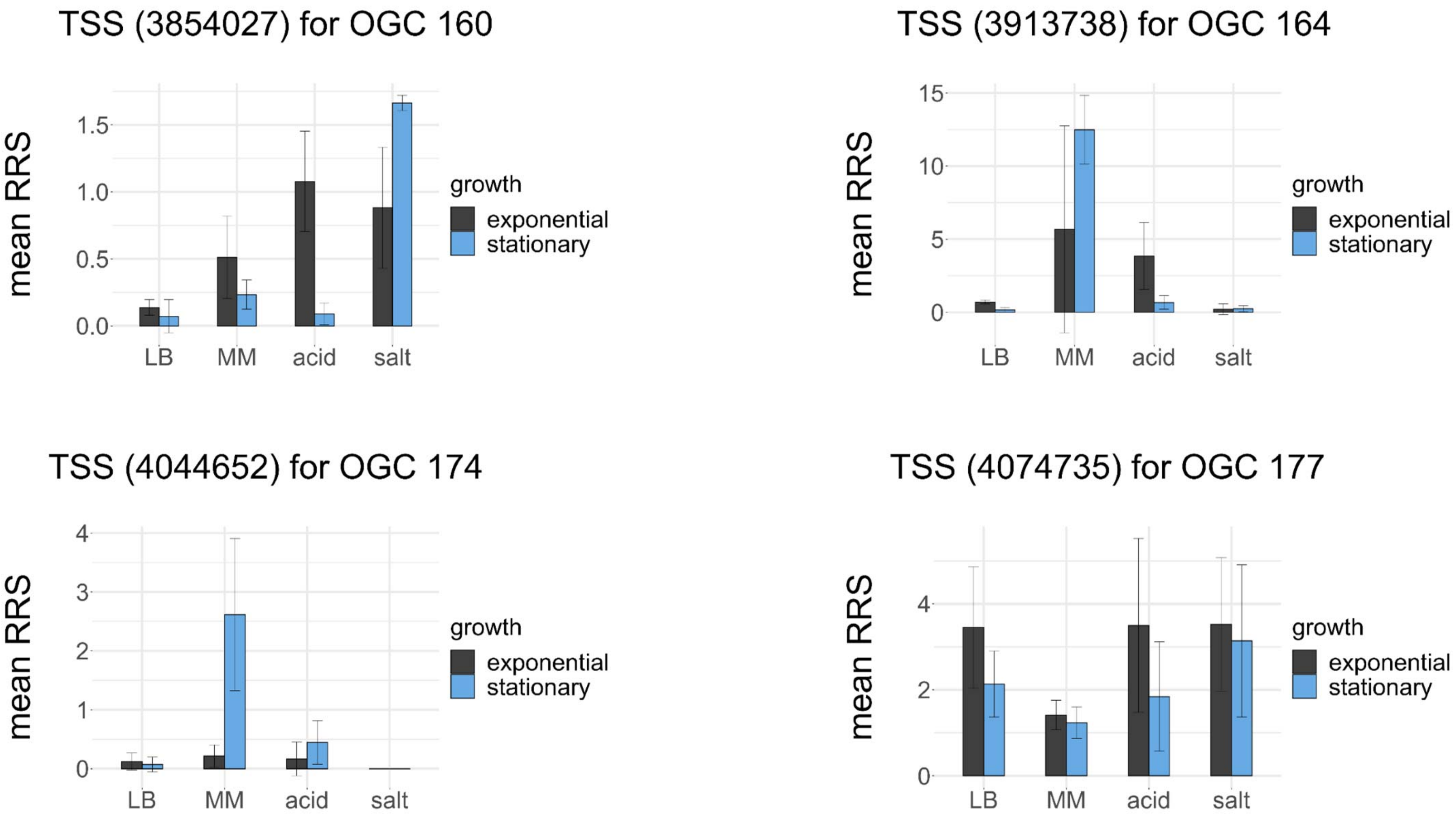
TSS (3854027) for OGC 160
mean RRS
0.0
0.5
1.0
1.5
LB
MM
acid
salt
growth
exponential
stationary
TSS (3913738) for OGC 164
mean RRS
0
5
10
15
LB
MM
acid
salt
growth
exponential
stationary
TSS (4044652) for OGC 174
mean RRS
0
1
2
3
4
LB
MM
acid
salt
growth
exponential
stationary
TSS (4074735) for OGC 177
mean RRS
0
2
4
LB
MM
acid
salt
growth
exponential
stationary

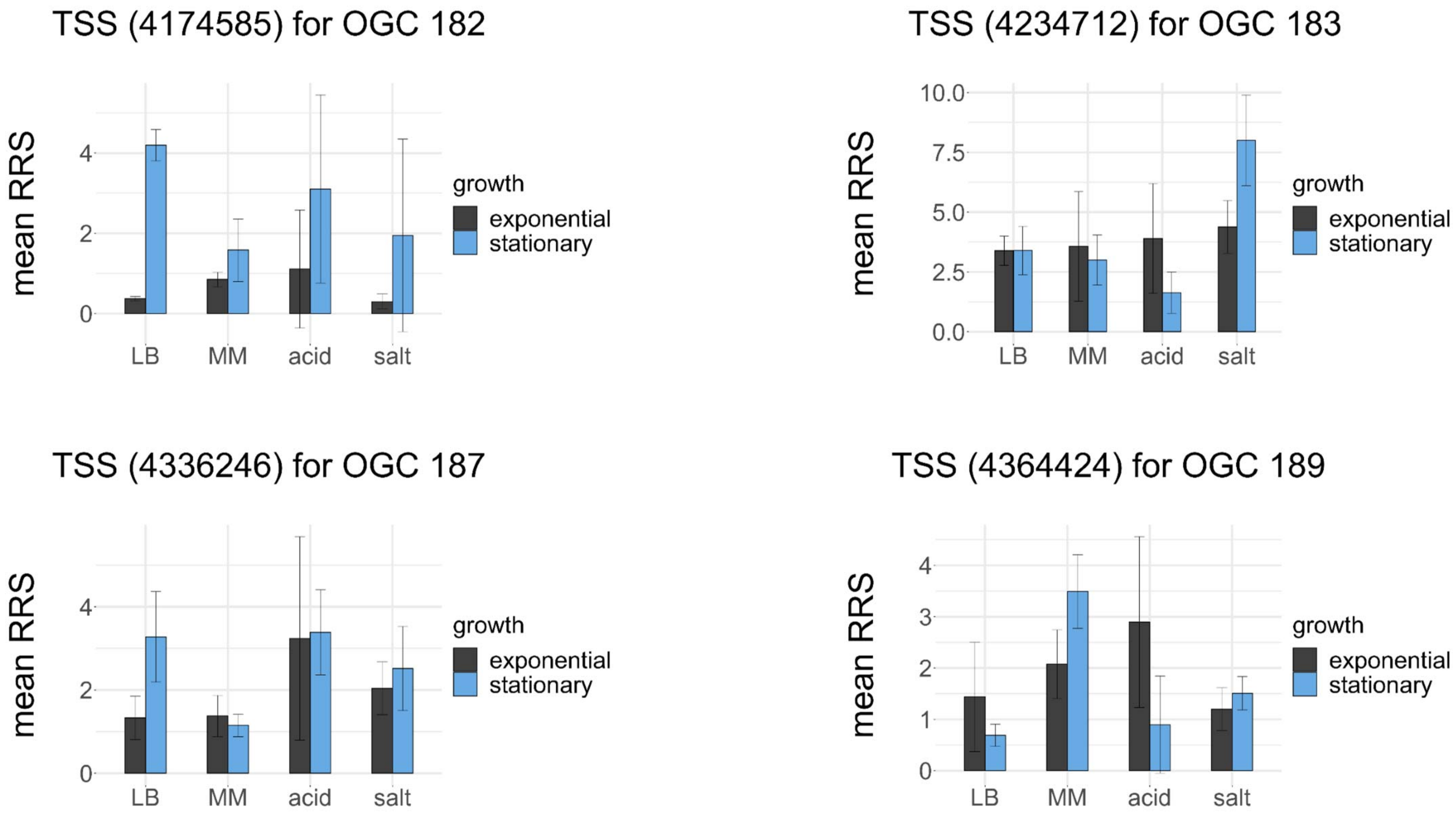
TSS (4174585) for OGC 182
mean RRS
0
2
4
LB
MM
acid
salt
growth
exponential
stationary
TSS (4234712) for OGC 183
mean RRS
0.0
2.5
5.0
7.5
10.0
LB
MM
acid
salt
growth
exponential
stationary
TSS (4336246) for OGC 187
mean RRS
0
2
4
LB
MM
acid
salt
growth
exponential
stationary
TSS (4364424) for OGC 189
mean RRS
0
1
2
3
4
LB
MM
acid
salt
growth
exponential
stationary

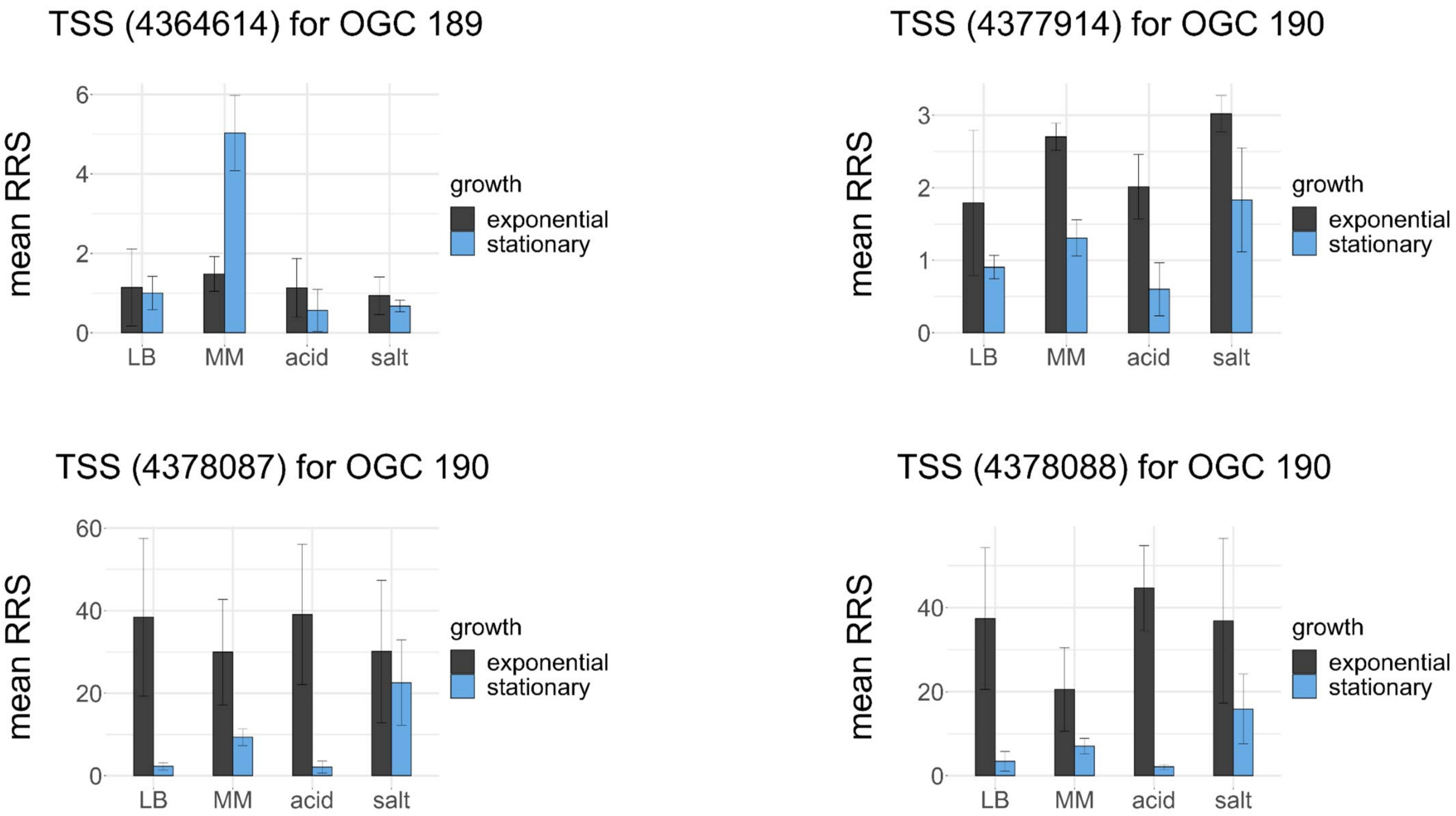
TSS (4364614) for OGC 189
TSS (4377914) for OGC 190
TSS (4378087) for OGC 190
TSS (4378088) for OGC 190
mean RRS
growth
exponential
stationary
LB
MM
acid
salt
0
2
4
6
1
3
20
40
60

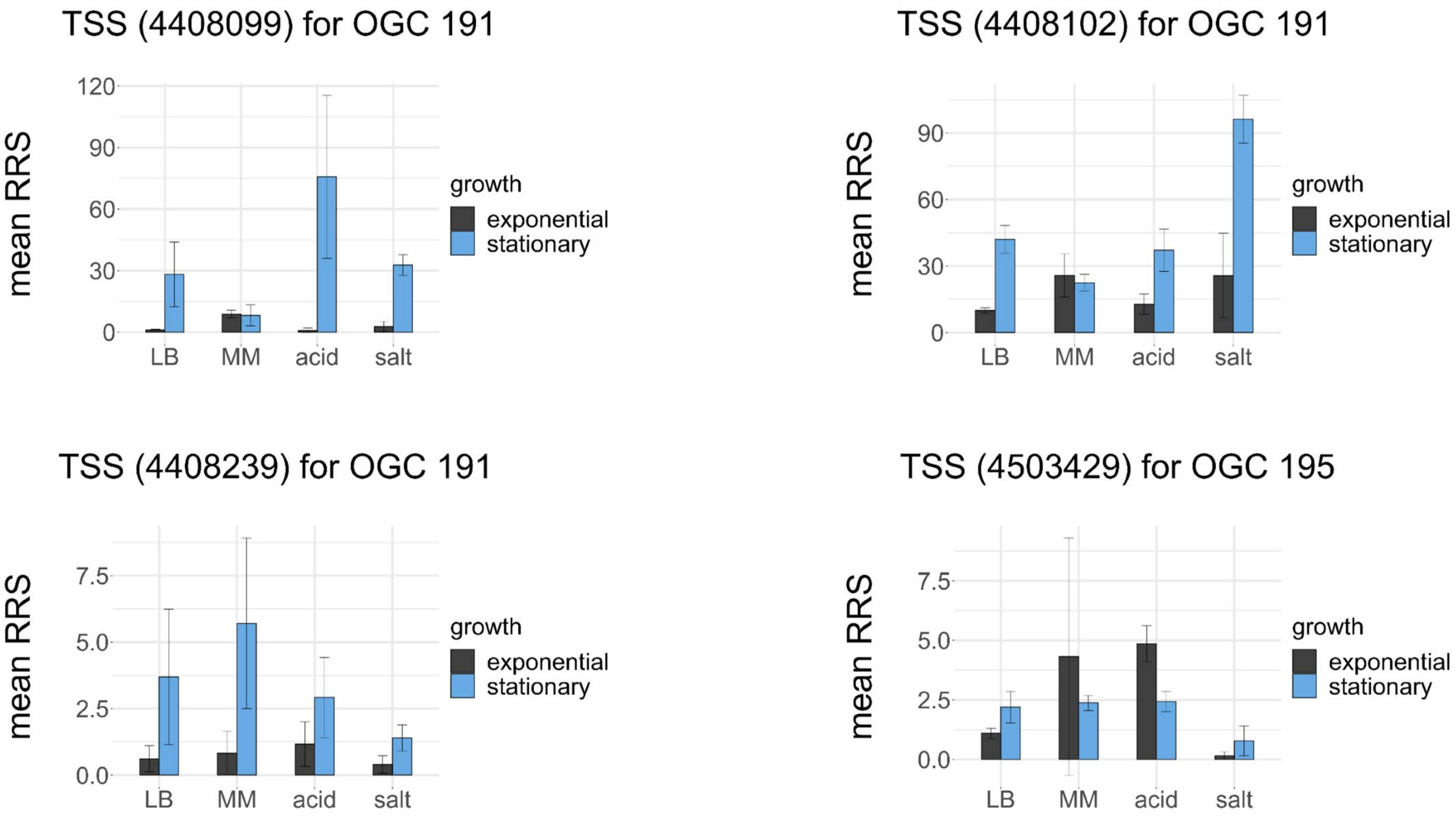
TSS (4408099) for OGC 191
mean RRS
120
90
60
30
0
LB
MM
acid
salt
growth
exponential
stationary
TSS (4408102) for OGC 191
mean RRS
90
60
30
0
LB
MM
acid
salt
growth
exponential
stationary
TSS (4408239) for OGC 191
mean RRS
7.5
5.0
2.5
0.0
LB
MM
acid
salt
growth
exponential
stationary
TSS (4503429) for OGC 195
mean RRS
7.5
5.0
2.5
0.0
LB
MM
acid
salt
growth
exponential
stationary

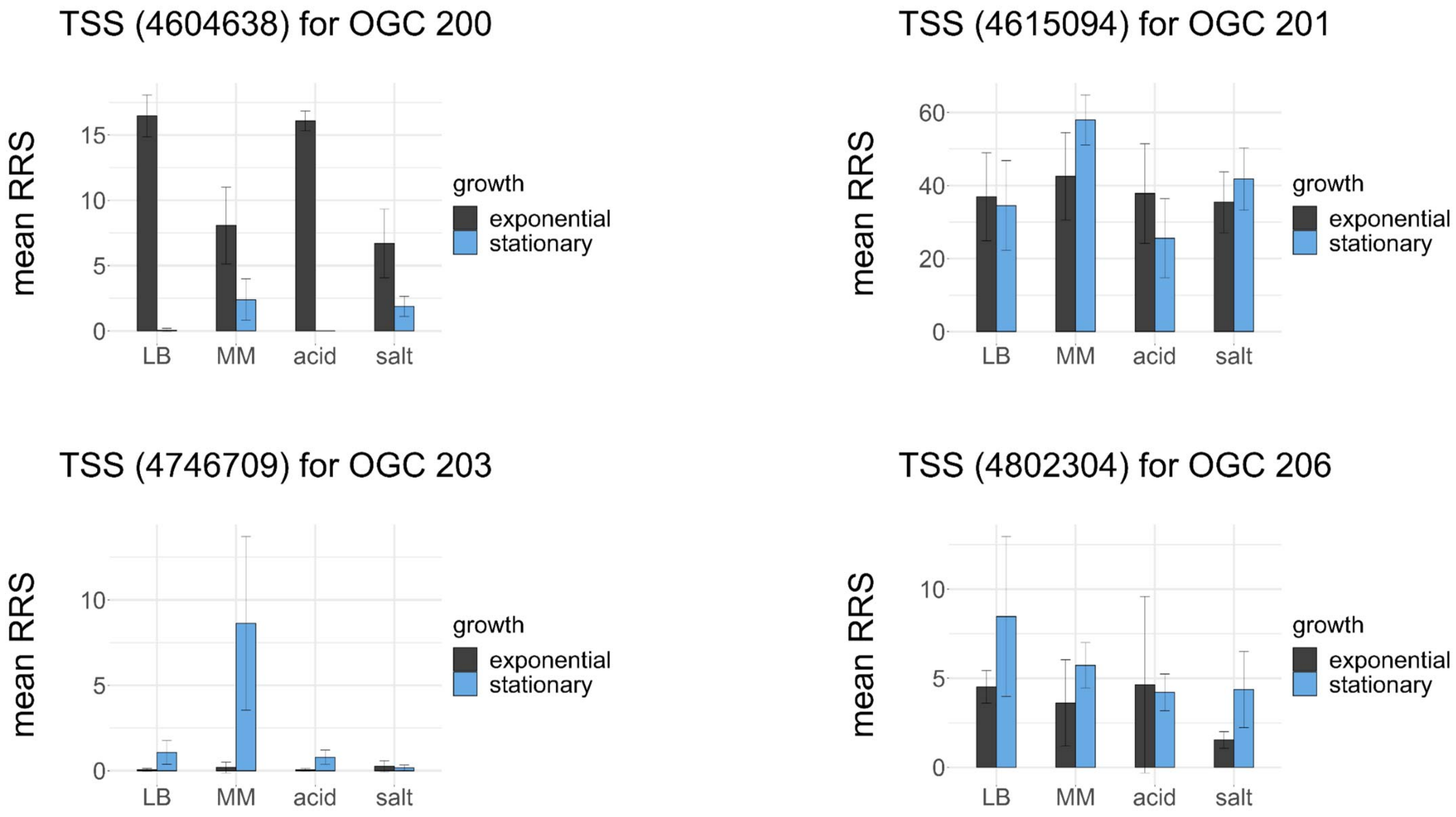

TSS (4604638) for OGC 200
TSS (4615094) for OGC 201
TSS (4746709) for OGC 203
TSS (4802304) for OGC 206
mean RRS
LB
MM
acid
salt
growth
exponential
stationary
0
5
10
15
20
40
60

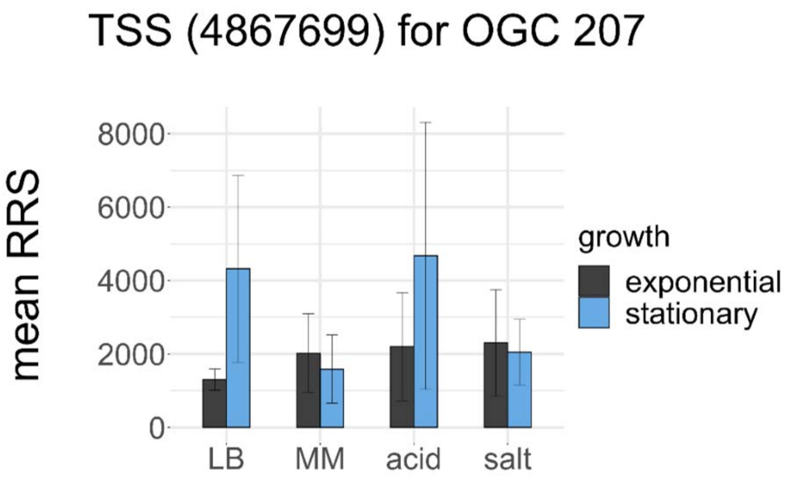
TSS (4867699) for OGC 207
mean RRS
8000
6000
4000
2000
0
LB
MM
acid
salt
growth
exponential
stationary

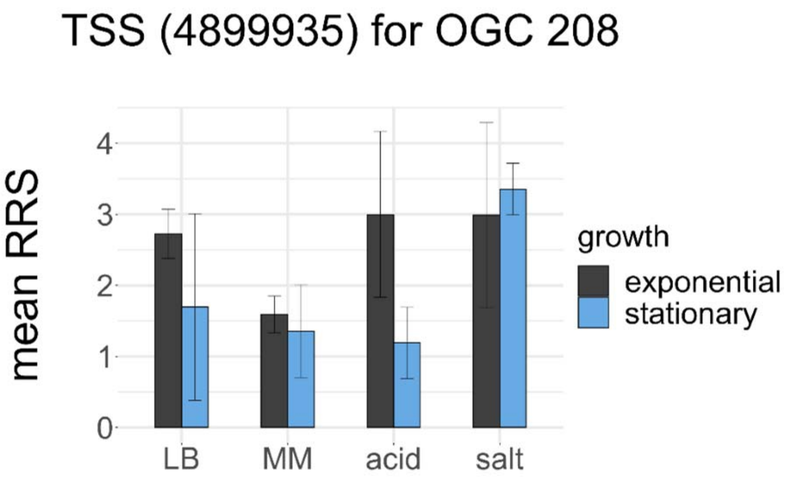
TSS (4899935) for OGC 208
mean RRS
4
3
2
1
0
LB
MM
acid
salt
growth
exponential
stationary

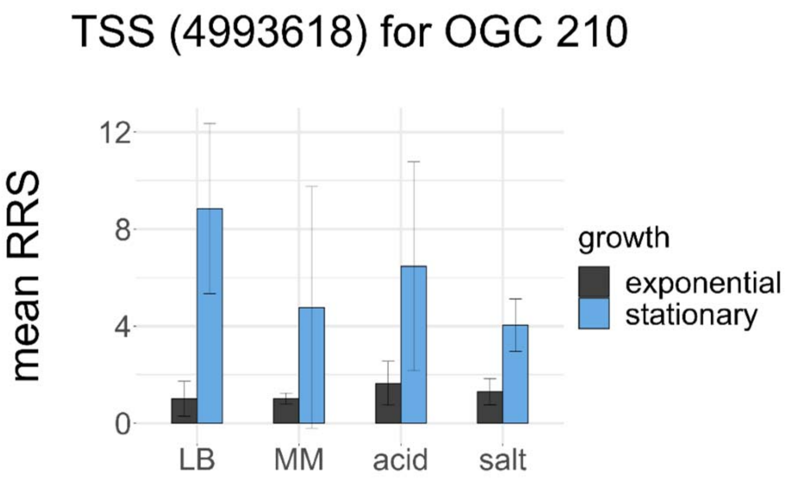
TSS (4993618) for OGC 210
mean RRS
12
8
4
0
LB
MM
acid
salt
growth
exponential
stationary

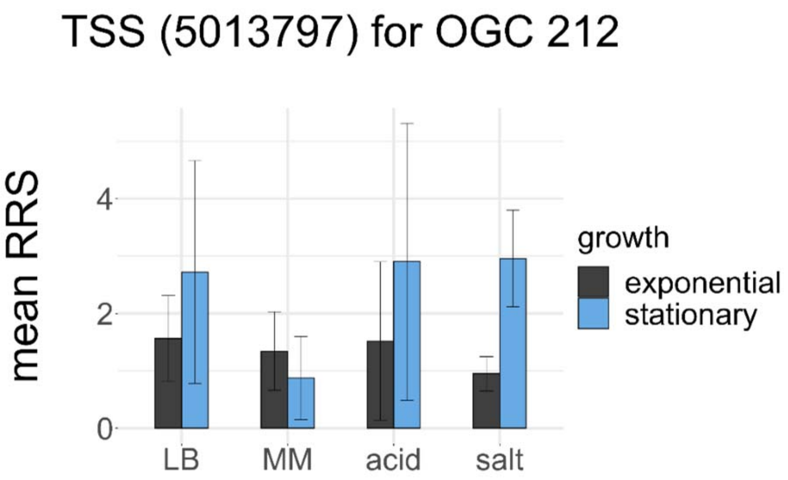
TSS (5013797) for OGC 212
mean RRS
4
2
0
LB
MM
acid
salt
growth
exponential
stationary

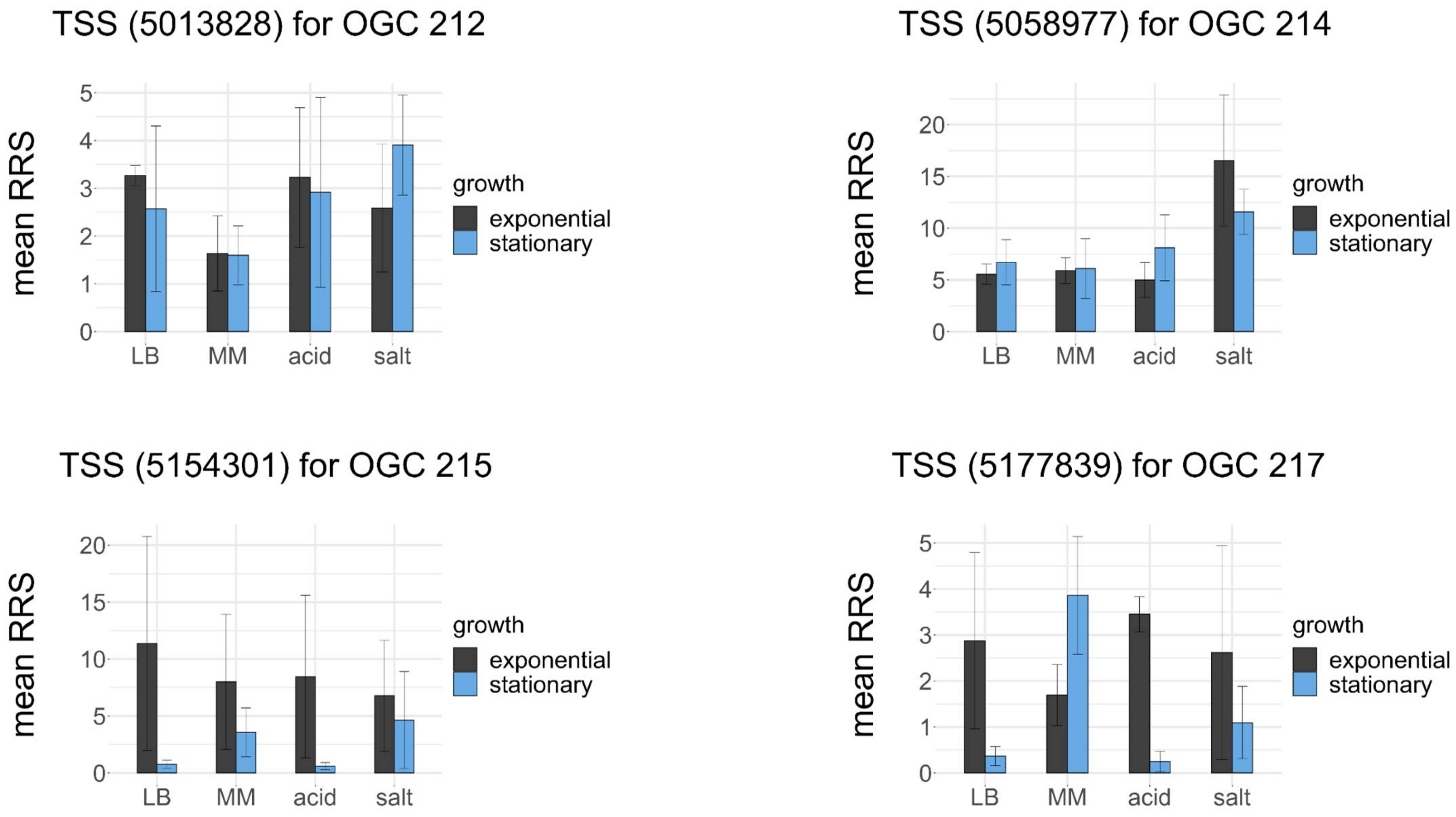
TSS (5013828) for OGC 212
TSS (5058977) for OGC 214
TSS (5154301) for OGC 215
TSS (5177839) for OGC 217
mean RRS
growth
exponential
stationary
LB
MM
acid
salt

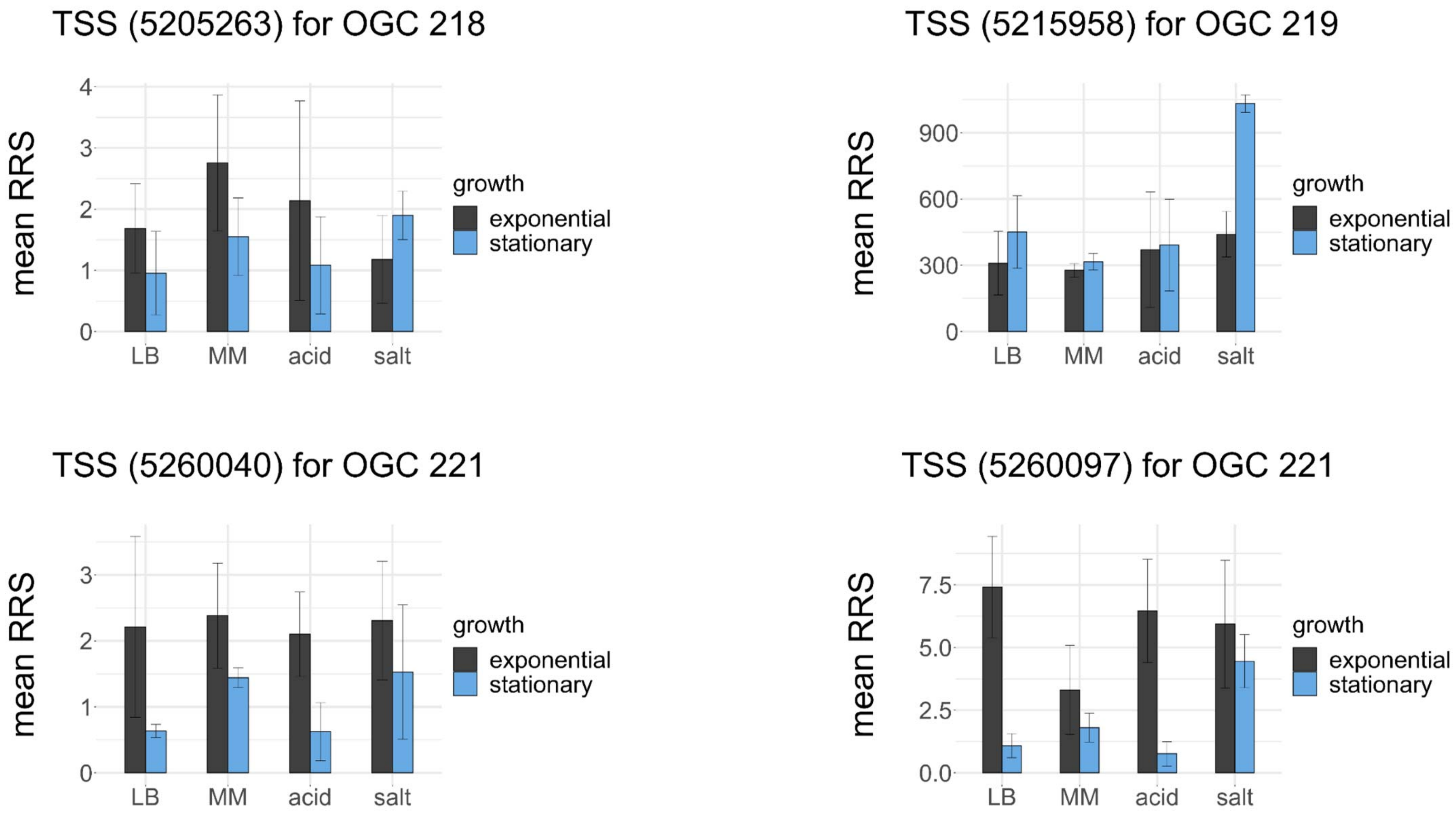
TSS (5205263) for OGC 218
TSS (5215958) for OGC 219
TSS (5260040) for OGC 221
TSS (5260097) for OGC 221
mean RRS
growth
exponential
stationary
LB
MM
acid
salt
0
1
2
3
4
300
600
900
0.0
2.5
5.0
7.5

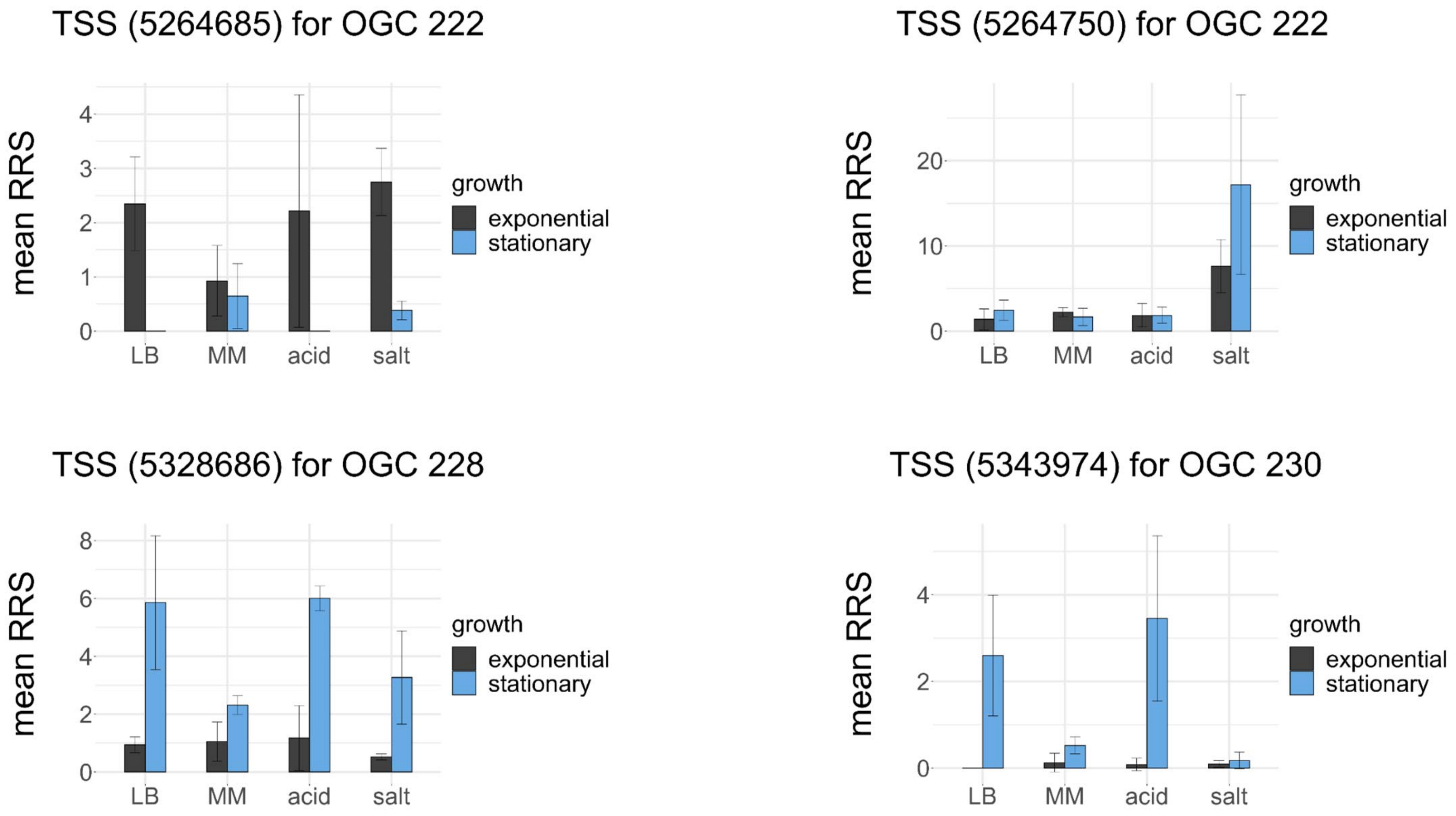
TSS (5264685) for OGC 222
TSS (5264750) for OGC 222
TSS (5328686) for OGC 228
TSS (5343974) for OGC 230
mean RRS
growth
exponential
stationary
LB
MM
acid
salt

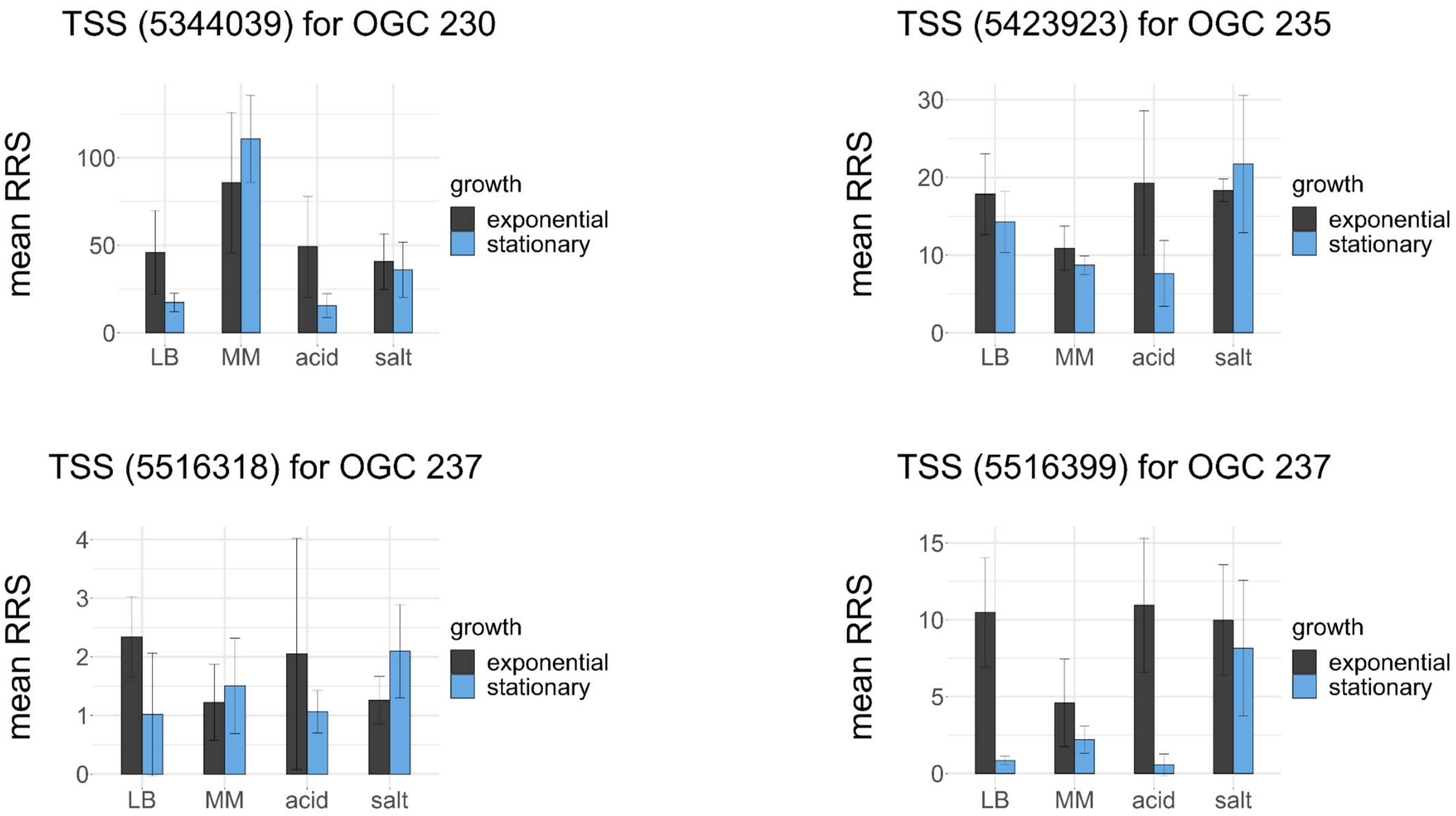
TSS (5344039) for OGC 230
mean RRS
100
50
0
LB
MM
acid
salt
growth
exponential
stationary
TSS (5423923) for OGC 235
mean RRS
30
20
10
0
LB
MM
acid
salt
growth
exponential
stationary
TSS (5516318) for OGC 237
mean RRS
4
3
2
1
0
LB
MM
acid
salt
growth
exponential
stationary
TSS (5516399) for OGC 237
mean RRS
15
10
5
0
LB
MM
acid
salt
growth
exponential
stationary

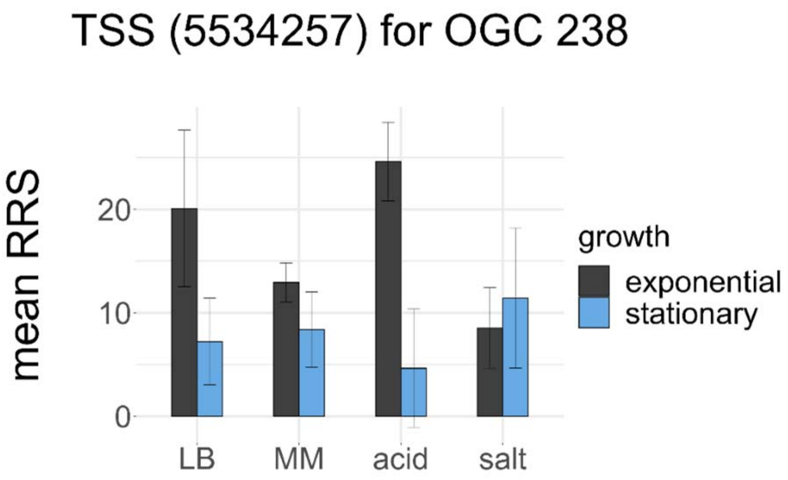

TSS (5534257) for OGC 238
mean RRS
0
10
20
LB
MM
acid
salt
growth
exponential
stationary


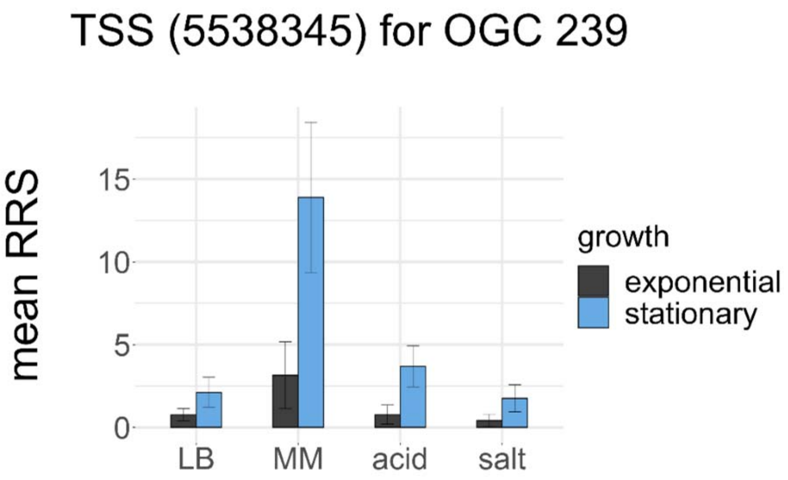

TSS (5538345) for OGC 239
mean RRS
0
5
10
15
LB
MM
acid
salt
growth
exponential
stationary


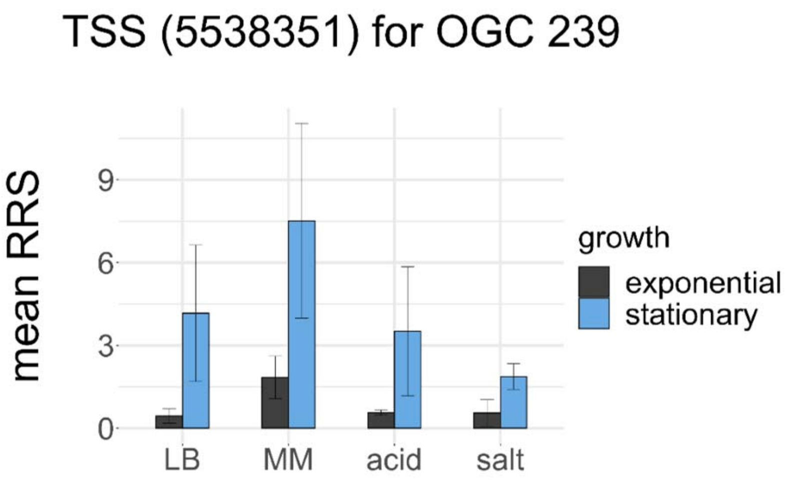

TSS (5538351) for OGC 239
mean RRS
0
3
6
9
LB
MM
acid
salt
growth
exponential
stationary


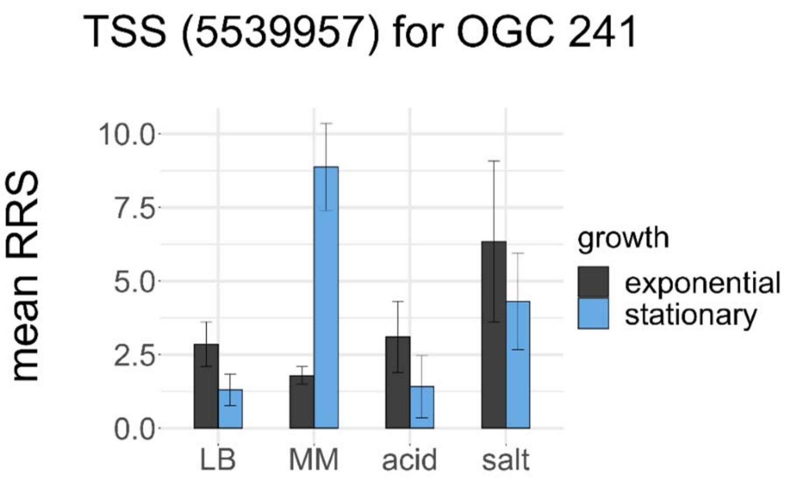

TSS (5539957) for OGC 241
mean RRS
0.0
2.5
5.0
7.5
10.0
LB
MM
acid
salt
growth
exponential
stationary
