## Supplementary Figure S3 for "Evidence for Numerous Embedded Antisense Overlapping Genes in Diverse *E. coli* Strains"

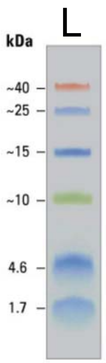


**Uncut Western blots of OGCs**, C-terminally fused to an SPA-tag (~8 kDa).

G: positive control (glutathione-S-transferase, 31 kDa incl. SPA-tag)
C: empty cells
T: empty vector
L: marker lane (see panel to the right for sizes)
Numbers correspond the OGC tested.
* indicates the expected protein size.
Some OGCs could not be cloned and are missing.

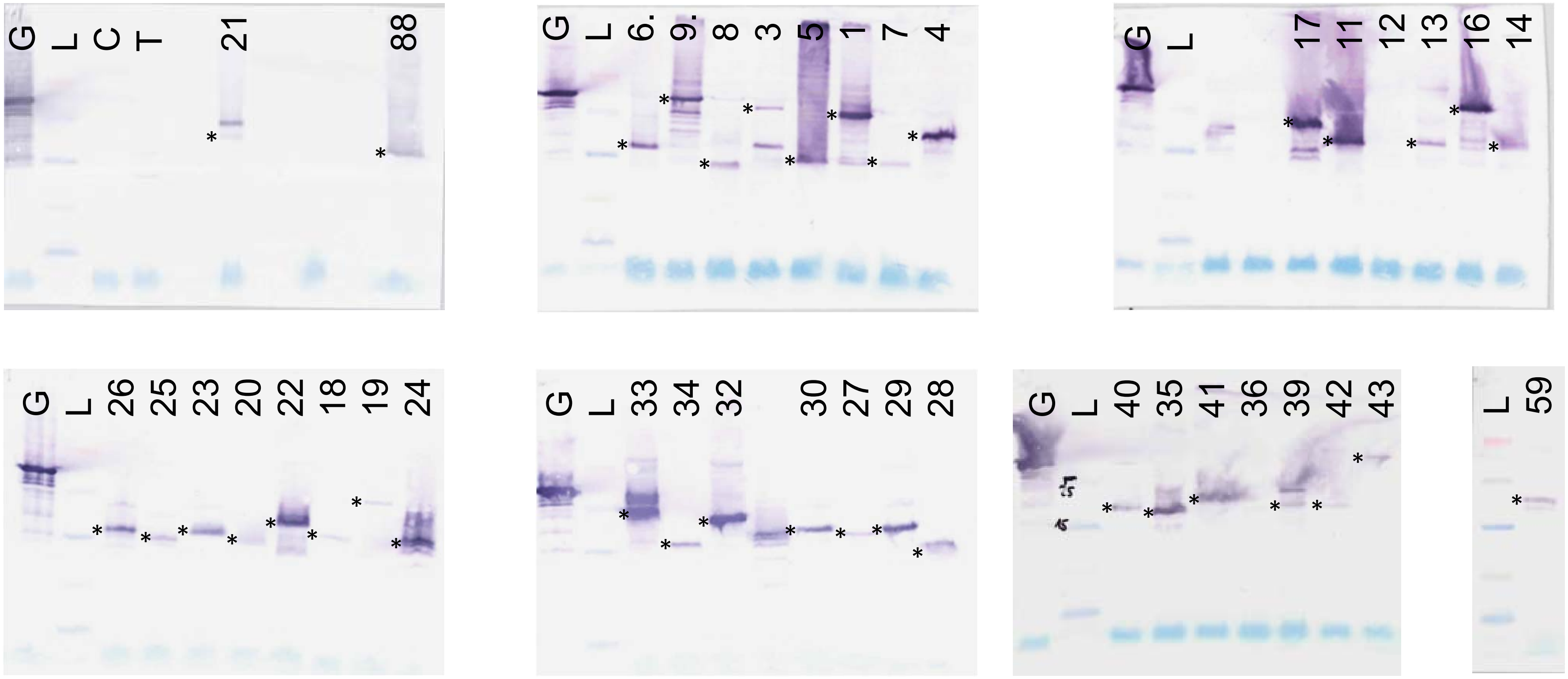

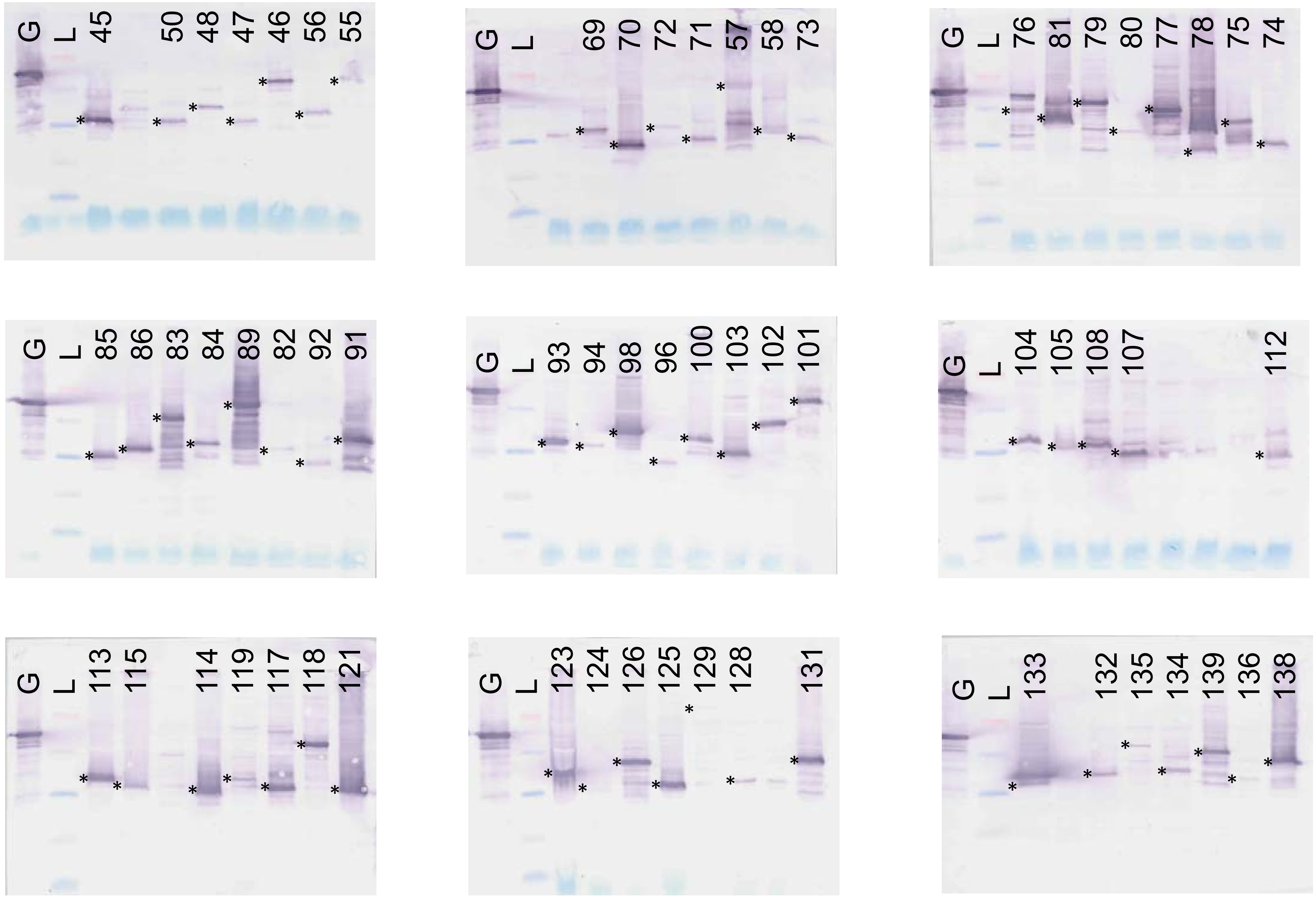
G L 45 50 48 47 46 56 55
G L 69 70 72 71 57 58 73
G L 76 81 79 80 77 78 75 74
G L 85 86 83 84 89 82 92 91
G L 93 94 98 96 100 103 102 101
G L 104 105 108 107 112
G L 113 115 114 119 117 118 121
G L 123 124 126 125 129 128 131
G L 133 132 135 134 139 136 138

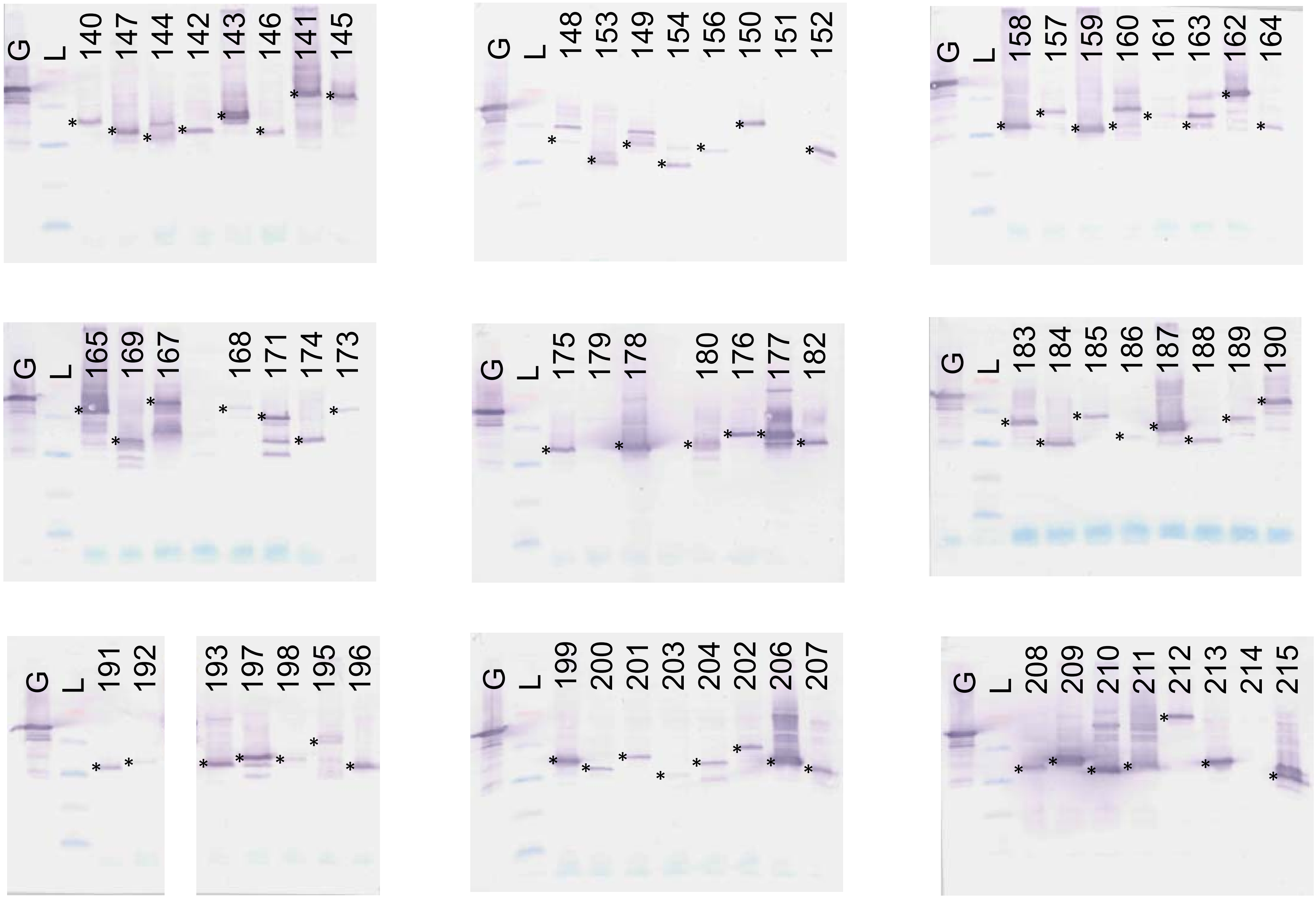

G L 140 147 144 142 143 146 141 145
G L 148 153 149 154 156 150 151 152
G L 158 157 159 160 161 163 162 164
G L 165 169 167 168 171 174 173
G L 175 179 178 180 176 177 182
G L 183 184 185 186 187 188 189 190
G L 191 192
193 197 198 195 196
G L 199 200 201 203 204 202 206 207
G L 208 209 210 211 212 213 214 215

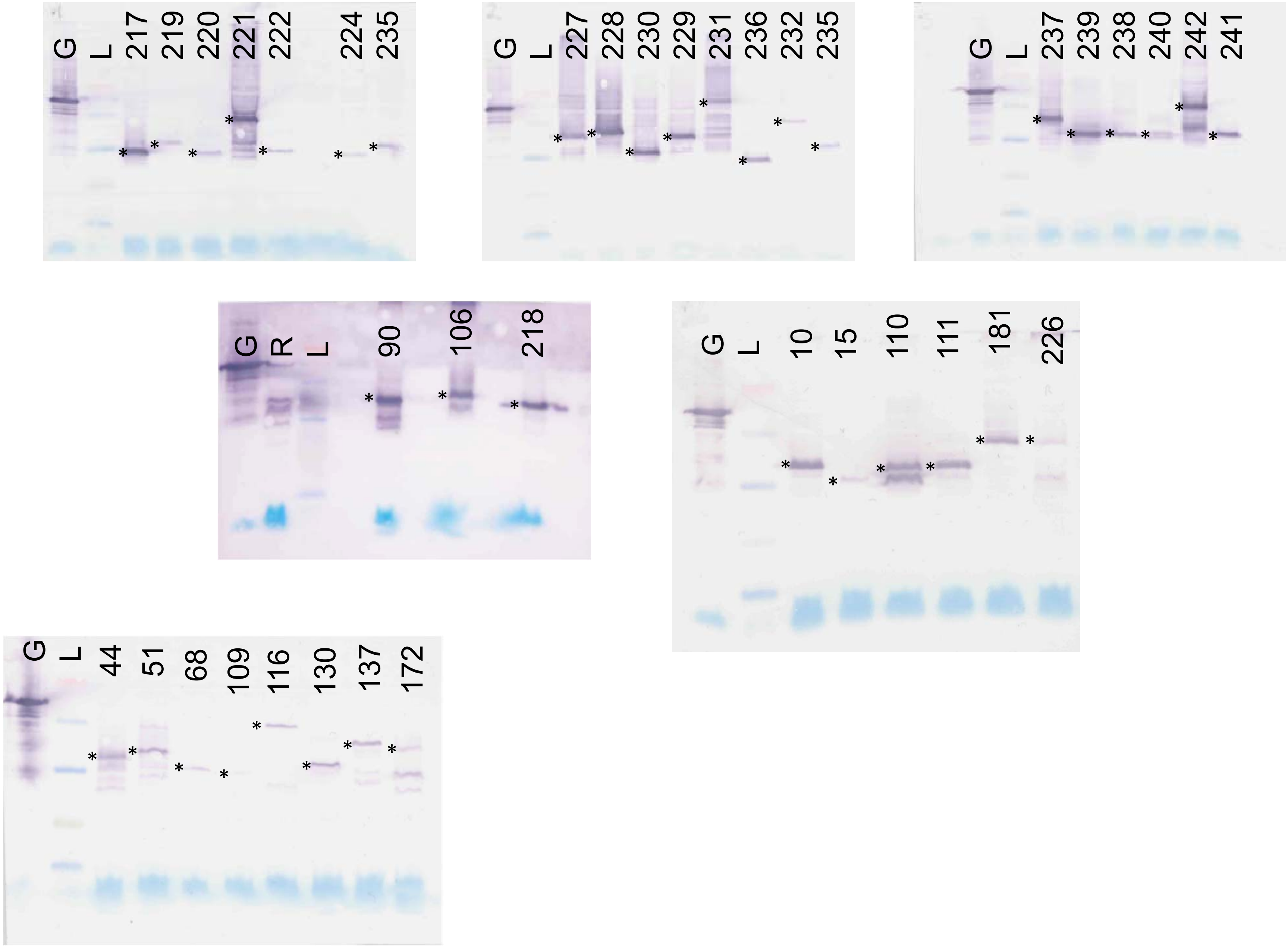

G L 217 219 220 221 222 224 235
G L 227 228 230 229 231 236 232 235
G L 237 239 238 240 242 241
G R L 90 106 218
G L 10 15 110 111 181 226
G L 44 51 68 109 116 130 137 172
