## Supplementary Figure S4 for "Evidence for Numerous Embedded Antisense Overlapping Genes in Diverse *E. coli* Strains"

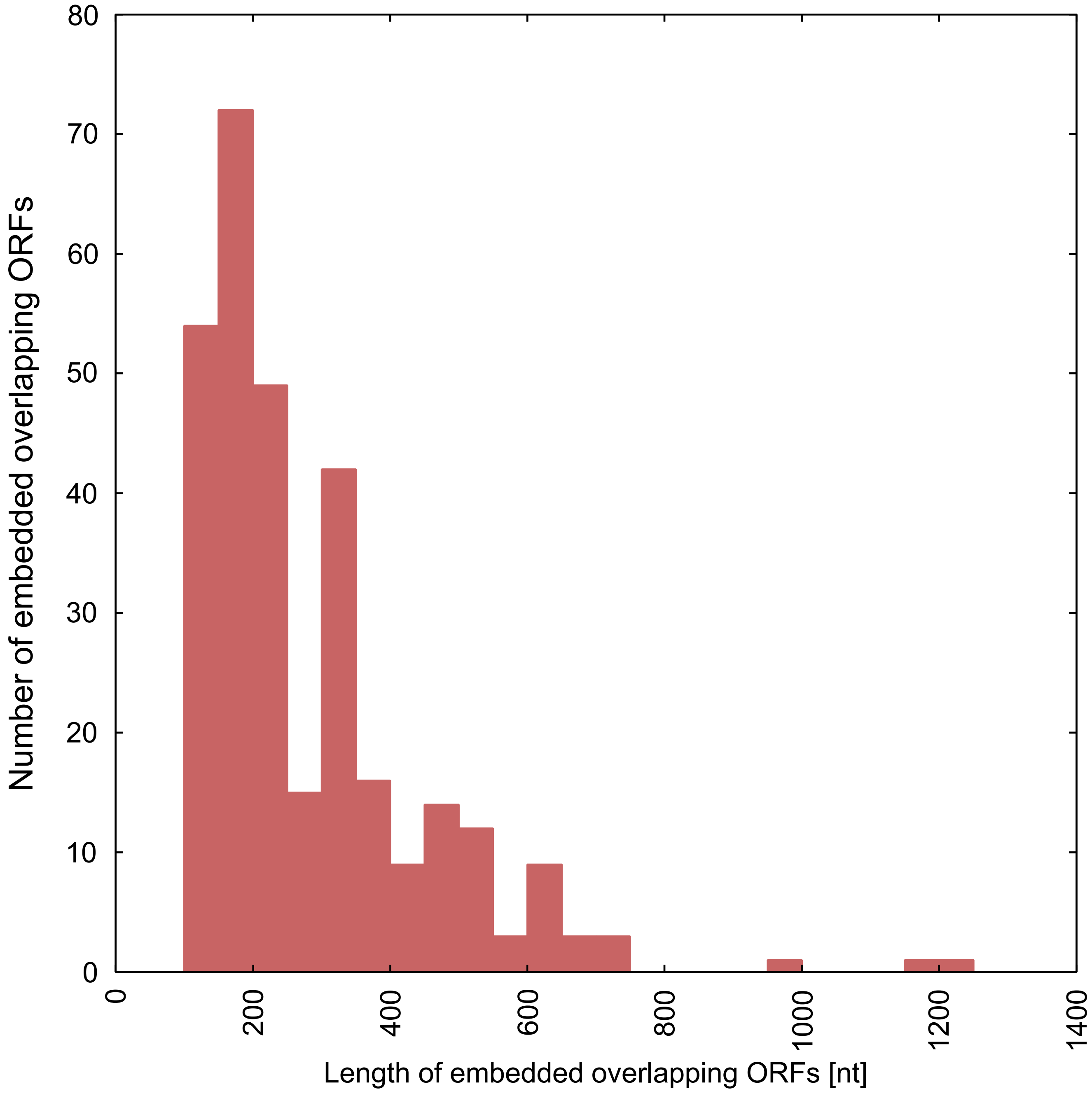


**Supplementary Figure S4. Length distributions of overlapping ORFs**

Length distribution of embedded overlapping ORFs in five *E. coli* strains (see Table 1), which have homologous protein sequences in other organisms (e-value ≤ $10^{-10}$) according to a BLASTp analysis of the RefSeq database.
