## Supplementary Figure S5 for "Evidence for Numerous Embedded Antisense Overlapping Genes in Diverse *E. coli* Strains"

**N-terminal peptide fragments of overlapping genes in *E. coli* MG1655**. N-terminal peptides for (A) embedded ORFs in sense and (B) embedded ORFs in antisense. Data are from Ndah, et al.[16].

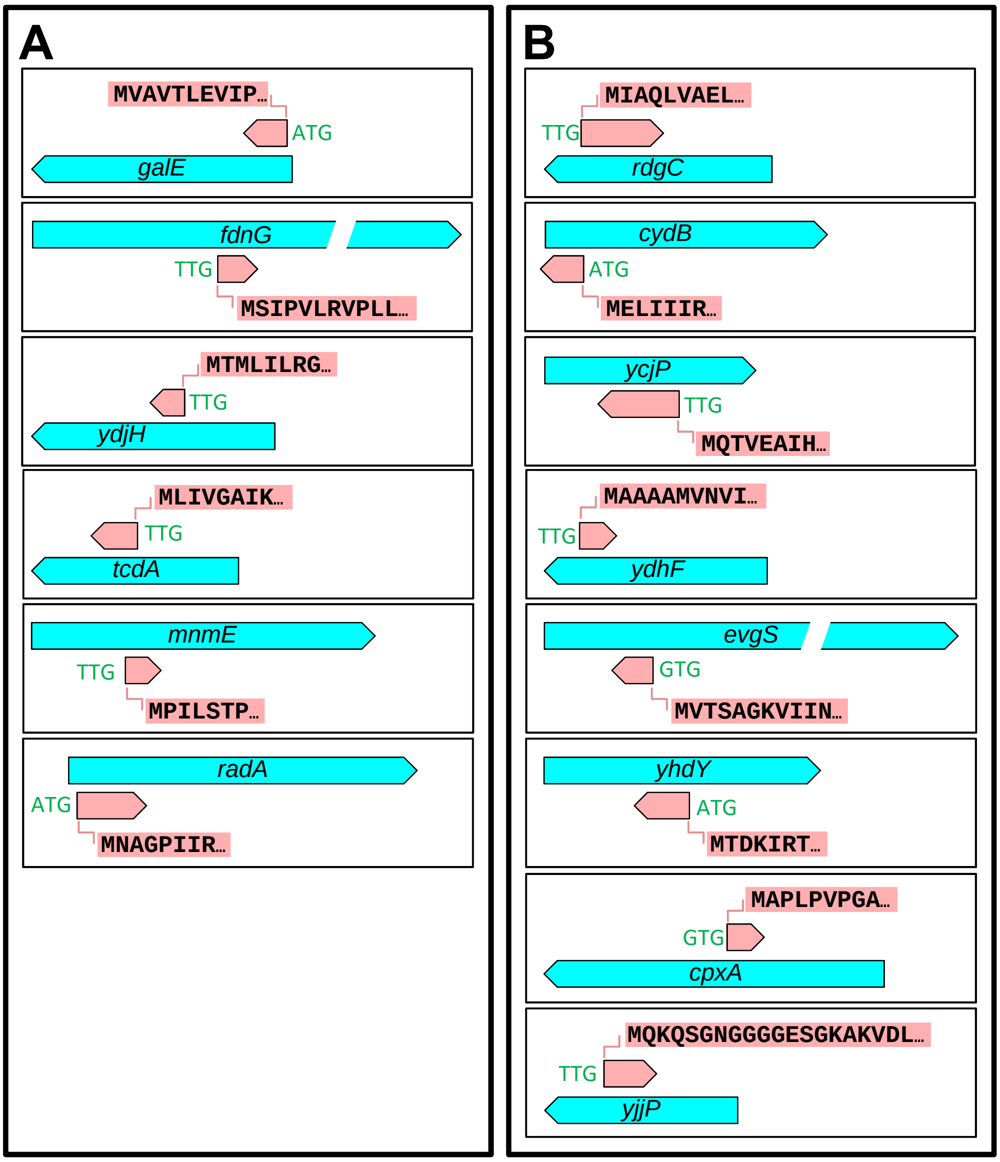
